# SMYD3 protein degraders in HPV-negative head and neck squamous cell carcinoma

**DOI:** 10.64898/2026.09.28.754983

**Authors:** Jawad Akhtar, Sai Reddy Doda, William J. Moore, Mohd Saleem Dar, Shafagh Valipour, Sohyoung Kim, Jianghong Wu, Yong Wan, Yuan Wang, Meiye Jiang, Arfa Moshiri, John Schneekloth, Sudipto Das, Thorkell Andresson, Rolf E. Swenson, Vassiliki Saloura

## Abstract

Despite the advent of immunotherapy in the treatment of human-papilloma-virus (HPV)-negative head and neck squamous cell carcinoma, the overall prognosis still remains dismal. Novel therapeutic approaches are thus urgently needed. We recently reported that SET and MYND containing 3 (SMYD3), a protein methyltransferase that is known to activate the transcription of its target genes through H3K4me3, is overexpressed in ∼60% of HPV-negative HNSCC tumors, it drives proliferation and invasion, and induces resistance to immunotherapy in HPV-negative HNSCC. These phenotypes are mediated partially through transcriptional regulation of cell cycle, EMT and immune-related target genes by SMYD3. Despite its enzymatic inhibitory potency in HPV-negative HNSC cell lines, the commercially available SMYD3 inhibitors EPZ031686 and BAY-6035 demonstrated no in vitro phenotypic efficacy in these cell lines, supporting that SMYD3 exerts its oncogenic functions predominantly through catalytically-independent mechanisms. Thus, to effectively target SMYD3, we designed a series of SMYD3 proteolysis targeting chimeras (PROTACs). Treatment of HPV-negative HNSCC cell lines with SMYD3 PROTACs at nanomolar concentrations degraded SMYD3 effectively and induced a drastic decrease in the proliferative and invasive potential of HPV-negative HNSCC cells. One of the designed compounds, IAP-08, showed a promising DC_50_ at ∼177nM with Dmax of ∼90%, and effectively induced the formation of ternary complexes with SMYD3 and the XIAP E3 ligase. Furthermore, IAP-08 showed a favorable pharmacokinetic profile in vivo, underscoring the feasibility for further clinical development of this compound. Mass spectrometry analysis of IAP-08 supported its high specificity towards SMYD3. SMYD3 depletion by IAP-08 in HNSCC cells revealed transcriptional regulation of cell cycle, EMT- and type I IFN response genes, in contrast to SMYD3 inhibition which had only a weak transcriptional effect. In summary, we show that inhibition of the enzymatic activity of SMYD3 is not sufficient to repress its oncogenic activity and provide evidence that depletion through PROTACs effectively targets the oncogenic functions of SMYD3 in HPV-negative HNSCC.

## INTRODUCTION

Head and neck squamous cell carcinoma (HNSCC) is the sixth most prevalent non-skin cancer in the world, with approximately 73,000 new cases and 17,000 deaths in the United States in 2025 [1]. HNSCC is categorized into two main pathogenetic types: human-papilloma-virus (HPV)-positive and HPV-negative, which has strong associations with tobacco use. HPV-negative HNSCC is associated with a worse prognosis and treatment is often multimodal and toxic, with the use of surgical intervention, platinum-based chemotherapy and radiation therapy. Despite the recent advent of pembrolizumab, which blocks the programmed-death-1 (PD-1)/programmed death-ligand 1 (PD-L1) axis, as first-line immunotherapy in the treatment of HPV-negative HNSCC in both the curative-intent and the recurrent/metastatic setting [2, 3], response rates are low and the overall survival remains dismal. Novel therapeutic approaches are thus urgently needed.

SET and MYND Domain-containing 3 (SMYD3) is a protein lysine methyltransferase that has been identified as an oncogene and its overexpression is associated with poor survival in multiple cancer types, such as colon cancer, hepatocellular carcinoma (HCC), ovarian, breast, lung, pancreatic and HPV-negative HNSCC [4-23]. Importantly, its function is dispensable for normal mouse development and adult life, supporting its promise as an anti-cancer drug target [24]. SMYD3 binds to and activates the transcription of various oncogenes through the methylation of its histone substrate lysine 4 of histone H3 (H3K4) to trimethyl H3K4 (H3K4me3) [4-11]. SMYD3 has also been reported to promote the methylation of H4K20 (H4K20me3), a repressive histone mark [12-14]. Additionally, it also methylates non-histone substrates, such as MAP3K2, VEGFR1, AKT1 and HER2, which mediate its cytoplasmic functions [15, 17-20].

Regarding the oncogenic role of SMYD3 in HPV-negative HNSCC, our group recently reported that SMYD3 is overexpressed in ∼60% of HPV-negative HNSCC tumors, and it is associated with poor CD8+ T-cell intratumoral infiltration and resistance to pembrolizumab in HPV-negative HNSCC patients [21]. Albeit its established function as a transcriptional activator, we found that this is mediated through an orchestrated repression of multiple type I IFN response genes in HPV-negative HNSCC cancer cells through the transcriptional regulation of Ubiquitin-like with plant homeodomain and ring finger domains 1 (UHRF1), a reader of H3K9me3, and by non-canonically promoting the deposition of the repressive mark H4K20me3 on immune-related genes [21]. Accordingly, Smyd3 depletion increased the intratumoral influx of CD8+ T-cells and sensitized HPV-negative HNSCC flank mouse tumors to anti-PD-1 therapy in vivo [21, 22]. Aside from its immunomodulatory function, we showed that SMYD3 depletion also decreases the proliferative and invasive capacity of HPV-negative HNSCC cells through direct transcriptional regulation of specific cell cycle and epithelial-mesenchymal-transition (EMT) gene sets [23]. These phenotypes were mediated, at least partially, through direct canonical activation and non-canonical repression of specific gene sets by SMYD3, suggesting a “duality” of the transcriptional function of SMYD3 within the same cell context, a phenomenon that is largely unexplored.

Herein, we show for the first time that inhibition of the enzymatic activity of SMYD3 is not sufficient to repress its oncogenic functions in HPV-negative HNSCC, supporting that pharmacologic depletion rather than enzymatic inhibition is a necessary strategy to effectively target SMYD3 in HPV-negative HNSCC. We present the first-in-class PROTACs targeting SMYD3, with high potency in decreasing the proliferative and invasive capacity of HPV-negative HNSCC cells and a highly favorable pharmacokinetic profile in vivo, supporting feasibility of translation towards the clinic.

## RESULTS

### Pharmacologic inhibition of SMYD3 does not hinder the proliferation and invasion of HPV-negative HNSCC cells

Using SMYD3 depletion cell systems and mouse models, we recently reported the role of SMYD3 in the proliferative potential, colony formation, invasive capacity and immune evasion of HPV-negative HNSCC cell lines, making it a rational drug target for HPV-negative HNSCC [21-23]. We thus sought to therapeutically target SMYD3 using two commercially available and specific SMYD3 inhibitors, EPZ031686 and BAY-6035 [25, 26]. EPZ031686 was discovered in 2016 as the first selective SMYD3 inhibitor with double-digit nanomolar inhibitory potency in both biochemical (IC_50_=3nM) and cellular assays (IC_50_=36nM), and it functions in a non-competitive, allosteric manner to both S-adenosyl-methionine (SAM) and the MAP3K2 substrate [25]. BAY-6035 was discovered in 2021 as a selective, substrate-competitive inhibitor, showing double-digit nanomolar IC_50_s in biochemical (IC_50_=88nM) and cellular assays (IC_50_=70nM) [26].

To evaluate the pharmacodynamic efficacy of EPZ031686 as an enzymatic inhibitor in HPV-negative HNSCC cells, the global levels of H3K4me3, one of the enzymatic end products of SMYD3, were evaluated by Western blotting in three HPV-negative HNSCC cell lines with confirmed endogenous expression of SMYD3 (HN-6, HN-SCC-151, PE/CA-PJ15, **Supplementary Fig.1**). Cells were treated with EPZ031686 or DMSO as control for 6 days at a range of concentrations from 1-10uM (1, 2.5, 5, 10 μM). Results showed that H3K4me3 levels were decreased in a dose-dependent manner in all three cell lines, specifically by ∼50-80% at 5-10 μM (**Fig. 1A**). Given that SMYD3 also methylates MAP3K2, which induces activation of the downstream ERK signaling pathway, pERK levels were also evaluated by Western blotting and results showed a similar dose-dependent response with a decrease by ∼50-80% at 5-10 μM of treatment (**Fig. 1A**). Similar results were obtained with BAY-6035 (**Supplementary Fig. 2**).

**Figure 1.**
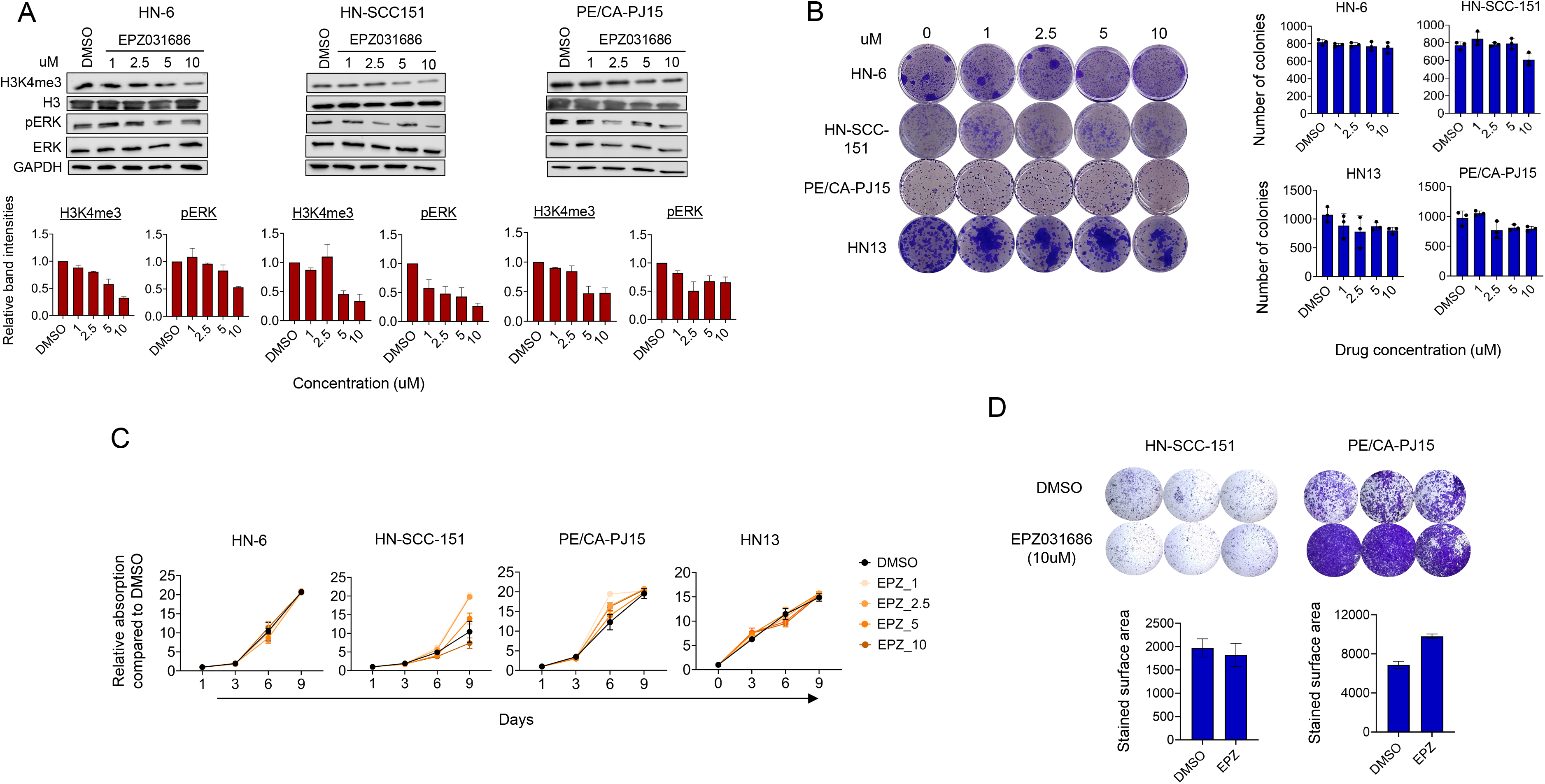
The SMYD3 inhibitor EPZ031686 does not hinder the proliferative and invasive capacity of HPV-negative HSNCC cells despite effective enzymatic inhibition. **(A)** Global H3K4me3 and pERK levels in HPV-negative HNSCC cells treated with EPZ031686. HN-6, HN-SCC-151 and PE/CA-PJ15 cells were treated with EPZ031686 versus DMSO at a concentration range of 1-10uM for 6 days. Nuclear and cytoplasmic extracts were obtained. 5ug of nuclear extracts were loaded and blotted for H3K4me3 and 20 μg of cytoplasmic extracts were loaded and blotted for pERK. H3 and GAPDH were used as loading controls. **Top:** Western blots for H3K4me3 and pERK. **Bottom:** Bar graphs of relative band intensities for H3K4me3 normalized to H3 and for pERK normalized to ERK. **(B)** Colony forming assays of HPV-negative HNSCC cells (HN-6, HN-SCC-151, PE/CA-PJ15 and HN13) treated with EPZ031686 versus DMSO at a concentration range of 1-10 μM for 10 days. Cells were seeded at ∼500-1000 cells/well in triplicates. After 10 days, cells were fixed and stained with crystal violet. Medium with the inhibitor was replenished every 48h. **Left:** Images of colonies in biological triplicates. **Right:** Bar graphs showing the average number of colonies for three biological replicates per condition. The number of colonies was counted using image J. Standard deviation (SD) is shown. **(C)** CCK8 assays of HPV-negative HNSCC cells (HN-6, HN-SCC-151, PE/CA-PJ15 and HN13) treated with EPZ031686 versus DMSO at a concentration range of 1-10uM for 9 days. Cells were seeded at ∼500 cells/well in biological quadruples. CCK8 assays were conducted at the indicated time points (days 1, 3, 6 and 9 of treatment). Curves are showing the average of four biological replicates per condition. Standard deviation (SD) is shown**. (D)** Invasion assays of HPV-negative HNSCC cells (HN-6, HN-SCC-151 and PE/CA-PJ15) treated with DMSO versus EPZ031686 at a concentration of 10uM for 6 days. Cells were seeded at ∼75,000 cells/transwell in biological triplicates. **Top:** Images of transwell in biological triplicates. **Bottom:** Bar graphs showing the average stained surface area of three biological replicates per condition.

To assess whether EPZ031686 decreases the proliferation of HPV-negative HSNCC cells, we conducted colony forming (CFAs) and CCK8 assays in four cell lines (HN-6, HN-SCC-151, PE/CA-PJ15, HN13) using a range of concentrations from 1-10 μM (1, 2.5, 5 and 10 μM). We found that EPZ031686 did not affect the proliferation of these cell lines (**Fig. 1B, C**). Furthermore, treatment of HPV-negative HNSCC cells with EPZ031686 did not hinder their invasive capacity (HN-SCC-151, PE/CA-PJ15) (**Fig. 1D**). Similar results were obtained with BAY-6035, with no effect in the colony forming and proliferative capacity of any of the four cell lines (**Supplementary Fig. 3**).

To further evaluate the importance of the enzymatic methyltransferase activity of SMYD3 in the proliferative and invasive capacity of HPV-negative HNSCC cells, we generated *SMYD3*-mutant cell lines stably transfected with a doxycycline-inducible plasmid expressing mutant SMYD3 (F183A). Stably expressing *SMYD3*-mutant cells exhibited decreased pERK protein levels by ∼60% compared to stably expressing *SMYD3* wild-type cells, confirming the enzymatic inactivity of SMYD3 in these cells (**Fig. 1E**). Interestingly, the global levels of H3K4me3 remained stable, a finding which we previously observed in SMYD3 KO cells possibly reflecting compensatory mechanisms (**Fig. 1E**, **Supplementary Fig. 4**) [23]. Importantly, the colony formation, proliferation and invasive capacity of these *SMYD3*-mutant cell lines were not significantly affected (**Fig.1 F-H**).

In contrast to these results, we previously reported that siRNA-mediated knockdown or knockout of SMYD3 led to a significant decrease in the colony formation, proliferation and invasion of HPV-negative HNSCC cells [23]. These findings support that inhibition of the catalytic activity of SMYD3 is not sufficient to abolish the oncogenic phenotypes of proliferation and invasion in HPV-negative HNSCC cells.

### Development and biochemical characterization of SMYD3 PROTACs

The above results support that the oncogenic functions of SMYD3 in HPV-negative HNSCC may be at least partially mediated through catalytically-independent mechanisms. As such, our group pursued the development of proteolysis-targeting chimeras (PROTACs) to suppress the catalytically-independent oncogenic effects of SMYD3 in HPV-negative HNSCC cells. Synthesis of PROTACs involves the conjugation of an E3-ubiquitin ligase ligand to a ligand targeting a protein of interest via a rigid or flexible chemical linker. For ligands targeting SMYD3, we considered the commercially available inhibitors EPZ031686, BAY-6035, BCI-121 and EPZ028862 (**Fig. 2A**, **Supplementary Fig. 5**) [25-29]. Among them, EPZ031686 is reported to have the highest enzymatic inhibitory potency, and was therefore chosen for further design of SMYD3 PROTACs (**Fig. 2A**). The bridged piperidine core of the EPZ031686 was chosen as a suitable linker attachment site based on the published crystal structure [25]. The trifluoro-group of the piperidine core in EPZ031686 was replaced with propyl amine to generate the modified inhibitor EPZ031686 propyl amine, enabling facile linker attachment and the development of PROTACs with diverse E3 ligase ligands (**Fig.2A**). A schema for the development of the SMYD3 PROTACs based on EPZ031686 propyl amine is shown in **Fig. 2A**.

**Figure 2.**
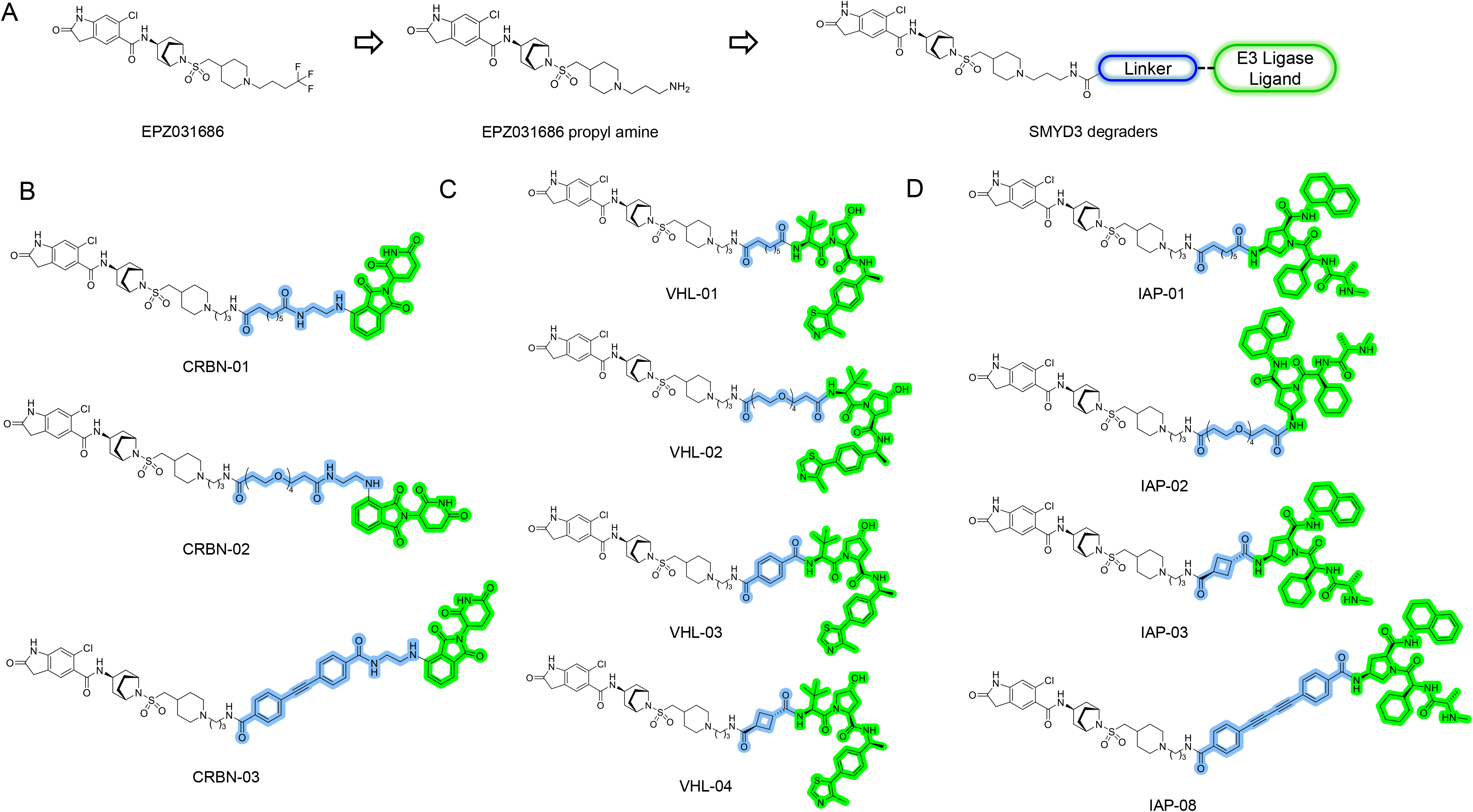
Development of SMYD3 PROTACs from the SMYD3 inhibitor EPZ031686. **(A)** Steps in the development of SMYD3 PROTACs based on EPZ031686. EPZ031686 propyl amine: replacement of the CF3 group of the piperidine core of EPZ031686 with propyl amine. POI: Protein Of Interest. **(B-D)** Chemical structures of the CRBN- **(B)**, VHL- **(C)** and IAP-based **(D)** SMYD3 PROTACs.

We then designed and developed a library of SMYD3 PROTAC compounds by coupling EPZ031686 to various E3-ubiquitin ligase ligands (Cereblon Cullin 4-Ring ubiquitin ligase CRBN, von Hippel-Lindau Cullin 2-Ring ubiquitin ligase VHL, and inhibitors of apoptosis proteins ubiquitin E3-ligase IAP) and diverse linkers for PROTAC conjugation. Each linker contained a bis-carboxyl group, with the second carboxyl group available to attach to various E3 ligase ligands. Initially, we synthesized PROTACs with CRBN, VHL and IAP employing a suberoyl linker, yielding the PROTACs CRBN-01, VHL-01 and IAP-01 (**Fig. 2B-D**). We then further modified these initial compounds by exploring different linker types, such as flexible (PEG) and rigid chemical linkers (aromatic, bridged and alkyne). Overall, we generated 15 SMYD3 PROTACs (**Supplementary Table 1**): (1) the CRBN-based PROTACs CRBN-01, -02 and -03 (**Fig. 2B**), (2) the VHL-based PROTACs VHL-01, -02, -03 and -04 (**Fig. 2C**), and (3) the IAP-based PROTACs IAP-01, -02, -03, -04, -05, -07, -08 and -09 (**Fig. 2D**), with IAP-01(neg) and IAP-08(neg) developed as negative controls with inversion of all stereocenters in the IAP ligand of PROTACs IAP-01 and IAP-08 respectively (**Supplementary Fig. 6**).

### IAP-08 efficiently degrades SMYD3

We then evaluated the pharmacodynamic efficacy of the SMYD3 PROTACs by evaluating SMYD3 protein levels in HPV-negative HNSCC cells (HN-6). Cells were treated with incremental concentrations of the PROTACs for 48h, nuclear extracts were obtained and Western blotting for SMYD3 was performed (**Supplementary Fig.7**). Among these compounds, IAP-08, with an 1,3-dialkyne-aromatic linker, was found to be the most potent, degrading SMYD3 protein levels by ∼ 85% at 0.1 μM and ∼95% at a concentration of 0.5 μM in HN-6 cells (**Fig. 3A**, **Supplementary Fig.7**, **Supplementary Fig.8**). The decrease in SMYD3 levels was associated with a dose-dependent decrease in global H3K4me3 and pERK levels (**Fig 3A**, **Supplementary Fig.8**). The protein levels of UHRF1, a direct gene target of SMYD3 [21], were also decreased in a dose-dependent manner, confirming the on-target effect of IAP-08 (**Fig. 3A**). Similar results were reproduced in three more HPV-negative HNSCC cell lines (HN-SCC151, PE/CA-PJ15, HN13) (**Fig.3A**, **Supplementary Fig.8**).

**Figure 3.**
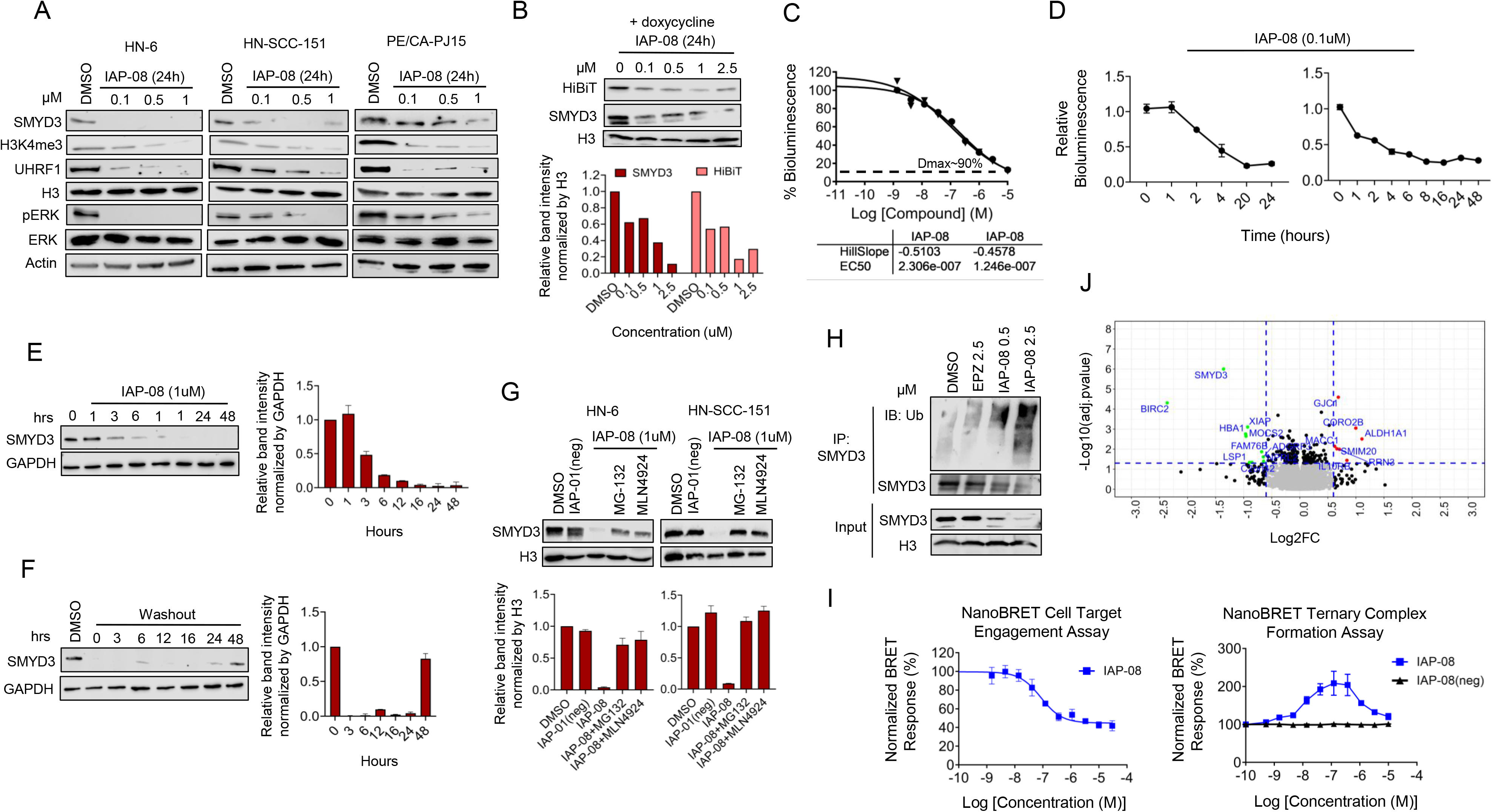
The SMYD3 PROTAC IAP-08 effectively and specifically degrades SMYD3 protein levels in HPV-negative HSNCC cells. **(A)** SMYD3, H3K4me3, UHRF1 and pERK protein levels in HPV-negative HNSCC cells treated with IAP-08. Cells (HN-6, HN-SCC-151, PE/CA-PJ15) were treated with IAP-08 versus DMSO at a concentration range of 0.1-1 μM for 48h. Nuclear and cytoplasmic extracts were obtained. 10ug of nuclear or 20ug of cytoplasmic extracts were loaded. H3 or actin were used as a loading control. **(B)** Western blotting for HiBiT-tagged SMYD3 in stably transfected HN13 cells exposed to doxycycline (1 μg/mL) and 24h later they were treated with IAP-08 at concentrations ranging from 0-2.5 μM for another 24h. Nuclear extracts were obtained, 10 μg were loaded and blotted for HiBiT and SMYD3. H3 was used as a loading control. Top: Western blots for HiBiT and SMYD3. Bottom: Bar graph of band intensities for total SMYD3 and HiBiT-tagged SMYD3. **(C)** HiBiT assay for SMYD3 in HN13 cells stably transfected with a doxycycline-inducible plasmid expressing HiBiT-tagged SMYD3. Cells were exposed to IAP-08 at a concentration range from 5e^-4^-10 μM for 24h. The DC_50_ curve was generated. The maximal level of degradation of SMYD3 is shown with the dotted line (D_max_). Biological duplicates per condition are shown. The y axis represents the % bioluminescence compared to DMSO (control) and the x axis represents the concentration of the compound (log[C]M). **(D)** Time-response curves obtained with HiBiT assays in HN13 cells stably expressing doxycycline-inducible HiBiT-tagged SMYD3. Cells were seeded in 6-well plates, and induction with doxycycline (1 μM/mL) was initiated. Next day, the cells were seeded into 96-well plates in triplicates (10,000 cells/well) and 24h later, treatment with IAP-08 at 0.1 μM was initiated. Bioluminescence was obtained at the indicated time points for a duration of 24h and 48h. **(E)** HN-SCC-151 cells were treated with IAP-08 at 1 μM and nuclear extracts were obtained at the indicated time points (0h-48h). 20ug of whole cell extracts were loaded and immunoblotting was conducted for SMYD3. GAPDH was used as a loading control. **(F)** HN-SCC-151 cells were treated with IAP-08 at 1uM for 24h, “washed” with PBS and then cell lysates were obtained at the indicated time points (0h-48h). 2 μg of whole cell extracts were loaded and blotted for SMYD3. GAPPDH was used as a loading control. **(G)** SMYD3 protein degradation is rescued through inhibition of the proteasomal pathway. HN-6 or HN-SCC-151 cells were treated with IAP-08 or its epimer control -IAP-01(neg) at 1 μM with or without the presence of the proteasomal inhibitor MG132 or the neddylation inhibitor MLN4924 for 48h. Nuclear extracts were obtained, 10 μg were loaded and blotted for SMYD3. H3 was used as a loading control. **(H)** HN-6 cells were treated for 6h with IAP-08 at 0.5 μM and 2.5 μM, the SMYD3 inhibitor EPZ031686 at 2.5 μM and DMSO. Whole-cell extracts were obtained and immunoprecipitation was conducted for SMYD3. Immunoprecipitates were blotted for ubiquitin and SMYD3. Input is shown for SMYD3, using H3 as a loading control. **(I)** NanoBRET assays. **Left:** NanoBRET target engagement cellular SMYD3 assay. HEK293 cells were transiently transfected with an N-terminal nanoluciferase (NanoLuc)-tagged SMYD3 construct and were treated with incremental concentrations of IAP-08 for 2h. MG-132 (10 μM) was used to inhibit proteasomal degradation of SMYD3. **Right:** NanoBRET cellular ternary complex formation assay for XIAP/SMYD3/IAP-08. IAP-08(neg) (negative control) has the same structure as IAP-08 except for the IAP ligand which is the enantiomer of the IAP ligand of IAP-08. HEK293 were co-transfected with plasmids encoding SMYD3-NanoLuc fused target protein and XIAP-HaloTag E3 ligase and incubated with incremental concentrations of IAP-08 for 2h. The BRET ratio was calculated as follows: BRET ratio = [(Acceptor sample)/(Donor sample)] – [(Acceptor no-tracer control)/(Donor no tracer control)]. Normalized BRET Response (%)=(BRET ratio of each treatment sample/ Average BRET ratio of DMSO control samples)*100%. The IC50 curves were plotted and IC_50_ values were calculated using the GraphPad Prism program based on a sigmoidal dose-response equation. SDs are shown from two independent biological replicates. **(J)** Volcano plot of protein levels in HN-SCC-151 cells treated with the SMYD3 PROTAC versus control. Cells were treated with IAP-08 at 1 μM or control DMSO. Protein extracts were obtained and quantified by tandem-mass-tag (TMT) mass spectrometry analysis. Results represent averages of four biological replicates.

To validate these results, we generated a doxycycline-inducible stable cell line (HN13 parental cells) expressing HiBiT-tagged SMYD3. This technology is based on a split luciferase system, whereby an 11-amino-acid epitope HiBiT-tagged protein fuses with its complementation partner LargeBit forming a functional NanoLuc luciferase, allowing for the relative quantification of intracellular SMYD3 protein levels through bioluminescence in living cells. Bioluminescence was increased upon exposure to doxycycline, confirming the functionality of this doxycycline-inducible cell system (**Supplementary Fig. 9**). To further validate that the bioluminescence was secondary to the expression of HiBiT-tagged SMYD3, the stably transfected HN13 cells were induced with doxycycline, treated with IAP-08 at a range of concentrations (0-2.5 μM) for 24h, and Western blotting for HiBiT-tagged and total SMYD3 was performed in nuclear extracts. HiBiT-tagged and total nuclear SMYD3 protein levels decreased in a dose-dependent manner upon incremental concentrations of the SMYD3 PROTAC, with ∼80% of HiBiT-SMYD3 protein levels depleted at 1 μM of SMYD3 PROTAC treatment for 24h (**Fig. 3B**).

Following the validation of this cell system, HiBiT-tagged SMYD3 expressing HN13 cells were induced with doxycycline and then exposed to all CRBL-, VHL- and IAP-based SMYD3 PROTACs at range of concentrations up to 10 μM for 24h. The half-maximal degradation concentration (DC_50_, uM) and the maximal degradation (D_max_, %) was calculated for all compounds, and IAP-08 was confirmed as the most potent compound with DC_50_ at 0.177 μM and D_max_ at ∼90% (**Fig. 3C**, **Supplementary Fig.10**). We thus decided to further focus our analysis on IAP-08.

To further assess the kinetics of the degradation of HiBiT-tagged SMYD3 by IAP-08, we conducted HiBiT assays in the stably expressing HN13 cells exposed to concentrations ranging from 0-5 μM of the PROTAC, and dose-response curves were generated. At 4h and 24h respectively, nearly 45% and ∼80% of HiBiT-tagged SMYD3 levels were degraded at 0.1 μM (**Supplementary Fig.11**). At 24h of treatment, protein degradation plateaued at 0.1 μM, while at 4h of treatment, further degradation was observed up to ∼65% at 5 μM of the SMYD3 PROTAC (**Supplementary Fig.11**). We then performed a HiBiT assay in stably expressing HN13 cells treated with 0.1 μM of IAP-08 at time points ranging from 0-24h and 0-48h, and time-response curves were generated (**Fig. 3D**). In accordance with the aforementioned results, ∼40-45% of HiBiT-tagged SMYD3 was degraded at 4h of PROTAC treatment as evidenced in both time-response curves (**Fig. 3D**), and a steady state was reached at ∼20-24h, with nearly 80-90% of the HiBiT-tagged SMYD3 degraded (**Fig. 3D**).

To further evaluate the kinetics of SMYD3 protein degradation after treatment with IAP-08, Western blotting was pursued at multiple time points in HN-SCC-151 cells treated with the SMYD3 PROTAC at 1 μM for a duration ranging from 0-48h (1, 3, 6, 12, 16, 24 and 48h). SMYD3 protein degradation occurred within 3 hours of treatment and was sustained up to the last examined time point of PROTAC treatment at 48h (**Fig. 3E**). To further assess the sustainability of the SMYD3 protein degradation induced by IAP-08, HN-SCC-151 cells were treated for 24h with the SMYD3 PROTAC at 1 μM, “washed” with PBS and then collected at time points ranging from 0-48h (**Fig. 3F**). SMYD3 protein levels remained degraded by ∼90% 24h after the removal of the SMYD3 PROTAC, and increased back to baseline at 48h.

### IAP-08 degrades SMYD3 in a proteasomal- and IAP-dependent manner

To assess whether IAP-08 mediated the degradation of SMYD3 specifically through the ubiquitination pathway, HN-6 and HN-SCC-151 cells were treated with IAP-08 (1uM) for 48h, with or without the presence of the proteasomal inhibitor MG-132 or MLN4924 (pevonedistat), a neddylation inhibitor of NEDD8 which activates the cullin-RING E3 ubiquitin ligases to attach ubiquitin chains to specific target proteins. As a control, we treated cells with IAP-01(neg), which carries a biologically-inactive cis-epimer of the IAP ligand that cannot trigger the IAP ligase and thus is not expected to degrade SMYD3 (**Supplementary Fig. 6B**). SMYD3 degradation by IAP-08 was rescued with bortezomib or MLN4924, confirming that IAP-08 degrades SMYD3 through the proteasomal pathway (**Fig. 3G**). Furthermore, IAP-01(neg) did not induce degradation of SMYD3, supporting the necessity of the IAP ligand moiety for IAP-08-induced degradation of SMYD3 (**Fig. 3G**). We further assessed whether treatment with IAP-08 induced polyubiquitination of SMYD3 by blotting whole-cell extracts from HN-6 cells treated with incremental doses of IAP-08 for 6h for ubiquitin. Results showed that IAP-08 induced dose-dependent polyubiquitination of SMYD3, whereas the controls DMSO and EPZ031686 did not (**Fig. 3H**).

### IAP-08 effectively engages and forms intracellular ternary complexes with SMYD3 and XIAP

To evaluate the engagement of intracellular SMYD3 with IAP-08, we established a NanoBRET target engagement cellular SMYD3 assay using HEK293 cells. Specifically, an N-terminal nanoluciferase (NanoLuc)-tagged SMYD3 construct was generated, transiently transfected into HEK293 cells and the assay was validated through competitive displacement by a fluorescent tracer. The transfected HEK293 cells were first treated with incremental concentrations of EPZ031686 as a positive control, which fully engaged with cellular SMYD3 with an EC_50_ at ∼13 nM (100% cell target engagement, **Supplementary Fig. 12**). The transfected HEK293 cells were then treated with incremental concentrations of IAP-08 in combination with MG132 to stabilize SMYD3 and enable accurate target engagement readouts. IAP-08 engaged SMYD3 with an EC_50_ at ∼85 nM, though complete engagement was not achieved and saturation occurred at 60% (**Fig. 3I**). The lower target engagement for IAP-08 compared to EPZ031686 was expected, given that PROTACs are substantially larger molecules compared to their counterpart inhibitors, thus reducing their cell permeability and intracellular free concentration.

To assess the effectiveness of IAP-08 in inducing the formation of ternary complexes with IAP E3-ligases, we generated a NanoBRET cellular ternary complex assay by co-expressing Halo-tagged XIAP with NanoLuc-tagged SMYD3 in HEK293 cells (**Fig. 3I**). The transfected cells were incubated with NanoBRET HaloTag 618 Ligand, a fluorescence tracer with an XIAP ligand. Results indicated that treatment of these cells with incremental concentrations of IAP-08 was associated with an increasing BRET ratio signal compared to its negative control (inverse IAP ligand of IAP-08) (**Fig.3I**). The formation of a cellular ternary complex between XIAP and SMYD3 through IAP-08 was confirmed with an EC_50_ at ∼11 nM. The familiar hook effect was observed at higher concentrations. As expected, ternary complex formation was not observed with the IAP-08 negative control compound (**Fig. 3I**, **Supplementary Fig. 6B**), consistent with diminished binding of XIAP and SMYD3.

### IAP-08 specifically targets SMYD3

To assess the specificity of IAP-08 towards SMYD3, HN-SCC-151 cells were treated with IAP-08 at 1 μM for 48h, while IAP-01(neg) and the acetylated benzoyl-IAP (Bz-IAP), which is a biologically-active IAP antagonist ligand, were used as controls. Protein extracts were obtained and tandem mass tag (TMT) mass spectrometry was conducted. Results showed that SMYD3 was amongst the top two most degraded proteins in HN-SCC-151 cells treated with the SMYD3 PROTAC (log2 (IAP-08/DMSO)=-1.358, decrease by ∼96%, adj. p-value=1e^-6^), while the control epimer IAP-01(neg) and the Bz-IAP did not degrade SMYD3 (**Supplementary Tables 2, 3**). Furthermore, defining degradation as significant at -log10(adj.p-value)>1.3 (<0.05) and log2FC<(-0.6) (>75%) decrease, only seven off-target proteins were detected, including E3 ubiquitin-protein ligase X-linked inhibitor of apoptosis protein (XIAP), Baculoviral IAP repeat-containing protein 2 (BIRC2), Hemoglobin subunit alpha (HBA1), Molybdopterin synthase catalytic subunit (MOCS2), Lymphocyte-specific protein 1 (LSP1), the GATOR complex protein NPRL2 and Family With Sequence Similarity 76 Member B (FAM76B) (**Fig.3J**). These results support that IAP-08 specifically targets SMYD3 through an IAP-dependent proteasomal manner.

### IAP-08 significantly decreases the proliferative and invasive capacity of HPV-negative HNSCC cells

To evaluate the effect of IAP-08 on the proliferative capacity of HPV-negative HNSCC cells, four HPV-negative HNSCC cell lines (HN-6, HN-SCC-151, PE/CA-PJ15 and HN13) were treated with IAP-08 for 9-10 days at concentrations ranging from 0.05-2.5 μM, and CFAs and CKK8 assays were pursued. Treatment with IAP-08 induced a significant decrease by ∼90% in the colony forming capacity and relative number of these cells at concentrations as low as 0.05-1 μM (**Fig. 4A, B**). This was in contrast to the enzymatic inhibition through EPZ031686 which did not affect the proliferation of these cells at concentrations as high as 10 μM with the same duration of exposure (**Fig.1B, C**). Importantly, the colony forming capacity of the normalized bronchial mucosal cell line BEAS-2B was unaffected by IAP-08 despite the high expression of SMYD3 and XIAP in these cells (**Fig.4A**, **Supplementary Fig. 1**), signifying the specific effect of IAP-08 on HPV-negative HNSCC cancer cells but not in normalized epithelial cells. Furthermore, markers of apoptosis were affected in a dose-dependent manner, with a decrease in PARP and an increase in cleaved PARP and cleaved caspase 3, supporting the induction of apoptosis in HPV-negative HSNCC cells treated with IAP-08 (**Fig. 4C**).

**Figure 4.**
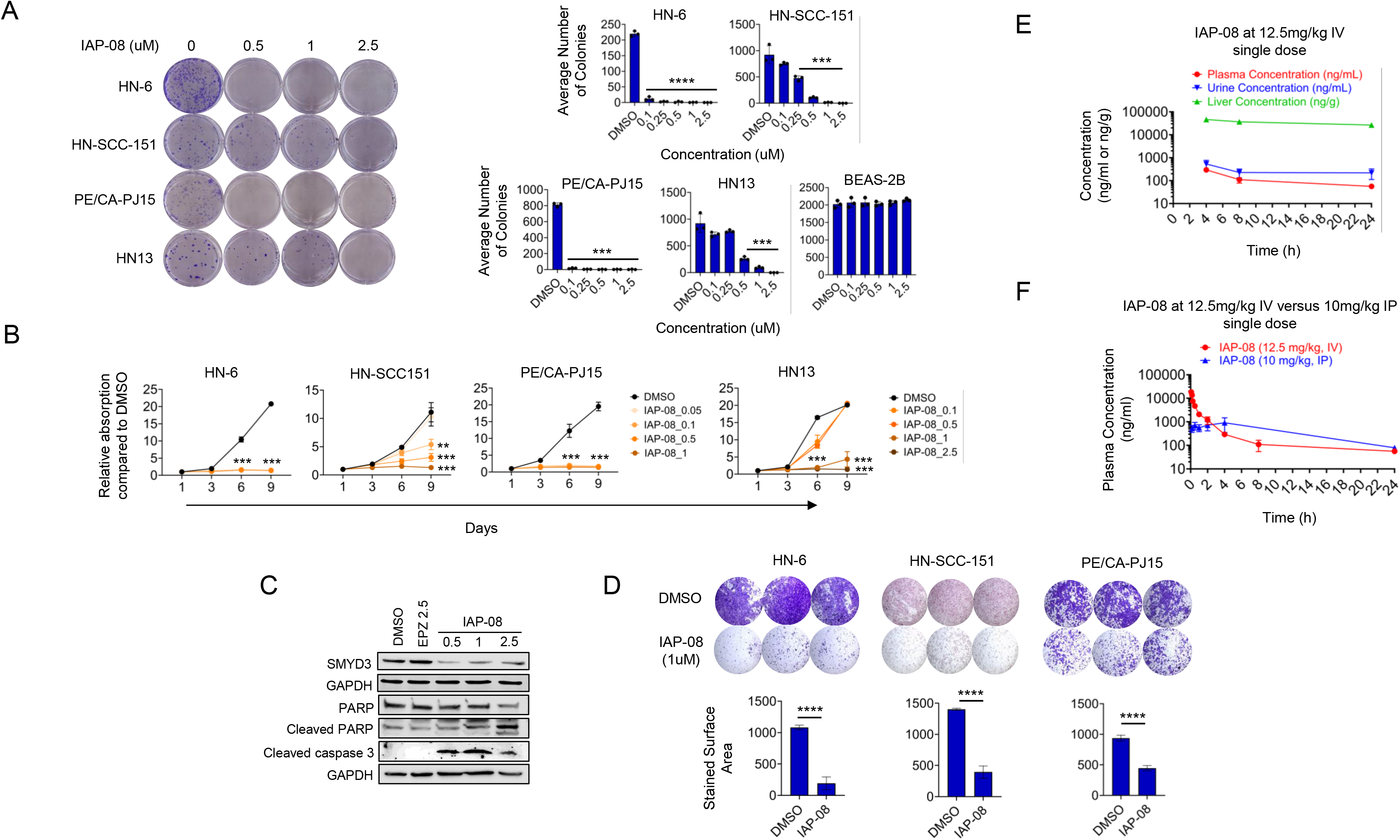
The SMYD3 PROTAC IAP-08 decreases the proliferative and invasive capacity of HPV-negative HNSCC cell lines in vitro and restrains tumor growth in vivo. **(A)** Colony forming assays of HPV-negative HNSCC cells (HN-6, HN-SCC-151, PE/CA-PJ15 and HN13) treated with IAP-08 versus DMSO at a concentration range of 0.1-2.5 μM for 9-12 days. Cells were seeded at ∼500-1000 cells/well in triplicates. After 9-12 days, cells were fixed and stained with crystal violet. Left: Images of colonies in biological triplicates. Right: Bar graphs showing the average of three biological replicates per condition. The number of colonies was counted using Image J. Standard deviation (SD) is shown. Student t-test, *p<0.05, **p<0.01, ***p<0.005, ****p<0.001. **(B)** CCK8 assays of HPV-negative HNSCC cells (HN-6, HN-SCC-151, PE/CA-PJ15 and HN13) treated with IAP-08 versus DMSO at a concentration range of 0.05-2.5 μM for 9 days. Cells were seeded at ∼500 cells/well in biological quadruples. CCK8 assays were conducted at the indicated time points (days 1, 3, 6 and 9 of treatment). Curves are showing the average of four biological replicates per condition. Standard deviation (SD) is shown. Student t-test, * p<0.05, **p<0.01, ***p<0.005, ****p<0.001. **(C)** Markers of apoptosis in HN-SCC-151 cells treated with IAP-08 (0.5-2.5uM) versus EPZ031686 (2.5uM) or DMSO for 24h. Cytoplasmic extracts were obtained and blotted for PARP, cleaved PARP, cleaved caspase 3 (50ug) and for SMYD3 (30ug). GAPDH was used as a loading control. **(D)** Invasion assays of HPV-negative HNSCC cells (HN-6, HN-SCC-151, PE/CA-PJ15) treated with IAP-08 versus DMSO at a concentration of 1uM for 24h. Cells were seeded at ∼75,000 cells/transwell in biological triplicates. Top: Images of transwells in biological triplicates. Bottom: Bar graphs showing the average of three biological replicates per condition. Unpaired t-test, * p<0.05, **p<0.01, ***p<0.005, ****p<0.001. **(E)** Mouse plasma, urine and liver homogenate concentrations of IAP-08 after a single dose of 12.5mg/kg administered IV in nude BALB/C mice. Samples were obtained from three mice at the indicated time points (4h, 8h, 24h). **(F)** Mouse plasma concentrations of IAP-08 after a single dose of 12.5mg/kg administered IV or 10mg/kg administered IP in nude BALB/C mice. Samples were obtained from three mice at the indicated time points (0.08h, 0.17h, 0.25h, 0.5h, 1h, 2h 4h, 8h and 24h).

Invasion assays were also conducted in HN-6, HN-SCC-151 and PE/CA-PJ15 cells treated with IAP-08 and showed a significant decrease in the invasive capacity of these cells (**Fig. 4D**), in contrast to EPZ031686 which had no effect on this phenotype (**Fig. 1D**).

### IAP inhibition does not contribute to the antiproliferative effects of IAP-08 at concentrations lower than 1 μM

Inhibitors of apoptosis proteins (IAPs), such as XIAP and BIRC2, have established antiapoptotic effects and are known to function as oncogenes in HPV-negative HNSCC [30-32]. Our MS analysis revealed that IAP-08 was associated with degradation not only of SMYD3, but also of XIAP and BIRC2, raising the possibility that the antiproliferative effects observed with IAP-08 may be due not only to SMYD3 depletion but also to XIAP and BIRC2 degradation. To evaluate this, we treated two HPV-negative HNSCC cell lines (HN-SCC-151, HN13) with incremental concentrations of a highly potent IAP antagonist (A 410099.1) which has an identical structure as the IAP ligand moiety of IAP-08 and compared its efficacy to IAP-08 at the same concentrations through colony forming assays (**Supplementary Fig. 13**). Results showed that while IAP-08 significantly decreased the number of colonies and the relative cell number of both cell lines at concentrations less than 1uM, the IAP antagonist did not significantly affect it, supporting that the antiproliferative efficacy of IAP-08 is due to the degradation of SMYD3 and not of XIAP/BIRC2.

### IAP-08 has a favorable pharmacokinetic profile in vivo

To evaluate the pharmacokinetic (PK) properties and safety profile of IAP-08, an in vivo study with nude BALB/C mice was initially conducted with the administration of 3 different single doses of the PROTAC intravenously (IV) (12.5mg/kg, n=9; 25mg/kg, n=2; 50mg/kg, n=2). From the three doses, the single IV dose of 12.5mg/kg IV was tolerated very well with no deaths (n=9 mice). Blood, urine and liver homogenate samples were obtained at given time points after the dose administration (0.08h, 0.17h, 0.25h, 0.5h, 1h, 2h 4h, 8h and 24h) for quantitative analysis of the PROTAC using liquid chromatography-mass spectrometry. The PK parameters of IAP-08 revealed an AUC_last_ of 12,542 h*ng/ml and a favorable half-life of ∼9.7h, similar to that of irinotecan (t_1/2_ ∼6-12h in humans) [31]. Comparing the concentration of IAP-08 in the mouse liver, urine and plasma compartments, the highest concentrations were observed in the liver, supporting predominant uptake by the liver (**Supplementary Fig. 14**, **Fig. 4E**).

To further evaluate and compare the PK profile of IAP-08 administered through IV versus intraperitoneal (IP) route, nude BALB/C mice (n=9 per route) were dosed with a single dose of 12.5 mg/kg IV versus a single dose of 10 mg/kg IP, and the aforementioned measurements were pursued. The AUC levels were comparable between the 10 mg/kg IP (AUC_last_=12,808 h*ng/ml) and 12.5 mg/kg IV doses (**Fig. 4F**), indicating similar systemic exposure via both administration routes. To evaluate the metabolic profile of IAP-08, in vitro plasma protein binding assays and liver microsome assays were pursued (**Supplementary Fig.15**). The protein binding capacity of IAP-08 in human and mouse plasma was assessed at 99%. Using human or mouse liver microsomes in vitro and after 60min of incubation with the compound, approximately 56% and 59% of the compound was remaining, with an intrinsic clearance of 97.6 and 89.1 uL/min/mg respectively and a half-life of more than 60min, indicating a moderate clearance and a stable microsomal metabolic profile of IAP-08.

These results support a favorable PK profile for IAP-08, with a half life of ∼9h, a high plasma protein binding capacity and a stable liver metabolic profile.

### Depletion but not enzymatic inhibition of SMYD3 regulates the transcription of cell cycle-, EMT- and immune-related genes in HPV-negative HNSCC cells

We then sought to evaluate the effect of the IAP-08 on the transcriptional profile of HPV-negative HNSCC cells, and to compare it with the effect of EPZ031686. To this purpose, we treated HN-SCC-151 cells with IAP-08 (0.5 μM) or EPZ031686 (10 μM) versus DMSO for 48h and RNA-seq was conducted. IAP-08 induced transcriptional regulation of 3,539 genes (1,930 upregulated, 1,609 genes downregulated), whereas EPZ031686 affected only 385 genes (283 upregulated, 102 downregulated) (log2FC>0.3, FDR<0.1), supporting that SMYD3 depletion has a greater impact on the transcriptional profile of HPV-negative HNSCC cells compared to SMYD3 enzymatic inhibition (**Fig.5A-B**, **Supplementary Table 4**). Interestingly, the majority of the genes regulated by EPZ031686 (74%) were also regulated by IAP-08, while only 10% of the genes regulated by IAP-08 were commonly regulated by EPZ031686 (**Fig. 5C**).

**Figure 5.**
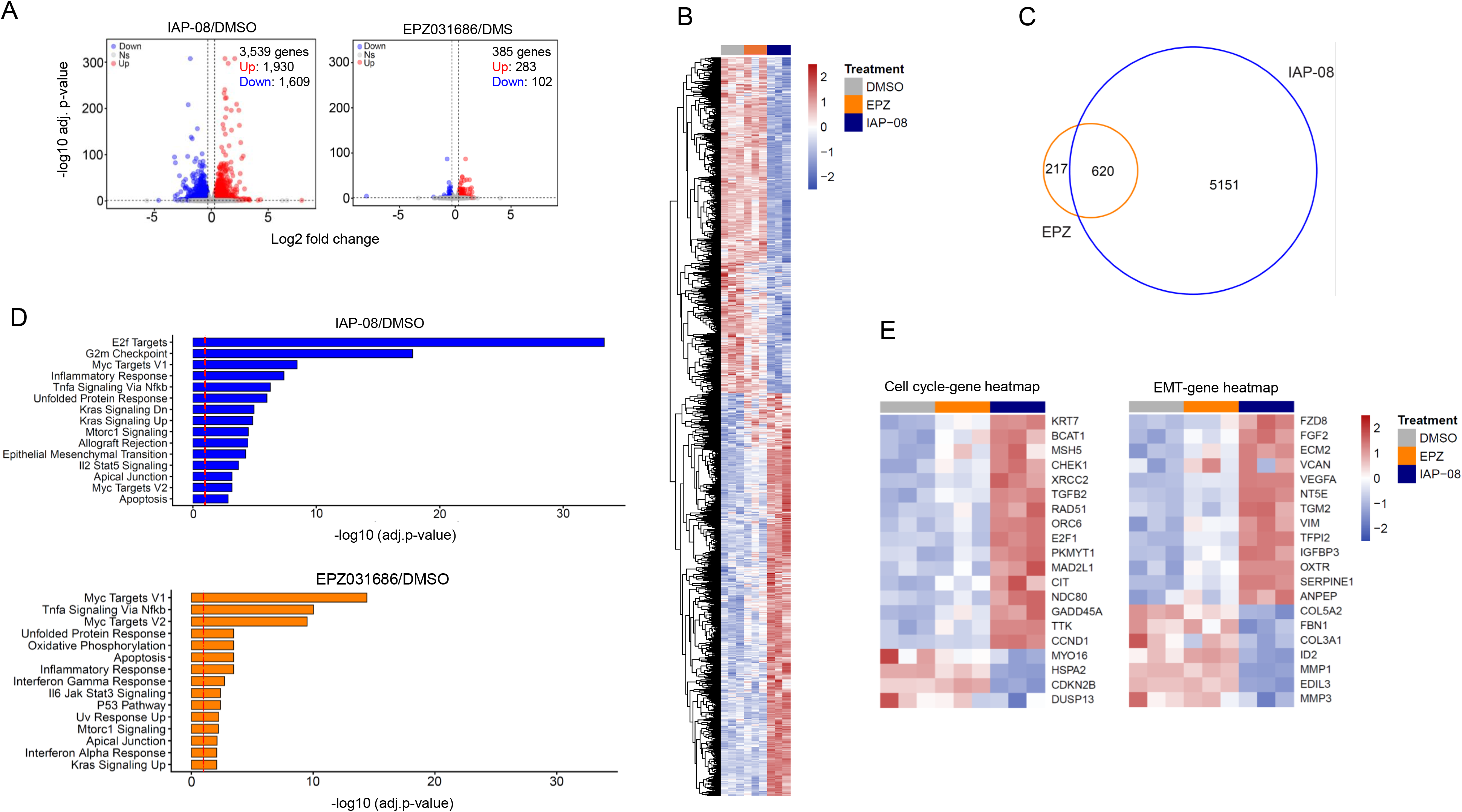
Effects of IAP-08 versus EPZ031686 on the transcriptomic profile of HPV-negative HNSCC cells. **(A)** Volcano plots of RNA-seq data derived from HN-SCC-151 cells treated with by IAP-08 (0.5uM) or EPZ031686 (10uM) compared to DMSO for 48h. FDR<0.1, log2FC>±0.3. NS: not significant. Left plot: 3,539 were significantly affected, with 1,930 upregulated and 1,609 downregulated. Right plot: 385 genes were significantly affected, with 283 upregulated and 102 downregulated. **(B)** Heatmap derived from RNA-seq of HN-SCC-151 cells treated with IAP-08 (0.5uM), EPZ031686 (10 μM) or DMSO for 48h. FDR<0.1, log2FC>±0.3. **(C)** Overlap of genes regulated by IAP-08 (0.5 μM) or EPZ031686 (10 μM) in HN-SCC-151 cells treated for 48h. FDR<0.1. **(D)** Gene Set Enrichment Analysis (GSEA) of RNA-seq data obtained from HN-SCC-151 cells treated with IAP-08 (0.5uM) or EPZ031686 (10 μM) versus DMSO for 48h. Pre-ranked GSEA using all tested genes. Y axis: -log10 (adjusted p-value). Red dotted line indicating –log10 (adj.p-value)=0.1. Top 15 Hallmark pathways shown with FDR (adjusted p-value)<0.1. **(E)** Heatmaps of top 20 cell-cycle- and EMT-related genes with the greatest log2 fold change induced by treatment of HN-SCC-151 cells with IAP-08 (0.5 μM) or EPZ031686 (10 μM) versus DMSO for 48h. The ratio of up- over down-regulated genes represents the one observed in the respective gene list. FDR<0.1.

GSEA of IAP-08 treated HNSCC cells revealed enrichment in cell cycle- (E2F targets, G2M checkpoint, Mitotic spindle, MYC targets V1, V2), epithelial mesenchymal transition (EMT) and immune-related (Inflammatory response, IL2_STAT5 signaling, IFNA and IFNG response) Hallmark pathways (**Fig. 5D**, **Supplementary Table 5**), in accordance with our previous reports showing that genetic SMYD3 depletion regulates these pathways [20-21]. In contrast, treatment with the SMYD3 inhibitor was associated with a comparatively lower degree of pathway enrichment (**Fig. 5D**), consistent with the lack of effect in the proliferation and invasion phenotypes of HPV-negative HNSCC cells (**Fig. 1B-D**). Accordingly, focusing on the genes comprising the cell-cycle-, EMT- and IFN response HALLMARK pathways (**Supplementary Table 6**), many of these were found transcriptionally regulated by IAP-08 but not by EPZ031686 (**Fig. 5E**, **Supplementary Table 7**, **Supplementary Fig. 16**).

These results further support that SMYD3 affects the expression of these genes in a predominantly catalytically-independent manner.

## DISCUSSION

Despite the recent advent of pembrolizumab immunotherapy as first-line treatment, HPV-negative HNSCC has a dismal prognosis [1-3]. Novel therapeutic approaches are thus urgently needed. Our group recently reported that SMYD3, which is overexpressed in ∼60% of HPV-negative HNSCC, drives proliferation and invasiveness in HPV-negative HNSCC, and is a major epigenetic repressor of cancer-cell intrinsic type I IFN responses inducing resistance to anti-PD-1 immunotherapy [21-23]. SMYD3 has also emerged as an important oncogenic driver in multiple other solid tumor types, including HCC, colon, breast and ovarian carcinoma [4-20].

As such, SMYD3 inhibitors with excellent biochemical and cellular inhibitory potency (≤1 μM) have been developed [25-29, 36-39]. However, when we tested the in vitro efficacy of two of these inhibitors, EZP031686 and BAY-6035, in our HPV-negative HNSCC cell lines, their proliferative and invasive capacity was not affected despite effective enzymatic inhibition of SMYD3 (**Fig. 1**, **Supplementary Fig. 3**). These results indicated that inhibition of the enzymatic activity of SMYD3 is not sufficient to hinder its oncogenic functions in HPV-negative HNSCC cells. In accordance to our findings, Thomenius et al tested the highly potent SMYD3 inhibitor EPZ028862 (biochemical IC_50_=1.8 nM, cellular IC_50_=32 nM) across 240 cancer cell lines and found no impact on the cellular proliferation of these cell lines at concentrations up to 25 μM [39]. Importantly, while a number of groups have reported that enzymatic SMYD3 inhibition with BCI-121, one of the first ever reported SMYD3 inhibitors in the field [27], induces a modest decrease in the cellular proliferation of various cancer cell lines [7-11, 13, 14], the high concentrations required to induce a cellular pharmacodynamic effect (∼100 μM) raise concerns for possible off-target effects of this inhibitor [27]. Importantly, while these data support that enzymatic inhibition of SMYD3 is not sufficient to effectively hinder the proliferative capacity of cancer cells, a few studies have highlighted the importance of the enzymatic activity of SMYD3 in the phenotypes of invasion and chemosensitivity. Specifically, Ikram et al [15] showed that SMYD3 promotes invasion, migration and anchorage-independent growth in a catalytically-dependent manner through methylation of MAP3K2 in prostate cancer cell lines and orthotopic xenograft mouse models expressing the catalytically-inactive SMYD3_F183A_ or treated with the SMYD3 inhibitor EPZ031686. According to our findings, they also found that SMYD3 enzymatic inhibition did not affect the proliferative capacity of prostate cancer cell lines. Furthermore, Sanese et al [41] showed that SMYD3 inhibition with EM127 increased the chemosensitivity of colorectal, breast and gastric cancer cell lines to the DNA-damaging agents 5-fluorouracil, irinotecan and oxaliplatin both in vitro and in vivo, consistent with the reported role of SMYD3 in the regulation of DNA-damage-response and homologous recombination repair [16]. Overall, the above data support that the importance of the enzymatic activity of SMYD3 in exerting its oncogenic functions is cell-context dependent; while it may be important in promoting invasion in prostate cancer and chemoresistance in breast, colorectal and gastric cancer cell lines, enzymatic inhibition is not sufficient to repress the invasive and proliferative capacity of HPV-negative HNSCC cells.

We thus reasoned that effective therapeutic targeting for SMYD3 in HPV-negative HNSCC, as well as other cancer types, would require depletion/degradation of SMYD3 since this approach would abrogate both its catalytically-independent and -dependent mechanisms. To this purpose, we designed a series of 15 novel SMYD3 PROTACs utilizing one of the most potent commercially available SMYD3 inhibitors, EPZ031686, and coupling it to all three types of E3 ligase warheads (IAP, VHL, CRBN) and various linkers (suberoyl, PEG, aromatic, bridged, and alkyne linkers) (**Fig. 2**, **Supplementary Table 1**). Of these PROTACs, IAP-08 with an alkyne-aromatic linker was the most potent at degrading SMYD3 (DC50=177nM, Dmax ∼90%) (**Fig.3**, **4**). IAP-08 induced SMYD3 protein degradation specifically through the proteasome system, as inhibition of the proteasome or the neddylation enzyme NEDD8 rescued the degradation of SMYD3 by IAP-08 (**Fig. 3**). Additionally, SMYD3 was amongst the top two most significantly and deeply degraded protein targets (**Fig. 3**), while treatment of HPV-negative HNSCC cells with IAP-01(neg), which does not activate the proteasomal pathway, and the Bz-IAP antagonist ligand alone did not induce SMYD3 protein degradation (**Supplementary Table 2, 3**), further supporting the specificity of IAP-08.

In accordance with our hypothesis that SMYD3-driven oncogenic phenotypes in HPV-negative HNSCC cells are predominantly catalytically-independent, IAP-08 drastically decreased the proliferation of four HPV-negative HNSCC cell lines at concentrations as low as 50-500 nM (**Fig. 4A-B**). This was associated with a dose-dependent decrease in SMYD3 protein levels, as well as known gene targets, enzymatic endproducts and downstream effectors of SMYD3, such as UHRF1 [21], H3K4me3 [21] and pERK [18], supporting these proteins as on-target pharmacodynamic biomarkers of response to IAP-08. Interestingly, these proteins were decreased by ∼80-100% in all HPV-negative HNSCC cell lines treated with IAP-08 at 1 μM for 48h, indicating effective depletion of SMYD3 and its downstream targets. In contrast, the SMYD3 inhibitors EPZ031686 and BAY-6035 did not hinder the proliferation of the same cell lines even at concentrations as high as 10 μM, despite enzymatic inhibition evidenced by a decrease in H3K4me3 and pERK levels by ∼60-80% (**Fig. 1**, **Supplementary Fig. 2**, **3**). The invasiveness of HPV-negative HNSCC cells was also hindered after treatment with 1 μM of the SMYD3 PROTAC IAP-08 (**Fig. 4D**), but not with 10 μM of the SMYD3 inhibitor EPZ031686 (**Fig. 1D**). Accordingly, in contrast to EPZ031686 which only affected the transcription of 837 genes, IAP-08 affected the transcription of 5,771 genes compared to control (DMSO) (**Fig. 5A-B**). The limited effect of SMYD3 inhibition on the transcriptomic profile of HPV-negative HNSCC cells was further supported by GSEA showing a lower level of enrichment of oncogenic pathways in cells treated with EPZ031686 (FDR=0.09-10^-15^), in contrast to the highly significant enrichment of cell cycle-, EMT- and immune-related pathways in cells treated with IAP-08 (FDR=0.08-10^-34^) (**Fig. 5C**, **Supplementary Table 5**).

Our finding that IAP-based PROTACs were more effective at degrading SMYD3 in HPV-negative HNSCC cells compared to CRBN- and VHL-based PROTACs may be because of the higher cellular availability of IAP E3 ligases compared to CRBN and VHL E3 ligases in these cells. Higher E3 ligase levels are expected to lead to higher amounts of ternary PROTAC complexes which allows for more effective target protein degradation. Indeed, XIAP, the E3 ligase that the IAP-ligand of IAP-08 binds, was found to be present at nearly 8 times higher compared to VHL and 4 times higher compared to CRBN in HN-SCC-151 cells (**Supplementary Table 8**). Furthermore, the potency of a PROTAC is heavily influenced by its ability to induce a stable and productive ternary complex with the target protein of interest and the E3 ligase. As such, the optimal linker length, flexibility, rigidity and hydrophobicity, as well as the spatial orientation of the target protein of interest and the E3 ligase are crucial for efficient ubiquitination. IAP-08 contains a rigid and hydrophobic linker, specifically consecutive alkyne-containing aromatic rings, which can form additional favorable non-covalent interactions (e.g., π-stacking or hydrophobic contacts) with surface residues of either the IAP E3 ligase or the SMYD3 protein within the ternary complex. We speculate that these interactions may improve the formation of ternary complexes and enhance cellular permeability, leading to the observed more efficient SMYD3 degradation with IAP-08 compared to other flexible linkers. Comparative 3D-crystallography amongst the SMYD3 PROTACs could further help solidify the above possibilities.

One of the major impediments in the translation of PROTACs to the clinic has been their unfavorable PK profile in vivo due to their high molecular weight, which has typically hindered further clinical development. In contrast, the bioavailability and PK profile of IAP-08 in mice was highly favorable, resembling that of irinotecan, an established chemotherapeutic drug (**Fig. 4E-F**, **Supplementary Fig. 14-15**). We attributed this favorable PK to the more lipophilic physicochemical properties of IAP-08. Specifically, the more lipophilic alkyne-containing linker may enable IAP-08 to bind to serum albumin, allowing for greater metabolic stability.

In summary, this study provides evidence that SMYD3 may be driving oncogenic phenotypes predominantly through catalytically-independent mechanisms in HPV-negative HNSCC cells and presents the first-in-class SMYD3 PROTACs. We show that enzymatic inhibition of SMYD3 is insufficient to hinder oncogenic phenotypes in HPV-negative HNSCC and that SMYD3 depletion represents the most effective therapeutic strategy to target SMYD3-driven HPV-negative HNSCC. We present IAP-08, which is highly efficacious in vitro in multiple HPV-negative HNSCC cell lines and has a favorable PK profile, providing a stepping stone for the translation of SMYD3 PROTACs in the treatment of patients with HPV-negative HNSCC, and potentially other squamous cell carcinomas of the upper aerodigestive tract [35].

### Limitations of the study

While our results support that inhibition of the enzymatic activity of SMYD3 is insufficient to repress the oncogenic phenotypes of cell proliferation, colony formation and invasion in HPV-negative HNSCC cells, the enzymatic inhibition induced by EPZ031686 and BAY-6035 (∼60-80%) was not as deep as that exerted by IAP-08 (∼80-100%); as such, further validation in HPV-negative HNSCC cell lines expressing enzymatically inactive SMYD3 will be necessary to further dissect the role of enzymatic inhibition of SMYD3 in driving oncogenic phenotypes in these cancer cells. Additionally, while IAP-08 hindered the proliferation and invasion of HPV-negative HNSCC cells at nanomolar concentrations in vitro, in vivo tolerability and efficacy studies are ongoing to determine its intratumoral penetration, pharmacodynamic effect and in vivo efficacy in xenograft mouse models. Further, as we previously published that Smyd3 depletion sensitizes syngeneic HPV-negative HNSCC mouse models to anti-PD-1 therapy [21], in vivo mouse experiments are also planned to combine IAP-08 with anti-PD-1. Importantly, XIAP and BIRC2 were found to be amongst the top 5 most degraded proteins after IAP-08 treatment of HN-SCC-151 cells, a finding that was expected as the degradation of XIAP and BIRC2 has been reported with IAP antagonism [42-44]. Given the established role of the IAP family as oncoproteins in HPV-negative HNSCC biology, further studies will be required to dissect the contribution of IAP antagonism in the antiproliferative and anti-invasive effect of IAP-08. Ongoing optimization is also aiming to improve the pharmacodynamic profile of the CRBN-based SMYD3 PROTAC compounds [45]. Moreover, genome-wide mapping studies would be necessary to investigate the impact of IAP-08 on the chromatin binding of SMYD3 and to decipher the mechanism through which SMYD3 regulates transcription. Additionally, given that SMYD3 is predominantly expressed in the cytoplasm of HPV-negative HNSCC cells, the oncogenic effects mediated through cytoplasmic SMYD3 and how these are disrupted by IAP-08 should be further investigated. Finally, it will be important to evaluate the expandability of these findings to other squamous cell carcinomas, such as lung, esophageal, bladder and cervical squamous cell carcinomas which have similar genetic backgrounds as HPV-negative HNSCC, as well as in cancer types with high frequency of SMYD3 amplification, such as breast carcinomas.

## SIGNIFICANCE

SMYD3 is a protein methyltransferase with important oncogenic roles in multiple solid tumor types, including HPV-negative HNSCC; however, inhibition of its catalytic activity imparts limited in vitro efficacy towards hindering the proliferative and invasive capacity of HPV-negative HNSCC cells. Using PROTACs, we demonstrate that degradation of SMYD3 has drastically higher in vitro potency than inhibiting its methyltransferase activity alone. This work provides the stepping stone for the translation of SMYD3 PROTACs in the clinic for patients with SMYD3-driven HPV-negative HNSCC and potentially other squamous cell carcinomas.

## AUTHOR CONTRIBUTIONS

JA: Data curation, investigation, formal analysis, methodology, validation, visualization, writing-original draft; SRD: Data curation, investigation, formal analysis, methodology, validation, visualization, writing-original draft; SK: formal analysis, investigation, methodology, software, visualization; MJ: formal analysis, investigation, methodology, software, visualization; MSD: data curation, methodology, visualization; JS: investigation, methodology; WJM: investigation, methodology; RS: conceptualization, data curation, formal analysis, investigation, methodology; VS: conceptualization, data curation, formal analysis, funding acquisition, investigation, methodology, project supervision, visualization, writing-original draft, review and editing.

## ACKNOWLEDGEMENTS

This research was supported by the Intramural Research Program of the National Institutes of Health (NIH). The contributions of the NIH author(s) were made as part of their official duties as NIH federal employees, are in compliance with agency policy requirements, and are considered Works of the United States Government. However, the findings and conclusions presented in this paper are those of the author(s) and do not necessarily reflect the views of the NIH or the U.S. Department of Health and Human Services.

## DECLARATION OF CONFLICT OF INTEREST

Drs. Sai Reddy Doda, Jawad Akthar, Rolf Swenson and Vassiliki Saloura are inventors on a patent application covering this work (PCT patent application WO2026050190 A1). The authors declare no other conflicts of interest.

## RESOURCE AVAILABILITY

### Lead contact

Further information and requests for resources and reagents should be directed to and will be fulfilled by the lead contact Vassiliki Saloura.

### Materials availability

HiBiT-tagged SMYD3 expressing cell lines generated in this study are available from the lead contact with a completed Materials Transfer Agreement.

### Data and code availability

- RNA-seq and CUT&RUN DNA-seq data have been deposited at GEO and are publicly available as of the date of publication. Accession numbers are listed in the key resources table.
- This paper does not report original code.
- All raw data reported in this paper will be shared by the lead contact upon request. Any additional information required to reanalyze the data reported in this paper is available from the lead contact upon request.

## EXPERIMENTAL MODEL AND SUBJECT DETAILS

### Cell lines

HPV-negative squamous cell carcinoma cell lines HN-6, HN13, PE/CA-PJ15 and HN-SCC-151 cells were derived from patients with locoregionally advanced HPV-negative HNSCC and were kindly provided by Dr. Tanguy Seiwert (University of Chicago). HN-6 and HN13 cells were maintained in DMEM medium with 10% fetal bovine serum, 1% penicillin/streptomycin, and 2 nM L-glutamine. PE/CA-PJ15 cells were maintained IDMEM medium with 10% fetal bovine serum, 1% penicillin/streptomycin, and 2 nM L-glutamine. HN-SCC-151 cells were maintained in DMEM/F12 medium, 10% fetal bovine serum, 1% penicillin/streptomycin and 2 nM L-glutamine. HEK293 cells were purchased from ATCC (Manassas, VA) and were cultured with EMEM plus 10% FBS. 100 µg/ml penicillin and 100 µg/ml streptomycin were added to the culture medium. Cultures were maintained at 37°C in a humidified atmosphere of 5% CO2 and 95% air.

### Generation of doxycycline-inducible HiBiT-tagged SMYD3 expressing cell line

HN13 cells were transfected with a doxycycline-inducible HiBiT-tagged SMYD3 expressing plasmid (pJB0160_iF59_SMYD3_HiBiT) and antibiotic selection with puromycin (1ug/ml) was performed. The expression of HiBiT-tagged SMYD3 upon exposure to doxycycline (1ug/ml) was confirmed by HiBiT assays and by Western blotting using a HiBiT tag antibody (**Supplementary Fig.9**).

### In vivo mouse experiments

In vivo mouse experiments were conducted at the AAALAC-accredited laboratory animal facility of Eurofins in general accordance with the “Guide for the Care and Use of Laboratory Animals: Eighth Edition” (The National Academies Press, Washington, DC, 2011). Female BALB/c nude mice were purchased from BioLASCO Taiwan Co., Ltd. (Charles River Laboratories Licensee) at 6 weeks of age. Mice were administered either intravenous (IV) or intraperitoneal (IP) doses of the SMYD3 PROTAC IAP-08. For the assessment of the pharmacokinetic (PK) profile of IAP-08, mice were dosed one time either IV=12.5mg/kg or IP=10mg/kg and 6 designated time points were evaluated with blood samples. The vehicle control consisted of 10% DMA/ 20% PG/ 40% PEG-400/ 30% of 20% HPβCD.

## METHOD DETAILS

### HiBiT assays

HiBiT assays were performed using Promega Nano-Glo^®^ HiBiT Lytic Detection System following the manufacturer’s protocol. Briefly, HiBiT tag-SMYD3 protein expressing cells were grown in a 96 well plate. After 24hrs of induction with doxycycline, the media was removed and cells were washed once with PBS. The Promega reagent was prepared using the lysing reagent and LgBiT protein and directly added to the 96 well plate to lyse the cells. The cells are incubated for 10min at room temperature, followed by luminescence reading using a plate reader. The intensity of luminescence is measured to quantify the amount of protein expressed.

### Experimental design with doxycycline-inducible HiBiT-tagged SMYD3 expressing HN13 cells

To generate a dose-response curve for the SMYD3 PROTAC IAP-08 using the HiBiT-tagged SMYD3 expressing HN13 cells, cells were seeded in 6-well plates for 24h, and next day, induction with doxycycline (1ug/ml) was initiated. After 24h, cells were trypsinized and seeded into the wells of a 96-well plate in triplicates (10,000 cells/well). Next day, treatment with a range of concentrations of the SMYD3 PROTAC (0-5uM) was initiated for 24h to 48h. Western blotting was conducted at 24h of treatment (**Fig. 3B**). Bioluminescence was captured after 4h and 24h of treatment (**Fig. 3C**), and at various time points in cells treated with 0.1uM for 24h or 48h (**Fig. 3D**).

### Quantitative bioanalysis (plasma, urine, liver homogenates) for the evaluation of the PK parameters

Blood samples were obtained in triplicates (n=3 mice) at 9 time points (0.08h, 0.17h, 0.25h, 0.5h, 1h, 2h 4h, 8h and 24h) after a single IV dose administration of the SMYD3 PROTAC IAP-08 for quantitative analysis of the PROTAC with liquid chromatography-mass spectrometry analysis. Urine samples were obtained at 0-4h, 4-8h and 8-24h, and liver homogenates were obtained 4h, 8h and 24h as a terminal collection of 3 mice for each time point.

The plasma, liver and urine homogenate samples were processed using protein precipitation and analyzed by LCMS/MS. A calibration curve with the respective same matrix of plasma, urine and liver homogenates was generated as follows; aliquots of drug-free plasma, urine and liver homogenates were spiked with the test compound at specified concentration levels. The spiked plasma, urine and liver homogenate samples were processed together with the respective samples.

The exposure levels (ng/mL) of the test compound in plasma, urine and liver homogenate were measured. Plots of plasma, urine, and liver concentrations of the test compound versus time were constructed. The fundamental PK parameters of the test compound post IV dosing were obtained from non-compartmental analysis (NCA) of the plasma data using WinNonlin (best-fit mode). The exposure levels of urine/plasma and liver/plasma ratios were determined. PK parameters calculated included t_1/2_, C_0_, AUC_last_, AUC_Inf_, AUC/D, AUC Extr, MRT, Vss and CL.

### NanoBRET Target Engagement Intracellular SMYD3 Assays

The NanoBRET target engagement intracellular SMYD3 assays were developed and conducted at Reaction Biology Corporation (RBC). The SMYD3_pNLF1-N [CMV_Hygro] fusion vector was generated at RBC. The NanoBRET SMYD3 tracer was produced by the NCI. The FuGENE HD transfection reagent (Cat#E2311), the transfection carrier DNA (Cat# E4881), the NanoBRET HaloTag 618 Ligand (Cat# G9801), the NanoBRET Nano-Glo substrate (Cat# N1571), and the tracer dilution buffer (Cat# N2191) were purchased from Promega. HEK293 cells were transiently transfected with the SMYD3_pNLF1-N [CMV_Hygro] vector using the FuGENE HD transfection reagent. Incremental concentrations of IAP-08 and the proteasome inhibitor MG132 (10uM) were delivered into a 384 well assay plate by Echo 550 liquid handler (Labcyte Inc, Sunnyvale, CA). 24 hours post transfection, the transfected cells were harvested, resuspended in Opti-MEM I Reduced Serum Medium, mixed with cell-permeable NanoBRET SMYD3 tracer and dispensed into 384 well plates. Plates were incubated at 37°C in 5% CO2 cell culture incubator for 2 hours. NanoBRET® Nano-Glo® substrate plus extracellular NanoLuc® inhibitor solution (Cat# N2162) was added into the wells of the assay plate and incubated for 20 minutes at room temperature. The donor emission wavelength (460nm) and acceptor emission wavelength (600nm) were measured on Envision 2104 Multilabel Reader (PerkinElmer, Santa Clara, CA). The BRET ratio was calculated as follows: BRET ratio = [(Acceptor sample)/(Donor sample)] – [(Acceptor no-tracer control)/(Donor no tracer control)]. The BRET ratio of the background was subtracted from each control and treatment sample, and the normalized BRET Response (%) was calculated as follows: (BRET ratio of each treatment sample/Average BRET ratio of DMSO control samples)*100%. The IC50 curves were plotted and IC50 values were calculated using the GraphPad Prism program based on a sigmoidal dose-response equation.

### NanoBRET Cellular Ternary Complex Formation Assays

The NanoBRET cellular ternary complex formation assays were developed and conducted at Reaction Biology Corporation (RBC). SMYD3-NanoLuc fusion vector and XIAP-HaloTag fusion vector were generated at RBC. FuGENE HD transfection reagent (Cat#E2311), transfection carrier DNA (Cat# E4881), NanoBRET HaloTag 618 Ligand (Cat# G9801), and NanoBRET Nano-Glo substrate (Cat# N1571) were purchased from Promega. HEK293 were seeded in 6-well plate and co-transfected with plasmids encoding SMYD3-NanoLuc fused target protein and XIAP-HaloTag E3 ligase using FuGENE HD transfection reagent. 24 hours post transfection, cells were harvested and resuspended in culture medium (Opti-MEM I Reduced Serum Medium +4% FBS), then were dispensed into white 384-well plates in the presence or absence of the HaloTag NanoBRET 618 Ligand, and incubated overnight. The IAP-08 compound was supplied by NCI as a powdered stock, dissolved in DMSO at 10 mM, and was added at serial concentrations along with a vehicle control. Plates were incubated for 2 hours at 37 °C. The Nano-Glo substrate diluted in assay buffer (Opti-MEM I Reduced Serum Medium) was added to each well, the plates were incubated for 5 minutes. The BRET signal was measured and the normalized BRET Response (%) was calculated as described above.

### CCK8 assays

HNSCC cells were plated overnight in 24- or 96-well plates and on the next day, they were treated either with the SMYD3 PROTAC EPZ031686-IAP-08 or the SMYD3 inhibitor EPZ031686. The number of viable cells was measured using the Cell Counting Kit-8 (Dojindo, Kumamoto, Japan) at the indicated time points.

### Colony formation assays (CFAs)

HNSCC cells were plated overnight in 6-well plates (500 cells/well) and on the next day, they were treated either with the SMYD3 PROTAC IAP-08 or the SMYD3 inhibitor EPZ031686. Once colonies were visible, they were stained with 0.01% (w/v) crystal violet (Sigma-Adrich, cat # HT901-8FOZ), washed with dH20 to remove excess stain, and left to dry. Colonies were counted using the ImageJ software (version 1.53k).

### Transwell migration and invasion assays

For the invasion assays, 50 μL of Matrigel (Corning™, cat # 354234) were placed on top of the transwell inserts and was solidified in a 37 °C incubator for 15-30 minutes to form a thin gel layer before seeding 80,000 cells/100uL DMEM. Following the same 24-hour incubation period at 37°C, the cells that invaded through the Matrigel and transwell pores were similarly fixed and stained with crystal violet, then visualized and counted using ImageJ. The assays were performed in biological triplicates and were replicated twice.

### Western blotting

Nuclear extracts were prepared from cultured cell lines using the Nuclear Complex Co-IP kit (Active Motif) and 10-15 μg were loaded to examine protein levels of SMYD3 or HiBiT-tagged SMYD3, with histone H3 used as a loading control. Primary antibodies used were anti-SMYD3 (ab187149, Abcam, dilution 1:2000), anti-SMYD3 (ab187149, Abcam, dilution 1:2000), anti-ubiqutin (Sc-8017, Santa-Cruz, dilution 1:1000), anti-HiBiT (N7200, Promega, dilution 1:2000), anti-H3K4me3 (ab8580, abcam, dilution 1:2000), anti-p-ERK (9101, cell signaling, dilution 1:500), anti-ERK (sc-1647, santa-Cruz, dilution 1:1000) anti-UHRF1 (D6G8E, Cell Signaling, dilution 1:1000), anti-PARP (46D11, Cell Signaling, Dilution 1:500), anti-capsase-3 (D175, Cell Signaling, Dilution 1:500), anti-histone 3 (ab1791, Abcam, dilution 1:125000), anti-GAPDH (Sc-5174, Santa-Cruz, dilution 1:2000), anti-actin (A2228, SIGMA, Dilution 1:2000) antibodies. Amersham ECL prime Western Blotting Detection Reagent (Cytiva, cat # 45-002-401) or Pierce ECL Western Blotting Substrate (Thermo Scientific, cat# 32106) were used as detection reagents. Blots were imaged using a Chemi-fluorescent Odyssey FC machine after applying a detection reagent. Densitometry of all blots was performed using ImageJ software (1.53k, NIH, Bethesda, MD).

### In vitro metabolic assays for IAP-08

For the evaluation of the plasma protein binding capacity of IAP-08, equilibrium dialysis was conducted using human or mouse plasma and the compound at a concentration of 1e-05M. Duplicate samples were incubated for 4h at 37°C and high performance liquid chromatography(HPLC)-MS/MS analysis was pursued to quantify the unbound compound. The peak areas of the test compound in the buffer and test samples were used to calculate % binding and recovery according to the following formula:

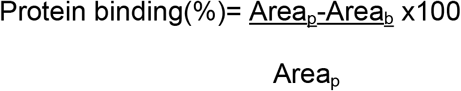

To assess the liver metabolic profile of IAP-08, liver microsome intrinsic clearance assays, which assess oxidation by cytochrome P450, were conducted by incubating the compound at a concentration of 1e-07M with liver microsomes and NAPDH at 37°C for various time periods (0, 15min, 30min, 45min, 60min). The compound was quantified using HPLC-MS/MS. The metabolic stability was expressed as % of the parent compound remaining was calculated by comparing the peak area of the compound at the time point relative to that at time 0. These assays were conducted by Eurofins Discovery.

### RNA-sequencing

RNA-seq was performed in HN-SCC-151 cells. The cells were seeded in a 10cm dish, allowed to settle for 24hrs, and then treated with either 10uM of the SMYD3 inhibitor EPZ031686 or 0.5uM of the SMYD3 PROTAC IAP-08 for 48h.

Following completion of incubation, cells were trypsinized, washed twice with PBS, centrifuged and processed for RNA extraction (Direct-zol RNA miniprep kit, Zymo Research). Three biological replicates for each sample were processed to extract RNA, quantified using Qubit and sequenced. Samples were pooled and sequenced on NextSeq or NovaSeq using the Illumina TruSeq Stranded mRNA Library Prep kit (Illumina) and paired-end sequencing. The samples have 68 to 130 million pass filter reads. Reads of the samples were trimmed for adapters and low-quality bases using Cutadapt before alignment with the reference genome (hg38) and the annotated transcripts using STAR. The samples had 62-75% non-duplicate reads. In addition, the gene expression quantification analysis was performed for all samples using STAR/RSEM tools. The raw counts are provided as part of the data delivery.

### Tandem Mass Tag (TMT) labeling, fractionation and data analysis

#### Protein Digestion and TMT labeling

The cell pellets were lysed in EasyPrep Lysis buffer (Thermo Fisher, CA) according to manufacturer’s protocol. Lysates were clarified by centrifugation and protein concentration was quantified using BCA protein estimation kit (Thermo Fisher, CA). Fifteen micrograms of lysate were reduced, alkylated and digested by addition of trypsin at a ratio of 1:50 (Promega) and incubating overnight at 37°C. For TMT labeling 100mg of TMTpro label (Thermo Fisher, CA) in 100% ACN was added to each sample. After incubating the mixture for 1 hr at room temperature with occasional mixing, the reaction was terminated by adding 50 ml of 5% hydroxylamine, 20% formic acid. The peptide samples for each condition were pooled and peptide clean-up was performed using the proprietary peptide clean up columns from the EasyPEP Mini MS Sample Prep kit (Thermo Fisher, CA).

#### High pH reverse phase fractionation

The first dimensional separation of the peptides was performed using a Waters Acquity UPLC system coupled with a fluorescence detector (Waters, Milford, MA) using a 150mm x 3.0mm Xbridge Peptide BEM^TM^ 2. 5 mm C18 column (Waters, MA) operating at 0.35 ml/min. The dried peptides were reconstituted in 100 ul of mobile phase A solvent (10 mM Ammonium Formate, pH 9.4). Mobile phase B was 10 mM Ammonium Formate /90% acetonitrile, pH9.4. The column was washed with mobile phase A for 5 min followed by gradient elution 10-50% B (5-60 min) and 50-75 %B (60-70 min). The fractions were collected every minute. These 60 fractions were pooled into 24 fractions. The fractions were vacuum centrifuged to dryness and stored at -80°C until analysis by mass spectrometry.

#### Mass Spectrometry acquisition and data analysis

The dried peptide fractions were reconstituted in 0.1%TFA and subjected to nanoflow liquid chromatography (Thermo Ultimate^TM^ 3000RSLC nano LC system, Thermo Scientific) coupled to an Orbitrap Eclipse mass spectrometer (Thermo Scientific, CA). Peptides were separated using a low pH gradient using a 5-50% ACN over 120 minutes in mobile phase containing 0.1% formic acid at 300 nl/min flow rate. MS scans were performed in the Orbitrap analyser at a resolution of 120,000 with an ion accumulation target set at 4e^5^ and max IT set at 50ms over a mass range of 400-1600 m/z. Ions with determined charge states between 2 and 5 were selected for MS2 scans. A cycle time of 3 sec was used, and a quadrupole isolation window of 0.4 m/z was used for MS/MS analysis. An Orbitrap at 15,000 resolution with a normalized AGC set at 250 followed by maximum injection time set as “Auto” with a normalized collision energy setting of 38 was used for MS/MS analysis. The node “Turbo TMT” was switched on for high resolution acquisition of TMT reporter ions.

Acquired MS/MS spectra were searched against a human uniprot protein database using a SEQUEST HT and percolator validator algorithms in the Proteome Discoverer 2.4 software (Thermo Scientific, CA). The precursor ion tolerance was set at 10 ppm and the fragment ions tolerance was set at 0.02 Da along with methionine oxidation included as dynamic modification. Carbamidomethylation of cysteine residues and TMT16 plex (304.2071Da) was set as a static modification of lysine and the N-termini of the peptide. Trypsin was specified as the proteolytic enzyme, with up to 2 missed cleavage sites allowed. Searches used a reverse sequence decoy strategy to control for the false peptide discovery and identifications were validated using percolator software. Only peptides with less the 50% co-isolation interference were used for quantitative analysis.

Reporter ion intensities were adjusted to correct for the impurities according to the manufacturer’s specification and the abundances of the proteins were quantified using the summation of the reporter ions for all identified peptides. The reporter abundances were normalized across all the channels to account for equal peptide loading. Data analysis and visualization were performed in Microsoft Excel or R.

## QUANTIFICATION AND STATISTICAL ANALYSIS

### RNA-seq Heatmaps

RNA-Seq data were quantitated to obtain raw tag counts at the gene level using either HTSeq or featureCount. The raw tag count data was variance stabilizing transformed using VST function in DESeq2 R library, and z-score of the transformed data was obtained to color code for heatmap. For clustered heatmaps, pheatmap R library was used with Euclidean distance and ward.D2 clustering options. Significance of gene expression changes was evaluated using DESeq2 R library and determined based on Wald-statistics (FDR<0.1) and shrunken log2 fold-change (>log2(1.3), <-log2(1.3)) using ahsr method available from DESeq2 library.

### Volcano plots

For all volcano plots, EnhancedVolcano R library was used.

### Mass spectrometry analysis

Two tail t-test was performed considering equal variance on both treated and control samples.

### Chemical synthesis and characterization

NMR data are provided in **Supplementary Table 9**.

### General information for chemical synthesis

General information for chemical synthesis Starting materials were used as received unless otherwise noted. All moisture sensitive reactions were performed in an inert atmosphere of argon with oven dried glassware. Reagent grade solvents were used for extractions and flash chromatography. Reaction progress was monitored by LC-MS analysis performed on an Agilent UPLC/MS instrument equipped with a RP-C18 column (Poroshell 120 SB-C18, 4.6 350mm,2.7mmorZorbax300SB-C18,4.6350mm,3.5mm),dualatmosphericpressurechemical ionization (APCI)/electrospray (ESI) mass spectrometry detector, and photodiode array detector. Flash chromatography was performed by using a RediSepRf NP-silica (40–63 mm60A °) or a Teledyne RediSepRf Gold RP-C18 column (20–40 mm 100 A °) in a Teledyne ISCO CombiFlash Rf200 purification system unless otherwise specified.^1^H NMR spectra were recorded on an Agilent 400MHz spectrometer and are reported in parts per million (ppm) on the d scale relative to CDCl_3_ (d 7.26) and DMSO-d6 (d 2.50) as internal standards. Data are reported as follows: chemical shift, multiplicity (s = singlet, d = doublet, t = triplet, q = quartet, b = broad, m = multiplet), coupling constants(Hz), andintegration. ^13^C-NMR spectra were recorded on an Agilent 100 MHz and are reported in parts per million (ppm) on the d scale relative to CDCl_3_ (d 77.00), CD_3_OD and DMSO-d6 (d 39.52).

### Synthesis and characterization data

#### Schema S1: Synthesis of common intermediate (S7)

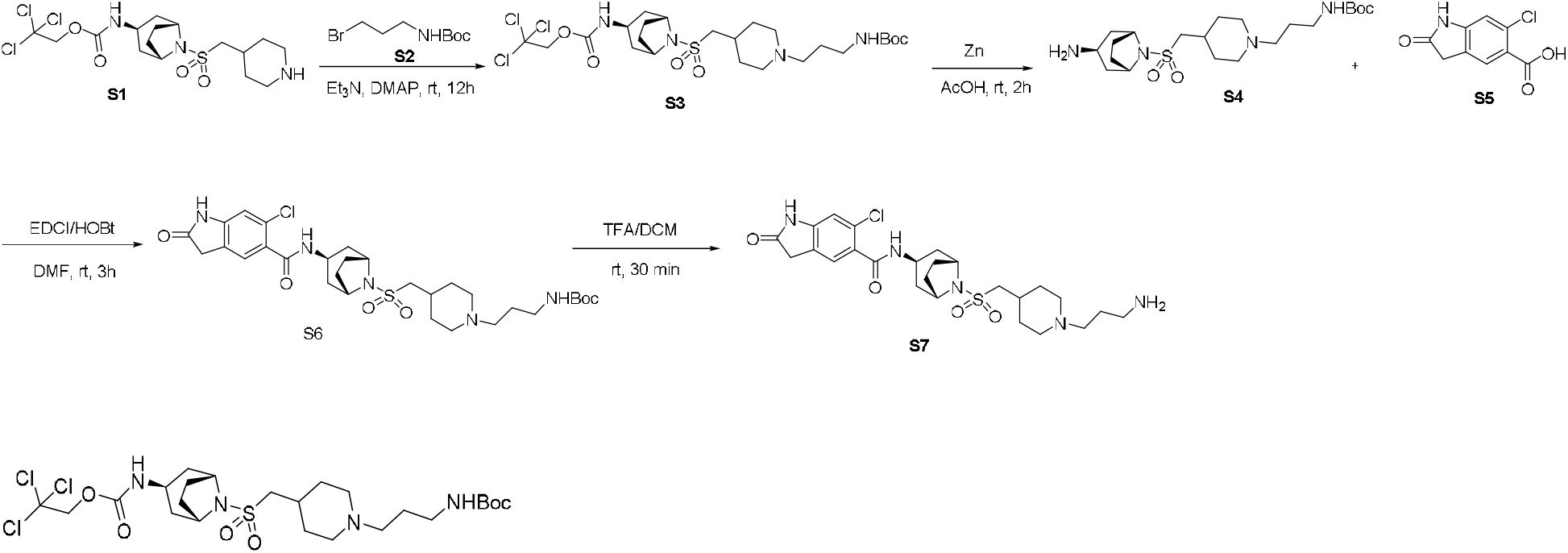

2,2,2-Trichloroethyl ((*1R,3r,5S*)-8-(((1-(3-((*tert*-butoxycarbonyl)amino)propyl)piperidin-4-yl)methyl)sulfonyl)-8-azabicyclo[3.2.1]octan-3-yl)carbamate **(S3):** To a stirred solution of benzyl 4-((((*1R,3r,5S*)-3-(((2,2,2-trichloroethoxy)carbonyl)amino)-8-azabicyclo[3.2.1]octan-8-yl)sulfonyl)methyl)piperidine-1-carboxylate (1.2 g, 2.01 mmol) in DCM (5 mL), and acetonitrile (20 mL), were added sequentially *tert*-butyl (3-bromopropyl)carbamate (957.4 mg, 4.021 mmol), Et_3_N (0.84 mL, 6.031 mmol) and DMAP (24.56 mg, 0.2 mmol) at 0 °C. The resulting mixture was stirred at room temperature for overnight. After the completion of the reaction (the progress of the reaction was monitored by LCMS) the solution was concentrated and purified by flash chromatography on silica gel (5% MeOH in DCM) to give of title compound 0.95 g (76%) as a white solid. ^1^H NMR (400 MHz, CDCl_3_) δ 5.38 (s, 1H), 5.27 (d, *J* = 6.4 Hz, 1H), 4.70 (s, 2H), 4.22 (dq, *J* = 5.6, 3.0 Hz, 2H), 3.93 (q, *J* = 6.7 Hz, 1H), 3.15 (q, *J* = 6.3 Hz, 2H), 2.89 (dd, *J* = 6.5, 4.3 Hz, 4H), 2.37 (t, *J* = 6.8 Hz, 2H), 2.30 – 2.08 (m, 4H), 2.02 – 1.85 (m, 9H), 1.63 (p, *J* = 6.7 Hz, 2H), 1.41 (s, 9H); ^13^C NMR (101 MHz, cdcl_3_) δ 156.1, 153.8, 95.5, 78.8, 77.4, 77.2, 77.0, 76.7, 74.4, 59.4, 56.9, 55.6, 53.3, 43.4, 39.8, 37.5, 32.1, 32.0, 28.9, 28.4, 26.5; LC-MS (ESI): (m/z) = 619 [M+H].

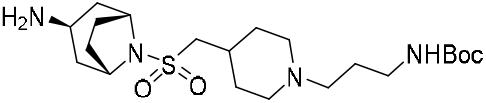

*tert*-Butyl(3-(4-((((*1R,3r,5S*)-3-amino-8-azabicyclo[3.2.1]octan-8-yl)sulfonyl)methyl)piperidin-1-yl)propyl)carbamate (S**4**): To a stirred solution of 2,2,2-trichloroethyl 2,2,2-trichloroethyl ((*1R,3r,5S*)-8-(((1-(4,4,4-trifluorobutyl)piperidin-4-yl)methyl)sulfonyl)-8-azabicyclo[3.2.1]octan-3-yl)carbamate (850 mg, 1.48 mmol) in glacial acetic acid (15 mL), was added zinc (970 mg, 10 eq, 14.8 mmol) at room temperature, and the resulting heterogenous solution was stirred at room temperature for the 2 hrs. After completion of the reaction (as indicated by LCMS), the glacial acetic acid was removed under vacuum and the crude residue was neutralized with saturated NaHCO_3_ solution (10 mL). The resulting solution was extracted with DCM (50 mL x 5), and the combined organics were dried over anhydrous Na_2_SO_4_ and filtered. The solution was concentrated under vacuum to get crude product (**4**), which was used without further purification. LC-MS (ESI): (m/z) = 445 [M+H].

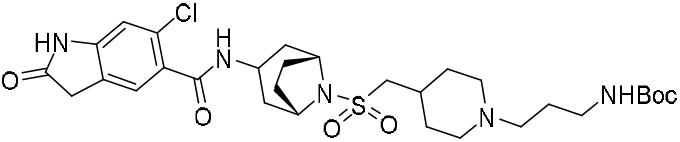

*tert*-Butyl-(3-(4-((((1R,5S)-3-(6-chloro-2-oxoindoline-5-carboxamido)-8-azabicyclo[3.2.1]octan-8-yl)sulfonyl)methyl)piperidin-1-yl)propyl)carbamate **(S6)**: To a stirred solution of (*1R,3r,5S*)-8-(((1-(4,4,4-trifluorobutyl)piperidin-4-yl)methyl)sulfonyl)-8-azabicyclo[3.2.1]octan-3-amine (700 mg, 1.76 mmol) in DMF (10 mL), were added 6-chloro-2-oxoindoline-5-carboxylic acid (559 mg, 2.64 mmol), EDCI (675 mg, 3.52 mmol), HOBt (539 mg, 3.52 mmol) and Et_3_N (982 μL, 7.04 mmol) at 0 °C. The resulting mixture was stirred at room temperature for 3h. After completion of the reaction (as indicated by LCMS), the volatiles were removed under vacuum and the crude material was purified by reverse phase chromatography without workup using a gradient of 10-55% acetonitrile in water with the addition of 0.05% trifluoroacetic acid (TFA) to give title compound 550 mg (55%) as white solid. ^1^H NMR (400 MHz, CD_3_OD-d_4_) δ 11.42 (s, 1H), 9.01 (d, *J* = 4.6 Hz, 1H), 8.02 (s, 1H), 7.75 (t, *J* = 5.8 Hz, 1H), 7.63 (s, 1H), 4.91 (s, 2H), 4.76 (d, *J* = 6.1 Hz, 1H), 4.43-4.04 (m, 4H), 3.90 (d, *J* = 6.1 Hz, 2H), 3.76 (q, *J* = 7.0 Hz, 6H), 2.89 (dt, *J* = 29.8, 10.8 Hz, 6H), 2.73 (d, *J* = 14.6 Hz, 4H), 2.55 (q, *J* = 7.7 Hz, 2H), 2.30 (t, *J* = 12.4 Hz, 2H), 2.18 (s, 9H); ^13^C NMR (101 MHz, DMSO-d6) δ 176.8, 166.9, 156.2, 146.0, 130.1, 129.5, 125.3, 125.1, 110.1, 78.3, 56.6, 55.4, 54.5, 51.9, 48.2, 42.0, 37.6, 36.6, 35.7, 30.1, 29.0, 28.7, 28.3, 24.7; LC-MS (ESI): (m/z) = 637 [M+H].

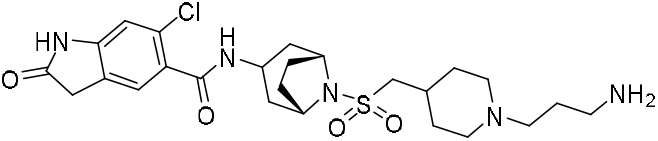

N-((1*R*,5S)-8-(((1-(3-Aminopropyl)piperidin-4-yl)methyl)sulfonyl)-8-azabicyclo[3.2.1]octan-3-yl)-6-chloro-2-oxoindoline-5-carboxamide **(S7):** The compound *tert*-butyl (3-(4-((((1*R*,5*S*)-3-(6-chloro-2-oxoindoline-5-carboxamido)-8-azabicyclo[3.2.1]octan-8-yl)sulfonyl)methyl)piperidin-1-yl)propyl)carbamate (500 mg, 0.78 mmol) was dissolved in 20% TFA in DCM (10 ml) and stirred for 1h at room temperature, After the completion of the reaction (as indicated by LCMS), the solution was concentrated and dried to give the title product, which was used for next step without further purification (**7**). LC-MS (ESI): (m/z) = 537 [M+H].

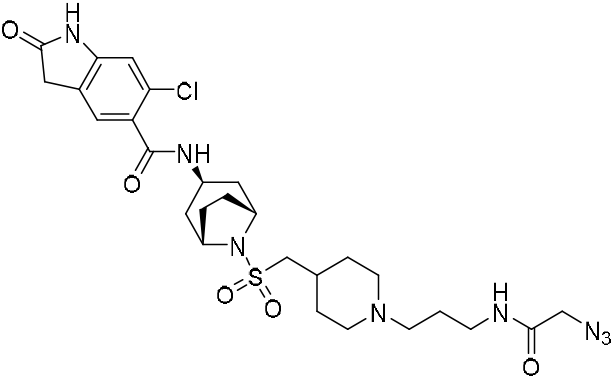

N-((*1R,3r,5S*)-8-(((1-(3-(2-Azidoacetamido)propyl)piperidin-4-yl)methyl)sulfonyl)-8-azabicyclo[3.2.1]octan-3-yl)-6-chloro-2-oxoindoline-5-carboxamide **(S8):** The compound N-((*1R,3r,5S*)-8-(((1-(3-aminopropyl)piperidin-4-yl)methyl)sulfonyl)-8-azabicyclo[3.2.1]octan-3-yl)-6-chloro-2-oxoindoline-5-carboxamide **7** (100 mg, 1 eq, 0.186 mmol) was dissolved in DMF (2 mL), were added 2,5-dioxopyrrolidin-1-yl 2-azidoacetate (73.6 mg, 2 eq, 0.372 mmol) and DIPEA (60.0 mg, 80.9 μL, 2.5 eq, 0.465 mmol) at 0 °C. After completion of the reaction (as indicated by LCMS), the volatiles were removed under vacuum and the crude material was purified by reverse phase chromatography without workup using a gradient of 10-65% acetonitrile in water with the addition of 0.05% TFA to give title compound 65 mg (56%) as white solid. ^1^H NMR (400 MHz, DMSO-d_6_) δ 10.61 (s, 1H), 9.27 (s, 1H), 8.26 (t, *J* = 5.7 Hz, 1H), 8.20 (d, *J* = 4.7 Hz, 1H), 7.21 (s, 1H), 6.82 (s, 1H), 4.11 (d, *J* = 5.6 Hz, 2H), 3.96 (d, *J* = 10.0 Hz, 1H), 3.82 (s, 2H), 3.46 (d, *J* = 23.9 Hz, 9H), 3.12 (dt, *J* = 18.5, 6.1 Hz, 5H), 3.00 – 2.90 (m, 5H), 2.57 (s, 1H), 2.21 – 1.71 (m, 17H), 1.49 (q, *J* = 13.2 Hz, 2H); ^13^C NMR (101 MHz, DMSO) δ 176.8, 173.2, 168.0, 166.9, 146.0, 130.1, 129.5, 125.3, 125.1, 110.1, 56.6, 55.4, 54.4, 51.9, 51.3, 42.0, 36.6, 36.4, 35.7, 30.1, 29.0, 28.3, 25.7, 24.2; LC-MS (ESI): (m/z) = 621 [M+H].

#### Coupling of liner to E3 ligase ligand

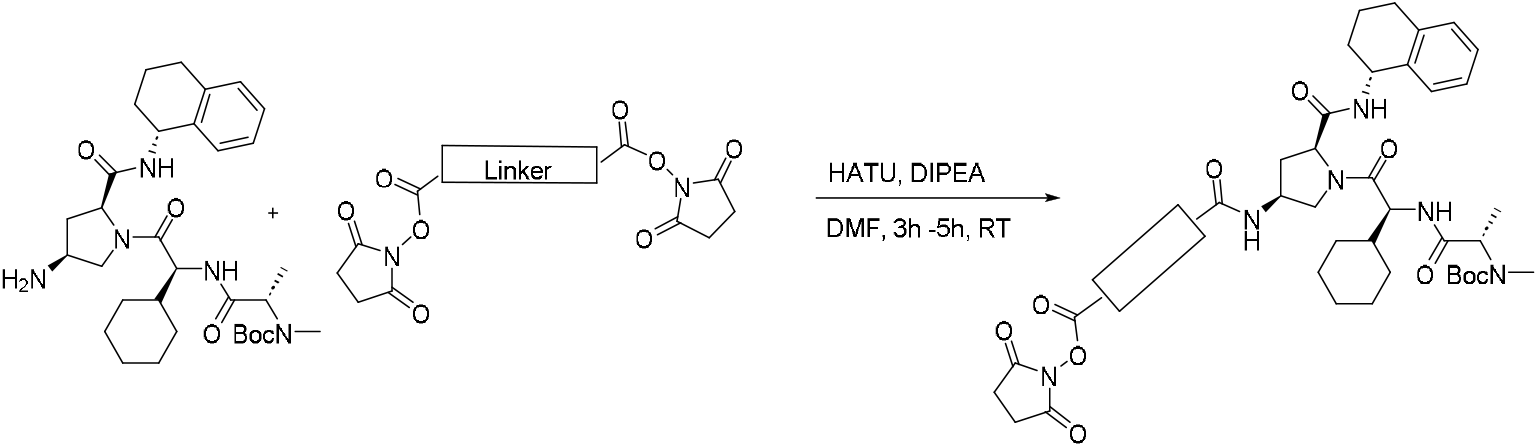

##### General Method 1A

To a stirred solution of the *tert*-butyl ((*S*)-1-(((*S*)-2-((2*S*,4*S*)-4-amino-2-(((*R*)-1,2,3,4-tetrahydronaphthalen-1-yl)carbamoyl)pyrrolidin-1-yl)-1-cyclohexyl-2-oxoethyl)amino)-1-oxopropan-2-yl)(methyl)carbamate hydrochloric salt (1.0 eq) in DMF (2 mL), were sequentially added bis(2,5-dioxopyrrolidin-1-yl) decanedioate (2 eq) and DIPEA ( 2 eq) at were added 0 °C. The resulting mixture was stirred at room temperature for 1h. After completion of the reaction (as indicated by LCMS) the volatiles were removed under vacuum and the crude material was purified by reverse phase chromatography without workup to give corresponding product.

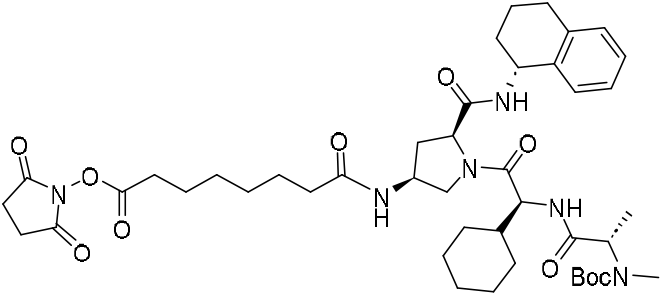

2,5-Dioxopyrrolidin-1-yl-8-(((3*S*,5*S*)-1-((S)-2-((S)-2-((*tert*-butoxycarbonyl)(methyl)amino)propanamido)-2-cyclohexylacetyl)-5-(((*R*)-1,2,3,4-tetrahydronaphthalen-1-yl)carbamoyl)pyrrolidin-3-yl)amino)-8-oxooctanoate **(S9a)**: General procedure 1A was followed on 150 mg (0.25 mmol) scale of the ((*S*)-1-(((*S*)-2-((2*S*,4*S*)-4-amino-2-(((R)-1,2,3,4-tetrahydronaphthalen-1-yl)carbamoyl)pyrrolidin-1-yl)-1-cyclohexyl-2-oxoethyl)amino)-1-oxopropan-2-yl)(methyl)carbamate hydrochloric salt. The crude material was purified by reverse phase chromatography without workup using a gradient of 10-85% acetonitrile in water with the addition of 0.05% TFA to afford 120 mg (66%) of title compound. ^1^H NMR (400 MHz, DMSO-d_6_) δ 8.37 (d, *J* = 8.6 Hz, 1H), 8.10 (d, *J* = 7.6 Hz, 1H), 7.29 (d, *J* = 7.5 Hz, 1H), 7.17 – 7.03 (m, 3H), 4.96 – 4.86 (m, 1H), 4.35 – 4.18 (m, 3H), 4.03 (dd, *J* = 9.9, 6.8 Hz, 1H), 3.29 (dd, *J* = 9.8, 7.6 Hz, 1H), 2.71 (d, *J* = 8.3 Hz, 5H), 2.63 (t, *J* = 7.2 Hz, 2H), 2.35 (dt, *J* = 12.7, 7.6 Hz, 1H), 2.03 (t, *J* = 7.4 Hz, 2H), 1.90 – 1.54 (m, 8H), 1.48 (p, *J* = 7.3 Hz, 2H), 1.39 (d, *J* = 11.2 Hz, 11H), 1.30 – 0.83 (m, 8H); ^13^C NMR (101 MHz, DMSO-d6) δ 172.3, 171.3, 170.7, 170.2, 169.4, 137.8, 137.4, 129.0, 128.8, 127.1, 126.1, 79.4, 58.9, 55.3, 52.7, 48.1, 47.1, 35.8, 34.9, 30.6, 30.5, 30.3, 29.2, 29.1, 28.6, 28.5, 28.5, 28.2, 26.3, 26.2, 26.0, 25.9, 25.3, 24.6, 20.7; LC-MS (ESI): (m/z) = 837 [M+H].

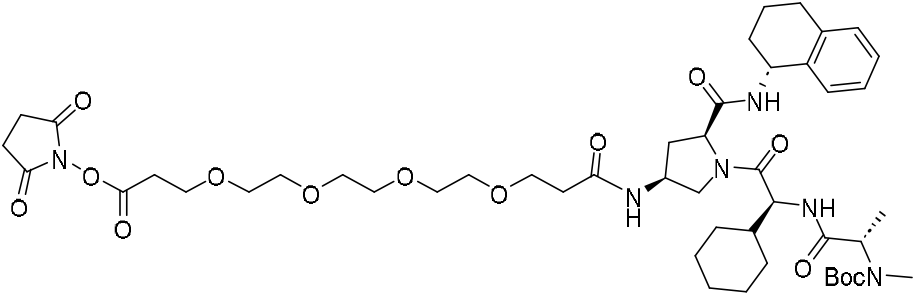

2,5-dioxopyrrolidin-1-yl 16-(((3*S*,5*S*)-1-((*S*)-2-((*S*)-2-((*tert*-butoxycarbonyl)(methyl)amino)propanamido)-2-cyclohexylacetyl)-5-(((*R*)-1,2,3,4-tetrahydronaphthalen-1-yl)carbamoyl)pyrrolidin-3-yl)amino)-16-oxo-4,7,10,13-tetraoxahexadecanoate **(S9b):** General procedure 1A was followed on 100 mg (0.17 mmol) scale of the ((*S*)-1-(((*S*)-2-((2*S*,4*S*)-4-amino-2-(((*R*)-1,2,3,4-tetrahydronaphthalen-1-yl)carbamoyl)pyrrolidin-1-yl)-1-cyclohexyl-2-oxoethyl)amino)-1-oxopropan-2-yl)(methyl)carbamate hydrochloric salt. After the completion of the reaction, crude material was purified by reverse phase chromatography without workup using a gradient of 10-85% acetonitrile in water with the addition of 0.05% TFA to afford 110 mg (67%) of title compound. ^1^H NMR (500 MHz, CDCl_3_) δ 8.18 (s, 1H), 7.50 (d, *J* = 8.3 Hz, 1H), 7.06 (t, *J* = 6.8 Hz, 2H), 7.00 (dd, *J* = 12.7, 7.3 Hz, 2H), 5.01 (q, *J* = 6.0 Hz, 1H), 4.75 – 4.44 (m, 2H), 4.42 – 4.19 (m, 5H), 3.99 (dd, *J* = 10.9, 5.3 Hz, 1H), 3.85 – 3.64 (m, 4H), 3.57 (d, *J* = 6.2 Hz, 13H), 2.81 (t, *J* = 6.3 Hz, 2H), 2.77 – 2.63 (m, 9H), 2.46 (q, *J* = 6.2 Hz, 2H), 2.27 (d, *J* = 13.8 Hz, 1H), 2.15 (ddd, *J* = 14.4, 8.9, 6.5 Hz, 1H), 1.93 (td, *J* = 10.4, 5.3 Hz, 1H), 1.75 (h, *J* = 6.5 Hz, 3H), 1.66 – 1.42 (m, 5H), 1.38 (s, 9H), 1.20 (d, *J* = 7.1 Hz, 3H), 1.01 (t, *J* = 11.6 Hz, 3H), 0.94 – 0.74 (m, 2H); ^13^C NMR (126 MHz, CDCl_3_) δ 172.7, 171.7, 170.9, 169.1, 166.8, 137.3, 136.0, 129.2, 128.3, 127.4, 126.1, 70.6, 70.6, 70.5, 70.4, 70.4, 70.3, 66.9, 65.7, 60.1, 55.4, 55.3, 49.5, 48.0, 36.9, 32.1, 31.6, 29.9, 29.2, 29.1, 28.5, 28.3, 25.9, 25.7, 25.6, 20.0; LC-MS (ESI): (m/z) = 957 [M+H].

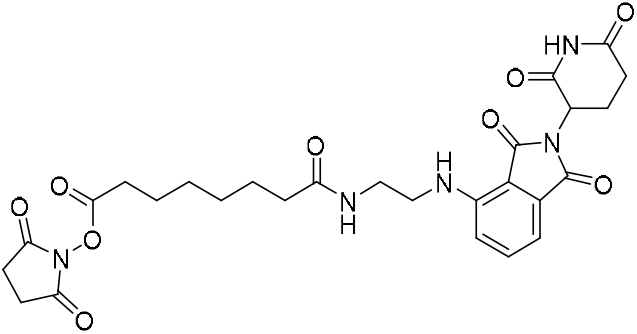

2,5-dioxopyrrolidin-1-yl-8-((2-((2-(2,6-dioxopiperidin-3-yl)-1,3-dioxoisoindolin-4-yl)amino)ethyl)amino)-8-oxooctanoate **(S9c):** General procedure 1A was followed on 100 mg (0.316 mmol) scale of the 4-((2-aminoethyl)amino)-2-(2,6-dioxopiperidin-3-yl)isoindoline-1,3-dione. After the completion of the reaction, crude material was purified by reverse phase chromatography without workup using a gradient of 10-85% acetonitrile in water with the addition of 0.05% TFA to afford 79 mg (44%) of title compound as pale yellow solid. ^1^H NMR (500 MHz, CDCl_3_) δ 8.60 (s, 1H), 7.43 (dd, *J* = 8.5, 7.2 Hz, 1H), 7.02 (d, *J* = 7.1 Hz, 1H), 6.91 (d, *J* = 8.6 Hz, 1H), 6.33 (q, *J* = 5.9 Hz, 2H), 4.86 (dd, *J* = 11.9, 5.4 Hz, 1H), 3.39 (q, *J* = 4.7 Hz, 4H), 2.92 – 2.63 (m, 7H), 2.58 – 2.46 (m, 3H), 2.12 – 2.00 (m, 3H), 1.59 (dp, *J* = 40.3, 7.3 Hz, 4H), 1.42 – 1.20 (m, 5H); ^13^C NMR (126 MHz, CDCl_3_) δ 174.0, 171.6, 169.6, 169.5, 168.9, 168.7, 167.6, 146.8, 136.3, 132.5, 116.8, 111.9, 110.2, 48.9, 42.1, 39.0, 36.0, 31.4, 30.9, 28.3, 28.0, 25.6, 25.1, 24.3, 22.7; LC-MS (ESI): (m/z) = 570 [M+H].

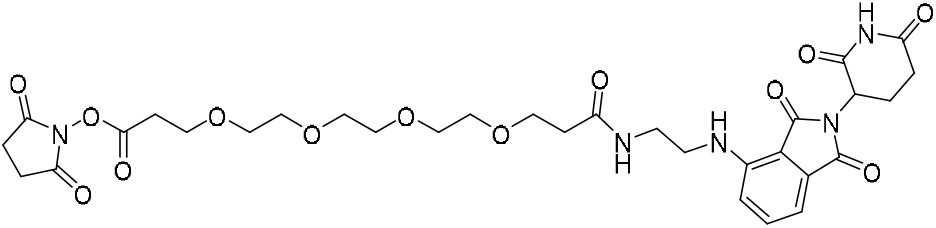

2,5-dioxopyrrolidin-1-yl 1-((2-(2,6-dioxopiperidin-3-yl)-1,3-dioxoisoindolin-4-yl)amino)-4-oxo-7,10,13,16-tetraoxa-3-azanonadecan-19-oate **(S9d):** General procedure 1A was followed on 200 mg (0.316 mmol) scale of the 4-((2-aminoethyl)amino)-2-(2,6-dioxopiperidin-3-yl)isoindoline-1,3-dione. After the completion of the reaction, crude material was purified by reverse phase chromatography without workup using a gradient of 10-85% acetonitrile in water with the addition of 0.05% TFA to afford 130 mg (59%) of title compound as pale yellow solid. ^1^H NMR (500 MHz, CDCl_3_) δ 8.99 (s, 1H), 7.42 (t, *J* = 7.8 Hz, 1H), 7.08 (d, *J* = 5.7 Hz, 1H), 6.98 (dd, *J* = 17.7, 7.9 Hz, 2H), 6.40 (d, *J* = 5.9 Hz, 1H), 4.89 – 4.81 (m, 1H), 3.74 (t, *J* = 6.3 Hz, 2H), 3.62 (t, *J* = 5.7 Hz, 2H), 3.58 – 3.48 (m, 13H), 3.38 (q, *J* = 2.9 Hz, 4H), 2.84 – 2.65 (m, 10H), 2.39 (t, *J* = 5.7 Hz, 2H); ^13^C NMR (126 MHz, CDCl_3_) δ 172.6, 171.8, 169.3, 169.3, 168.9, 167.6, 166.8, 146.8, 136.2, 132.4, 116.9, 111.6, 110.1, 70.6, 70.5, 70.4, 70.3, 70.1, 70.1, 67.0, 65.7, 53.5, 48.9, 42.0, 38.8, 36.7, 32.1, 31.4, 25.6, 22.7; LC-MS (ESI): (m/z) = 690 [M+H].

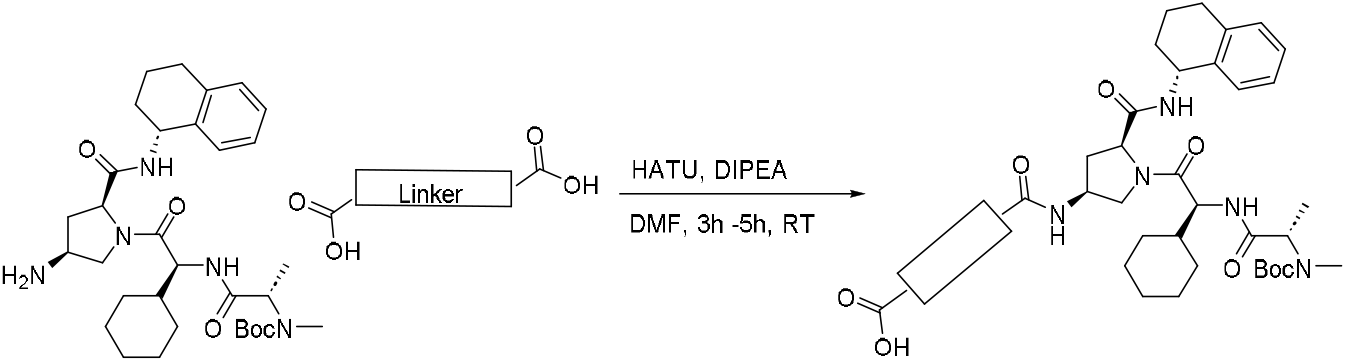

##### General Procedure 1B

To a stirred solution of *tert*-butyl ((*S*)-1-(((*S*)-2-((2*S*,4*S*)-4-amino-2-(((*R*)-1,2,3,4-tetrahydronaphthalen-1-yl)carbamoyl)pyrrolidin-1-yl)-1-cyclohexyl-2-oxoethyl)amino)-1-oxopropan-2-yl)(methyl)carbamate (1 eq, 0.13 mmol) in DMF (2 mL), were added sequentially dicarboxylic acid (1.5 eq, 0.19 mmol), HATU (1.5 eq, 0.19 mmol) DIPEA ( 3.0 eq, 0.39 mmol) at 0 °C. The resulting mixture was stirred at rt for 4h. After completion of the reaction (as indicated by LCMS) crude material was purified by reverse phase chromatography without workup using a gradient of 10-75% acetonitrile in water with the addition of 0.05% TFA to get the corresponding product.

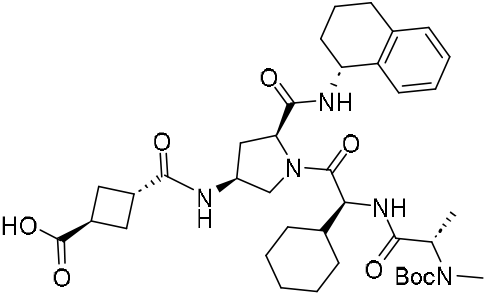

(1*S,*3*r*)-3-(((3*S*,5*S*)-1-((*S*)-2-((S)-2-((*tert*-butoxycarbonyl)(methyl)amino)propanamido)-2-cyclohexylacetyl)-5-(((*R*)-1,2,3,4-tetrahydronaphthalen-1-yl)carbamoyl)pyrrolidin-3-yl)carbamoyl)cyclobutane-1-carboxylic acid **(S9e):** General procedure 1B was followed on 75 mg (0.13 mmol) scale of the *tert*-butyl ((*S*)-1-(((*S*)-2-((2*S*,4*S*)-4-amino-2-(((*R*)-1,2,3,4-tetrahydronaphthalen-1-yl)carbamoyl)pyrrolidin-1-yl)-1-cyclohexyl-2-oxoethyl)amino)-1-oxopropan-2-yl)(methyl)carbamate. After the completion of the reaction, crude material was purified by reverse phase chromatography without workup using a gradient of 10-85% acetonitrile in water with the addition of 0.05% TFA to afford of title compound 91 mg (67%). ^1^H NMR (400 MHz, DMSO-d_6_) δ 8.39 (d, *J* = 8.7 Hz, 1H), 8.11 (d, *J* = 7.6 Hz, 1H), 7.29 (d, *J* = 7.4 Hz, 1H), 7.17 – 7.03 (m, 3H), 4.91 (q, *J* = 8.0 Hz, 1H), 4.28 (h, *J* = 8.1 Hz, 3H), 4.02 (dd, *J* = 9.9, 6.6 Hz, 1H), 3.31 (dd, *J* = 9.8, 7.3 Hz, 1H), 2.96 (qd, *J* = 9.2, 3.3 Hz, 2H), 2.71 (d, *J* = 9.5 Hz, 5H), 2.41 – 2.17 (m, 5H), 1.96 – 1.49 (m, 10H), 1.39 (d, *J* = 10.3 Hz, 9H), 1.26 – 0.82 (m, 8H); ^13^C NMR (101 MHz, DMSO-d6) δ 176.8, 174.1, 171.4, 170.2, 137.8, 137.4, 129.0, 128.8, 127.1, 126.1, 79.4, 58.9, 55.3, 52.8, 48.3, 47.1, 36.0, 34.9, 34.8, 30.5, 30.3, 29.2, 29.1, 28.5, 27.7, 27.6, 26.3, 26.2, 26.0, 20.7; LC-MS (ESI): (m/z) = 710 [M+H].

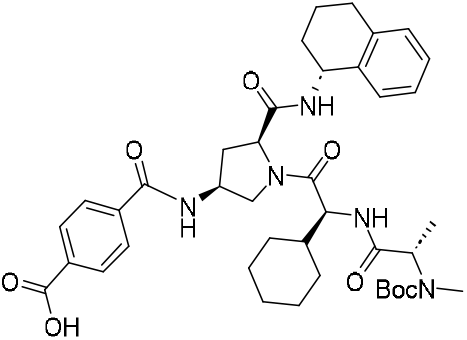

4-(((3*S*,5*S*)-1-((*S*)-2-((*S*)-2-((*tert*-butoxycarbonyl)(methyl)amino)propanamido)-2-cyclohexylacetyl)-5-(((*R*)-1,2,3,4-tetrahydronaphthalen-1-yl)carbamoyl)pyrrolidin-3-yl)carbamoyl)benzoic acid **(S9f)**: General procedure 1B was followed on 75 mg scale of the *tert*-butyl ((*S*)-1-(((*S*)-2-((*2S,4S*)-4-amino-2-(((*R*)-1,2,3,4-tetrahydronaphthalen-1-yl)carbamoyl)pyrrolidin-1-yl)-1-cyclohexyl-2-oxoethyl)amino)-1-oxopropan-2-yl)(methyl)carbamate hydrochloric salt. After the completion of the reaction, crude material was purified by reverse phase chromatography without workup using a gradient of 10-75% acetonitrile in water with the addition of 0.05% TFA to afford 45 mg (48%) of title compound as white solid. ^1^H NMR (400 MHz, DMSO-d6) δ 9.06 (d, *J* = 7.9 Hz, 1H), 8.54 (d, *J* = 8.7 Hz, 1H), 8.00 (d, *J* = 8.2 Hz, 2H), 7.94 (d, *J* = 8.3 Hz, 2H), 7.33 (d, *J* = 7.5 Hz, 1H), 7.16 – 6.99 (m, 3H), 5.00 – 4.90 (m, 1H), 4.55 (q, *J* = 6.8 Hz, 1H), 4.34 (dt, *J* = 14.5, 7.6 Hz, 2H), 4.05 (dd, *J* = 10.1, 6.2 Hz, 1H), 3.58 (dd, *J* = 10.2, 6.2 Hz, 1H), 2.72 (d, *J* = 8.4 Hz, 5H), 1.88 (ddq, *J* = 22.6, 12.1, 6.7 Hz, 4H), 1.78 – 1.46 (m, 5H), 1.38 (s, 9H), 1.20 (d, *J* = 7.5 Hz, 3H), 1.01 (dq, *J* = 41.2, 10.5 Hz, 2H); ^13^C NMR (101 MHz, DMSO-d6) δ 171.8, 170.4, 167.2, 165.5, 138.2, 137.7, 137.4, 133.7, 129.8, 129.0, 128.8, 127.8, 127.1, 126.1, 79.4, 59.0, 55.4, 53.1, 49.1, 47.3, 34.6, 30.5, 30.2, 29.2, 29.1, 28.5, 26.3, 26.2, 26.0, 20.7; LC-MS (ESI): (m/z) = 732 [M+H].

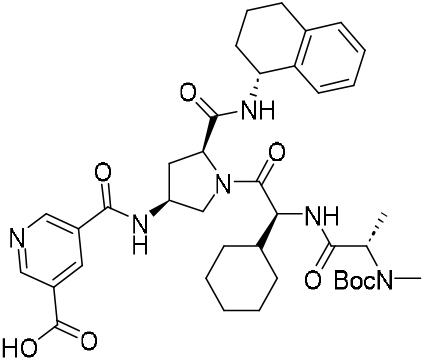

5-(((3*S*,5*S*)-1-((*S*)-2-((*S*)-2-((*tert*-butoxycarbonyl)(methyl)amino)propanamido)-2-cyclohexylacetyl)-5-(((*R*)-1,2,3,4-tetrahydronaphthalen-1-yl)carbamoyl)pyrrolidin-3-yl)carbamoyl)nicotinic acid **(S9g):** General procedure 1B was followed on 75 mg scale of the *tert*-butyl ((*S*)-1-(((*S*)-2-((2*S*,4*S*)-4-amino-2-(((*R*)-1,2,3,4-tetrahydronaphthalen-1-yl)carbamoyl)pyrrolidin-1-yl)-1-cyclohexyl-2-oxoethyl)amino)-1-oxopropan-2-yl)(methyl)carbamate hydrochloric salt. After the completion of the reaction, crude material was purified by reverse phase chromatography without workup using a gradient of 10-65% acetonitrile in water with the addition of 0.05% TFA to afford 60 mg (64%) of title compound as white solid. ^1^H NMR (400 MHz, DMSO-d_6_) δ 9.22 (d, *J* = 7.7 Hz, 2H), 8.69 (s, 1H), 8.50 (d, *J* = 8.7 Hz, 1H), 7.32 (d, *J* = 7.5 Hz, 1H), 7.10 (dt, *J* = 23.0, 7.5 Hz, 4H), 4.95 (q, *J* = 7.3 Hz, 1H), 4.55 (q, *J* = 7.1 Hz, 2H), 4.35 (q, *J* = 7.8 Hz, 2H), 4.11 (dd, *J* = 10.1, 6.6 Hz, 1H), 3.57 (dd, *J* = 10.0, 7.1 Hz, 1H), 2.72 (d, *J* = 10.4 Hz, 6H), 2.01 – 1.47 (m, 13H), 1.39 (d, *J* = 11.7 Hz, 11H), 1.24 – 0.80 (m, 11H); ^13^C NMR (101 MHz, dmso) δ 171.5, 170.3, 166.2, 164.0, 152.8, 152.3, 137.7, 137.4, 136.0, 129.0, 128.9, 127.1, 126.1, 79.4, 58.9, 55.4, 52.7, 49.0, 47.2, 34.6, 30.5, 30.2, 29.2, 29.1, 28.5, 26.3, 26.2, 26.0, 20.7; LC-MS (ESI): (m/z) = 733 [M+H].

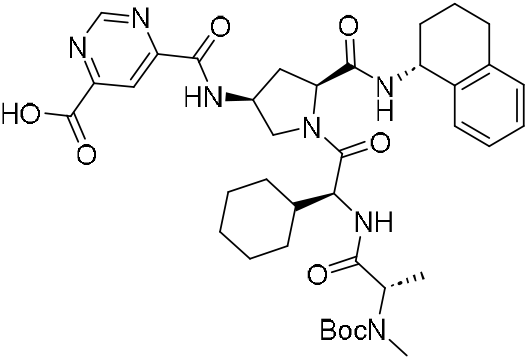

6-(((3*S*,5*S*)-1-((*S*)-2-((*S*)-2-((*tert*-butoxycarbonyl)(methyl)amino)propanamido)-2-cyclohexylacetyl)-5-(((*R*)-1,2,3,4-tetrahydronaphthalen-1-yl)carbamoyl)pyrrolidin-3-yl)carbamoyl)pyrimidine-4-carboxylic acid **(S9h).** General procedure 1B was followed on 50 mg (0.086%) scale of the *tert*-butyl ((*S*)-1-(((*S*)-2-((2*S*,4*S*)-4-amino-2-(((R)-1,2,3,4-tetrahydronaphthalen-1-yl)carbamoyl)pyrrolidin-1-yl)-1-cyclohexyl-2-oxoethyl)amino)-1-oxopropan-2-yl)(methyl)carbamate hydrochloric salt. After the completion of the reaction, crude material was purified by reverse phase chromatography without workup using a gradient of 10-75% acetonitrile in water with the addition of 0.05% TFA to afford 41 mg (65%) of title compound as white solid. ^1^H NMR (400 MHz, DMSO-d_6_) δ 9.70 (d, *J* = 8.4 Hz, 1H), 9.47 (d, *J* = 1.3 Hz, 1H), 8.46 (d, *J* = 8.7 Hz, 1H), 8.40 (d, *J* = 1.3 Hz, 1H), 7.30 (d, *J* = 7.5 Hz, 1H), 7.16 – 7.00 (m, 3H), 4.94 (td, *J* = 8.3, 4.6 Hz, 1H), 4.64 (p, *J* = 6.5 Hz, 1H), 4.43 – 4.26 (m, 2H), 4.03 (dd, *J* = 10.4, 6.1 Hz, 1H), 3.68 (dd, *J* = 10.3, 5.0 Hz, 1H), 2.71 (d, *J* = 8.8 Hz, 5H), 1.97 (dt, *J* = 13.0, 5.3 Hz, 1H), 1.89 – 1.45 (m, 4H), 1.39 (d, *J* = 12.1 Hz, 10H), 1.19 (d, *J* = 7.2 Hz, 4H), 1.11 – 0.83 (m, 6H); ^13^C NMR (101 MHz, DMSO-d6) δ 171.6, 170.4, 165.2, 162.4, 159.0, 158.9, 158.2, 137.8, 137.4, 129.0, 128.8, 127.1, 126.0, 118.0, 79.4, 59.1, 55.3, 53.3, 49.1, 47.2, 34.5, 30.5, 30.2, 29.2, 29.1, 28.5, 26.2, 26.0, 20.7; LC-MS (ESI): (m/z) = 734 [M+H].

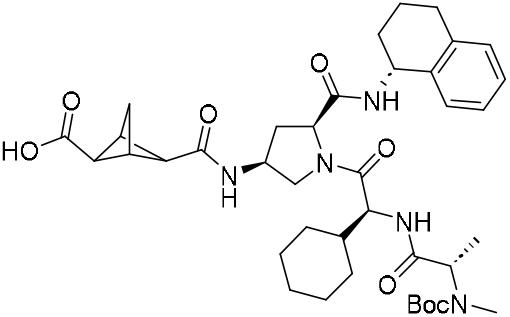

(1*R*,2*R*,3*S*,4r)-4-(((3*S*,5*S*)-1-((*S*)-2-((*S*)-2-((*tert*-butoxycarbonyl)(methyl)amino)propanamido)-2-cyclohexylacetyl)-5-(((R)-1,2,3,4-tetrahydronaphthalen-1-yl)carbamoyl)pyrrolidin-3-yl)carbamoyl)bicyclo[1.1.1]pentane-2-carboxylic acid **(S9i):** General procedure 1B was followed on 45 mg (0.077 mmol) scale of the *tert*-butyl ((S)-1-(((S)-2-((2S,4S)-4-amino-2-(((R)-1,2,3,4-tetrahydronaphthalen-1-yl)carbamoyl)pyrrolidin-1-yl)-1-cyclohexyl-2-oxoethyl)amino)-1-oxopropan-2-yl)(methyl)carbamate hydrochloric salt. After completion of the reaction, the crude material was purified by reverse phase chromatography without workup using a gradient of 10-85% acetonitrile in water with the addition of 0.05% TFA to afford 30 mg (64%) of title compound as white solid. ^1^H NMR (400 MHz, DMSO-d_6_) δ 12.45 (s, 1H), 8.46 (d, *J* = 8.6 Hz, 1H), 8.27 (d, *J* = 8.1 Hz, 1H), 7.30 (d, *J* = 7.5 Hz, 1H), 7.10 (dt, *J* = 23.3, 7.5 Hz, 3H), 4.91 (t, *J* = 7.0 Hz, 1H), 4.28 (t, *J* = 7.8 Hz, 3H), 4.06 – 3.88 (m, 1H), 3.44 – 3.33 (m, 1H), 2.71 (d, *J* = 5.8 Hz, 5H), 2.33 (dt, *J* = 13.9, 7.4 Hz, 1H), 2.08 (s, 6H), 1.94 – 1.51 (m, 9H), 1.39 (d, *J* = 12.4 Hz, 9H), 1.28 – 0.80 (m, 8H); ^13^C NMR (101 MHz, DMSO) δ 171.6, 171.2, 170.3, 168.7, 137.7, 137.4, 129.0, 128.9, 127.1, 126.1, 79.4, 62.6, 62.0, 58.9, 53.0, 51.9, 48.1, 47.2, 37.0, 34.6, 30.5, 30.2, 29.2, 29.0, 28.5, 26.3, 26.0, 20.7; LC-MS (ESI): (m/z) = 722 [M+H].

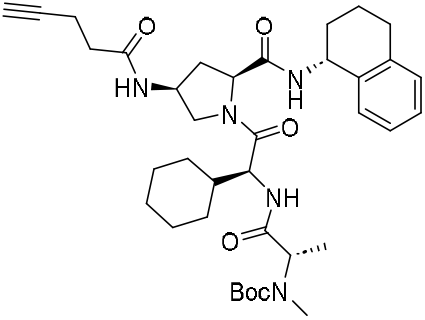

*tert*-butyl ((*S*)-1-(((*S*)-1-cyclohexyl-2-oxo-2-((2*S*,4*S*)-4-(pent-4-ynamido)-2-(((*R*)-1,2,3,4-tetrahydronaphthalen-1-yl)carbamoyl)pyrrolidin-1-yl)ethyl)amino)-1-oxopropan-2-yl)(methyl)carbamate **(S9j):** General procedure 1B was followed on 50 mg (0.086 mmol) scale of the *tert*-butyl ((*S*)-1-(((*S*)-2-((2*S*,4*S*)-4-amino-2-(((*R*)-1,2,3,4-tetrahydronaphthalen-1-yl)carbamoyl)pyrrolidin-1-yl)-1-cyclohexyl-2-oxoethyl)amino)-1-oxopropan-2-yl)(methyl)carbamate hydrochloric salt. After the completion of the reaction, crude material was purified by reverse phase chromatography without workup using a gradient of 10-85% acetonitrile in water with the addition of 0.05% TFA to afford 45 mg (79%) of title compound as white solid. ^1^H NMR (400 MHz, DMSO-d_6_) δ 8.36 (d, *J* = 8.6 Hz, 1H), 8.21 (d, *J* = 7.5 Hz, 1H), 7.30 (d, *J* = 7.4 Hz, 1H), 7.17 – 6.93 (m, 3H), 4.92 (td, *J* = 8.3, 4.2 Hz, 1H), 4.26 (tq, *J* = 15.3, 7.9 Hz, 3H), 4.06 (dd, *J* = 9.8, 6.9 Hz, 1H), 3.28 (t, *J* = 8.9 Hz, 1H), 2.80 – 2.62 (m, 7H), 2.35 (ddd, *J* = 9.5, 5.6, 2.8 Hz, 3H), 2.24 (t, *J* = 7.2 Hz, 2H), 1.91 – 1.51 (m, 9H), 1.39 (d, *J* = 10.9 Hz, 11H), 1.28 – 0.82 (m, 9H); ^13^C NMR (101 MHz, DMSO-d_6_) δ 171.2, 170.6, 170.2, 137.8, 137.4, 129.0, 128.8, 127.1, 126.1, 84.0, 79.4, 79.4, 71.8, 71.7, 58.9, 55.3, 52.5, 48.2, 47.1, 34.9, 34.7, 30.5, 30.3, 29.2, 29.1, 28.5, 26.3, 26.2, 26.0, 20.7, 14.5; LC-MS (ESI): (m/z) = 664 [M+H].

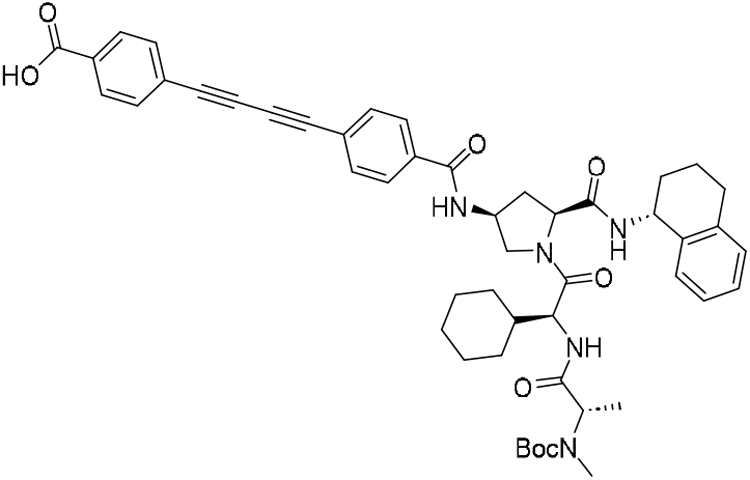

4-((4-(((3*S,*5*S*)-1-((*S*)-2-((*S*)-2-((*tert*-butoxycarbonyl)(methyl)amino)propanamido)-2-cyclohexylacetyl)-5-(((*R*)-1,2,3,4-tetrahydronaphthalen-1-yl)carbamoyl)pyrrolidin-3-yl)carbamoyl)phenyl)buta-1,3-diyn-1-yl)benzoic acid **(S9k):** General procedure 1B was followed on 75 mg (0.13 mmol) scale of the *tert*-butyl ((*S*)-1-(((*S*)-2-((2*S*,4*S*)-4-amino-2-(((*R*)-1,2,3,4-tetrahydronaphthalen-1-yl)carbamoyl)pyrrolidin-1-yl)-1-cyclohexyl-2-oxoethyl)amino)-1-oxopropan-2-yl)(methyl)carbamate hydrochloric salt. After the completion of the reaction, crude material was purified by reverse phase chromatography without workup using a gradient of 10-85% acetonitrile in water with the addition of 0.05% TFA to afford 40 mg (36%) of title compound as white solid. ^1^HNMR (400 MHz, DMSO-d_6_) δ 9.05 (d, *J* = 7.8 Hz, 1H), 8.55 (d, *J* = 8.6 Hz, 1H), 7.99 – 7.93 (m, 2H), 7.92 – 7.86 (m, 2H), 7.73 (ddd, *J* = 8.3, 3.4, 1.8 Hz, 4H), 7.33 (d, *J* = 7.6 Hz, 1H), 7.11 (dt, *J* = 23.1, 7.2 Hz, 3H), 4.95 (d, *J* = 5.7 Hz, 1H), 4.55 (q, *J* = 6.7 Hz, 1H), 4.34 (dt, *J* = 16.7, 7.8 Hz, 2H), 4.03 (t, *J* = 8.0 Hz, 1H), 3.58 (t, *J* = 8.1 Hz, 1H), 2.72 (d, *J* = 5.8 Hz, 5H), 1.96 – 1.78 (m, 4H), 1.78 – 1.48 (m, 4H), 1.37 (s, 9H), 1.19 (d, *J* = 7.4 Hz, 3H), 1.13 – 0.83 (m, 4H); LC-MS (ESI): (m/z) = 855 [M+H].

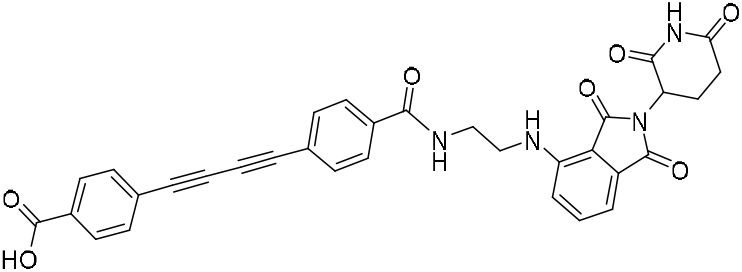

4-((4-((2-((2-(2,6-dioxopiperidin-3-yl)-1,3-dioxoisoindolin-4-yl)amino)ethyl)carbamoyl)phenyl)buta-1,3-diyn-1-yl)benzoic acid **(S9l):** General procedure 1B was followed on 150 mg (0.47 mmol) scale of the *tert*-butyl 4-((2-aminoethyl)amino)-2-(2,6-dioxopiperidin-3-yl)isoindoline-1,3-dione hydrochloric salt. After the completion of the reaction, crude material was purified by reverse phase chromatography without workup using a gradient of 10-75% acetonitrile in water with the addition of 0.05% TFA to afford 108 mg (38%) of title compound as white solid. ^1^H NMR (500 MHz, DMSO-d_6_) δ 13.11 (s, 1H), 11.03 (s, 1H), 8.76 (t, *J* = 5.4 Hz, 1H), 7.91 (d, *J* = 8.0 Hz, 2H), 7.81 (d, *J* = 8.0 Hz, 2H), 7.72 – 7.61 (m, 4H), 7.51 (t, *J* = 7.8 Hz, 1H), 7.17 (d, *J* = 8.6 Hz, 1H), 6.96 (d, *J* = 7.0 Hz, 1H), 6.78 (t, *J* = 6.2 Hz, 1H), 4.99 (dd, *J* = 12.8, 5.4 Hz, 1H), 3.43 (dq, *J* = 27.8, 6.8 Hz, 4H), 2.82 (ddd, *J* = 17.9, 13.9, 5.4 Hz, 1H), 2.56 – 2.45 (m, 2H), 2.02 – 1.91 (m, 1H); ^13^C NMR (126 MHz, DMSO-d_6_) δ 173.3, 170.6, 169.2, 167.8, 167.0, 166.3, 146.8, 136.7, 135.8, 133.2, 133.2, 133.0, 132.7, 132.2, 130.1, 128.1, 125.0, 123.3, 117.7, 111.1, 109.8, 82.8, 82.2, 76.0, 75.3, 49.0, 41.7, 39.3, 31.5, 22.6; LC-MS (ESI): (m/z) = 589 [M+H].

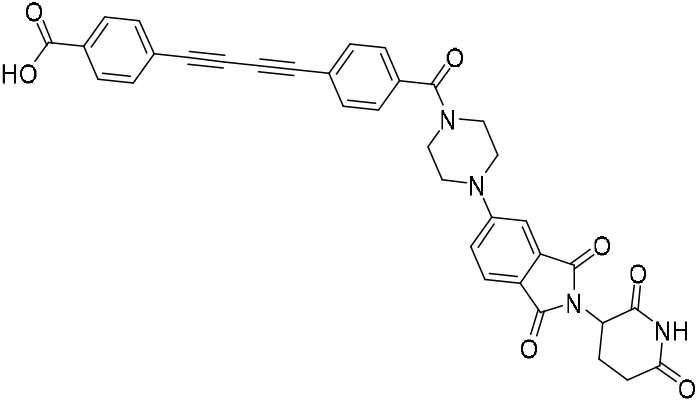

4-((4-(4-(2-(2,6-dioxopiperidin-3-yl)-1,3-dioxoisoindolin-5-yl)piperazine-1-carbonyl)phenyl)buta-1,3-diyn-1-yl)benzoic acid (**S9m**): General procedure 1B was followed on 100 mg (0.26 mmol) 2-(2,6-dioxopiperidin-3-yl)-5-(piperazin-1-yl)isoindoline-1,3-dione hydrochloric salt. After the completion of the reaction, crude material was purified by reverse phase chromatography without workup using a gradient of 10-75% acetonitrile in water with the addition of 0.05% TFA to afford 80 mg (49%) of title compound as yellow solid ^1^H NMR (500 MHz, DMSO) δ 11.09 (s, 1H), 7.98 (d, *J* = 8.3 Hz, 2H), 7.74 (td, *J* = 9.6, 7.7 Hz, 5H), 7.54 (d, *J* = 8.2 Hz, 2H), 7.37 (d, *J* = 2.3 Hz, 1H), 7.26 (dd, *J* = 8.6, 2.4 Hz, 1H), 5.08 (dd, *J* = 12.7, 5.5 Hz, 1H), 3.77 (s, 2H), 3.60 (s, 1H), 3.50 (s, 3H), 2.89 (ddd, *J* = 16.8, 13.7, 5.4 Hz, 1H), 2.64 – 2.52 (m, 2H), 2.02 (ddd, *J* = 17.4, 8.0, 4.7 Hz, 1H); ^13^C NMR (126 MHz, DMSO) δ 173.3, 170.5, 168.6, 168.0, 167.4, 167.0, 155.3, 137.5, 134.3, 133.2, 133.1, 132.2, 130.1, 128.1, 125.4, 125.0, 121.9, 119.2, 118.4, 108.6, 82.8, 82.0, 76.1, 74.7, 49.3, 31.4, 22.6; LC-MS (ESI): (m/z) = 615 [M+H].

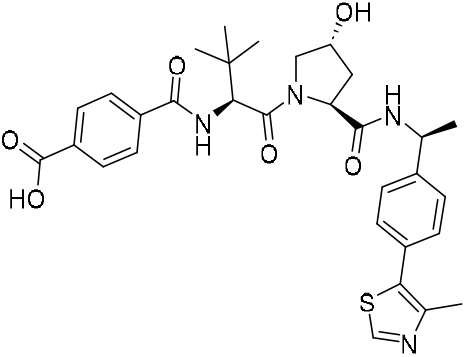

4-(((*S)-*1-((2*S*,4*R*)-4-hydroxy-2-(((*S*)-1-(4-(4-methylthiazol-5-yl)phenyl)ethyl)carbamoyl)pyrrolidin-1-yl)-3,3-dimethyl-1-oxobutan-2-yl)carbamoyl)benzoic acid **(S9m)**: General procedure 1B was followed on 150 mg (0.337 mmol) scale of the (2*S*,4*R*)-1-((*S*)-2-amino-3,3-dimethylbutanoyl)-4-hydroxy-N-((*S*)-1-(4-(4-methylthiazol-5-yl)phenyl)ethyl)pyrrolidine-2-carboxamide hydrochloric salt. After the completion of the reaction, crude material was purified by reverse phase chromatography without workup using a gradient of 10-65% acetonitrile in water with the addition of 0.05% TFA to afford 130 mg (67%) of title compound. ^1^H NMR (400 MHz, DMSO-d_6_) δ 9.04 (s, 1H), 8.40 (d, *J* = 7.8 Hz, 1H), 8.14 (d, *J* = 9.0 Hz, 1H), 8.04 – 7.96 (m, 2H), 7.93 (d, *J* = 8.4 Hz, 2H), 7.42 (d, *J* = 8.3 Hz, 2H), 7.37 (d, *J* = 8.3 Hz, 2H), 4.92 (p, *J* = 7.0 Hz, 1H), 4.75 (d, *J* = 9.1 Hz, 1H), 4.45 (t, *J* = 8.1 Hz, 1H), 4.29 (t, *J* = 3.6 Hz, 1H), 3.66 (d, *J* = 3.4 Hz, 2H), 2.44 (s, 3H), 2.07 – 1.97 (m, 1H), 1.79 (ddd, *J* = 12.9, 8.6, 4.6 Hz, 1H), 1.36 (d, *J* = 7.0 Hz, 3H), 1.02 (s, 9H); ^13^C NMR (101 MHz, DMSO-d_6_) δ 171.0, 169.7, 167.2, 166.3, 159.4, 159.0, 158.6, 158.2, 152.3, 147.6, 145.3, 138.4, 133.5, 131.9, 129.9, 129.6, 129.3, 128.3, 126.8, 69.3, 59.2, 58.0, 56.9, 48.2, 38.2, 36.1, 27.0, 26.9, 22.9, 16.1; LC-MS (ESI): (m/z) = 593 [M+H].

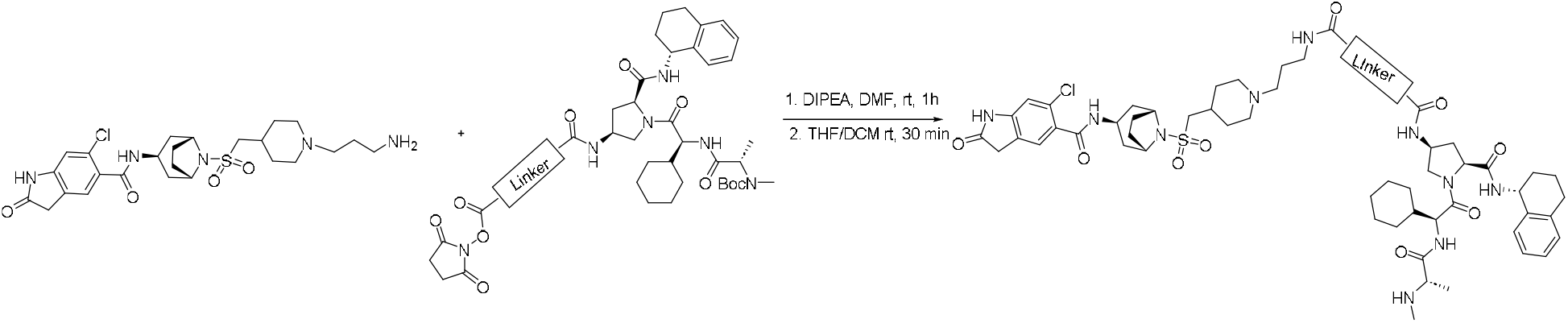

##### General Procedure 2A

A stirred solution of N-((*1R,3r,5S*)-8-(((1-(3-aminopropyl)piperidin-4-yl)methyl)sulfonyl)-8-azabicyclo[3.2.1]octan-3-yl)-6-chloro-2-oxoindoline-5-carboxamide (1.0 eq, 0.037 mmol) in DMF (1 mL), were sequentially added NHS ester of corresponding compound (1.1 eq, 0.040 mmol) and DIPEA (2.0 eq, 0.74 mmol) at 0 °C. The resulting mixture was stirred at room temperature for 1h. After completion of the reaction (as indicated by LCMS) the volatiles were removed under vacuum and the crude material was purified by reverse phase chromatography without workup the resulting product was used for next step without characterization. The resulting above product (0.037 mml) was dissolved 20% TFA in DCM (2 mL) and the resulting solution was stirred at room temperature for 1h. After completion of the reaction (as indicated by LCMS) the volatiles were removed under vacuum and the crude material was purified by reverse phase chromatography without workup to give corresponding product.

##### IAP-01

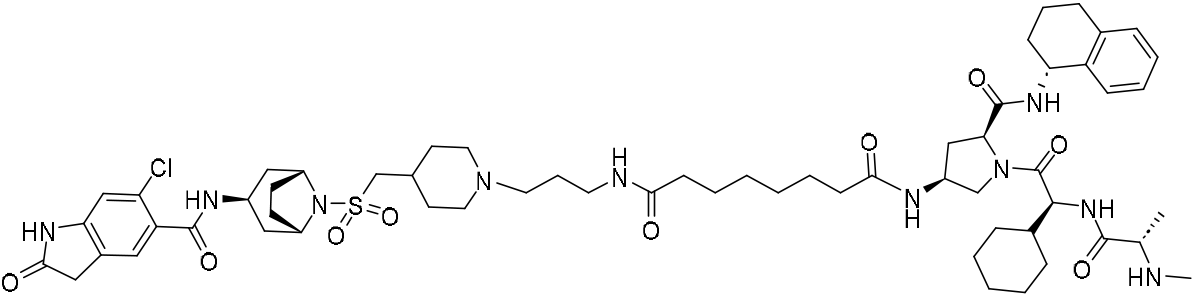

N1-(3-(4-((((*1R,3r,5S*)-3-(6-chloro-2-oxoindoline-5-carboxamido)-8-azabicyclo[3.2.1]octan-8-yl)sulfonyl)methyl)piperidin-1-yl)propyl)-N8-((3*S*,5*S*)-1-((*S*)-2-cyclohexyl-2-((S)-2-(methylamino)propanamido)acetyl)-5-(((*R*)-1,2,3,4-tetrahydronaphthalen-1-yl)carbamoyl)pyrrolidin-3-yl)octanediamide **(EPZ031686-IAP01)**: General procedure 2A was followed on 20 mg (0.037 mmol) scale of the N-((*1R,3r,5S*)-8-(((1-(3-aminopropyl)piperidin-4-yl)methyl)sulfonyl)-8-azabicyclo[3.2.1]octan-3-yl)-6-chloro-2-oxoindoline-5-carboxamide. After the completion of the reaction, crude material was purified by reverse phase chromatography without workup using a gradient of 10-70% acetonitrile in water with the addition of 0.05% TFA to afford 21 mg (44%) of title compound as white solid. ^1^H NMR (400 MHz, CD_3_OD-d_6_) δ 8.47 (d, *J* = 8.4 Hz, 1H), 7.40 – 7.34 (m, 1H), 7.30 (s, 1H), 7.11 (dtd, *J* = 16.9, 7.9, 5.9 Hz, 3H), 6.94 (s, 1H), 5.06 (t, *J* = 7.2 Hz, 1H), 4.52 – 4.38 (m, 3H), 4.31 – 4.09 (m, 5H), 3.87 (q, *J* = 6.9 Hz, 1H), 3.65 – 3.50 (m, 4H), 3.26 (d, *J* = 6.5 Hz, 1H), 3.05 (dd, *J* = 45.3, 14.7 Hz, 7H), 2.87 – 2.73 (m, 2H), 2.66 (s, 4H), 2.50 (dt, *J* = 14.7, 7.8 Hz, 1H), 2.40 – 2.11 (m, 10H), 2.10 – 1.57 (m, 14H), 1.47 (d, *J* = 7.0 Hz, 3H), 1.41 – 1.05 (m, 10H); LC-MS (ESI): (m/z) = 1159 [M+H].

##### IAP-02

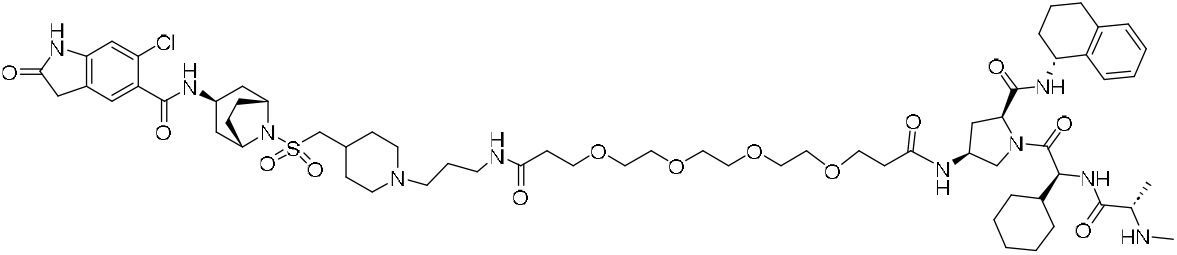

N1-(3-(4-((((*1R,3r,5S*)-3-(6-chloro-2-oxoindoline-5-carboxamido)-8-azabicyclo[3.2.1]octan-8-yl)sulfonyl)methyl)piperidin-1-yl)propyl)-N16-((3*S*,5*S*)-1-((*S*)-2-cyclohexyl-2-((S)-2-(methylamino)propanamido)acetyl)-5-(((*R*)-1,2,3,4-tetrahydronaphthalen-1-yl)carbamoyl)pyrrolidin-3-yl)-4,7,10,13-tetraoxahexadecanediamide **(EPZ031686-IAP02):** General procedure 2A was followed on 25 mg (0.046 mmol) scale of the N-((*1R,3r,5S*)-8-(((1-(3-aminopropyl)piperidin-4-yl)methyl)sulfonyl)-8-azabicyclo[3.2.1]octan-3-yl)-6-chloro-2-oxoindoline-5-carboxamide. After the completion of the reaction, crude material was purified by reverse phase chromatography without workup using a gradient of 10-70% acetonitrile in water with the addition of 0.05% TFA to afford 18 mg (33%) of title compound as white solid. ^1^H NMR (400 MHz, MeOD-d_4_) δ 5.05 (t, *J* = 7.1 Hz, 1H), 4.56 – 4.38 (m, 3H), 4.24 (t, *J* = 8.4 Hz, 3H), 4.13 (d, *J* = 6.6 Hz, 1H), 3.89 (q, *J* = 7.0 Hz, 1H), 3.74 (td, *J* = 6.1, 3.4 Hz, 4H), 3.62 (d, *J* = 6.0 Hz, 13H), 3.55 – 3.48 (m, 3H), 3.32 (d, *J* = 7.9 Hz, 2H), 3.12 (dd, *J* = 16.1, 7.0 Hz, 3H), 3.00 (t, *J* = 12.7 Hz, 2H), 2.78 (dd, *J* = 11.0, 6.2 Hz, 1H), 2.66 (d, *J* = 0.9 Hz, 3H), 2.47 (t, *J* = 6.0 Hz, 4H), 2.28 (t, *J* = 12.1 Hz, 4H), 2.16 (d, *J* = 7.9 Hz, 1H), 2.09 – 1.63 (m, 12H), 1.47 (d, *J* = 7.0 Hz, 3H), 1.38 – 1.01 (m, 5H); ^13^C NMR (101 MHz, MeOD-d_4_) δ 178.1, 174.0, 172.1, 171.7, 170.7, 168.7, 168.5, 145.8, 137.1, 136.2, 130.2, 129.5, 128.5, 128.4, 126.8, 125.7, 124.8, 124.7, 110.4, 70.1, 70.1, 70.0, 69.9, 69.9, 66.8, 66.7, 59.2, 56.8, 56.2, 55.6, 53.9, 52.7, 52.1, 48.4, 42.3, 39.8, 36.3, 36.1, 35.3, 34.1, 30.4, 30.2, 29.9, 29.1, 28.9, 28.8, 28.5, 28.1, 25.8, 25.6, 24.1, 20.2, 14.8; HRMS (ESI) calcd for C_64_H_95_ClN_10_O_13_SNa^+^ (MNa^+^) 1301.6387, found 1301.6388.

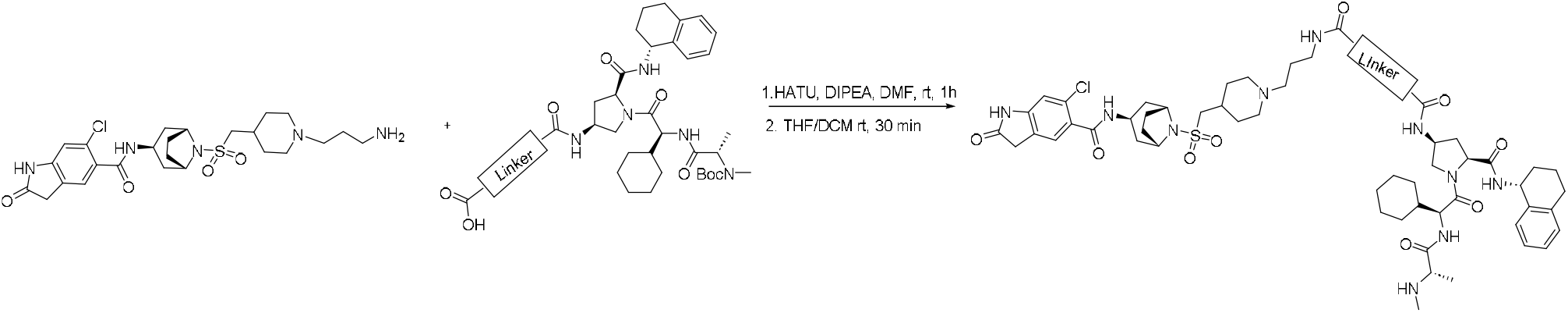

##### General Procedure 2B

A stirred solution of N-((*1R,3r,5S*)-8-(((1-(3-aminopropyl)piperidin-4-yl)methyl)sulfonyl)-8-azabicyclo[3.2.1]octan-3-yl)-6-chloro-2-oxoindoline-5-carboxamide (1.0 eq, 0.037 mmol) in DMF (1 mL), were sequentially added corresponding IAP carboxylic acid (1.1 eq, 0.040 mmol), HATU (1.5 eq, 0.055 mmol) and DIPEA (3.0 eq, 0.111 mmol) at 0 °C. The resulting mixture was stirred at room temperature for 3h to 4h. After completion of the reaction (as indicated by LCMS) the volatiles were removed under vacuum and the crude material was purified by reverse phase chromatography without workup the resulting product was used for next step without characterization. The resulting above product (0.037 mmL) was dissolved 20% TFA in DCM (2 mL) and the resulting solution was stirred at room temperature for 1h. After completion of the reaction (as indicated by LCMS) the volatiles were removed under vacuum and the crude material was purified by reverse phase chromatography without workup to give corresponding product.

##### IAP-03

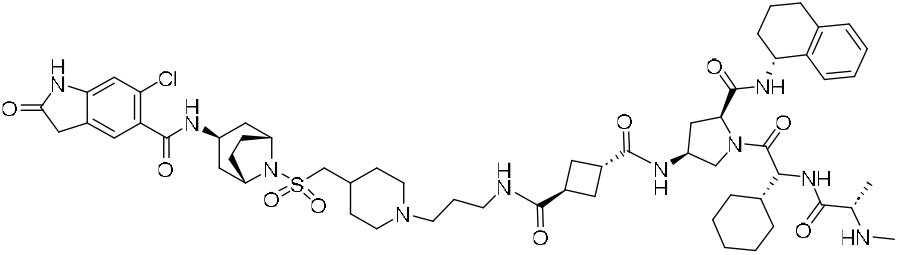

(1S,3R)-N1-(3-(4-((((*1R,3r,5S*)-3-(6-chloro-2-oxoindoline-5-carboxamido)-8-azabicyclo[3.2.1]octan-8-yl)sulfonyl)methyl)piperidin-1-yl)propyl)-N3-((3*S*,5*S*)-1-((*S*)-2-cyclohexyl-2-((*S*)-2-(methylamino)propanamido)acetyl)-5-(((*R*)-1,2,3,4-tetrahydronaphthalen-1-yl)carbamoyl)pyrrolidin-3-yl)cyclobutane-1,3-dicarboxamide **(EPZ031686-IAP03):** General procedure 2B was followed on 17 mg (0.031 mmol) scale of the N-((*1R,3r,5S*)-8-(((1-(3-aminopropyl)piperidin-4-yl)methyl)sulfonyl)-8-azabicyclo[3.2.1]octan-3-yl)-6-chloro-2-oxoindoline-5-carboxamide. After the completion of the reaction, crude material was purified by reverse phase chromatography without workup using a gradient of 10-70% acetonitrile in water with the addition of 0.05% TFA to afford 8 mg (23%) of title compound. ^1^H NMR (400 MHz, MeOD-d_4_) δ 7.73 (s, 1H), 7.37 (d, *J* = 7.2 Hz, 1H), 7.30 (d, *J* = 2.3 Hz, 1H), 7.11 (dq, *J* = 16.8, 9.3 Hz, 3H), 6.98 – 6.91 (m, 1H), 6.53 (s, 0H), 5.04 (s, 1H), 4.58 – 4.31 (m, 3H), 4.31 – 4.06 (m, 5H), 3.89 (t, *J* = 7.3 Hz, 1H), 3.58 (t, *J* = 13.2 Hz, 4H), 3.30 (tt, *J* = 4.4, 1.8 Hz, 6H), 3.22 – 2.97 (m, 7H), 2.79 (d, *J* = 11.4 Hz, 2H), 2.57 – 2.39 (m, 6H), 2.34 – 1.62 (m, 25H), 1.47 (dd, *J* = 6.9, 2.4 Hz, 3H), 1.37 – 0.99 (m, 5H); ^13^C NMR (101 MHz, MeOD-d_4_) δ 1777.0, 175.9, 171.9, 170.7, 168.7, 145.6, 137.1, 136.1, 130.2, 129.5, 128.5, 128.4, 126.8, 125.6, 124.6, 110.4, 59.3, 56.8, 56.2, 55.6, 54.3, 53.0, 52.1, 48.6, 42.3, 39.8, 36.3, 36.2, 35.9, 35.7, 34.1, 30.4, 30.1, 29.8, 29.0, 28.8, 28.8, 28.5, 28.1, 27.4, 27.3, 25.8, 25.6, 24.2, 20.2; HRMS (ESI) calcd for C_58_H_82_ClN_10_O_9_S^+^ (MH+) 1129.5675, found 1129.5667.

##### IAP-04

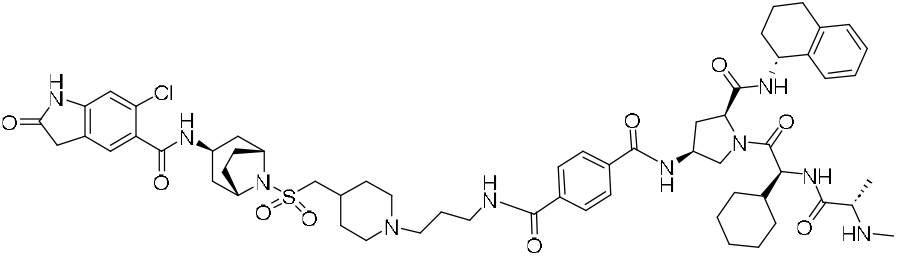

N1-(3-(4-((((*1R,3r,5S*)-3-(6-chloro-2-oxoindoline-5-carboxamido)-8-azabicyclo[3.2.1]octan-8-yl)sulfonyl)methyl)piperidin-1-yl)propyl)-N4-((3*S*,5*S*)-1-((*S*)-2-cyclohexyl-2-((*S*)-2-(methylamino)propanamido)acetyl)-5-(((*R*)-1,2,3,4-tetrahydronaphthalen-1-yl)carbamoyl)pyrrolidin-3-yl)terephthalamide **(EPZ031686-IAP04):** General procedure 2B was followed on 15 mg (0.023 mmol) scale of the N-((*1R,3r,5S*)-8-(((1-(3-aminopropyl)piperidin-4-yl)methyl)sulfonyl)-8-azabicyclo[3.2.1]octan-3-yl)-6-chloro-2-oxoindoline-5-carboxamide. After the completion of the reaction, crude material was purified by reverse phase chromatography without workup using a gradient of 10-70% acetonitrile in water with the addition of 0.05% TFA to afford 11 mg (42%) of title compound. ^1^H NMR (400 MHz, MeOD-d4) δ 8.60 (d, *J* = 8.5 Hz, 1H), 7.97 (q, *J* = 8.3 Hz, 4H), 7.40 (d, *J* = 7.3 Hz, 1H), 7.29 (s, 1H), 7.11 (dt, *J* = 18.2, 7.0 Hz, 3H), 6.94 (s, 1H), 5.09 (d, *J* = 7.3 Hz, 1H), 4.76 (s, 1H), 4.57 – 4.45 (m, 2H), 4.23 (s, 2H), 4.13 (dd, *J* = 10.5, 5.2 Hz, 2H), 3.96 – 3.79 (m, 2H), 3.62 (d, *J* = 12.2 Hz, 2H), 3.51 (d, *J* = 9.8 Hz, 4H), 3.21 – 3.09 (m, 3H), 3.03 (t, *J* = 12.8 Hz, 2H), 2.88 – 2.71 (m, 1H), 2.66 (s, 3H), 2.65 – 2.54 (m, 1H), 2.38 – 1.53 (m, 21H), 1.48 (d, *J* = 7.0 Hz, 3H), 1.29 – 0.96 (m, 7H); ^13^C NMR (101 MHz, MeOD-d4) δ 172.4, 170.9, 168.7, 168.4, 166.7, 137.1, 136.7, 136.6, 136.1, 128.5, 128.4, 127.3, 127.2, 126.8, 125.6, 124.6, 59.4, 56.8, 56.1, 55.5, 54.5, 53.8, 52.2, 49.5, 42.3, 40.0, 36.5, 36.2, 33.9, 30.4, 30.1, 29.8, 28.9, 28.8, 28.6, 28.1, 25.8, 25.7, 25.6, 24.2, 20.2, 14.8; LC-MS (ESI): (m/z) = 1151 [M+H].

##### IAP-05

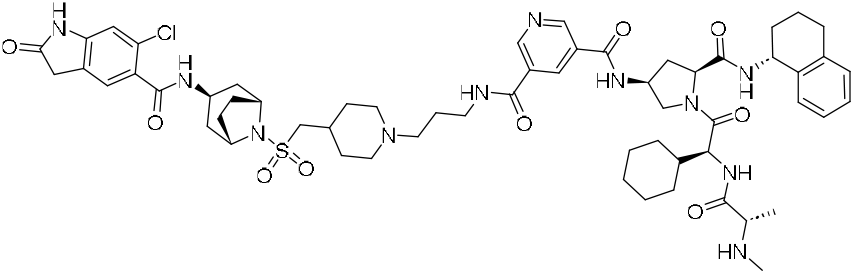

N3-(3-(4-((((*1R,3r,5S*)-3-(6-chloro-2-oxoindoline-5-carboxamido)-8-azabicyclo[3.2.1]octan-8-yl)sulfonyl)methyl)piperidin-1-yl)propyl)-N5-((3*S*,5*S*)-1-((S)-2-cyclohexyl-2-((*S*)-2-(methylamino)propanamido)acetyl)-5-(((*R*)-1,2,3,4-tetrahydronaphthalen-1-yl)carbamoyl)pyrrolidin-3-yl)pyridine-3,5-dicarboxamide **(EPZ031686-IAP05):** General procedure 2B was followed on 20 mg (0.031 mmol) scale of the N-((*1R,3r,5S*)-8-(((1-(3-aminopropyl)piperidin-4-yl)methyl)sulfonyl)-8-azabicyclo[3.2.1]octan-3-yl)-6-chloro-2-oxoindoline-5-carboxamide. After the completion of the reaction, crude material was purified by reverse phase chromatography without workup using a gradient of 10-70% acetonitrile in water with the addition of 0.05% TFA to afford 13 mg (37%) of title compound. ^1^H NMR (400 MHz, MeOD-d4) δ 9.18 (d, *J* = 22.0 Hz, 2H), 8.71 (s, 1H), 8.59 (d, *J* = 8.6 Hz, 1H), 7.46 – 7.36 (m, 1H), 7.30 (s, 0H), 7.11 (dt, *J* = 18.1, 7.1 Hz, 3H), 6.94 (d, *J* = 4.4 Hz, 1H), 5.19 – 5.03 (m, 1H), 4.79 – 4.72 (m, 1H), 4.60 – 4.47 (m, 2H), 4.30 – 4.09 (m, 4H), 3.94 – 3.71 (m, 2H), 3.62 (d, *J* = 11.9 Hz, 2H), 3.53 (d, *J* = 5.0 Hz, 3H), 3.19 (t, *J* = 7.9 Hz, 2H), 3.13 (d, *J* = 5.8 Hz, 1H), 3.01 (t, *J* = 12.5 Hz, 1H), 2.79 (td, *J* = 14.8, 8.4 Hz, 2H), 2.66 (s, 4H), 2.34 – 1.56 (m, 18H), 1.48 (d, *J* = 7.0 Hz, 3H), 1.33 – 0.99 (m, 6H); ^13^C NMR (101 MHz, MeOD-d4) δ 172.2, 170.8, 168.7, 168.5, 166.3, 164.7, 150.2, 150.1, 137.1, 136.2, 134.6, 130.2, 129.5, 128.5, 128.4, 126.8, 125.6, 124.6, 110.4, 59.4, 56.8, 56.1, 55.5, 54.5, 53.4, 52.2, 49.5, 42.3, 40.0, 36.5, 36.3, 33.9, 30.4, 30.1, 29.8, 29.0, 28.8, 28.8, 28.5, 28.2, 25.8, 25.7, 25.6, 24.1, 20.2, 14.8; HRMS (ESI) calcd for C_59_H_79_ClN_11_O_9_S^+^ (MH+) 1152.5471, found 1152.5482.

##### IAP-07

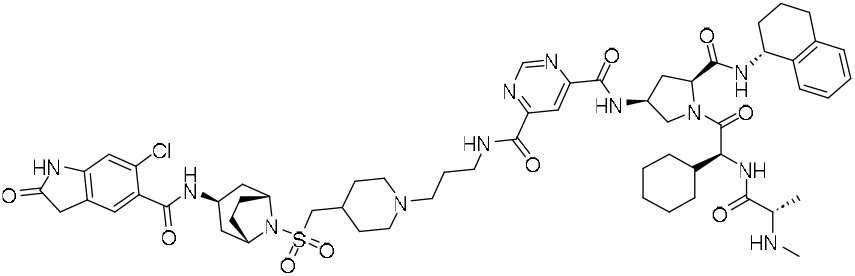

N4-(3-(4-((((*1R,3r,5S*)-3-(6-chloro-2-oxoindoline-5-carboxamido)-8-azabicyclo[3.2.1]octan-8-yl)sulfonyl)methyl)piperidin-1-yl)propyl)-N6-((3*S*,5*S*)-1-((*S*)-2-cyclohexyl-2-((*S*)-2-(methylamino)propanamido)acetyl)-5-(((*R*)-1,2,3,4-tetrahydronaphthalen-1-yl)carbamoyl)pyrrolidin-3-yl)pyrimidine-4,6-dicarboxamide **(EPZ031686-IAP07):** General procedure 2B was followed on 15 mg (0.028 mmol) scale of the N-((*1R,3r,5S*)-8-(((1-(3-aminopropyl)piperidin-4-yl)methyl)sulfonyl)-8-azabicyclo[3.2.1]octan-3-yl)-6-chloro-2-oxoindoline-5-carboxamide. After the completion of the reaction, crude material was purified by reverse phase chromatography without workup using a gradient of 10-70% acetonitrile in water with the addition of 0.05% TFA to afford 10 mg (54 %) of title compound. ^1^H NMR (400 MHz, DMSO-d6) δ 10.59 (s, 1H), 9.71 (d, *J* = 8.3 Hz, 1H), 9.44 (d, *J* = 2.5 Hz, 1H), 9.38 (s, 1H), 8.50 (d, *J* = 8.6 Hz, 1H), 8.44 (d, *J* = 2.2 Hz, 1H), 8.17 (d, *J* = 4.7 Hz, 1H), 7.96 (d, *J* = 8.5 Hz, 1H), 7.31 (d, *J* = 7.6 Hz, 1H), 7.21 (s, 1H), 7.08 (dd, *J* = 20.6, 7.3 Hz, 4H), 6.81 (d, *J* = 2.4 Hz, 1H), 4.95 (s, 1H), 4.67 (s, 1H), 4.45 – 4.28 (m, 2H), 4.07 (d, *J* = 18.7 Hz, 4H), 3.94 (s, 1H), 3.70 (d, *J* = 10.4 Hz, 1H), 3.34 (d, *J* = 27.1 Hz, 16H), 3.01 (dd, *J* = 14.0, 6.4 Hz, 4H), 2.92 – 2.83 (m, 3H), 2.70 (s, 2H), 2.48 (s, 2H), 2.33 (d, *J* = 6.5 Hz, 3H), 2.13 – 1.44 (m, 28H), 1.36 (d, *J* = 12.0 Hz, 2H), 1.17 – 0.87 (m, 12H); ^13^C NMR (101 MHz, DMSO-d6) δ 176.8, 174.5, 171.7, 170.5, 166.9, 162.4, 162.3, 159.4, 159.1, 158.0, 146.0, 137.8, 137.4, 130.2, 129.6, 129.0, 128.8, 127.1, 126.0, 125.3, 125.1, 115.6, 110.1, 59.4, 59.1, 57.7, 57.0, 55.3, 54.7, 53.5, 49.1, 47.3, 42.0, 36.7, 35.7, 34.5, 32.5, 31.8, 30.1, 29.1, 28.3, 26.3, 26.0, 20.7, 19.4; HRMS (ESI) calcd for C_58_H_78_ClN_12_O_9_S^+^ (MH+) 1153.5424, found 1153.5442.

##### IAP-08

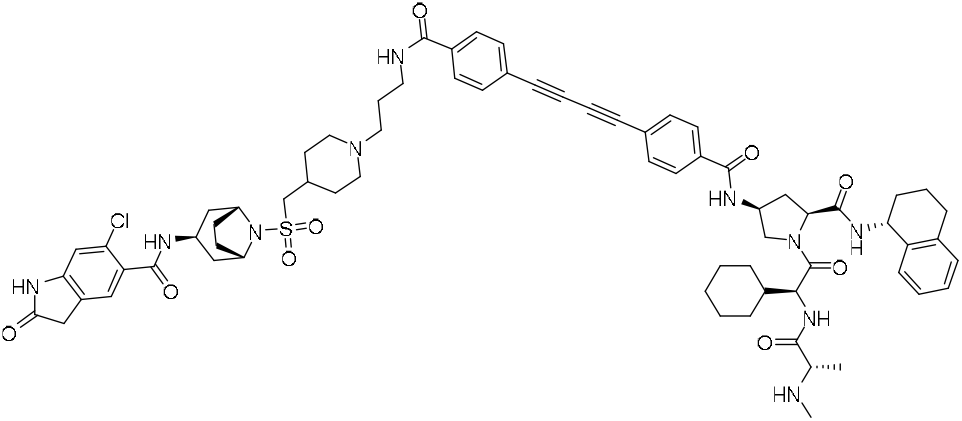

6-chloro-N-((*1R,3r,5S*)-8-(((1-(3-(4-((4-(((3*S*,5*S*)-1-((*S*)-2-cyclohexyl-2-((*S*)-2-(methylamino)propanamido)acetyl)-5-(((*R*)-1,2,3,4-tetrahydronaphthalen-1-yl)carbamoyl)pyrrolidin-3-yl)carbamoyl)phenyl)buta-1,3-diyn-1-yl)benzamido)propyl)piperidin-4-yl)methyl)sulfonyl)-8-azabicyclo[3.2.1]octan-3-yl)-2-oxoindoline-5-carboxamide **(EPZ031686-IAP08):** General procedure 2B was followed on 15 mg (0.028 mmol) scale of the N-((*1R,3r,5S*)-8-(((1-(3-aminopropyl)piperidin-4-yl)methyl)sulfonyl)-8-azabicyclo[3.2.1]octan-3-yl)-6-chloro-2-oxoindoline-5-carboxamide. After the completion of the reaction, crude material was purified by reverse phase chromatography without workup using a gradient of 10-70% acetonitrile in water with the addition of 0.05% TFA to afford 8 mg (22%) of title compound. ^1^H NMR (400 MHz, DMSO-d6) δ 10.59 (s, 1H), 9.06 (d, *J* = 7.8 Hz, 1H), 8.66 – 8.54 (m, 2H), 8.18 (d, *J* = 4.7 Hz, 1H), 7.96 – 7.80 (m, 5H), 7.76 – 7.66 (m, 4H), 7.37 – 7.30 (m, 1H), 7.20 (d, *J* = 1.2 Hz, 1H), 7.15 – 7.04 (m, 3H), 6.81 (s, 1H), 5.00 – 4.90 (m, 1H), 4.55 (h, *J* = 6.5 Hz, 1H), 4.43 – 4.33 (m, 2H), 4.11 – 4.00 (m, 3H), 3.94 (d, *J* = 5.8 Hz, 1H), 3.59 (dd, *J* = 10.1, 6.0 Hz, 1H), 3.49 (s, 2H), 3.26 (d, *J* = 8.7 Hz, 4H), 2.97 (dd, *J* = 13.0, 6.3 Hz, 3H), 2.82 (d, *J* = 11.2 Hz, 2H), 2.76 – 2.64 (m, 2H), 2.29 (t, *J* = 7.1 Hz, 2H), 2.16 (s, 3H), 2.08 (dd, *J* = 25.0, 6.9 Hz, 2H), 1.95 – 1.46 (m, 21H); ^13^C NMR (101 MHz, DMSO-d6) δ 176.8, 174.7, 171.8, 170.5, 166.9, 165.5, 165.1, 145.9, 137.7, 137.4, 136.1, 135.3, 133.0, 132.9, 130.2, 129.6, 129.0, 128.9, 128.0, 128.0, 127.1, 126.1, 125.3, 125.1, 123.6, 123.1, 110.1, 82.5, 82.3, 75.5, 75.3, 59.5, 59.0, 57.7, 56.4, 55.3, 54.7, 53.5, 53.2, 49.1, 47.3, 42.0, 38.5, 36.7, 35.7, 34.6, 32.4, 31.9, 30.2, 29.2, 28.4, 28.3, 26.8, 26.3, 26.2, 26.0, 20.7, 19.5; HRMS (ESI) calcd for C_70_H_84_ClN_10_O_9_S^+^ (MH+) 1275.5832, found 1275.5841.

##### IAP-09

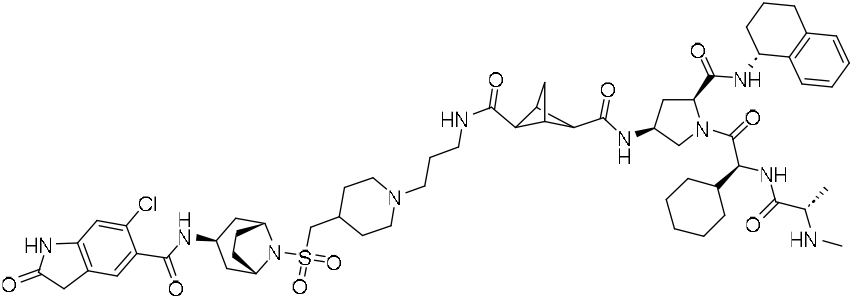

(1R,2S,3S,4S)-N2-(3-(4-((((*1R,3r,5S*)-3-(6-chloro-2-oxoindoline-5-carboxamido)-8-azabicyclo[3.2.1]octan-8-yl)sulfonyl)methyl)piperidin-1-yl)propyl)-N4-((3*S*,5*S*)-1-((*S*)-2-cyclohexyl-2-((*S*)-2-(methylamino)propanamido)acetyl)-5-(((*R*)-1,2,3,4-tetrahydronaphthalen-1-yl)carbamoyl)pyrrolidin-3-yl)bicyclo[1.1.1]pentane-2,4-dicarboxamide **(EPZ031686-IAP09):** General procedure 2B was followed on 15 mg (0.028 mmol) scale of the N-((*1R,3r,5S*)-8-(((1-(3-aminopropyl)piperidin-4-yl)methyl)sulfonyl)-8-azabicyclo[3.2.1]octan-3-yl)-6-chloro-2-oxoindoline-5-carboxamide. After the completion of the reaction, crude material was purified by reverse phase chromatography without workup using a gradient of 10-70% acetonitrile in water with the addition of 0.05% TFA to afford 12 mg (37%) of title compound. ^1^H NMR (400 MHz, DMSO-d_6_) δ 10.60 (s, 1H), 8.51 (d, *J* = 8.6 Hz, 1H), 8.28 (d, *J* = 8.1 Hz, 1H), 8.19 (d, *J* = 4.4 Hz, 1H), 7.91 (d, *J* = 8.4 Hz, 1H), 7.80 (d, *J* = 6.1 Hz, 1H), 7.32 (d, *J* = 7.4 Hz, 1H), 7.22 (s, 1H), 7.17 – 7.04 (m, 3H), 6.83 (d, *J* = 1.7 Hz, 1H), 4.94 (s, 1H), 4.34 (dt, *J* = 19.7, 7.8 Hz, 3H), 4.10 (s, 2H), 3.95 (d, *J* = 6.9 Hz, 2H), 3.50 (s, 2H), 3.43 (t, *J* = 8.2 Hz, 1H), 3.11 – 2.92 (m, 6H), 2.85 – 2.65 (m, 4H), 2.43 – 2.31 (m, 1H), 2.26 – 2.00 (m, 12H), 1.98 – 1.46 (m, 16H), 1.36 – 0.89 (m, 14H); ^13^C NMR (101 MHz, DMSO-d_6_) δ 176.8, 174.8, 171.7, 170.5, 169.1, 169.0, 166.9, 145.9, 137.6, 137.4, 130.2, 129.6, 129.0, 127.1, 126.1, 125.3, 125.1, 110.1, 59.5, 58.9, 57.7, 56.2, 55.3, 54.7, 53.4, 53.1, 51.6, 48.1, 47.2, 42.0, 38.5, 38.4, 37.5, 36.7, 35.7, 34.7, 32.4, 31.8, 30.1, 29.2, 29.1, 28.5, 28.3, 26.9, 26.4, 26.3, 26.0, 20.6, 19.5; HRMS (ESI) calcd for C_59_H_82_ClN_10_O_9_S^+^ (MH+) 1141.5675, found 1141.5693.

##### IAP10

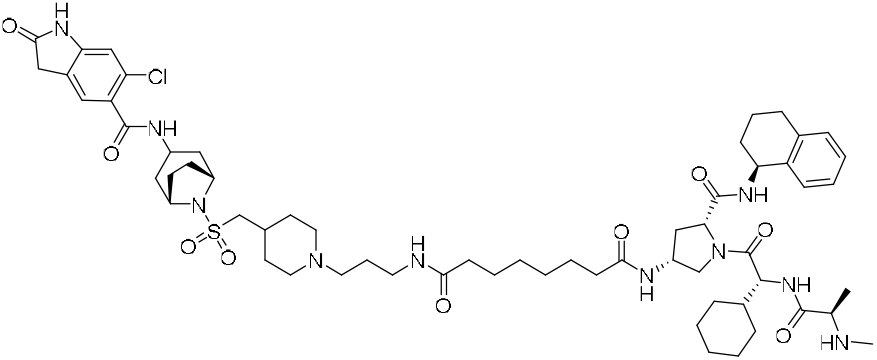

General procedure 2A was followed on 25 mg (0.046 mmol) scale of the N-((*1R,3r,5S*)-8-(((1-(3-aminopropyl)piperidin-4-yl)methyl)sulfonyl)-8-azabicyclo[3.2.1]octan-3-yl)-6-chloro-2-oxoindoline-5-carboxamide. After the completion of the reaction, crude material was purified by reverse phase chromatography without workup using a gradient of 10-70% acetonitrile in water with the addition of 0.05% TFA to afford 20 mg (34%) of title compound as white solid. ^1^H NMR (500 MHz, CD_3_OD_SPE) δ 7.39 (d, *J* = 7.4 Hz, 1H), 7.29 (s, 1H), 7.16 – 7.04 (m, 3H), 6.93 (d, *J* = 2.4 Hz, 1H), 5.01 (d, *J* = 6.8 Hz, 1H), 4.48 (d, *J* = 9.1 Hz, 2H), 4.39 (td, *J* = 11.0, 6.4 Hz, 1H), 4.22 (s, 2H), 4.12 (t, *J* = 7.1 Hz, 1H), 4.04 (d, *J* = 8.7 Hz, 0H), 3.91 – 3.76 (m, 1H), 3.61 – 3.47 (m, 4H), 3.26 (t, *J* = 6.7 Hz, 3H), 3.16 – 3.05 (m, 4H), 3.01 (t, *J* = 12.8 Hz, 2H), 2.66 (d, *J* = 5.0 Hz, 3H), 2.53 (tt, *J* = 15.2, 7.2 Hz, 1H), 2.33 – 2.11 (m, 10H), 2.07 – 1.89 (m, 6H), 1.84 – 1.58 (m, 8H), 1.49 (dd, *J* = 26.5, 6.9 Hz, 3H), 1.38 – 1.18 (m, 7H), 1.08 (dq, *J* = 25.2, 12.3 Hz, 2H); LC-MS (ESI): ^13^C NMR (126 MHz, CD_3_OD_SPE) δ 178.1, 175.7, 174.3, 172.1, 170.5, 168.7, 168.5, 145.8, 137.1, 136.2, 130.2, 129.5, 128.6, 128.1, 126.8, 125.7, 124.8, 124.7, 110.4, 59.1, 57.0, 56.9, 56.1, 55.6, 54.4, 53.0, 52.2, 42.3, 39.4, 36.3, 35.7, 35.5, 34.7, 30.6, 30.2, 30.0, 29.2, 28.8, 28.7, 28.6, 28.6, 28.1, 25.8, 25.6, 25.5, 25.3, 25.2, 24.1, 20.4, 15.1; (m/z) = 1259 [M+H].

##### IAP-11

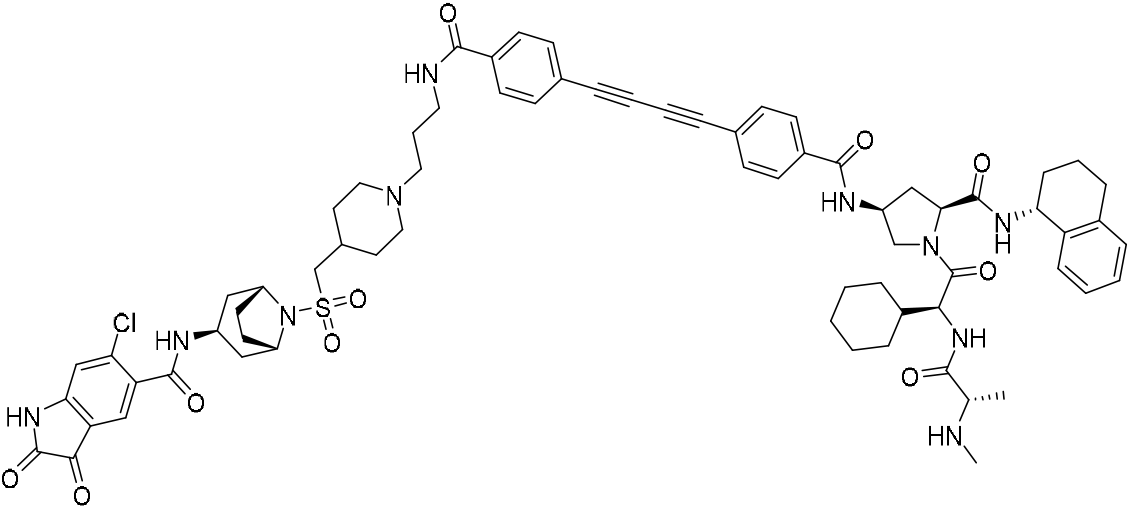

6-chloro-N-((*1R,3r,5S*)-8-(((1-(3-(4-((4-(((3*S*,5*S*)-1-((*S*)-2-cyclohexyl-2-((*S*)-2-(methylamino)propanamido)acetyl)-5-(((*R*)-1,2,3,4-tetrahydronaphthalen-1-yl)carbamoyl)pyrrolidin-3-yl)carbamoyl)phenyl)buta-1,3-diyn-1-yl)benzamido)propyl)piperidin-4-yl)methyl)sulfonyl)-8-azabicyclo[3.2.1]octan-3-yl)-2,3-dioxoindoline-5-carboxamide (**EPZ031686-IAP11**): To a stirred solution of the **EPZ031686-IAP08** (25 mg, 1 eq, 18 μmol), in DMF (0.5 mL), were added Cupric acetate, monohydrate (0. 36 mg,, 0.1 eq, 1.8 μmol), and K_2_CO_3_ (0.18 mg, 0.1 eq, 1.8 μmol) at room temperature. After completion of the reaction (as indicated by LCMS), the crude material was purified by reverse phase chromatography using a gradient of 10-65% acetonitrile in water with the addition of 0.05% trifluoracetic acid, after lyophilization of the solvent product was used dissolved in 20% TFA in DCM (1 mL), and stirred at room temperature for 1h. After completion of the reaction (as indicated by LCMS), the volatiles were removed under pressure and crude material was purified by reverse phase chromatography using a gradient of 10-65% acetonitrile in water with the addition of 0.05% trifluoracetic acid to give title product 15 mg (65%) as light brown color solid. ^1^H NMR (500 MHz, DMSO-d_6_) δ 11.29 (s, 1H), 9.29 (d, *J* = 58.2 Hz, 1H), 9.00 (d, *J* = 7.8 Hz, 1H), 8.91 – 8.68 (m, 5H), 8.52 (d, *J* = 8.6 Hz, 1H), 8.30 (d, *J* = 4.7 Hz, 1H), 7.84 (t, *J* = 8.0 Hz, 4H), 7.68 (t, *J* = 7.2 Hz, 4H), 7.43 (s, 1H), 7.26 (d, *J* = 7.6 Hz, 1H), 7.12 – 7.06 (m, 1H), 7.04 (t, *J* = 7.2 Hz, 2H), 6.93 (s, 1H), 4.95 – 4.87 (m, 1H), 4.51 (p, *J* = 6.9 Hz, 1H), 4.33 (dt, *J* = 16.0, 7.9 Hz, 2H), 4.05 (d, *J* = 13.6 Hz, 3H), 3.92 (q, *J* = 6.4 Hz, 1H), 3.80 (d, *J* = 6.6 Hz, 1H), 3.60 – 3.19 (m, 15H), 3.15 – 2.87 (m, 6H), 2.75 – 2.62 (m, 2H), 2.12 – 1.96 (m, 8H), 1.94 – 1.39 (m, 20H), 1.26 (d, *J* = 6.9 Hz, 4H), 1.17 – 0.88 (m, 6H).^13^C NMR (126 MHz, DMSO-d_6_) δ 183.2, 171.7, 169.9, 169.1, 166.0, 165.7, 165.2, 160.0, 152.0, 140.0, 137.7, 137.5, 135.7, 135.3, 133.1, 132.9, 131.8, 129.1, 128.8, 128.2, 128.0, 127.2, 126.1, 125.3, 123.6, 123.4, 118.9, 116.7, 116.5, 113.4, 82.5, 75.5, 75.4, 59.0, 56.6, 56.3, 55.9, 55.4, 54.6, 53.3, 51.9, 49.1, 47.3, 42.1, 37.1, 36.6, 34.5, 31.2, 30.2, 30.2, 29.2, 29.0, 28.9, 28.6, 28.3, 26.2, 26.2, 26.0, 24.4, 20.7, 16.2; LC-MS (ESI): (m/z) = 1289 [M+H].

##### EPZ031686-IAP12

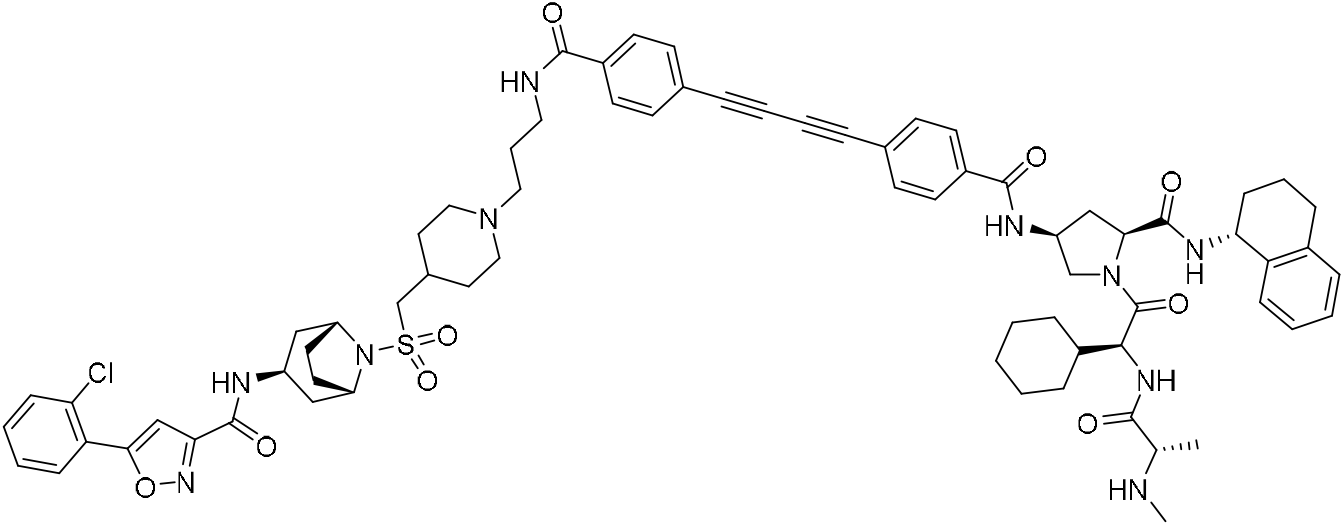

5-(2-chlorophenyl)-N-((1R,3r,5S)-8-(((1-(3-(4-((4-(((3S,5S)-1-((S)-2-cyclohexyl-2-((S)-2-(methylamino)propanamido)acetyl)-5-(((R)-1,2,3,4-tetrahydronaphthalen-1-yl)carbamoyl)pyrrolidin-3-yl)carbamoyl)phenyl)buta-1,3-diyn-1-yl)benzamido)propyl)piperidin-4-yl)methyl)sulfonyl)-8-azabicyclo[3.2.1]octan-3-yl)isoxazole-3-carboxamide **(EPZ031686-IAP12)**: General procedure 2B was followed on 30 mg (0.043 mmol) scale of the N-((1R,5S)-8-(((1-(3-aminopropyl)piperidin-4-yl)methyl)sulfonyl)-8-azabicyclo[3.2.1]octan-3-yl)-6-chloro-3-((Z)-4-cyano-2-fluoro-5-methoxybenzylidene)-2-oxoindoline-5-carboxamide. After the completion of the reaction, crude material was purified by reverse phase chromatography without workup using a gradient of 10-70% acetonitrile in water with the addition of 0.05% TFA to afford 24 mg (43%) of title compound as white solid. ^1^H NMR (500 MHz, DMSO) δ 9.01 (d, *J* = 7.8 Hz, 1H), 8.59 (t, *J* = 5.5 Hz, 1H), 8.52 (d, *J* = 8.7 Hz, 1H), 8.47 (d, *J* = 5.0 Hz, 1H), 8.19 – 8.11 (m, 1H), 7.84 (dt, *J* = 15.6, 7.4 Hz, 5H), 7.70 – 7.61 (m, 5H), 7.57 – 7.46 (m, 2H), 7.30 – 7.22 (m, 2H), 7.12 – 7.00 (m, 3H), 4.91 (q, *J* = 8.2 Hz, 1H), 4.50 (q, *J* = 6.9 Hz, 1H), 4.33 (dt, *J* = 14.5, 8.0 Hz, 2H), 4.08 (s, 2H), 4.05 – 3.96 (m, 2H), 3.54 (dd, *J* = 10.0, 6.2 Hz, 1H), 3.26 – 3.17 (m, 3H), 2.97 (d, *J* = 5.8 Hz, 2H), 2.84 (d, *J* = 11.0 Hz, 2H), 2.68 (d, *J* = 14.4 Hz, 2H), 2.44 (d, *J* = 3.2 Hz, 3H), 2.31 (d, *J* = 7.3 Hz, 2H), 2.08 – 1.85 (m, 10H), 1.78 (d, *J* = 13.9 Hz, 5H), 1.62 (dq, *J* = 26.0, 9.0 Hz, 3H), 1.49 (d, *J* = 10.9 Hz, 2H), 1.25 (d, *J* = 12.0 Hz, 2H), 1.17 – 1.09 (m, 3H), 1.08 – 0.88 (m, 2H); ^13^C NMR (126 MHz, DMSO) δ 171.8, 170.3, 167.6, 165.6, 165.2, 160.0, 158.9, 137.7, 137.5, 135.4, 133.0, 132.9, 132.8, 131.4, 130.5, 128.9, 128.5, 128.1, 128.0, 127.2, 126.1, 125.6, 123.6, 104.2, 59.0, 58.6, 57.6, 56.2, 55.3, 53.3, 47.3, 42.4, 38.4, 36.2, 33.6, 32.2, 31.6, 30.2, 29.2, 28.8, 28.5, 26.6, 26.3, 26.2, 26.0, 20.7, 18.5; LC-MS (ESI): (m/z) = 1287 [M+H].

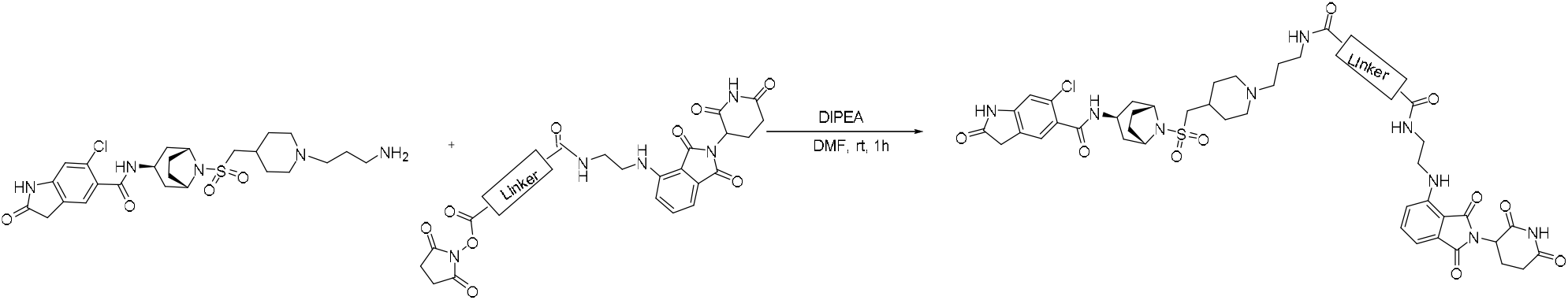

##### General Procedure 2C

A stirred solution of N-((*1R,3r,5S*)-8-(((1-(3-aminopropyl)piperidin-4-yl)methyl)sulfonyl)-8-azabicyclo[3.2.1]octan-3-yl)-6-chloro-2-oxoindoline-5-carboxamide (1.0 eq, 0.028 mmol) in DMF (1 mL), were sequentially added NHS ester of corresponding compound (1.1 eq, 0.031mmol) and DIPEA (2.0 eq, 0.56 mmol) at 0 °C. The resulting mixture was stirred at room temperature for 1h. After completion of the reaction (as indicated by LCMS) the volatiles were removed under vacuum and the crude material was purified by reverse phase chromatography without workup to get pure product.

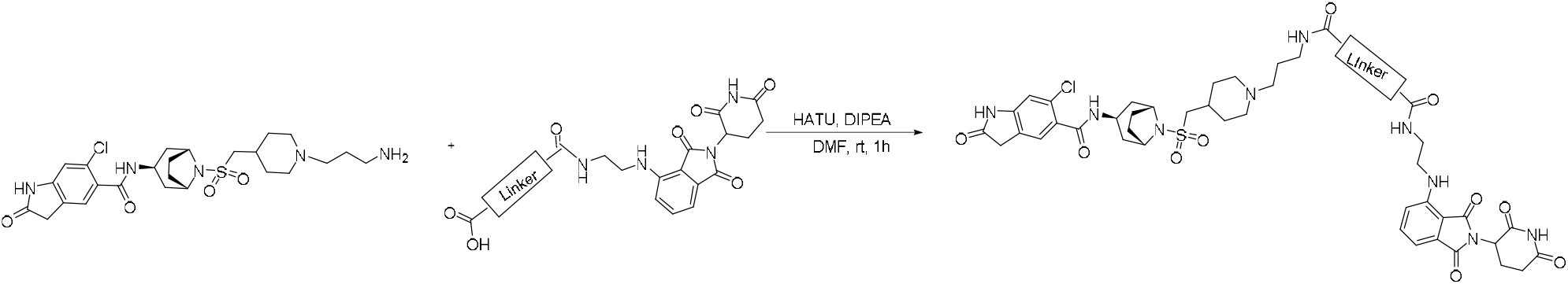

##### General Procedure 2D

A stirred solution of N-((*1R,3r,5S*)-8-(((1-(3-aminopropyl)piperidin-4-yl)methyl)sulfonyl)-8-azabicyclo[3.2.1]octan-3-yl)-6-chloro-2-oxoindoline-5-carboxamide (1.0 eq, 0.028mmol) in DMF (1 mL), were sequentially added corresponding IAP carboxylic acid (1.1 eq, 0.031 mmol), HATU (1.5 eq, 0.042 mmol) and DIPEA (3.0 eq, 0.84 mmol) at 0 °C. The resulting mixture was stirred at room temperature for 3h to 4h. After completion of the reaction (as indicated by LCMS) the volatiles were removed under vacuum and the crude material was purified by reverse phase chromatography without workup to get pure product.

##### CRBN-01

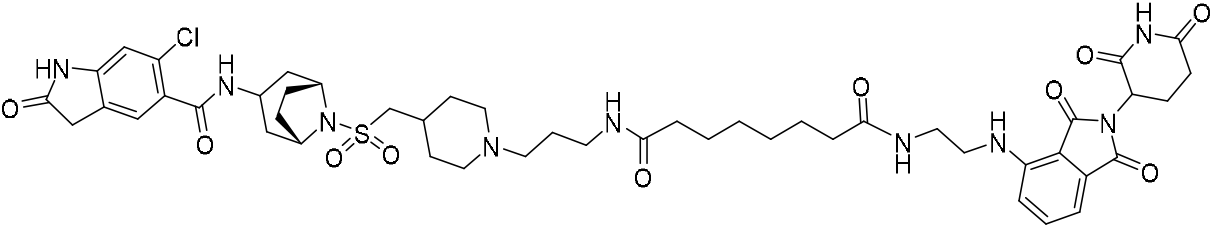

N1-(3-(4-((((1*R*,5*S*)-3-(6-chloro-2-oxoindoline-5-carboxamido)-8-azabicyclo[3.2.1]octan-8-yl)sulfonyl)methyl)piperidin-1-yl)propyl)-N8-(2-((2-(2,6-dioxopiperidin-3-yl)-1,3-dioxoisoindolin-4-yl)amino)ethyl)octanediamide **(EPZ031686-CRBN01):** General procedure 2C was followed on 15 mg (0.028 mmol) scale of the N-((*1R,3r,5S*)-8-(((1-(3-aminopropyl)piperidin-4-yl)methyl)sulfonyl)-8-azabicyclo[3.2.1]octan-3-yl)-6-chloro-2-oxoindoline-5-carboxamide. After the completion of the reaction, crude material was purified by reverse phase chromatography using a gradient of 10-60% acetonitrile in water with the addition of 0.05% TFA to afford 13 mg (47%) of title compound. ^1^H NMR (400 MHz, MeOD-d4) δ 7.56 (dd, *J* = 8.6, 7.1 Hz, 1H), 7.30 (d, *J* = 1.1 Hz, 1H), 7.12 (d, *J* = 8.5 Hz, 1H), 7.06 (dd, *J* = 7.1, 0.6 Hz, 1H), 6.94 (s, 1H), 5.06 (dd, *J* = 12.6, 5.5 Hz, 1H), 4.22 (s, 2H), 4.12 (t, *J* = 6.8 Hz, 1H), 3.55 (q, *J* = 10.4 Hz, 3H), 3.50 – 3.38 (m, 4H), 3.26 (t, *J* = 6.5 Hz, 2H), 3.16 – 2.95 (m, 5H); ^13^C NMR (101 MHz, MeOD-d4) δ 175.4, 173.3, 170.3, 169.2, 168.5, 167.9, 146.8, 135.9, 132.5, 129.5, 124.5, 116.7, 110.7, 110.3, 56.7, 55.6, 54.2, 52.2, 48.8, 42.3, 41.4, 38.3, 36.3, 35.5, 35.4, 30.8, 30.1, 28.9, 28.6, 28.4, 28.1, 25.3, 25.2, 24.2, 22.4; HRMS (ESI) calcd for C_48_H_63_ClN_9_O10_S_^+^ (MH+) 992.4107, found 992.4108.

##### CRBN-02

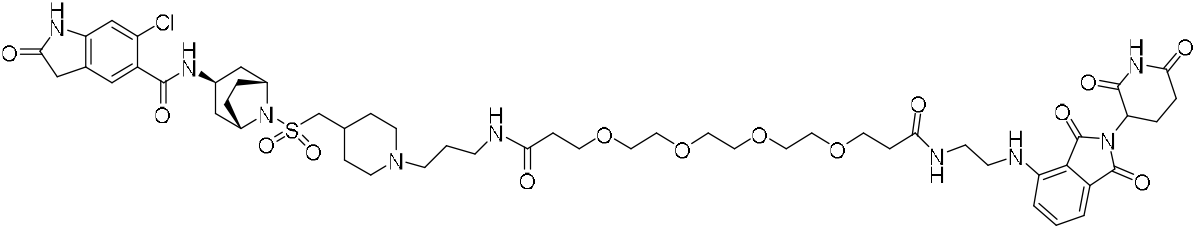

N1-(3-(4-((((*1R,3r,5S*)-3-(6-chloro-2-oxoindoline-5-carboxamido)-8-azabicyclo[3.2.1]octan-8-yl)sulfonyl)methyl)piperidin-1-yl)propyl)-N16-(2-((2-(2,6-dioxopiperidin-3-yl)-1,3-dioxoisoindolin-4-yl)amino)ethyl)-4,7,10,13-tetraoxahexadecanediamide **(EPZ031686-CRBN02).** General procedure 2C was followed on 18 mg (0.033 mmol) scale of the N-((*1R,3r,5S*)-8-(((1-(3-aminopropyl)piperidin-4-yl)methyl)sulfonyl)-8-azabicyclo[3.2.1]octan-3-yl)-6-chloro-2-oxoindoline-5-carboxamide. After the completion of the reaction, crude material was purified by reverse phase chromatography using a gradient of 10-60% acetonitrile in water with the addition of 0.05% TFA to afford 14 mg (38%) of title compound. ^1^H NMR (400 MHz, MeOD-d_4_) δ 7.57 (ddd, *J* = 8.9, 7.1, 1.8 Hz, 1H), 7.30 (d, *J* = 1.7 Hz, 1H), 7.14 (dd, *J* = 8.6, 1.7 Hz, 1H), 7.06 (dd, *J* = 7.0, 1.7 Hz, 1H), 6.94 (d, *J* = 1.7 Hz, 1H), 5.06 (ddd, *J* = 12.4, 5.5, 1.7 Hz, 1H), 4.22 (s, 3H), 4.11 (dt, *J* = 6.8, 4.0 Hz, 1H), 3.83 – 3.66 (m, 5H), 3.58 (t, *J* = 2.2 Hz, 18H), 3.51 – 3.41 (m, 5H), 3.17 – 3.06 (m, 4H), 2.98 (t, *J* = 12.7 Hz, 2H), 2.92 – 2.65 (m, 5H), 2.50 – 2.40 (m, 5H), 2.33 – 2.19 (m, 6H), 2.19 – 1.86 (m, 8H), 1.66 (q, *J* = 13.5 Hz, 2H); ^13^C NMR (101 MHz, MeOD-d4) δ 178.1, 174.1, 173.2, 173.2, 170.2, 169.2, 168.4, 167.8, 146.7, 145.8, 135.9, 132.5, 130.2, 129.5, 124.8, 124.7, 116.8, 110.8, 110.4, 110.0, 70.1, 70.0, 70.0, 69.9, 66.8, 66.7, 56.8, 55.6, 53.7, 52.1, 48.8, 42.3, 41.4, 38.4, 36.3, 36.2, 36.0, 35.2, 30.8, 30.2, 28.9, 28.1, 24.1, 22.4; HRMS (ESI) calcd for C_52_H_71_ClN_9_O_14_S^+^ (MH+) 1112.4530, found 1112.4547.

##### CRBN-03

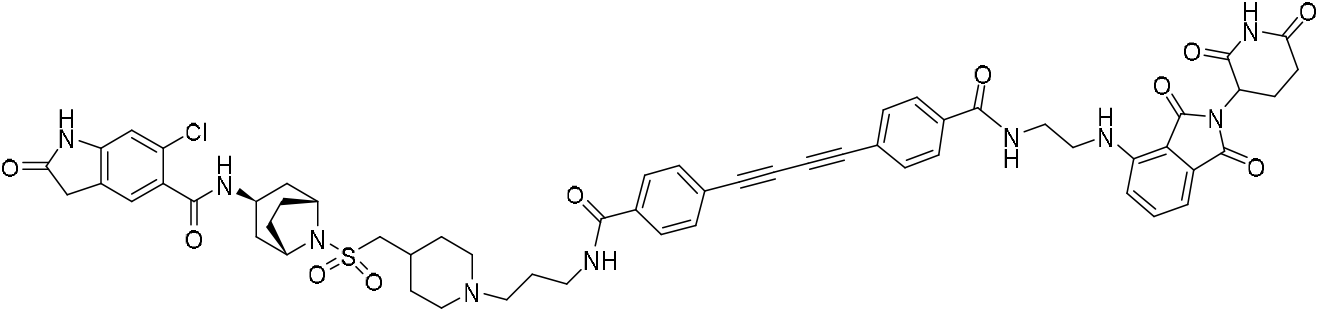

6-chloro-N-((*1R,3r,5S*)-8-(((1-(3-(4-((4-((2-((2-(2,6-dioxopiperidin-3-yl)-1,3-dioxoisoindolin-4-yl)amino)ethyl)carbamoyl)phenyl)buta-1,3-diyn-1-yl)benzamido)propyl)piperidin-4-yl)methyl)sulfonyl)-8-azabicyclo[3.2.1]octan-3-yl)-2-oxoindoline-5-carboxamide **(EPZ031686-CRBN03):** General procedure 2D was followed on 15 mg (0.028mmol) scale of the N-((*1R,3r,5S*)-8-(((1-(3-aminopropyl)piperidin-4-yl)methyl)sulfonyl)-8-azabicyclo[3.2.1]octan-3-yl)-6-chloro-2-oxoindoline-5-carboxamide. After the completion of the reaction, crude material was purified by reverse phase chromatography using a gradient of 10-55% acetonitrile in water with the addition of 0.05% TFA to afford 15 mg (49%) of title compound. ^1^H NMR (500 MHz, DMSO-d_6_) δ 11.04 (s, 1H), 10.57 (s, 1H), 9.16 (s, 1H), 8.75 (dt, *J* = 30.3, 5.7 Hz, 2H), 8.16 (d, *J* = 4.6 Hz, 1H), 7.83 (dd, *J* = 11.0, 8.2 Hz, 4H), 7.67 (dd, *J* = 11.1, 8.1 Hz, 4H), 7.51 (t, *J* = 7.8 Hz, 1H), 7.20 – 7.14 (m, 2H), 6.96 (d, *J* = 7.0 Hz, 1H), 6.78 (d, *J* = 5.5 Hz, 2H), 4.99 (dd, *J* = 12.8, 5.5 Hz, 1H), 4.06 (s, 2H), 3.90 (q, *J* = 6.2 Hz, 1H), 3.41 (d, *J* = 7.0 Hz, 3H), 3.26 (dq, *J* = 15.9, 6.6 Hz, 2H), 3.12 – 2.75 (m, 6H), 2.55 – 2.46 (m, 2H), 2.16 – 1.92 (m, 8H), 1.87 (d, *J* = 16.4 Hz, 7H), 1.46 (t, *J* = 13.1 Hz, 2H); ^13^C NMR (126 MHz, DMSO-d_6_) δ 176.8, 173.2, 170.5, 169.2, 167.8, 167.0, 166.3, 166.1, 158.5, 158.3, 146.9, 146.0, 136.7, 135.8, 135.7, 132.9, 132.7, 130.2, 129.6, 128.1, 125.3, 125.2, 123.5, 123.4, 118.7, 117.7, 116.3, 111.1, 110.2, 109.9, 82.5, 82.4, 75.5, 75.4, 56.7, 55.4, 54.6, 51.9, 49.0, 42.0, 41.8, 37.1, 36.7, 35.8, 31.5, 30.2, 29.0, 28.4, 24.4, 22.7; LC-MS (ESI): (m/z) = 1108 [M+H].

##### CRBN-04

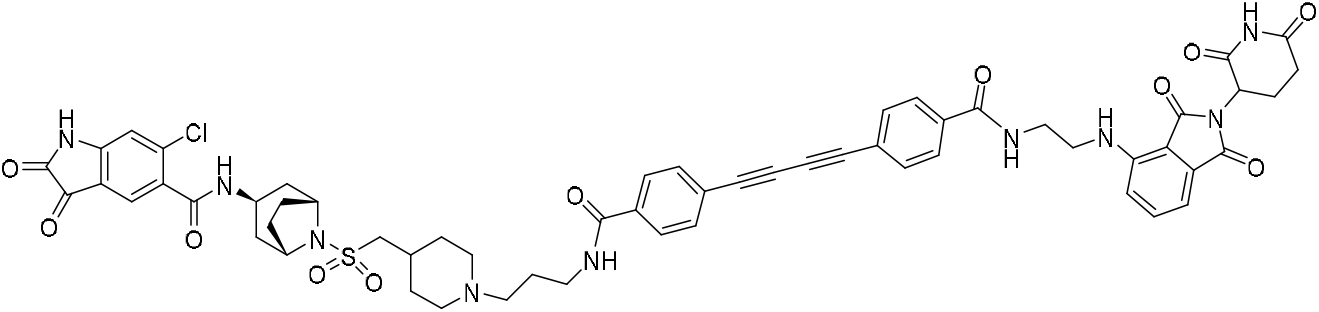

6-chloro-N-((*1R,3r,5S*)-8-(((1-(3-(4-((4-((2-((2-(2,6-dioxopiperidin-3-yl)-1,3-dioxoisoindolin-4-yl)amino)ethyl)carbamoyl)phenyl)buta-1,3-diyn-1-yl)benzamido)propyl)piperidin-4-yl)methyl)sulfonyl)-8-azabicyclo[3.2.1]octan-3-yl)-2,3-dioxoindoline-5-carboxamide **(EPZ031686-CRBN04):** To a stirred solution of the oxindole 6-chloro-N-((*1R,3r,5S*)-8-(((1-(3-(4-((4-((2-((2-(2,6-dioxopiperidin-3-yl)-1,3-dioxoisoindolin-4-yl)amino)ethyl)carbamoyl)phenyl)buta-1,3-diyn-1-yl)benzamido)propyl)piperidin-4-yl)methyl)sulfonyl)-8-azabicyclo[3.2.1]octan-3-yl)-2-oxoindoline-5-carboxamide (8 mg, 1 eq, 7 μmol), in DMF (0.5 mL), were added Cupric acetate, monohydrate (0.1 mg, 0.08 μL, 0.1 eq, 0.7 μmol), and K_2_CO_3_ (0.1 mg, 0.1 eq, 0.7 μmol) at room temperature. After completion of the reaction (as indicated by LCMS), the crude material was purified by reverse phase chromatography using a gradient of 10-65% acetonitrile in water with the addition of 0.05% TFA to give the title compound 5.5 mg (70%) as pale yellow solid. ^1^H NMR (500 MHz, CD_3_OD_SPE) δ 7.85 – 7.75 (m, 4H), 7.64 (d, *J* = 8.4 Hz, 1H), 7.60 (d, *J* = 8.3 Hz, 1H), 7.53 (dd, *J* = 8.6, 7.1 Hz, 1H), 7.19 (d, *J* = 8.7 Hz, 1H), 7.04 (d, *J* = 7.1 Hz, 1H), 6.80 (s, 1H), 5.04 (dd, *J* = 12.6, 5.5 Hz, 1H), 4.20 (s, 2H), 4.08 (s, 1H), 3.67 – 3.54 (m, 6H), 3.50 (dd, *J* = 11.2, 6.0 Hz, 1H), 3.45 – 3.40 (m, 2H), 3.16 (t, *J* = 1.6 Hz, 0H), 3.00 (d, *J* = 5.8 Hz, 2H), 2.86 – 2.79 (m, 1H), 2.73 (d, *J* = 17.8 Hz, 1H), 2.48 (s, 2H), 2.23 (s, 1H), 2.19 – 1.91 (m, 11H), 1.88 (s, 1H), 1.83 (t, *J* = 7.2 Hz, 2H); LC-MS (ESI): (m/z) = 1122 [M+H].

##### CRBN-05

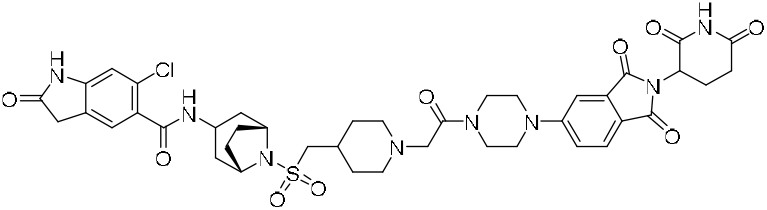

6-chloro-N-((1R,5S)-8-(((1-(2-(4-(2-(2,6-dioxopiperidin-3-yl)-1,3-dioxoisoindolin-5-yl)piperazin-1-yl)-2-oxoethyl)piperidin-4-yl)methyl)sulfonyl)-8-azabicyclo[3.2.1]octan-3-yl)-2-oxoindoline-5-carboxamide **(EPZ031686-CRBN05):**

To a stirred solution of 5-(4-(2-(4-((((1R,5S)-3-amino-8-azabicyclo[3.2.1]octan-8-yl)sulfonyl)methyl)piperidin-1-yl)acetyl)piperazin-1-yl)-2-(2,6-dioxopiperidin-3-yl)isoindoline-1,3-dione (50 mg, 0.074 mmol) in DMF (2 mL), were added 6-chloro-2-oxoindoline-5-carboxylic acid (24 mg, 0.074 mmol), EDCI (28 mg, 1.48 mmol), HOBt (23 mg, 1.48 mmol) and Et_3_N (40 μL, 70.296 mmol) at 0 °C. The resulting mixture was stirred at room temperature for 3h. After completion of the reaction (as indicated by LCMS), the volatiles were removed under vacuum and the crude material was purified by reverse phase chromatography without workup using a gradient of 10-55% acetonitrile in water with the addition of 0.05% trifluoroacetic acid (TFA) to give title compound **3**5 mg (54%) as pale yellow solid. ^1^H NMR (500 MHz, DMSO) δ 11.01 (s, 1H), 10.53 (s, 1H), 8.12 (d, *J* = 4.7 Hz, 1H), 7.63 (d, *J* = 8.5 Hz, 1H), 7.29 (d, *J* = 2.3 Hz, 1H), 7.19 (dd, *J* = 8.7, 2.3 Hz, 1H), 7.15 (s, 1H), 6.76 (s, 1H), 5.01 (dd, *J* = 12.9, 5.4 Hz, 1H), 4.03 (p, *J* = 3.2 Hz, 2H), 3.89 (q, *J* = 6.3 Hz, 1H), 3.65 (t, *J* = 5.1 Hz, 2H), 3.53 (t, *J* = 5.1 Hz, 5H), 3.44 (d, *J* = 6.5 Hz, 3H), 3.38 (t, *J* = 5.3 Hz, 2H), 3.13 (s, 2H), 2.94 (d, *J* = 5.9 Hz, 2H), 2.87 – 2.72 (m, 3H), 2.56 – 2.45 (m, 2H), 2.12 – 1.92 (m, 8H), 1.80 (dd, *J* = 48.6, 13.9 Hz, 4H), 1.26 (q, *J* = 11.9 Hz, 2H). LC-MS (ESI); ^13^C NMR (126 MHz, DMSO) δ 176.8, 173.3, 170.5, 168.2, 168.0, 167.4, 166.9, 155.4, 146.0, 134.3, 130.2, 129.6, 125.4, 125.2, 119.0, 118.4, 110.1, 108.5, 61.4, 57.5, 55.3, 53.1, 49.3, 47.7, 47.1, 44.8, 42.0, 41.2, 36.7, 35.7, 31.9, 31.7, 31.4, 28.3, 22.6; LC-MS (ESI): (m/z) = 863 [M+H].

##### CRBN-06

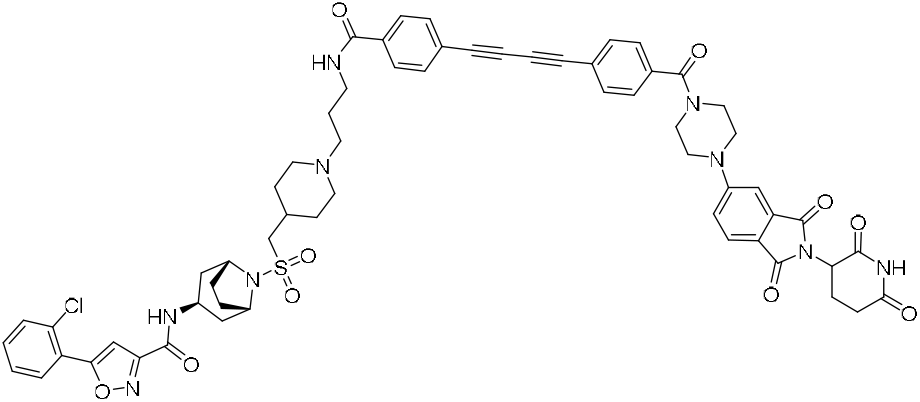

5-(2-chlorophenyl)-N-((1R,3r,5S)-8-(((1-(3-(4-((4-(4-(2-(2,6-dioxopiperidin-3-yl)-1,3-dioxoisoindolin-5-yl)piperazine-1-carbonyl)phenyl)buta-1,3-diyn-1-yl)benzamido)propyl)piperidin-4-yl)methyl)sulfonyl)-8-azabicyclo[3.2.1]octan-3-yl)isoxazole-3-carboxamide **(EPZ031686-CRBN06):** General procedure 2D was followed on 40 mg (0.072 mmol) scale of the N-((1R,3r,5S)-8-(((1-(3-aminopropyl)piperidin-4-yl)methyl)sulfonyl)-8-azabicyclo[3.2.1]octan-3-yl)-5-(2-chlorophenyl)isoxazole-3-carboxamide. After the completion of the reaction, crude material was purified by reverse phase chromatography using a gradient of 10-65% acetonitrile in water with the addition of 0.05% TFA to afford 30 mg (36%) of title compound as pale yellow solid. ^1^H NMR (500 MHz, DMSO) δ 11.02 (s, 1H), 9.26 (s, 1H), 8.49 (d, *J* = 4.9 Hz, 1H), 7.91 – 7.81 (m, 3H), 7.70 – 7.61 (m, 6H), 7.56 – 7.44 (m, 4H), 7.32 – 7.23 (m, 2H), 7.19 (dd, *J* = 8.7, 2.3 Hz, 1H), 5.01 (dd, *J* = 12.7, 5.5 Hz, 1H), 4.09 (s, 2H), 3.98 (s, 1H), 3.70 (s, 2H), 3.44 (d, *J* = 11.8 Hz, 5H), 3.27 (p, *J* = 9.2 Hz, 4H), 3.15 – 2.97 (m, 5H), 2.87 (dt, *J* = 56.5, 13.6 Hz, 2H), 2.56 – 2.44 (m, 2H), 2.01 (qd, *J* = 15.7, 5.7 Hz, 11H), 1.93 – 1.82 (m, 7H), 1.46 (q, *J* = 13.2 Hz, 2H); ^13^C NMR (126 MHz, DMSO) δ 173.2, 168.6, 168.0, 167.6, 167.4, 160.0, 158.9, 155.3, 137.5, 134.4, 133.1, 132.9, 132.8, 131.4, 130.5, 128.5, 128.1, 125.5, 123.5, 119.2, 118.5, 108.6, 104.2, 56.7, 55.4, 51.9, 42.4, 36.2, 31.5, 30.2, 29.0, 28.7, 22.6; LC-MS (ESI): (m/z) = 1146 [M+H].

##### VHL-01

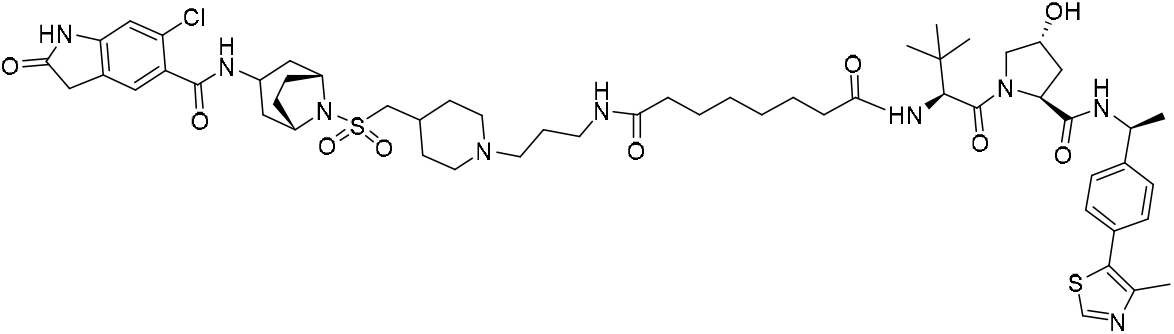

N1-(3-(4-((((1*R*,5*S*)-3-(6-chloro-2-oxoindoline-5-carboxamido)-8-azabicyclo[3.2.1]octan-8-yl)sulfonyl)methyl)piperidin-1-yl)propyl)-N8-((*S*)-1-((2*S*,4*R*)-4-hydroxy-2-(((S)-1-(4-(4-methylthiazol-5-yl)phenyl)ethyl)carbamoyl)pyrrolidin-1-yl)-3,3-dimethyl-1-oxobutan-2-yl)octanediamide **(EPZ031686-VHL01):** General procedure 2C was followed on 18 mg (0.033 mmol) scale of the N-((*1R,3r,5S*)-8-(((1-(3-aminopropyl)piperidin-4-yl)methyl)sulfonyl)-8-azabicyclo[3.2.1]octan-3-yl)-6-chloro-2-oxoindoline-5-carboxamide. After the completion of the reaction, crude material was purified by reverse phase chromatography using a gradient of 10-60% acetonitrile in water with the addition of 0.05% TFA to afford 16 mg (43%) of title compound. ^1^H NMR (400 MHz, MeOD-d_4_) δ 8.88 (d, *J* = 2.4 Hz, 1H), 8.51 (d, *J* = 7.5 Hz, 1H), 8.28 (d, *J* = 5.2 Hz, 1H), 7.78 (d, *J* = 9.0 Hz, 1H), 7.46 – 7.36 (m, 4H), 7.30 (s, 1H), 6.94 (s, 1H), 5.00 (t, *J* = 6.9 Hz, 1H), 4.66 – 4.52 (m, 2H), 4.43 (s, 1H), 4.24 (s, 2H), 3.87 (d, *J* = 11.1 Hz, 1H), 3.75 (dd, *J* = 11.0, 4.0 Hz, 1H), 3.55 (d, *J* = 12.5 Hz, 4H), 3.27 (d, *J* = 6.6 Hz, 2H), 3.18 – 2.94 (m, 5H), 2.47 (d, *J* = 1.1 Hz, 4H), 2.34 – 2.13 (m, 6H), 2.11 – 1.87 (m, 7H), 1.71 – 1.55 (m, 7H), 1.50 (d, *J* = 7.0 Hz, 3H), 1.38 – 1.31 (m, 5H), 1.04 (d, *J* = 1.1 Hz, 9H); LC-MS (ESI): (m/z) = 1120 [M+H].

##### VHL-02

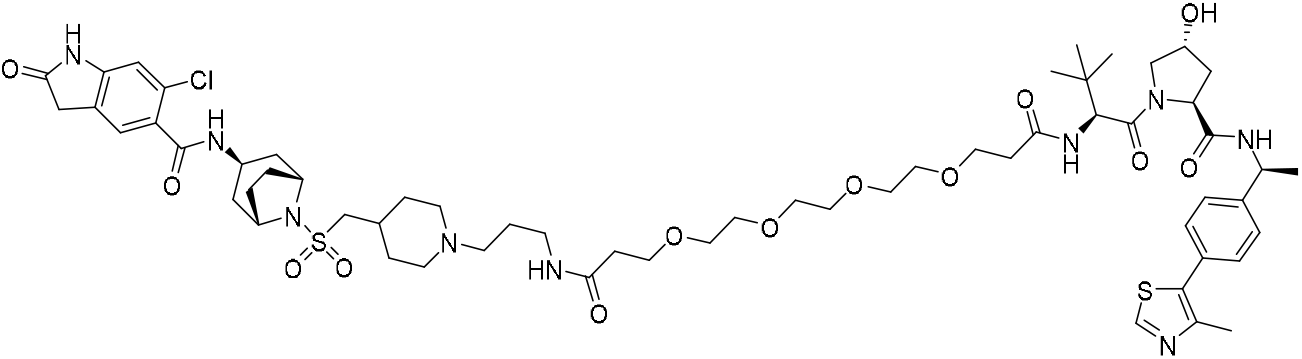

N1-(3-(4-((((*1R,3r,5S*)-3-(6-chloro-2-oxoindoline-5-carboxamido)-8-azabicyclo[3.2.1]octan-8-yl)sulfonyl)methyl)piperidin-1-yl)propyl)-N16-((*S*)-1-((2*S,*4*R*)-4-hydroxy-2-(((*S*)-1-(4-(4-methylthiazol-5-yl)phenyl)ethyl)carbamoyl)pyrrolidin-1-yl)-3,3-dimethyl-1-oxobutan-2-yl)-4,7,10,13-tetraoxahexadecanediamide **(EPZ031686-VHL02):** General procedure 2C was followed on 20 mg (0.037 mmol) scale of the N-((*1R,3r,5S*)-8-(((1-(3-aminopropyl)piperidin-4-yl)methyl)sulfonyl)-8-azabicyclo[3.2.1]octan-3-yl)-6-chloro-2-oxoindoline-5-carboxamide. After the completion of the reaction, crude material was purified by reverse phase chromatography using a gradient of 10-60% acetonitrile in water with the addition of 0.05% TFA to afford 18 mg (39%) of title compound. ^1^H NMR (400 MHz, MeOD-d_4_) δ 8.94 (s, 1H), 7.51 – 7.36 (m, 4H), 7.30 (d, *J* = 1.3 Hz, 1H), 6.94 (d, *J* = 1.4 Hz, 1H), 5.00 (q, *J* = 7.0 Hz, 1H), 4.64 (d, *J* = 1.3 Hz, 1H), 4.62 – 4.52 (m, 1H), 4.43 (s, 1H), 4.24 (s, 2H), 4.18 – 4.07 (m, 1H), 3.86 (d, *J* = 11.0 Hz, 1H), 3.79 – 3.69 (m, 5H), 3.62 (dq, *J* = 3.6, 1.8 Hz, 14H), 3.34 (d, *J* = 6.4 Hz, 2H), 3.19 – 3.08 (m, 3H), 3.00 (t, *J* = 12.8 Hz, 2H), 2.67 – 2.55 (m, 1H), 2.55 – 2.43 (m, 6H), 2.36 – 1.87 (m, 13H), 1.68 (q, *J* = 13.6 Hz, 2H), 1.50 (dd, *J* = 7.0, 1.4 Hz, 3H), 1.08 – 1.01 (m, 9H); ^13^C NMR (101 MHz, MeOD-d_4_) δ 178.0, 174.1, 172.2, 171.8, 170.7, 168.5, 151.7, 145.8, 144.3, 130.2, 130.0, 129.5, 129.1, 126.2, 124.8, 124.7, 110.4, 70.2, 70.1, 70.1, 70.1, 70.0, 69.9, 69.5, 66.9, 66.8, 59.2, 57.6, 56.9, 56.6, 55.6, 53.8, 52.1, 48.7, 42.3, 37.4, 36.3, 36.1, 35.9, 35.3, 35.2, 30.2, 28.9, 28.1, 25.7, 24.1, 21.0, 14.3; HRMS (ESI) calcd for C_60_H_87_ClN_9_O_13_S_2_^+^ (MH+) 1240.5553, found 1240.5575.

##### VHL-03

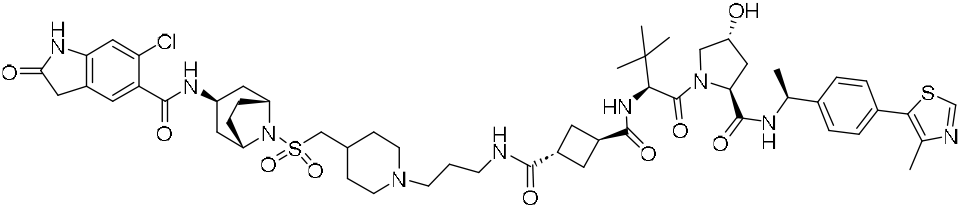

(1S,3R)-N1-(3-(4-((((*1R,3r,5S*)-3-(6-chloro-2-oxoindoline-5-carboxamido)-8-azabicyclo[3.2.1]octan-8-yl)sulfonyl)methyl)piperidin-1-yl)propyl)-N3-((*S*)-1-((2*S*,4*R*)-4-hydroxy-2-(((*S*)-1-(4-(4-methylthiazol-5-yl)phenyl)ethyl)carbamoyl)pyrrolidin-1-yl)-3,3-dimethyl-1-oxobutan-2-yl)cyclobutane-1,3-dicarboxamide **(EPZ031686-VHL-03):** General procedure 2D was followed on 15 mg (0.028mmol) scale of the N-((*1R,3r,5S*)-8-(((1-(3-aminopropyl)piperidin-4-yl)methyl)sulfonyl)-8-azabicyclo[3.2.1]octan-3-yl)-6-chloro-2-oxoindoline-5-carboxamide. After the completion of the reaction, crude material was purified by reverse phase chromatography using a gradient of 10-55% acetonitrile in water with the addition of 0.05% TFA to afford 11 mg (37%) of title compound. ^1^H NMR (400 MHz, MeOD-d_4_) δ 8.53 (d, *J* = 7.4 Hz, 1H), 8.28 (d, *J* = 5.2 Hz, 1H), 7.71 (d, *J* = 9.0 Hz, 1H), 7.52 – 7.36 (m, 4H), 7.30 (d, *J* = 1.6 Hz, 1H), 6.94 (d, *J* = 1.4 Hz, 1H), 4.99 (q, *J* = 6.9 Hz, 1H), 4.64 (dd, *J* = 7.4, 2.3 Hz, 1H), 4.56 (t, *J* = 8.3 Hz, 1H), 4.44 (s, 1H), 4.24 (s, 2H), 4.13 (s, 1H), 3.88 (d, *J* = 11.1 Hz, 1H), 3.83 – 3.72 (m, 1H), 3.59 (d, *J* = 12.7 Hz, 2H), 3.54 (s, 2H), 3.27 (d, *J* = 7.1 Hz, 3H), 3.18 – 2.95 (m, 6H), 2.58 – 1.87 (m, 21H), 1.66 (dt, *J* = 24.3, 10.6 Hz, 2H), 1.50 (dd, *J* = 7.1, 1.4 Hz, 3H), 1.03 (d, *J* = 1.5 Hz, 9H); ^13^C NMR (101 MHz, MeOD-d_4_) δ 177.3, 176.0, 171.8, 170.8, 145.8, 144.3, 130.2, 129.1, 126.2, 126.0, 124.7, 110.4, 69.6, 59.2, 57.7, 56.9, 56.6, 55.6, 54.3, 52.1, 48.7, 42.3, 37.4, 36.3, 36.0, 35.7, 35.5, 35.2, 30.2, 28.8, 28.1, 27.6, 27.0, 25.6, 24.2, 21.0, 14.4; LC-MS (ESI): (m/z) = 1090 [M+H].

##### VHL-04

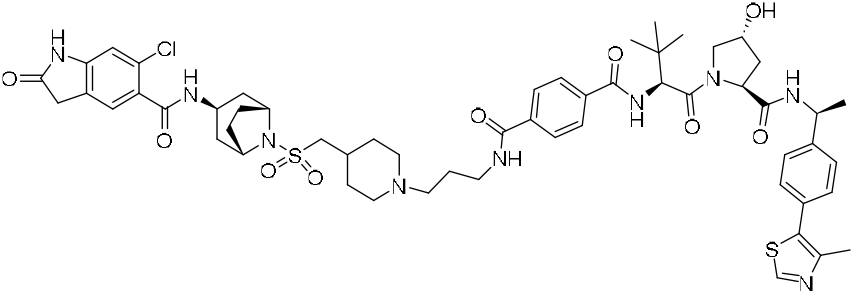

N1-(3-(4-((((*1R,3r,5S*)-3-(6-chloro-2-oxoindoline-5-carboxamido)-8-azabicyclo[3.2.1]octan-8-yl)sulfonyl)methyl)piperidin-1-yl)propyl)-N4-((*<u>S</u>*)-1-((2*S*,4*R*)-4-hydroxy-2-(((*S*)-1-(4-(4-methylthiazol-5-yl)phenyl)ethyl)carbamoyl)pyrrolidin-1-yl)-3,3-dimethyl-1-oxobutan-2-yl)terephthalamide **(EPZ031686-VHL04).** General procedure 2D was followed on 15 mg (0.033mmol) scale of the N-((*1R,3r,5S*)-8-(((1-(3-aminopropyl)piperidin-4-yl)methyl)sulfonyl)-8-azabicyclo[3.2.1]octan-3-yl)-6-chloro-2-oxoindoline-5-carboxamide. After the completion of the reaction, crude material was purified by reverse phase chromatography using a gradient of 10-55% acetonitrile in water with the addition of 0.05% TFA to afford 8 mg (20%) of title compound. ^1^H NMR (400 MHz, MeOD-d_4_) δ 8.98 (s, 1H), 8.59 (d, *J* = 7.5 Hz, 1H), 8.28 (d, *J* = 5.3 Hz, 1H), 8.02 (d, *J* = 9.0 Hz, 1H), 7.98 – 7.89 (m, 5H), 7.53 – 7.35 (m, 5H), 7.30 (s, 1H), 6.94 (d, *J* = 2.0 Hz, 1H), 5.11 – 4.97 (m, 1H), 4.95 – 4.88 (m, 2H), 4.60 (t, *J* = 8.5 Hz, 1H), 4.47 (s, 1H), 4.24 (s, 2H), 4.13 (s, 1H), 3.95 (d, *J* = 11.1 Hz, 1H), 3.81 (d, *J* = 10.8 Hz, 1H), 3.64 (d, *J* = 12.1 Hz, 2H), 3.52 (d, *J* = 8.0 Hz, 4H), 3.23 – 3.13 (m, 3H), 3.05 (t, *J* = 12.8 Hz, 2H), 2.48 (s, 3H), 2.35 – 1.90 (m, 19H), 1.67 (dt, *J* = 21.5, 10.5 Hz, 3H), 1.51 (dd, *J* = 7.1, 2.0 Hz, 3H), 1.20 – 1.05 (m, 9H); ^13^C NMR (101 MHz, MeOD-d_4_) δ 171.7, 170.7, 168.5, 167.6, 145.8, 144.3, 137.0, 136.5, 130.2, 129.5, 129.1, 127.4, 127.2, 126.2, 126.1, 124.6, 110.3, 69.6, 59.3, 58.2, 56.9, 55.5, 54.5, 52.2, 48.7, 42.3, 37.4, 36.4, 36.3, 35.8, 30.2, 28.9, 28.2, 25.7, 24.2, 21.0, 14.3; HRMS (ESI) calcd for C_56_H_71_ClN_9_O_9_S_2_^+^ (MH+) 1112.4505, found 1112.4497.

##### VHL-05

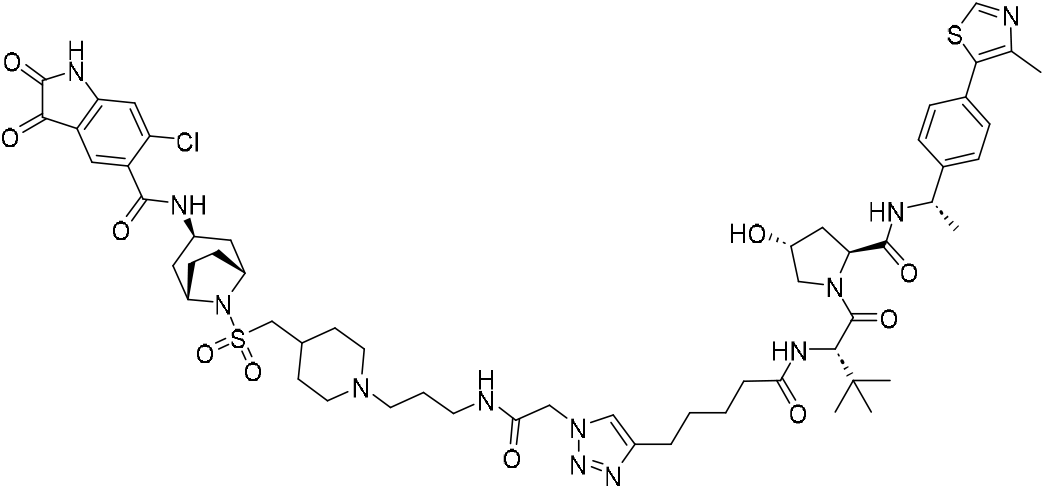

6-chloro-N-((*1R,3r,5S*)-8-(((1-(3-(2-(4-(5-(((*S*)-1-((2S,4R)-4-hydroxy-2-(((*S*)-1-(4-(4-methylthiazol-5-yl)phenyl)ethyl)carbamoyl)pyrrolidin-1-yl)-3,3-dimethyl-1-oxobutan-2-yl)amino)-5-oxopentyl)-1H-1,2,3-triazol-1-yl)acetamido)propyl)piperidin-4-yl)methyl)sulfonyl)-8-azabicyclo[3.2.1]octan-3-yl)-2,3-dioxoindoline-5-carboxamide **(EPZ031686-VHL05):** To a stirred solution of N-((*1R,3r,5S*)-8-(((1-(3-(2-azidoacetamido)propyl)piperidin-4-yl)methyl)sulfonyl)-8-azabicyclo[3.2.1]octan-3-yl)-6-chloro-2-oxoindoline-5-carboxamide (10 mg, 1 eq, 16 μmol)in DMSO (1 mL), were added (2*S*,4*R*)-1-((*S*)-2-(hept-6-ynamido)-3,3-dimethylbutanoyl)-4-hydroxy-N-((*S*)-1-(4-(4-methylthiazol-5-yl)phenyl)ethyl)pyrrolidine-2-carboxamide (8.9 mg, 1 eq, 16 μmol), CuI (0.31 mg, 0.1 eq, 1.6 μmol) and Et_3_N (1.6 mg, 2.2 μL, 1 eq, 16 μmol) at room temperature. The resulting solution was stirred at room temperature for 5h. After completion of the reaction, the crude material was purified by reverse phase chromatography using a gradient of 10-55% acetonitrile in water with the addition of 0.05% TFA to afford 10 mg (52%) of title compound. ^1^H NMR (400 MHz, dmso) δ 11.31 (s, 1H), 9.33 (d, *J* = 52.0 Hz, 1H), 8.97 (s, 1H), 8.44 (t, *J* = 5.9 Hz, 1H), 7.83 – 7.73 (m, 2H), 7.48 (s, 1H), 7.42 (d, *J* = 8.0 Hz, 2H), 7.36 (d, *J* = 8.1 Hz, 2H), 6.97 (s, 1H), 5.01 (s, 2H), 4.90 (t, *J* = 7.2 Hz, 1H), 4.50 (d, *J* = 9.2 Hz, 1H), 4.40 (t, *J* = 7.9 Hz, 1H), 4.26 (s, 1H), 4.11 (s, 3H), 3.97 (s, 2H), 3.58 (s, 2H), 3.44 (d, *J* = 11.8 Hz, 2H), 3.27 (d, *J* = 7.4 Hz, 1H), 3.12 (dd, *J* = 21.2, 6.1 Hz, 4H), 3.01 – 2.90 (m, 4H), 2.59 (d, *J* = 7.2 Hz, 3H), 2.44 (s, 3H), 2.28 (t, *J* = 10.9 Hz, 1H), 2.18 – 1.71 (m, 11H), 1.50 (dd, *J* = 24.5, 11.2 Hz, 6H), 1.36 (d, *J* = 7.0 Hz, 3H), 0.92 (s, 9H); ^13^C NMR (101 MHz, dmso) δ 183.2, 172.4, 171.1, 170.0, 166.3, 165.7, 159.9, 151.9, 148.2, 147.0, 145.1, 140.0, 131.7, 131.6, 130.1, 129.3, 126.8, 125.3, 123.7, 116.7, 113.4, 69.2, 59.0, 56.8, 56.7, 55.4, 54.3, 52.0, 51.9, 48.1, 42.1, 38.2, 36.6, 35.6, 35.0, 30.1, 29.0, 29.0, 28.3, 26.9, 25.4, 25.2, 24.2, 22.9, 16.4; LC-MS (ESI): (m/z) = 1187 [M+H].

## SUPPLEMENTAL INFORMATION

### Supplementary Table titles and legends

**Supplementary Table 1. Gene sets of cell cycle-, EMT-, type I IFN-response genes of the respective HALLMARK pathways. (A)** GSEA gene list of cell cycle-related genes (252 genes). **(B)** Gene set of the EMT Hallmark pathway (200 genes). **(C)** Gene sets of the type I IFN-response Hallmark pathway (97 genes).

## KEY RESOURCES TABLE

xxx

## DECLARATION OF INTERESTS

JA, SRD, RW and VS are co-inventors of the presented SMYD3 PROTACs and a patent has been filed (Patent application ID: 325900-8020). The other authors have no other relevant affiliations or financial involvement with any organization or entity with a financial interest in or financial conflict with the subject matter or materials discussed in the manuscript. This includes employment, consultancies, honoraria, stock ownership or options, expert testimony, grants, patents filed or pending or royalties.

## INCLUSION AND DIVERSITY

We support inclusive, diverse and equitable conduct of research.

## Notes

### Competing Interest Statement

The authors have declared no competing interest.

