## Supplementary Figure 1 to 4 for "SMYD3 protein degraders in HPV-negative head and neck squamous cell carcinoma"

**Supplementary Figures.**

**Supplementary Figure 1.** SMYD3 (top) and XIAP (bottom) protein expression levels in HPV-negative HNSCC cell lines and the normalized BEAS-2B epithelial cells. 20 ug and 30ug of cytoplasmic extracts were blotted for SMYD3 and XIAP respectively. Actin was used as a loading control.

FaDu

HN-6

HN13

PE/CA-PJ15

HN-SCC-151

YD-10B

BEAS-2B

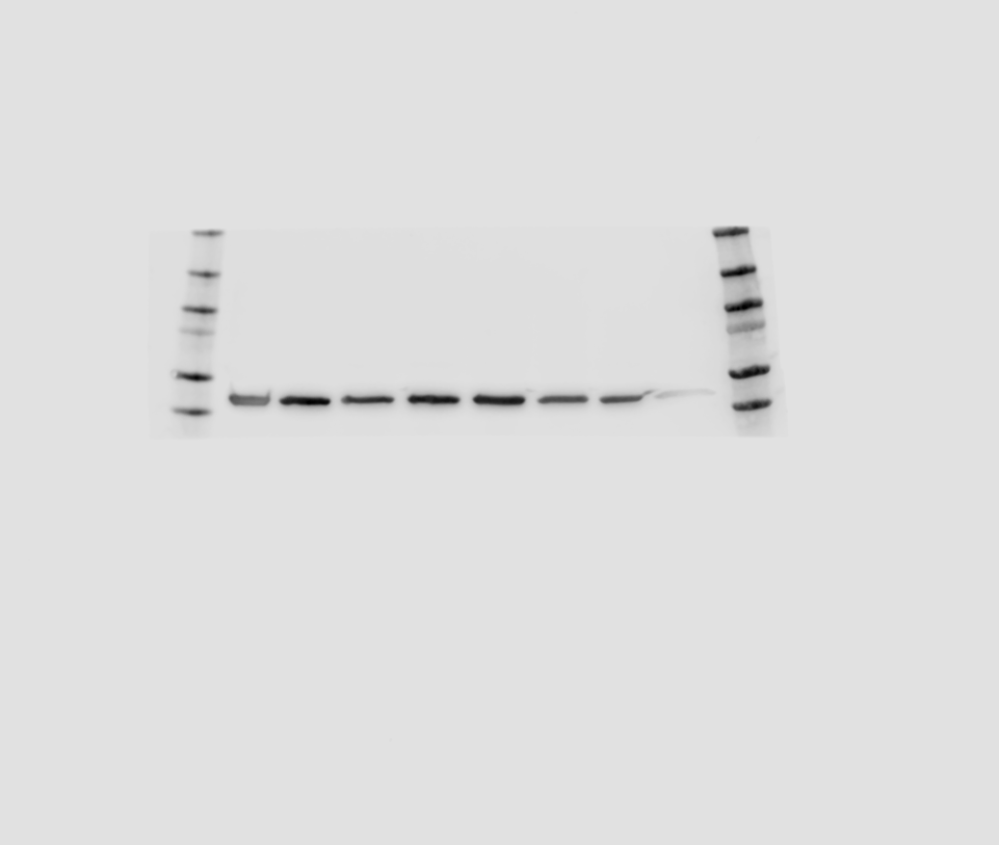

Actin

SMYD3

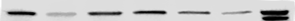

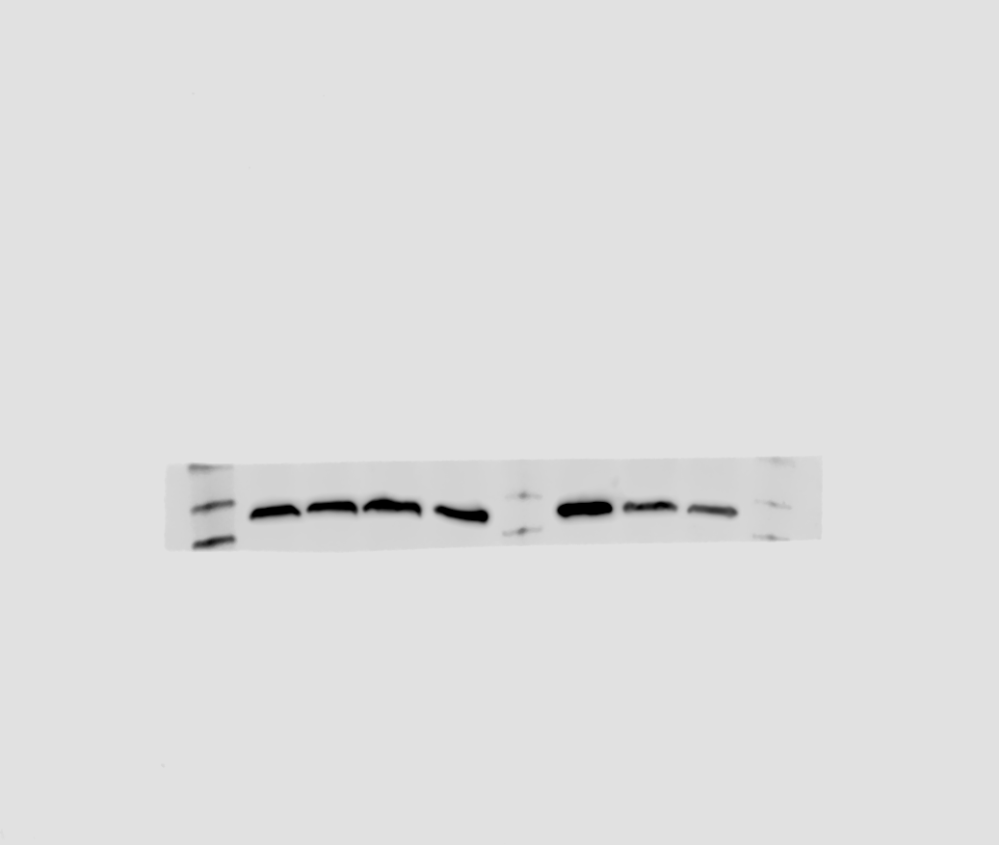

GAPDH

HN13

BEAS-2B

HEK293T

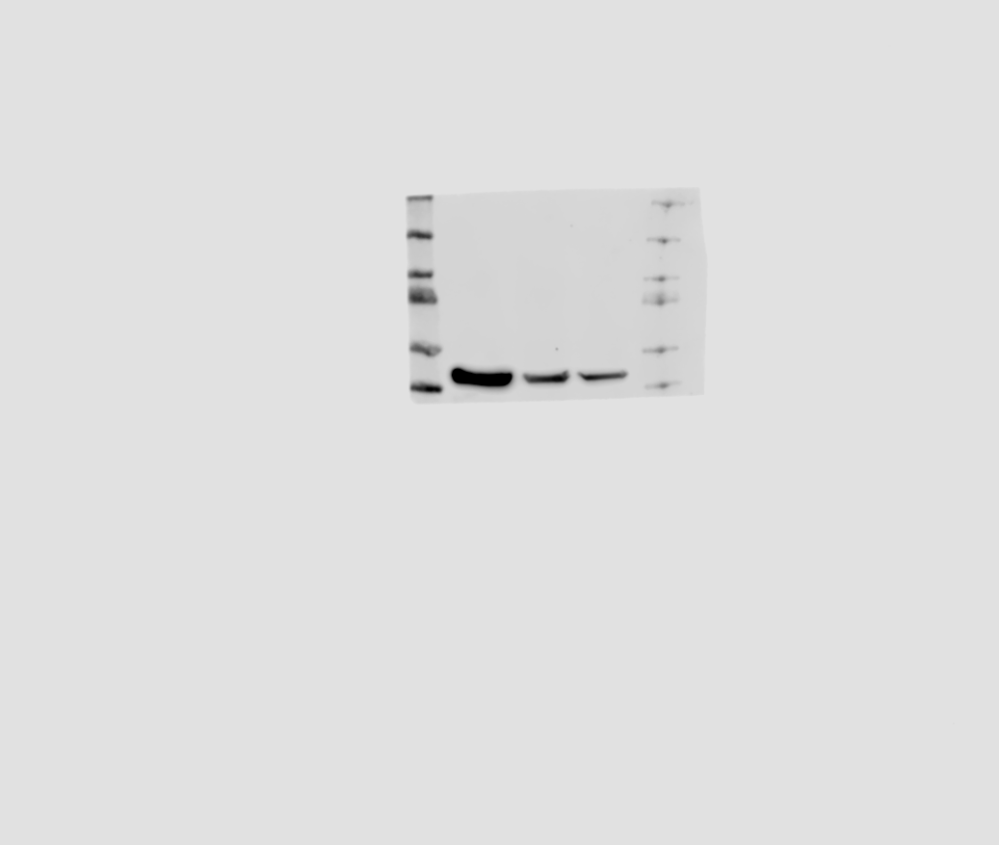

XIAP

**Supplementary Figure 2.** Western blotting for pERK, ERK and H3K4me3 in HPV-negative HNSCC cells (HN-6, HN-SCC-151, PE/CA-PJ15) treated with BAY-6035 at incremental concentrations for 6 days. 20ug of cytoplasmic extracts and 5ug of nuclear extracts were loaded for pERK and H3K4me3 respectively. H3K4me3 band intensities were normalized by H3 and pERK band intensities were normalized by ERK. Standard deviations are shown from two biological replicates.H3 and actin were used as loading controls.

DMSO

1

2.5

5

10

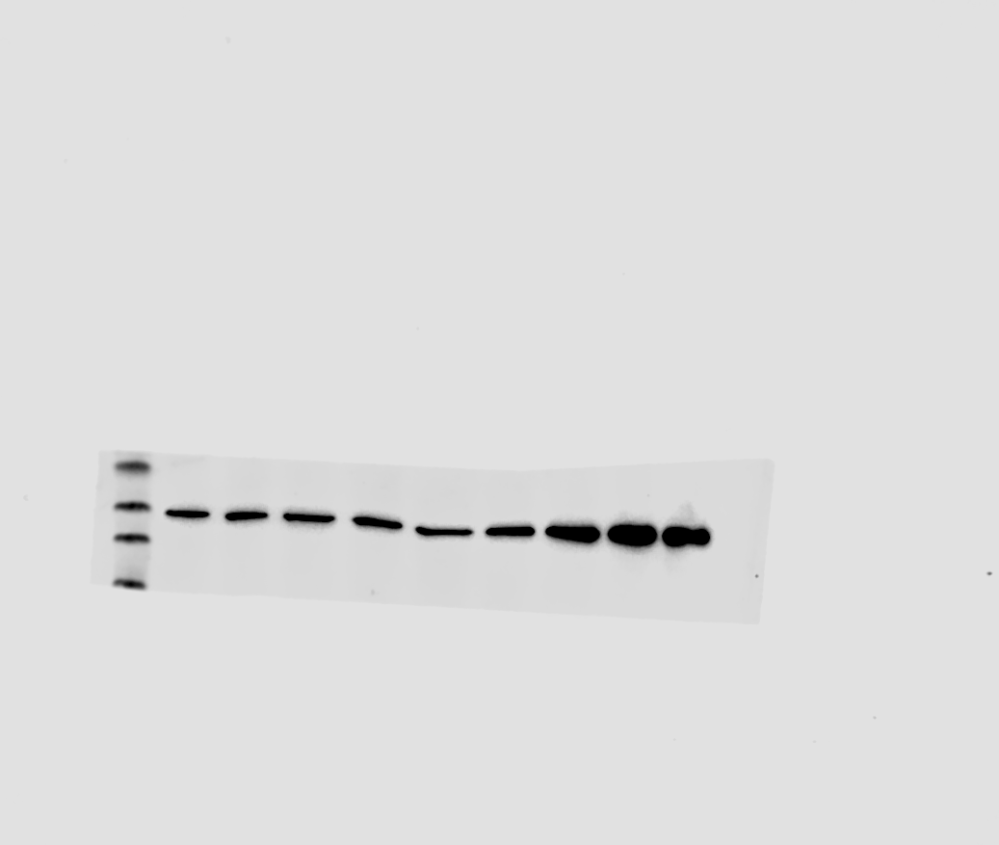

**PE/CA-PJ15**

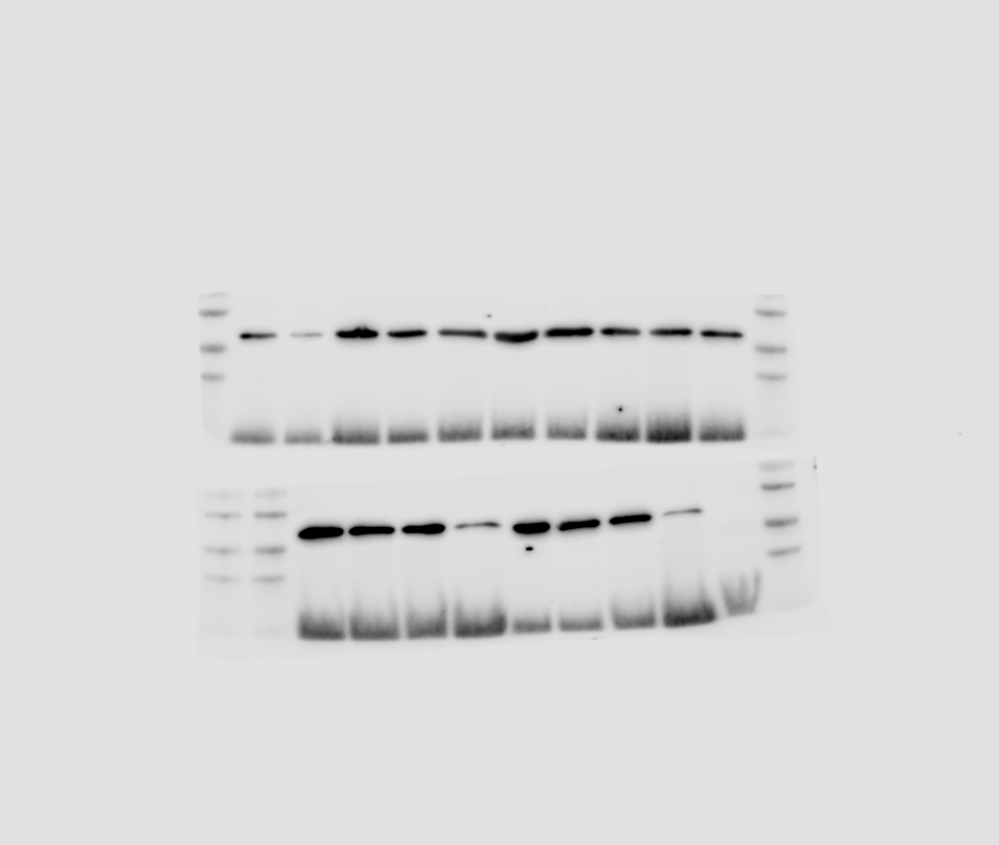

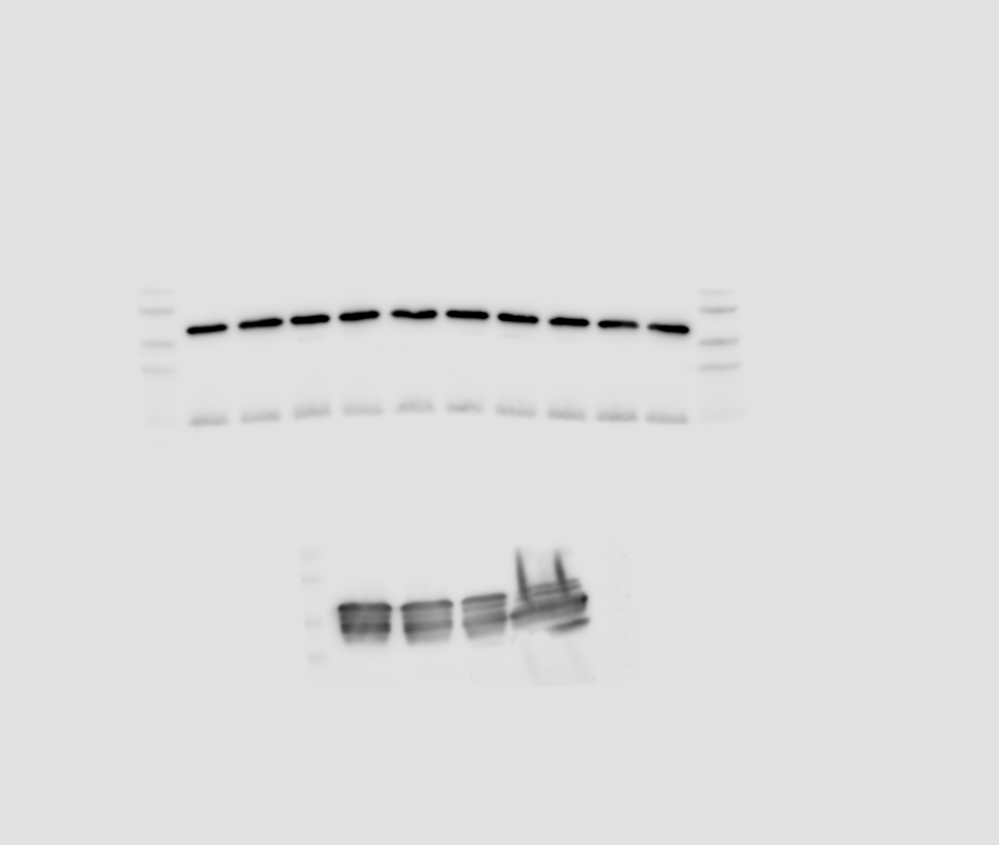

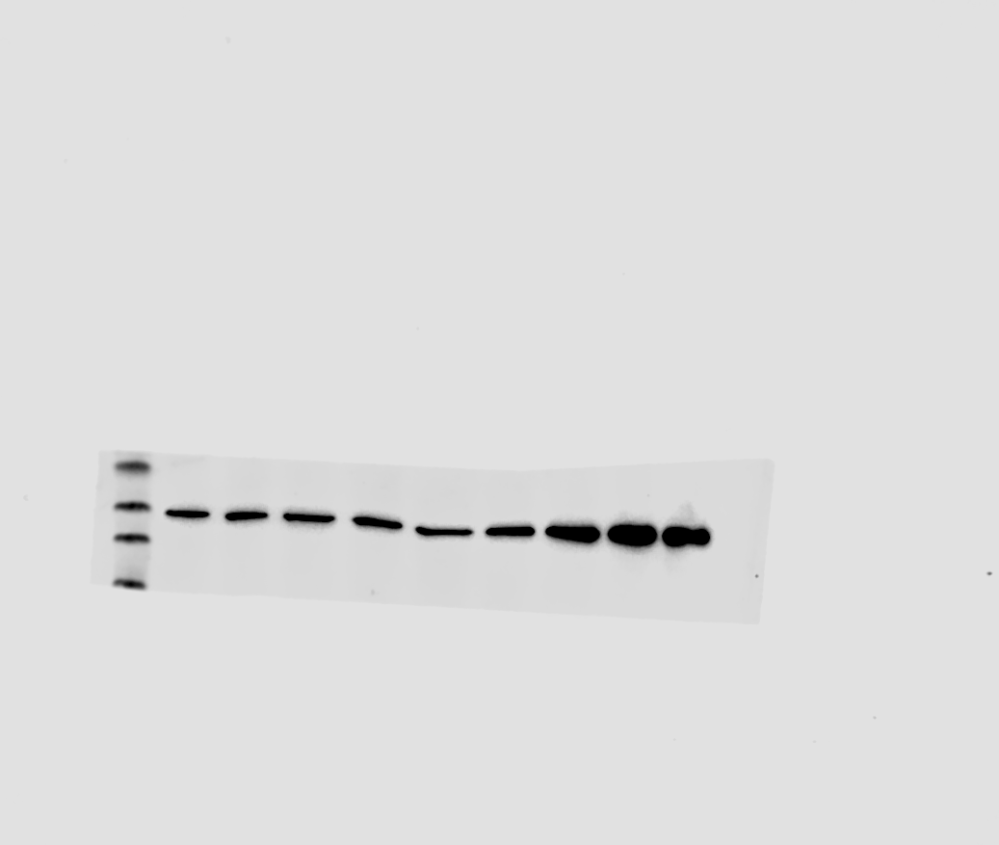

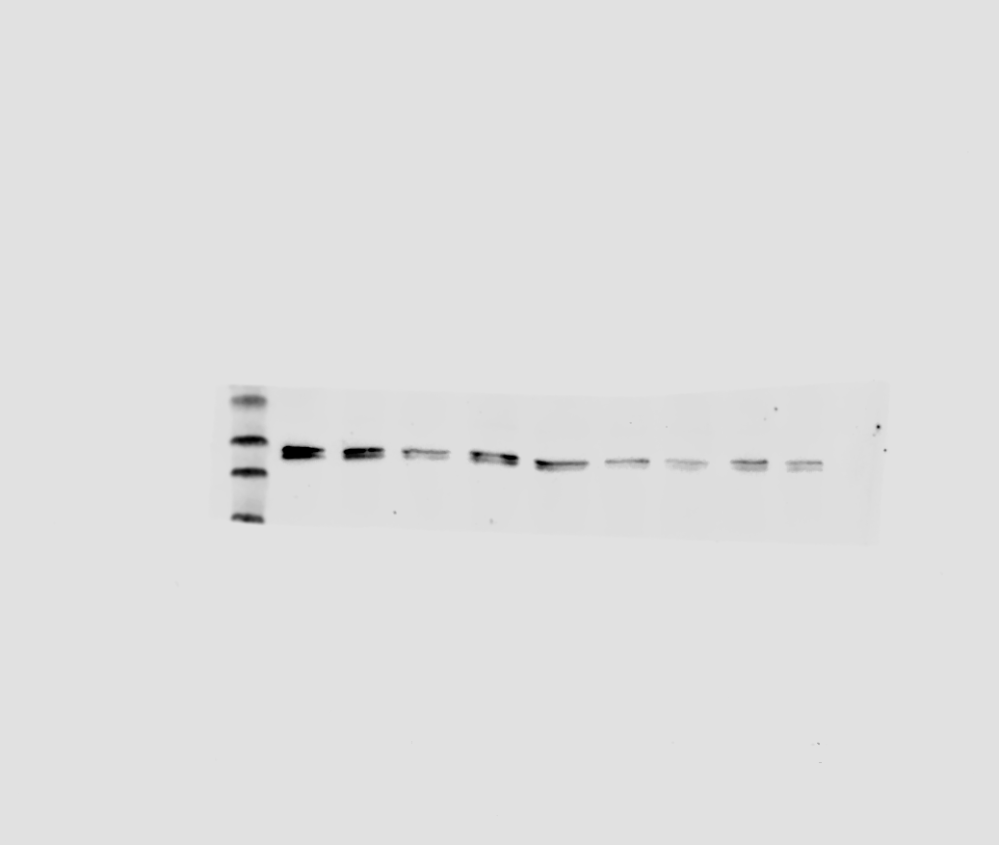

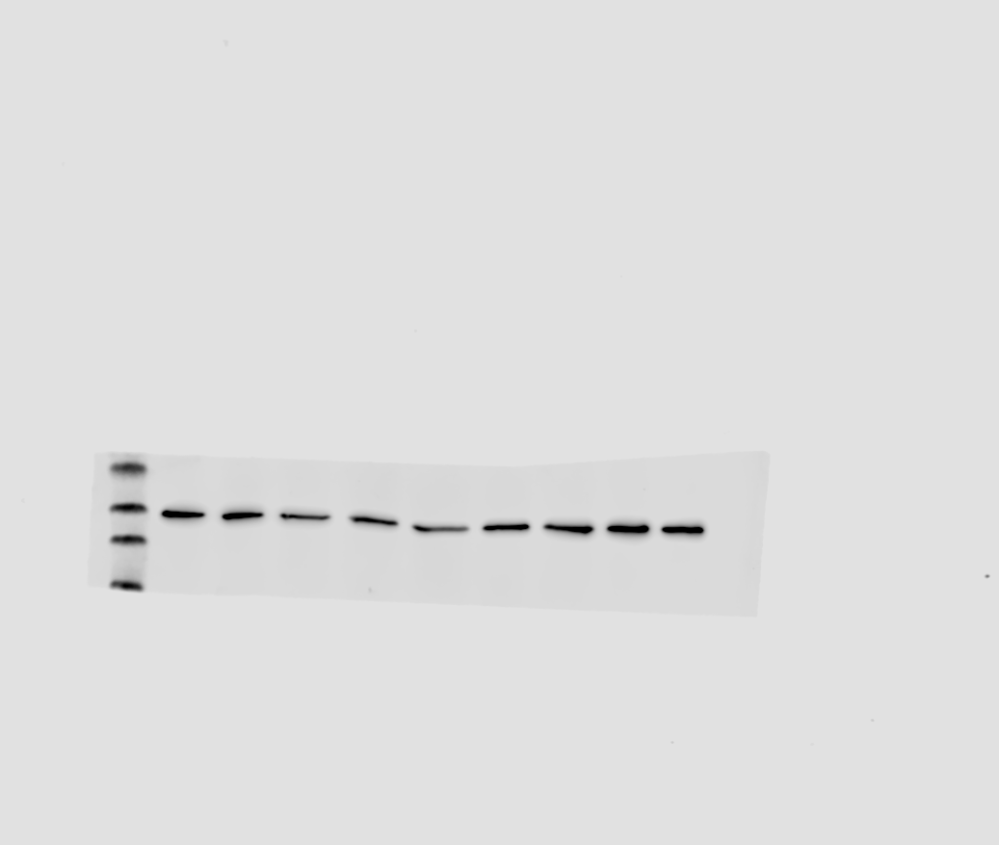

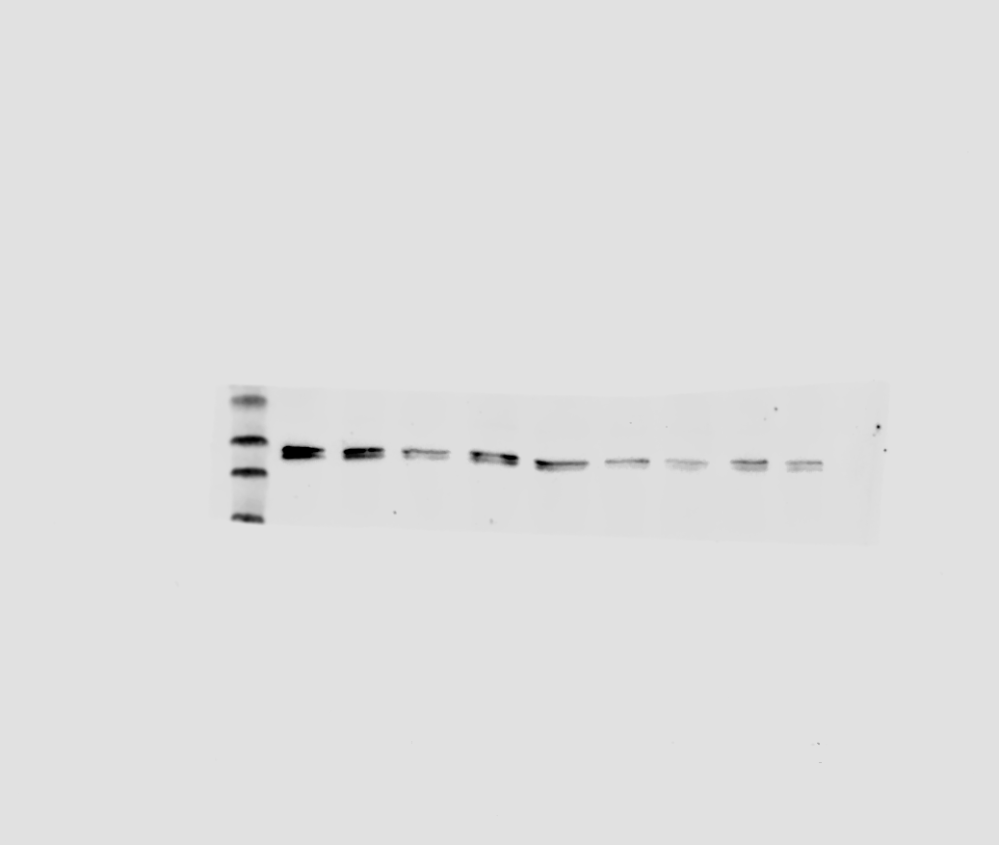

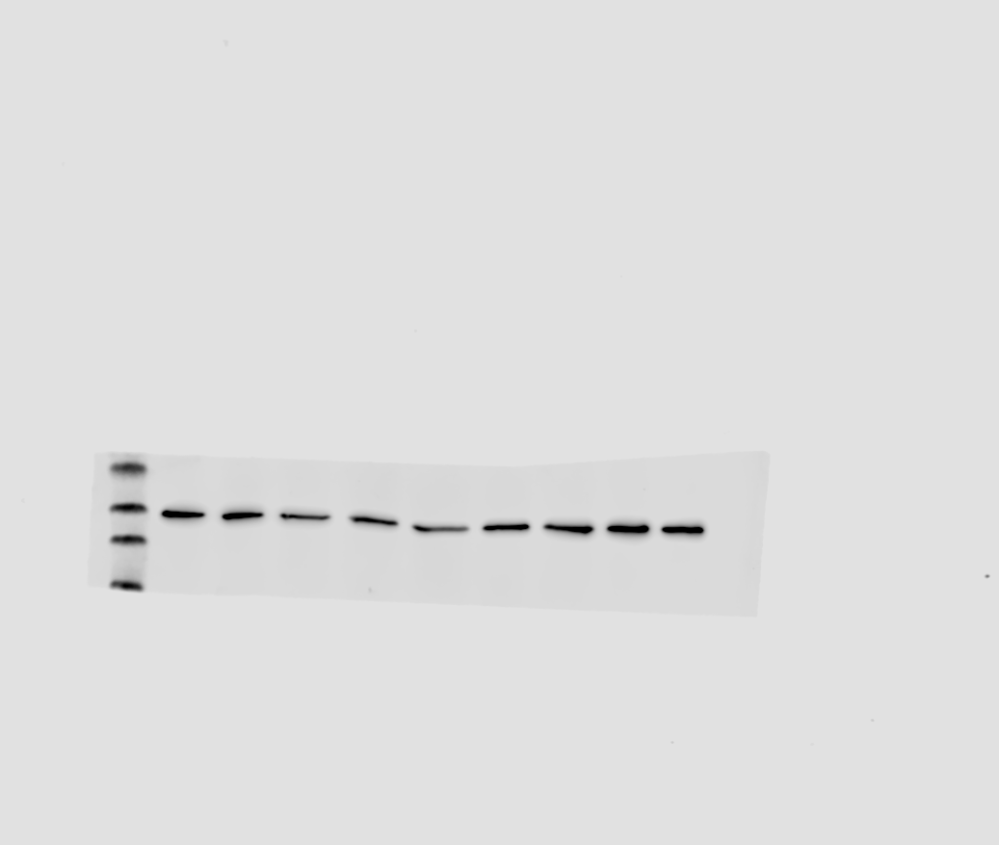

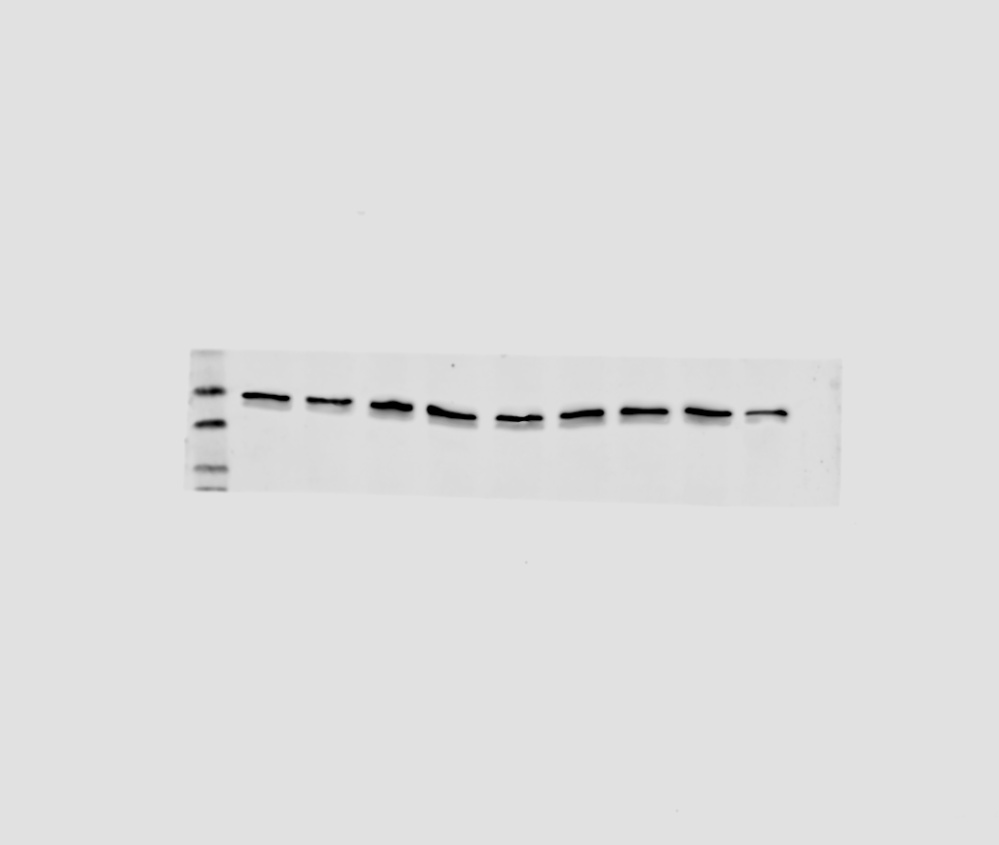

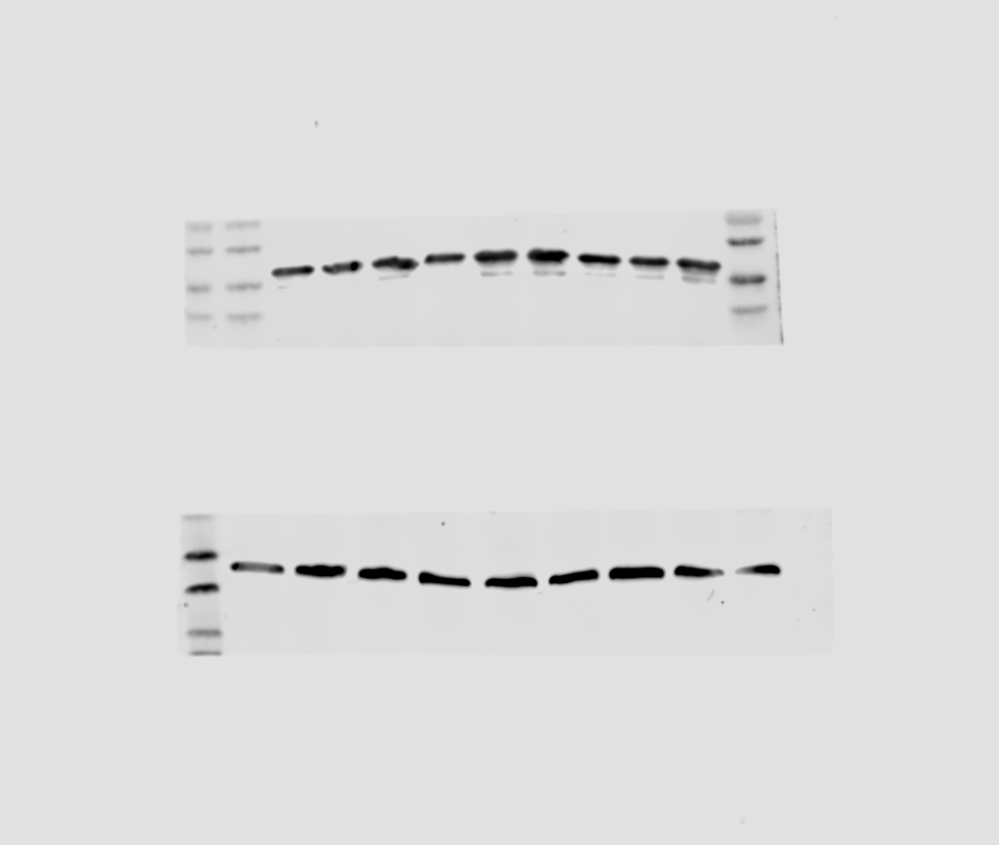

DMSO

**HN-SCC-151**

1

2.5

5

10

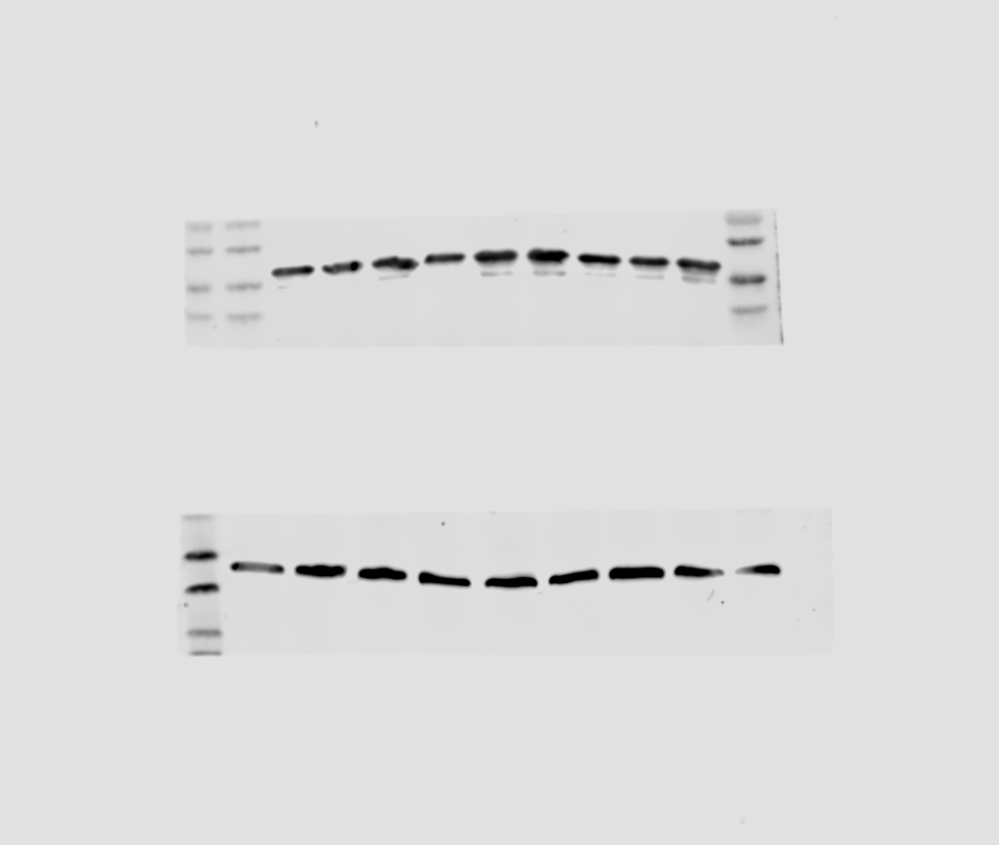

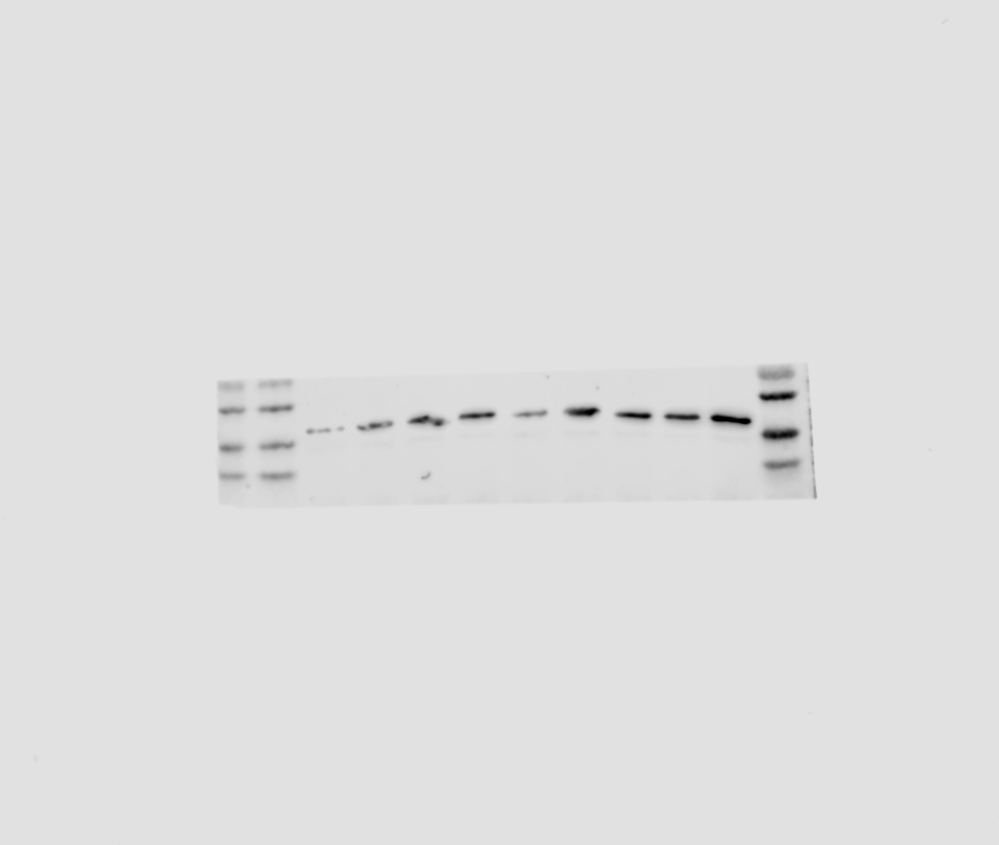

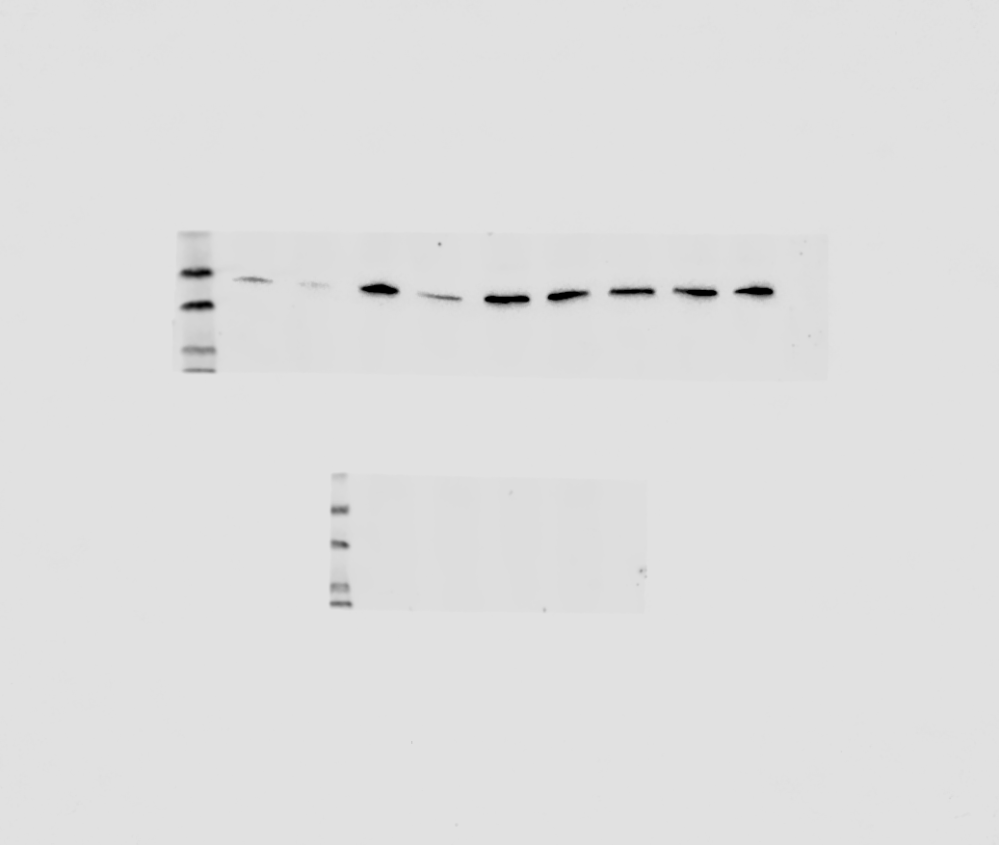

pERK

ERK

Actin

DMSO

**HN-6**

1

2.5

5

10

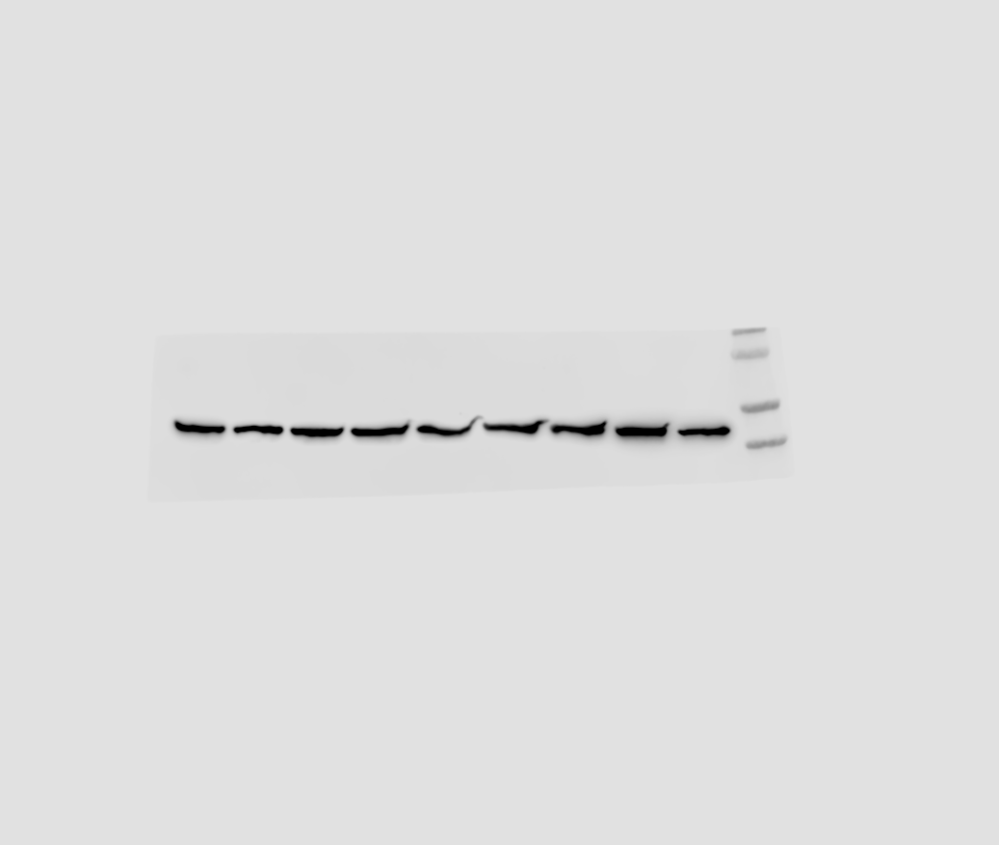

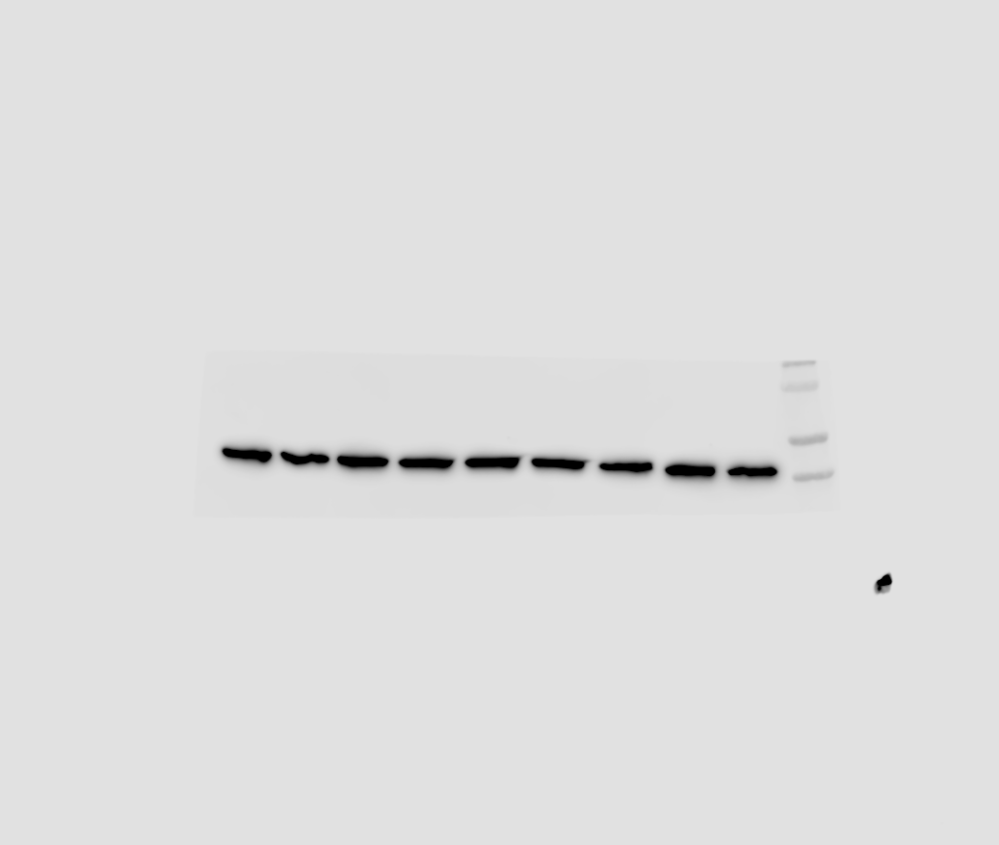

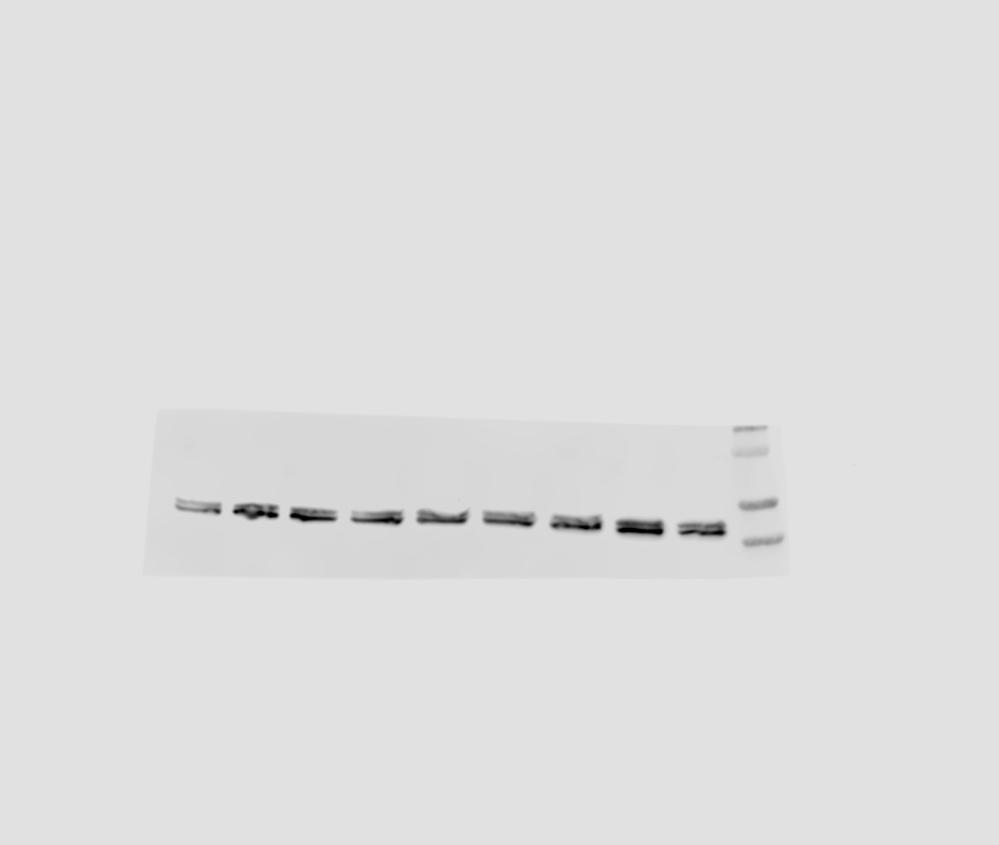

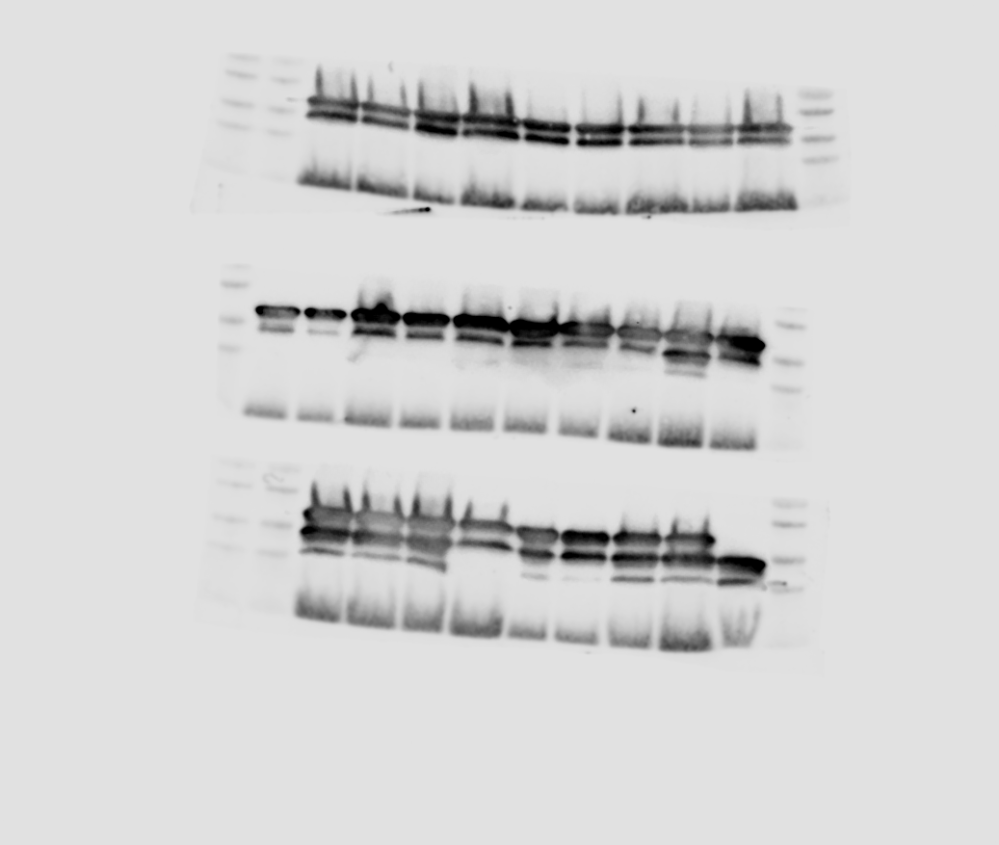

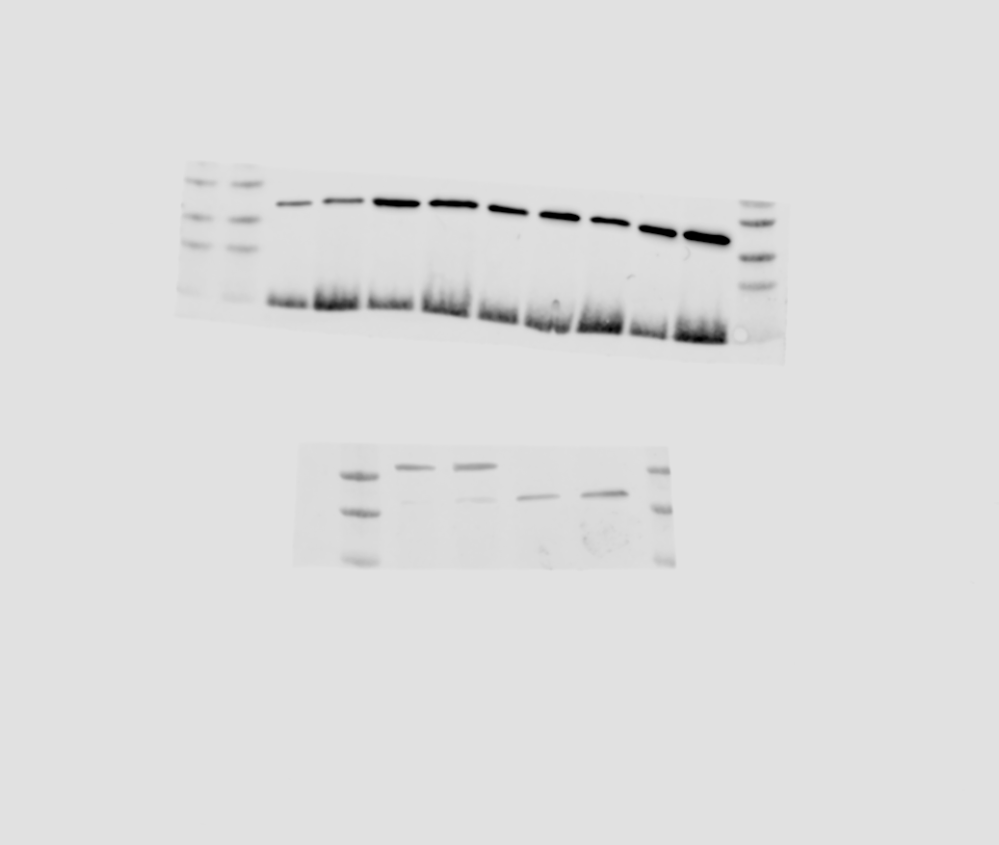

H3K4me3

H3

BAY-6035	BAY-6035	BAY-6035

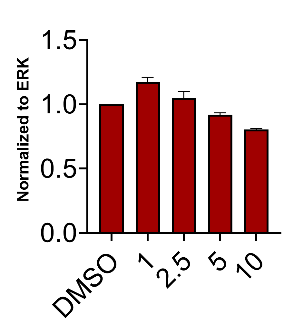

pERK

Relative band intensities

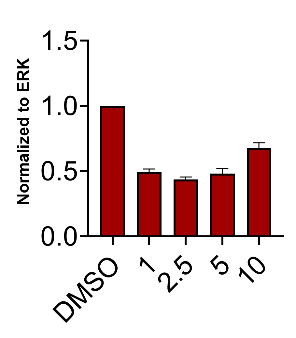

pERK

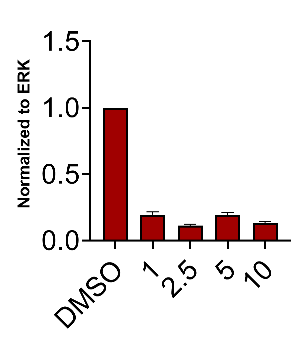

pERK

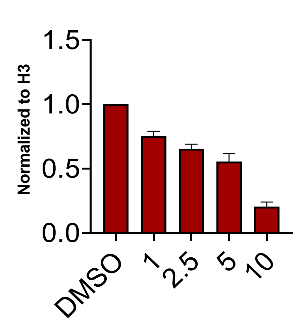

H3K4me3

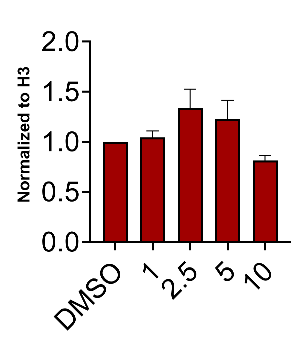

H3K4me3

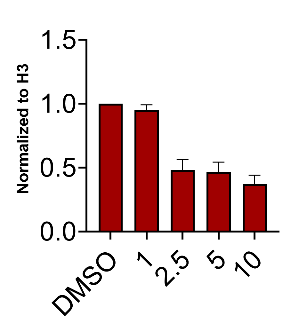

Relative band intensities

H3K4me3

**Supplementary Figure 3.** CCK8 assays **(A)** and CFAs **(B)** of HPV-negative HNSCC cells (HN-6, HN-SCC-151, PE/CA-PJ15 and HN13) treated with BAY-6035 versus DMSO at a concentration range of 1-10uM for 9 days. For the CCK8 assays, cells were seeded at ~500 cells/well in biological quadruples. Medium with the inhibitor was replenished every 48h. CCK8 assays were conducted at the indicated time points (days 1, 3, 6 and 9 of treatment). Curves are showing the average of four biological replicates per condition. Standard deviation (SD) is shown. For the CFAs, cells were seeded at ~100 cells/well in biological triplicates. CFAs were fixed and stained with crystal violet at 10 days of treatment. A representative example of one of the triplicates per cell line and condition is shown.

**(A)**

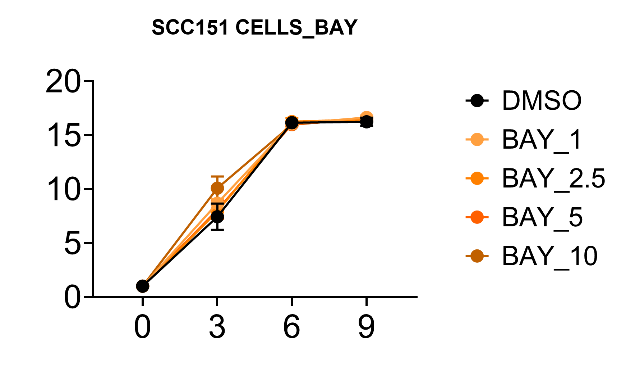

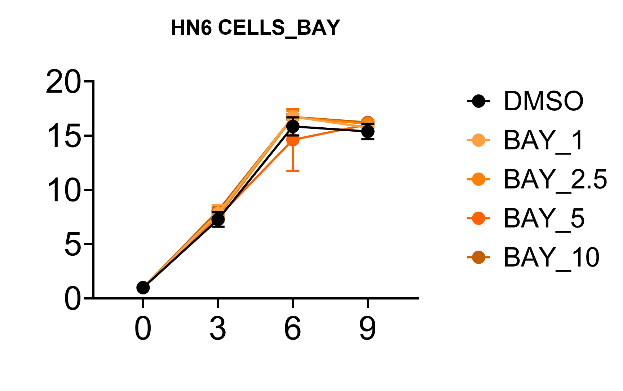
 **HN-6**	**HN-SCC-151**

Relative absorption compared to DMSO

Days

**PE/CA-PJ15	HN13**

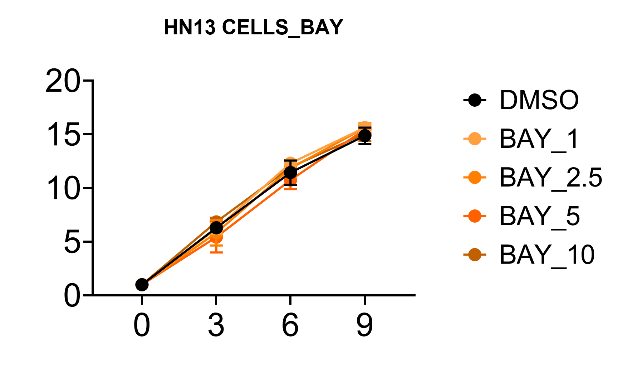

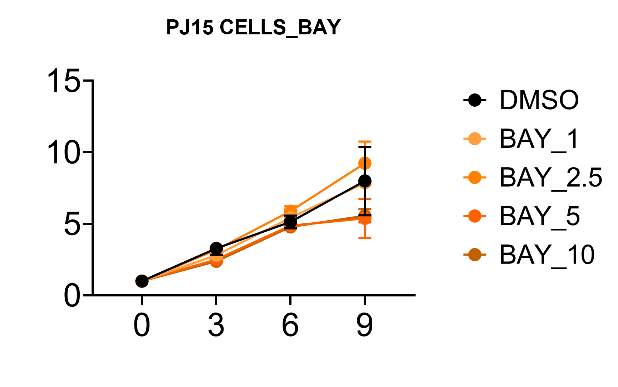

Relative absorption compared to DMSO

Days

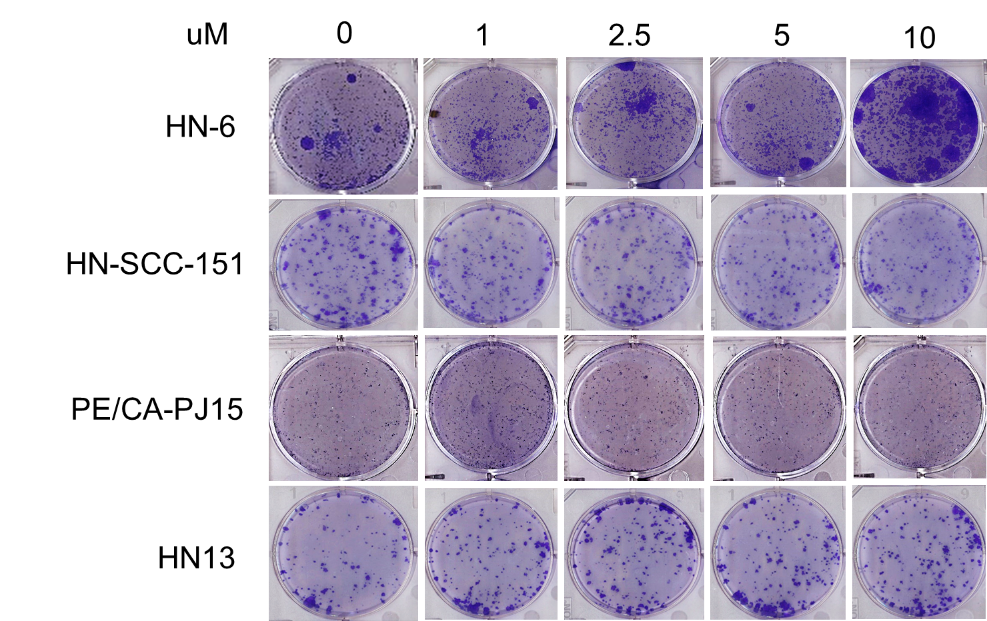
**(B)**

**Supplementary Figure 4.** Global H3K4me3 levels in SMYD3 KO cells before and after stable transfection with a doxycycline-inducible plasmid expressing wild-type versus F183A-mutant SMYD3. 5ug of nuclear extracts were loaded and blotted for H3K4me3 and H3 was used as loading control. H3K4me3 levels were normalized by H3.

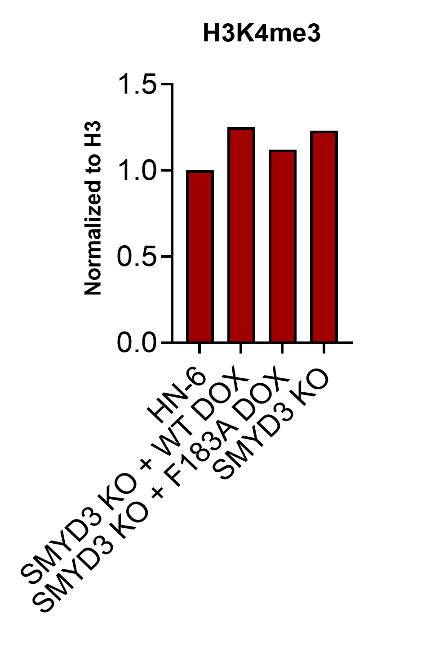

Relative band intensities
