## Supplementary Figure 5 to 7 for "SMYD3 protein degraders in HPV-negative head and neck squamous cell carcinoma"

**Supplementary Figure 5.** Chemical structures of the commercially available SMYD3 inhibitors BCI-121, EPZ028862 and BAY-6035.

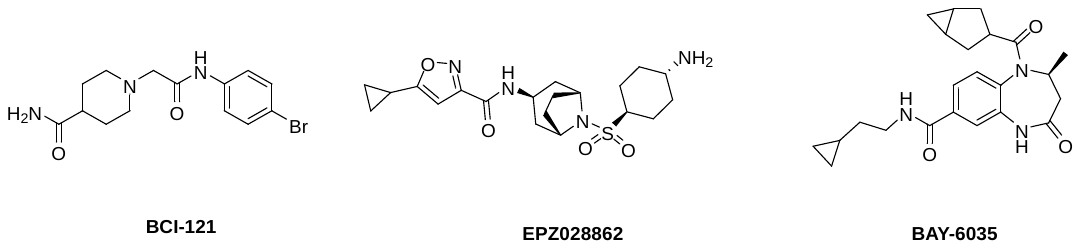

**Supplementary Figure 6.** SMYD3 PROTAC control structures. **(A)** IAP-01(neg) and **(B)** IAP-08(neg) epimer controls, developed with inversion of all stereocenters of the IAP ligand.

**(A)**

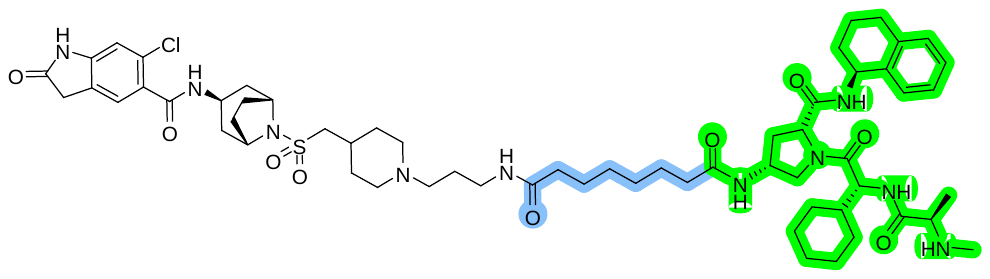

**(B)**

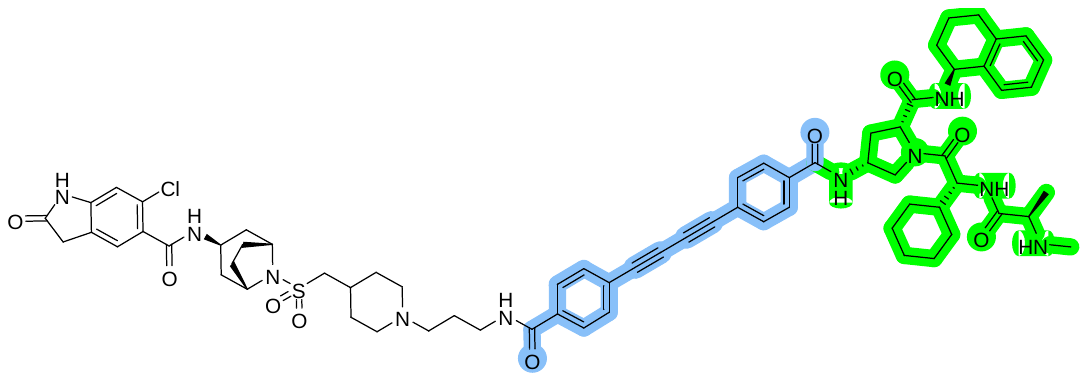

**Supplementary Figure 7.** SMYD3 degradation efficacy of SMYD3 PROTACs evaluated by Western blotting in HN-6 cells. **(A)** HN-6 cells were treated with incremental concentrations of each PROTAC for 48h, cells were collected and nuclear extraction was conducted. 10ug of nuclear extract was loaded and Western blotting for SMYD3 was performed. H3 was used as a control. Results from all PROTACs are shown. Densitometry values are shown at the bottom of each blot. Band intensities were normalized by H3, and comparisons were made with DMSO assigned a relative value of 1. **(B)** Table summarizing the degradation efficacy of SMYD3 PROTACs assessed by Western blotting in HN-6 cells.

**(A)**

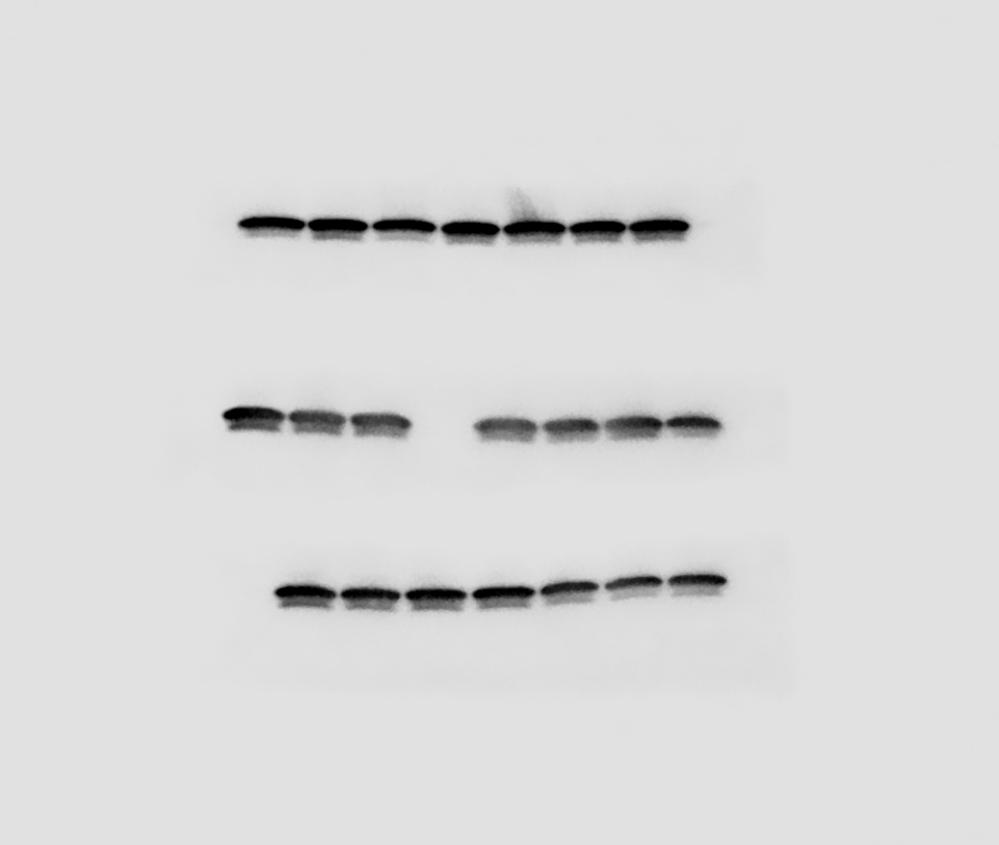

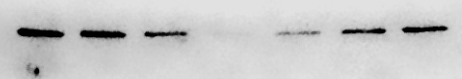

DMSO

0.5

1

2.5

1

0.85

0.05

0.001

0.5

1

2.5

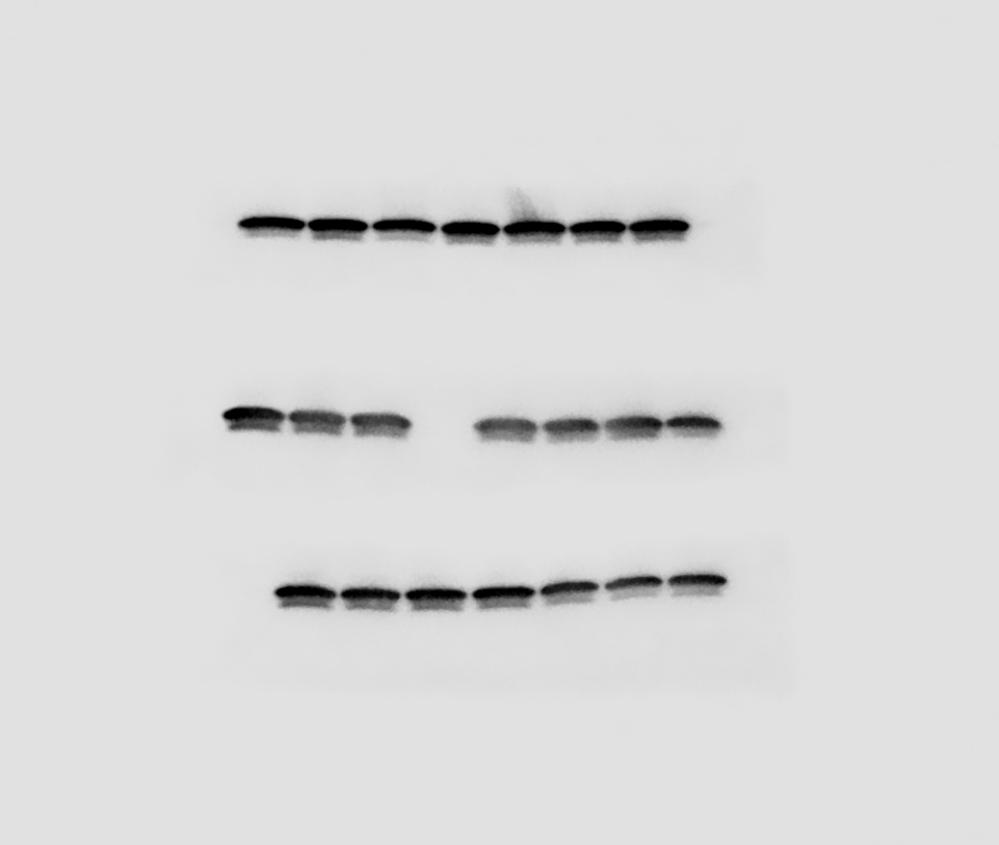

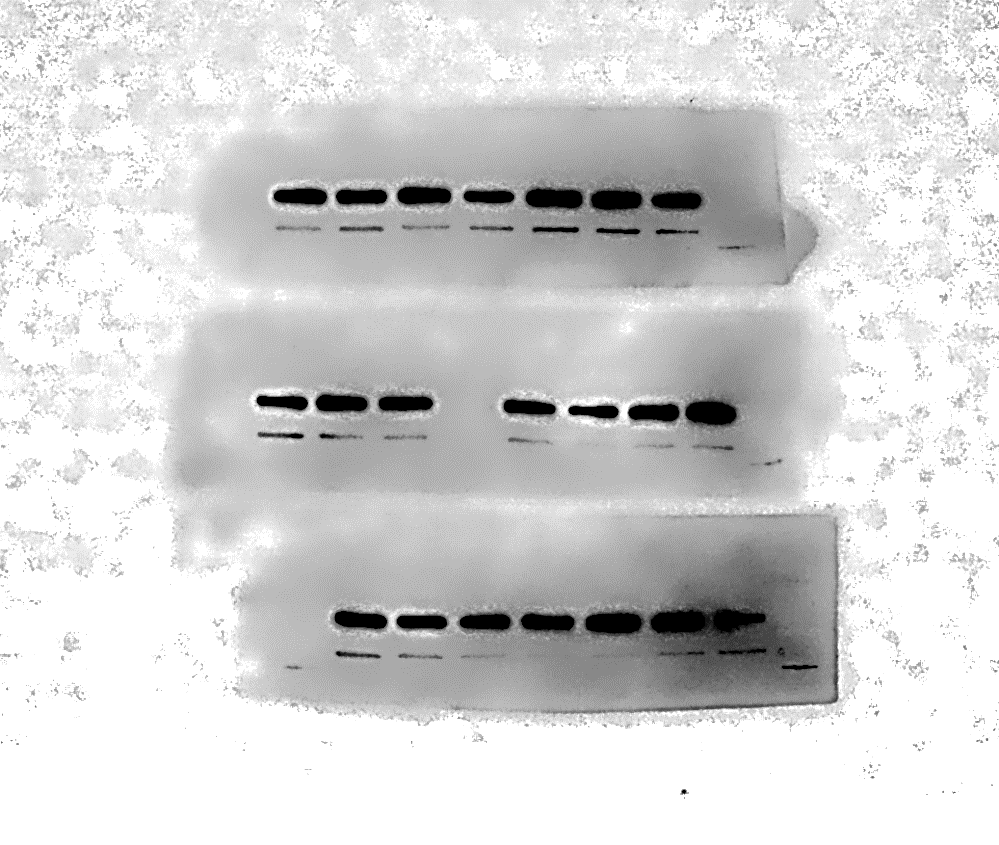

0.25

0.66

0.08

SMYD3

H3

µM

DMSO

1

IAP-01

IAP-02

1

0.95

0.8

0.2

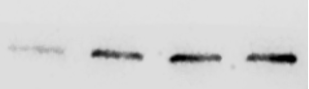

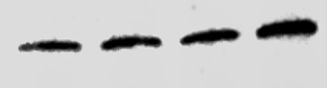

DMSO

1

2.5

0.5

IAP-05

IAP-03

1

2.5

0.5

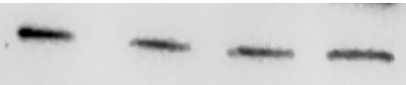

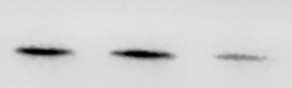

DMSO

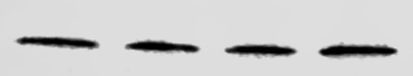

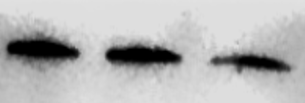

1

2.5

0.5

1

0.8

0.75

0.72

0.25

0.65

0.55

SMYD3

H3

µM

IAP-04

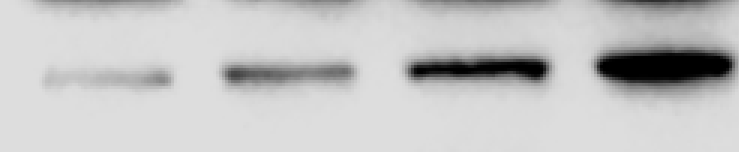

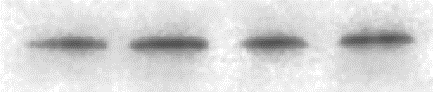

IAP-09

0.1

0.5

1

DMSO

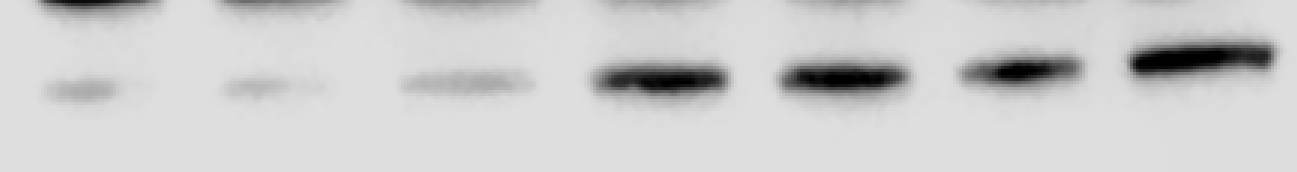

SMYD3

H3

IAP-07

DMSO

0.1

0.5

1

µM

IAP-08

0.1

0.5

1

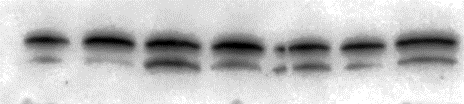

1

0.85

0.75

0.72

0.15

0.05

0.12

1

0.85

0.25

0.12

1.1

1.32

1.48

0.5

1

2.5

1

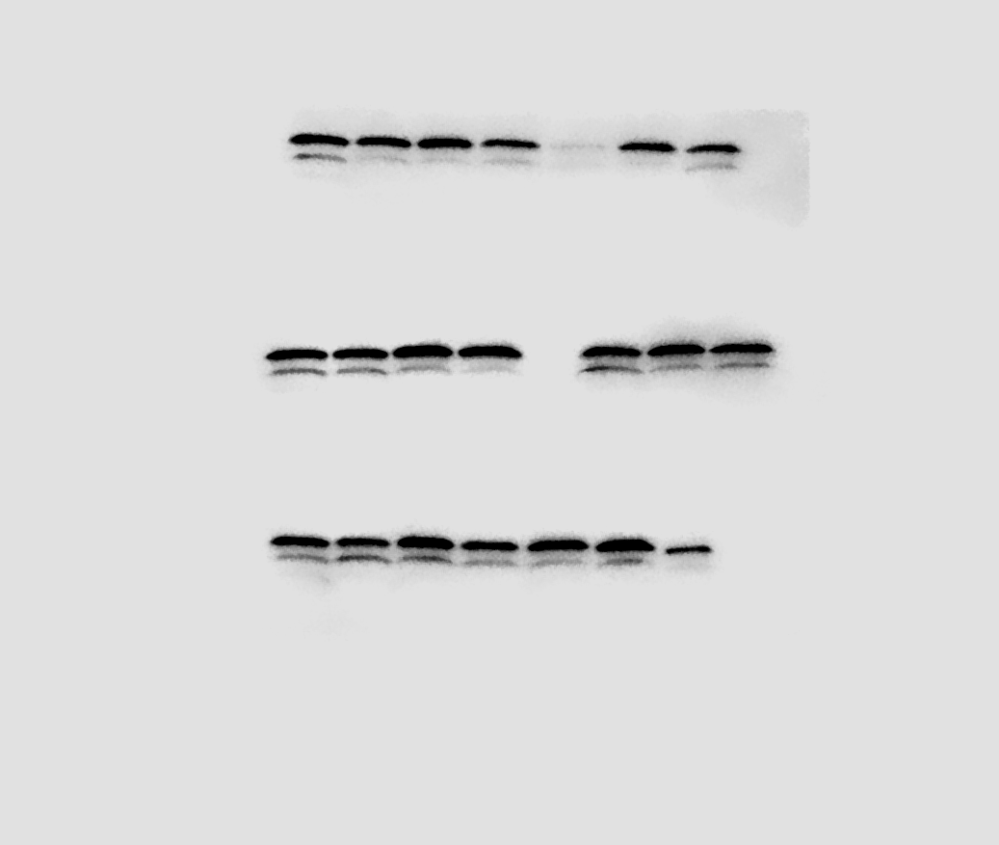

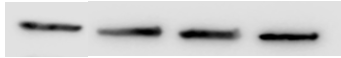

DMSO

CRBN-02

DMSO

0.5

1

2.5

1

CRBN-03

SMYD3

H3

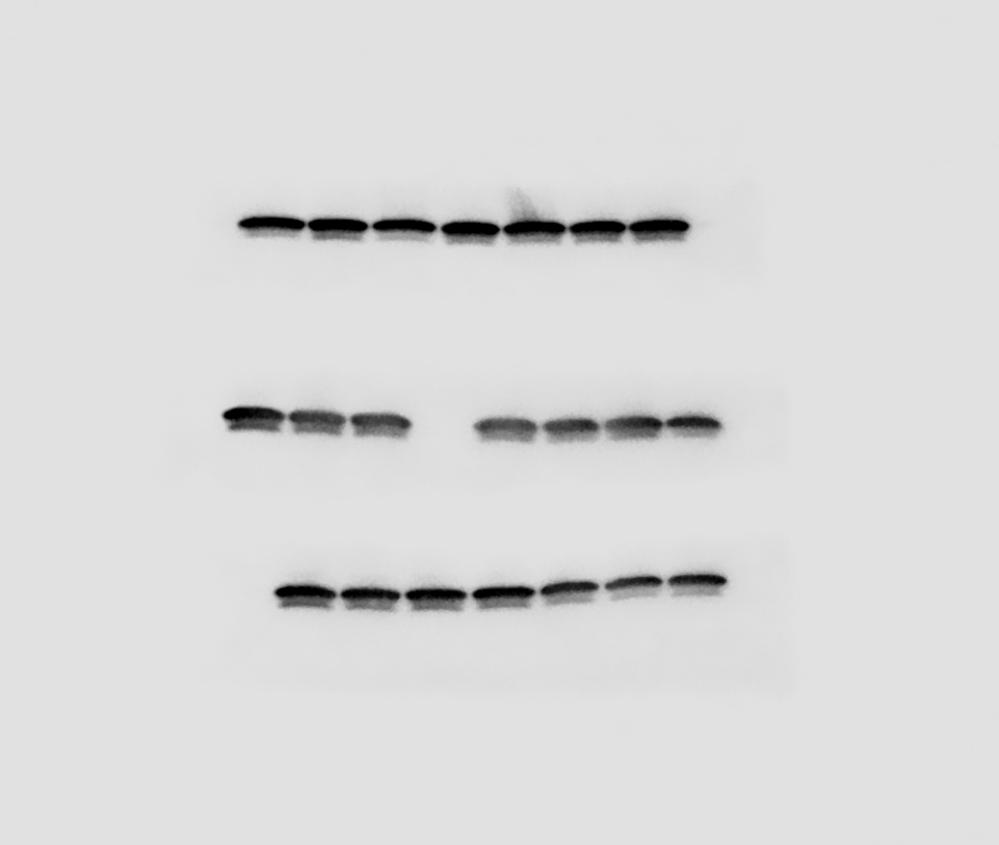

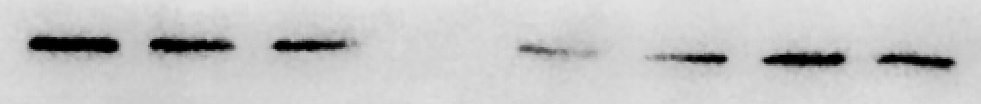

DMSO

0.5

1

2.5

µM

1

0.98

0.67

0.48

CRBN-01

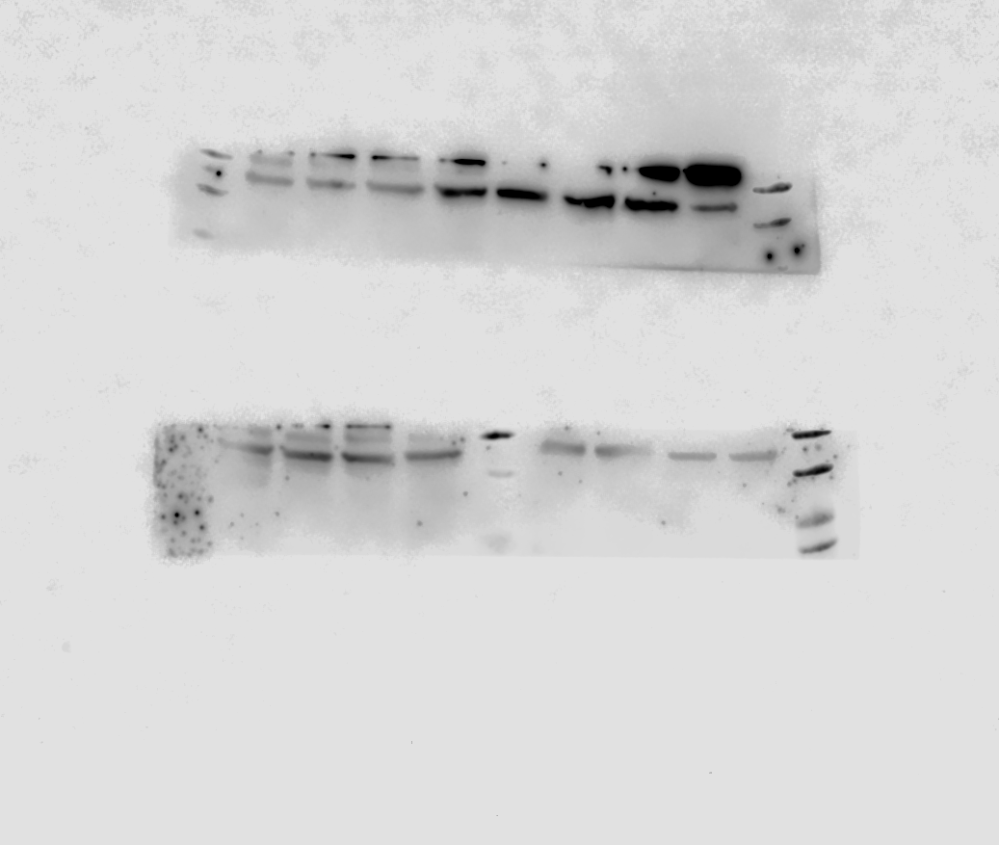

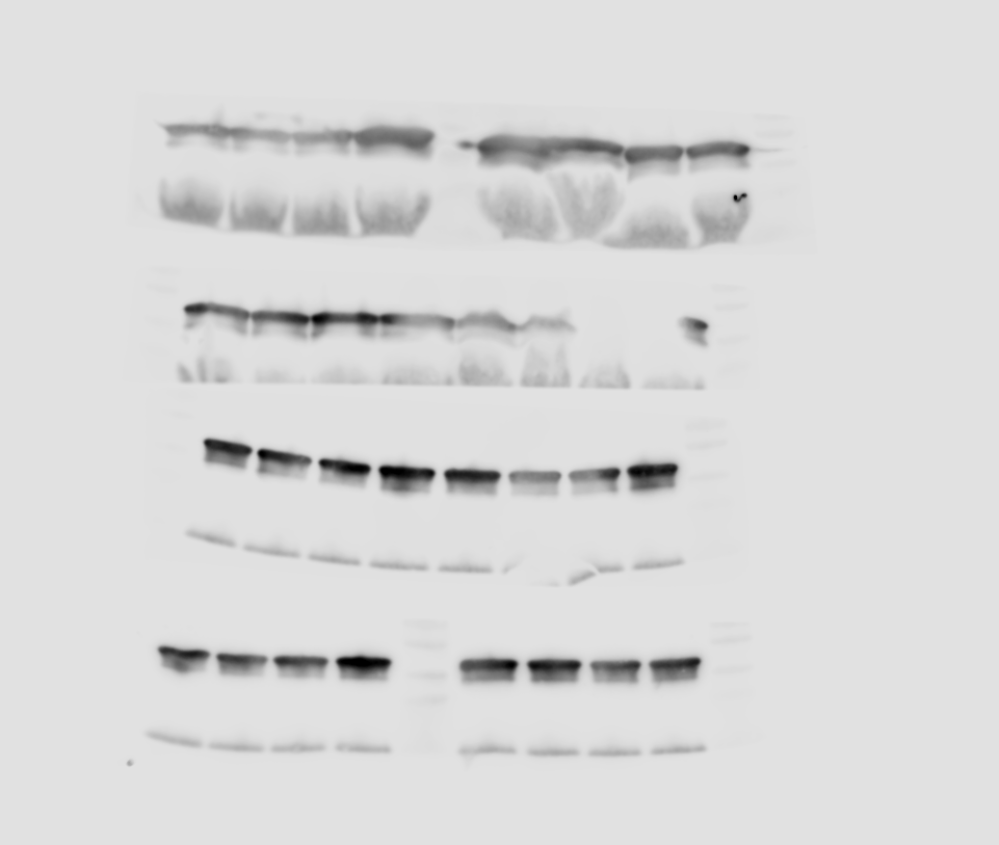

0.29

0.12

0.49

DMSO

1

1.05

1.05

0.5

1

2.5

VHL-03

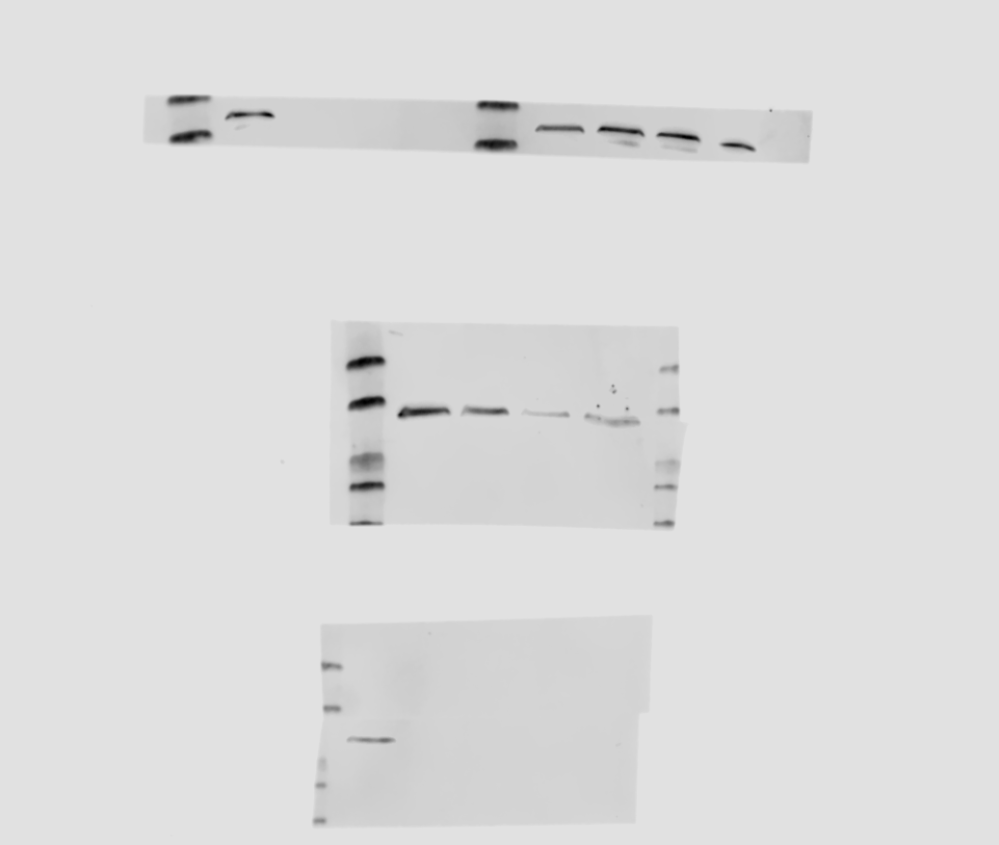

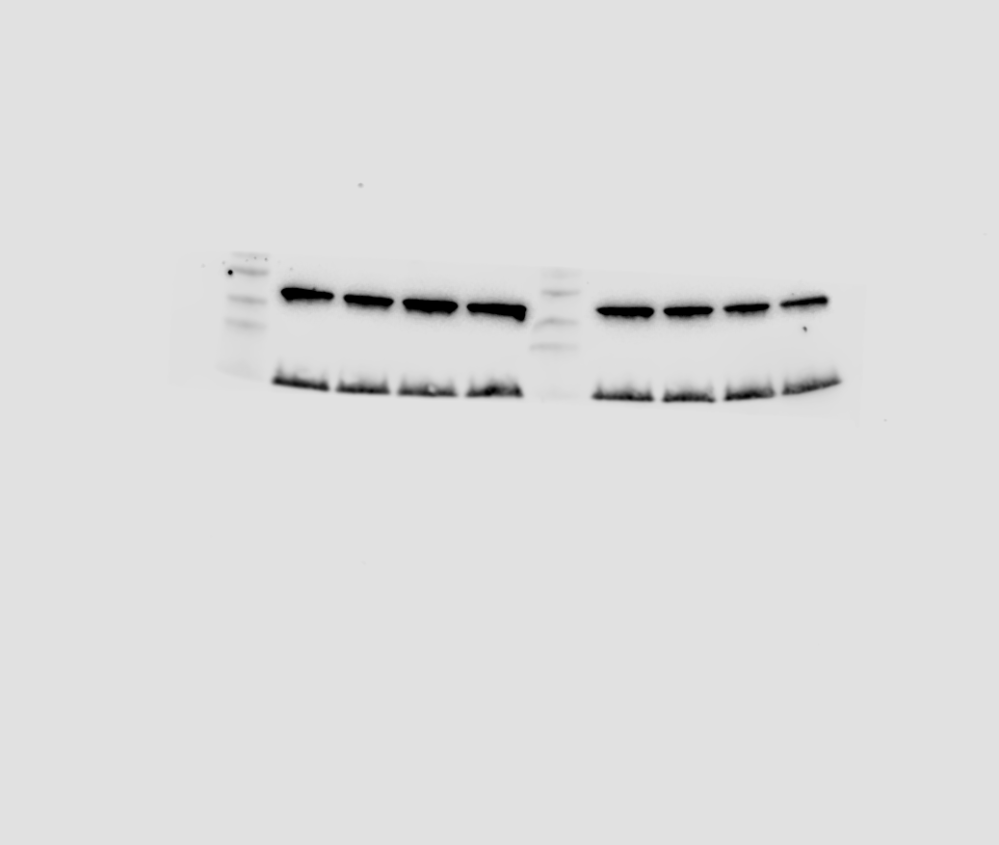

SMYD3

H3

DMSO

0.5

1

2.5

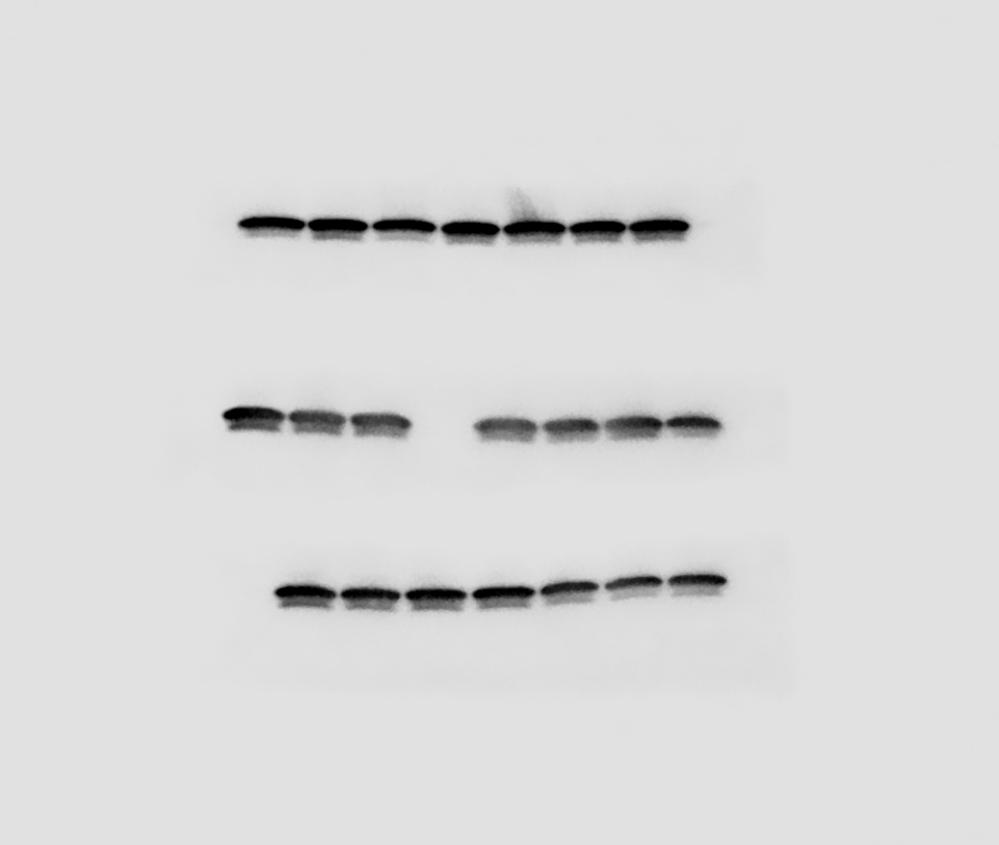

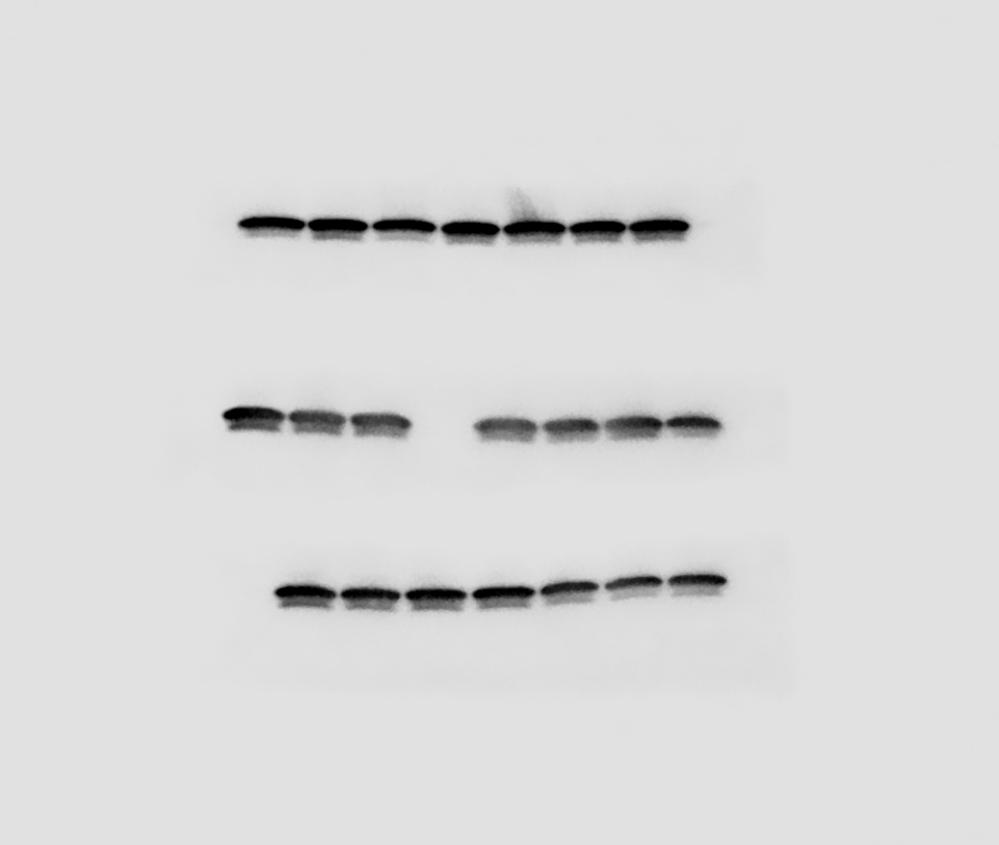

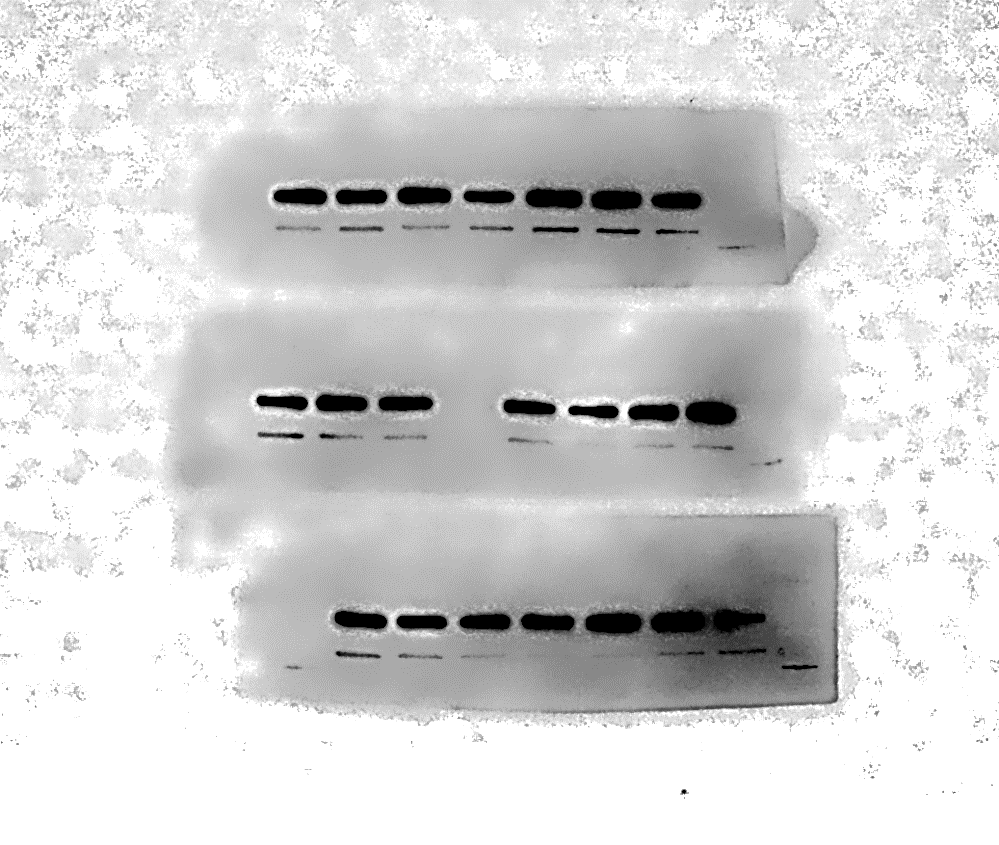

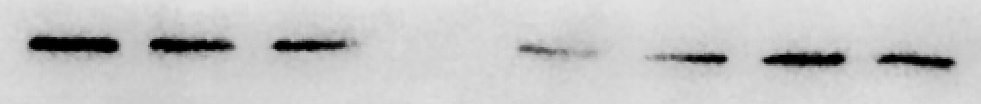

0.5

1

2.5

1

1.02

1

0.4

0.91

1.16

1.08

µM

VHL-01

VHL-02

0.63

DMSO

0.5

1

1

VHL-04

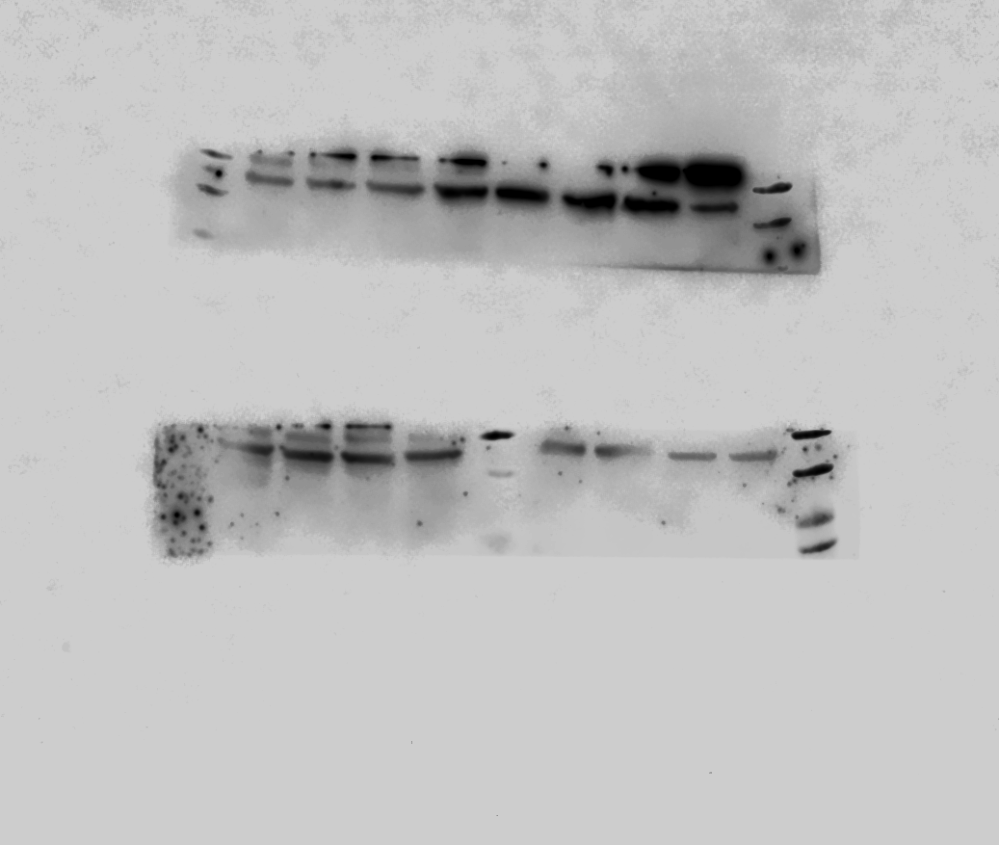

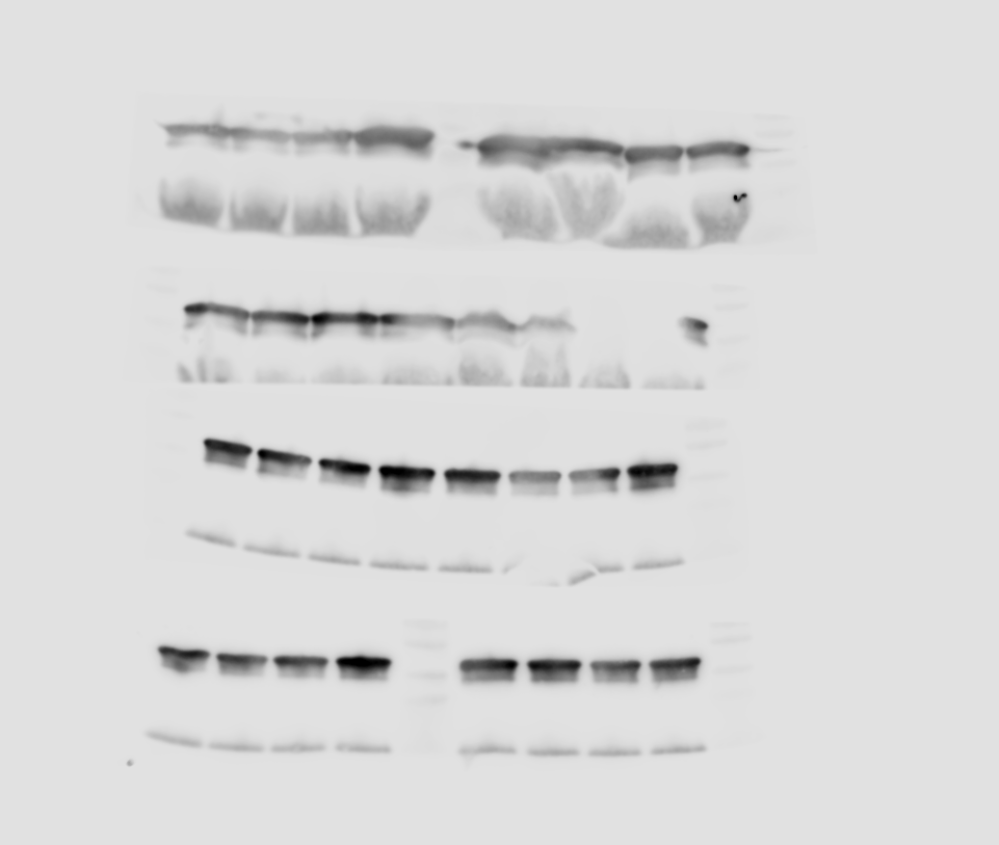

2.5

1.2

1.2

1.16

**(B)**

| **Compound Code** | **SMYD3 Degradation at 0.5uM for 48h (%), HN-6** | **Maximal SMYD3 Degradation at 48h (%, C at uM), HN-6** |
| --- | --- | --- |
| IAP-01 | 75% | 92%, 2.5uM |
| IAP-02 | 15% | 99%, 2.5uM |
| IAP-03 | 75% | 75%, 0.5uM |
| IAP-04 | 28% | 28%, 0.5uM |
| IAP-05 | 80% | 80%, 0.5uM |
| IAP-07 | 25% | 28%, 1uM |
| IAP-08 | 95% | 95%, 0.5uM |
| IAP-09 | 75% | 88%, 1uM |
| CRBN-01 | 2% | 52%, 2.5uM |
| CRBN-02 | No effect | No effect |
| CRBN-03 | 51% | 88%, 2.5uM |
| VHL-01 | No effect | 60%, 2.5uM |
| VHL-02 | 9% | No effect |
| VHL-03 | No effect | 37%, 2.5uM |
| VHL-04 | No effect | No effect |
