## Supplementary Figure 8 to 10 for "SMYD3 protein degraders in HPV-negative head and neck squamous cell carcinoma"

**Supplementary Figure 8.** Western blotting and densitometries for SMYD3, H3K4me3, UHRF1 and pERK protein levels in HPV-negative HNSCC cells treated with IAP-08. **(A)** HN13 cells were treated with IAP-08 versus DMSO at a concentration range of 0.1-1uM for 48h. Nuclear and cytoplasmic extracts were obtained. 10ug of nuclear or 20ug of cytoplasmic extracts were loaded. H3 or actin were used as a loading control. **(B)** Densitometry values of blots shown in Figure 3A and for HN13 cells. Band intensities were normalized by H3 or ERK, and comparisons were made with DMSO assigned a relative value of 1.

**(A)**

H3K4me3

HN13

SMYD3

H3

IAP-08 (48h)

DMSO

0.1

0.5

1

µM

UHRF1

pERK

ERK

Actin

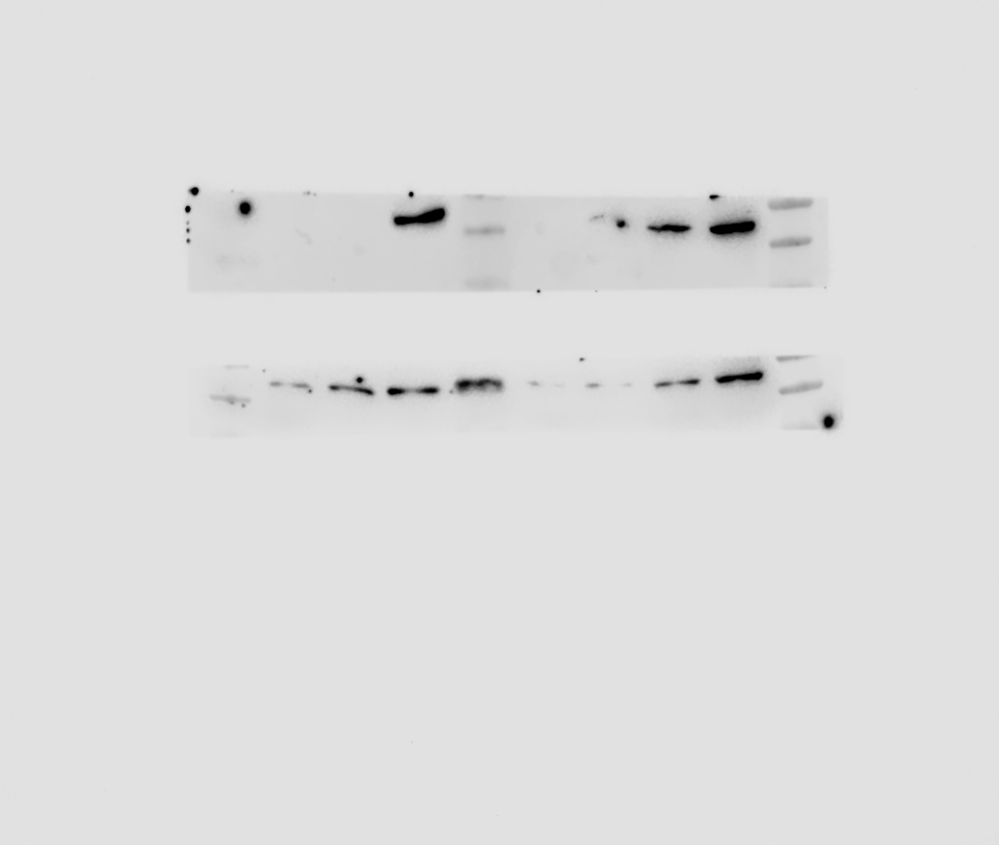

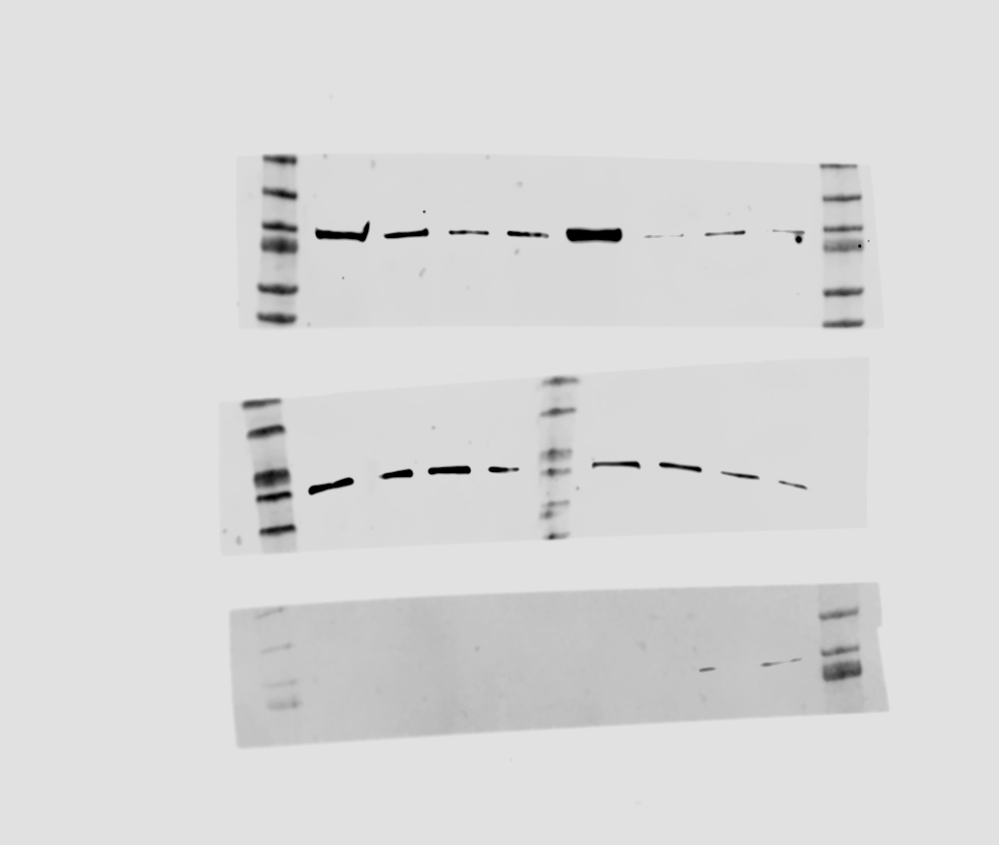

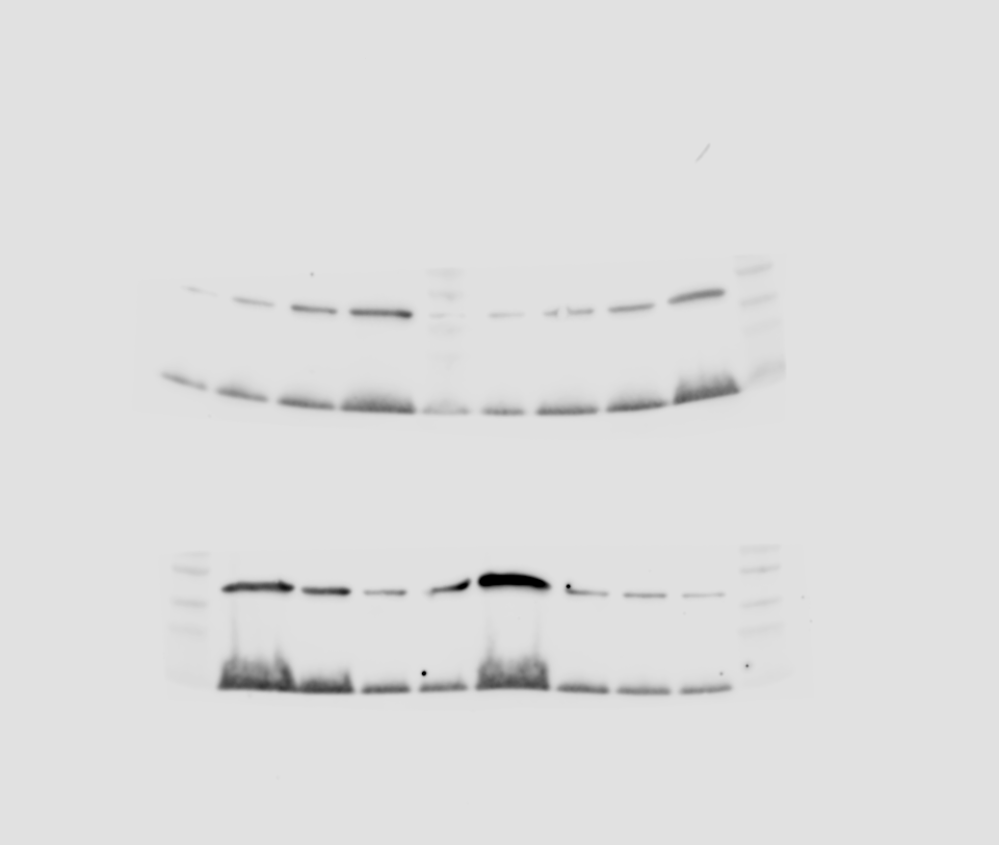

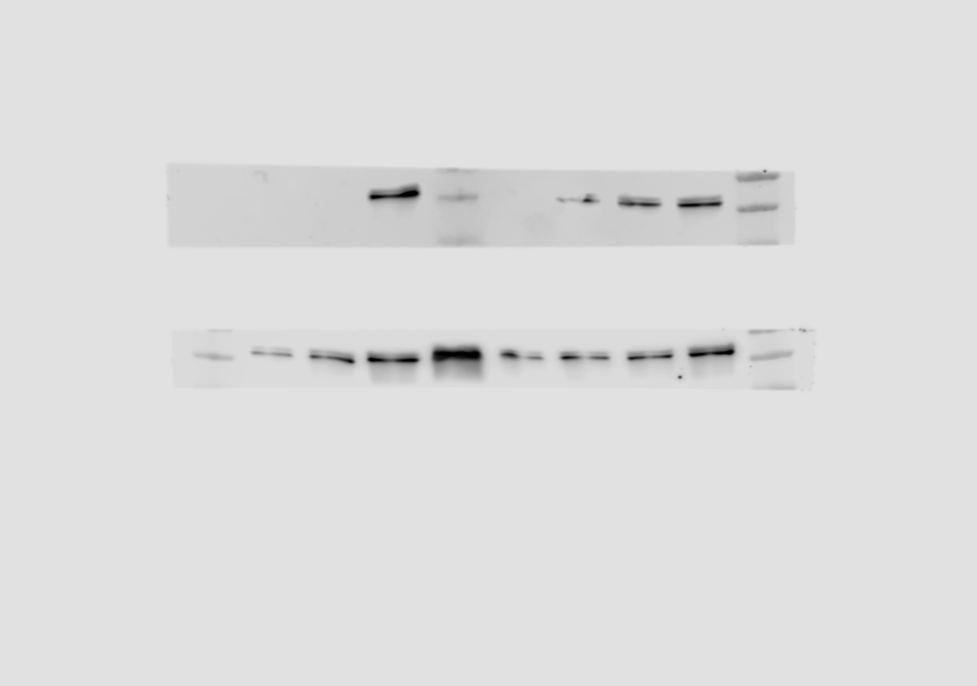

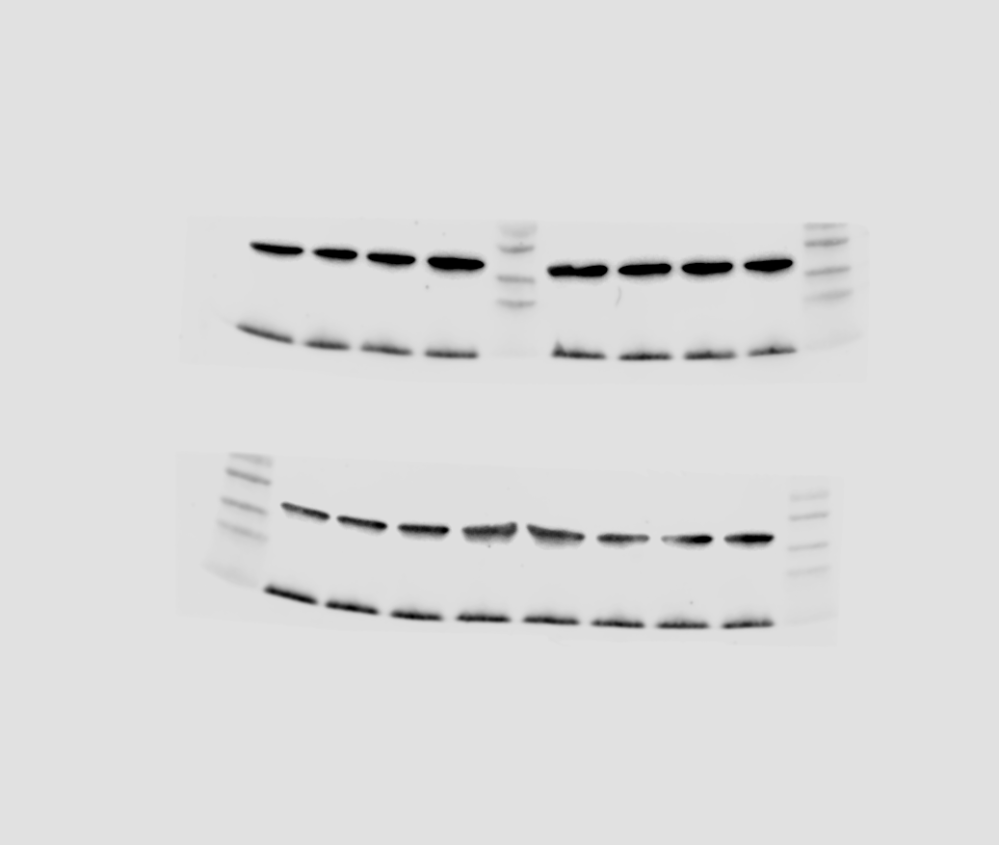

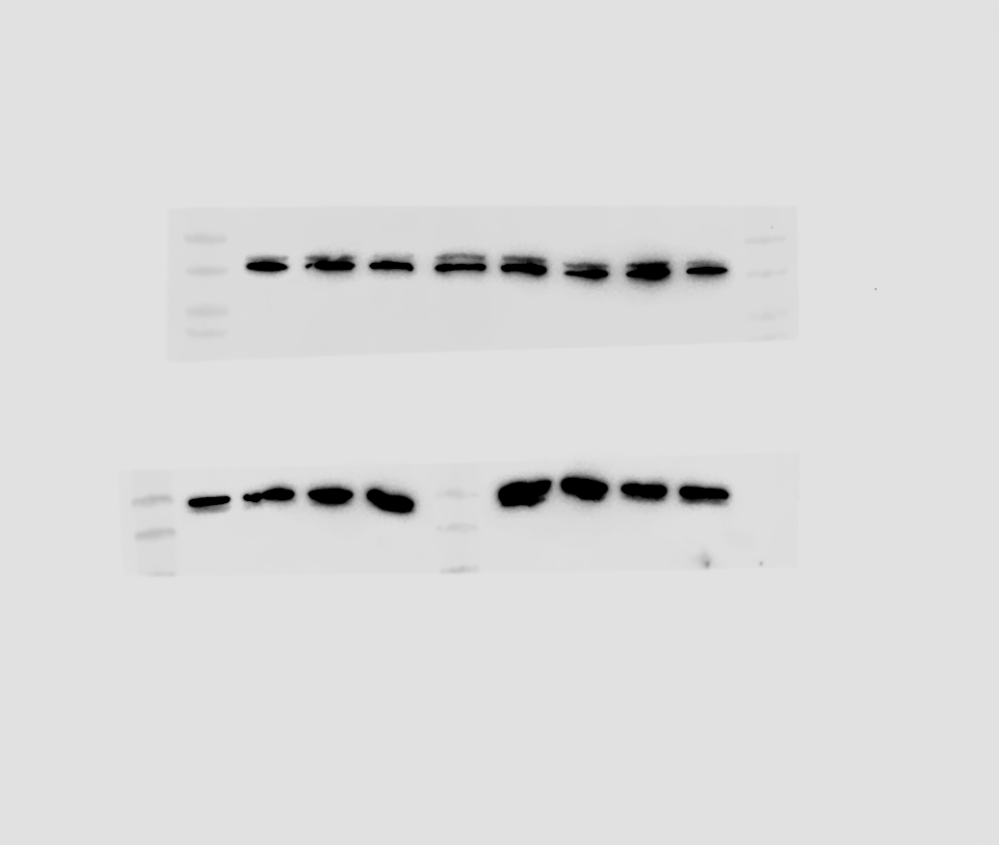

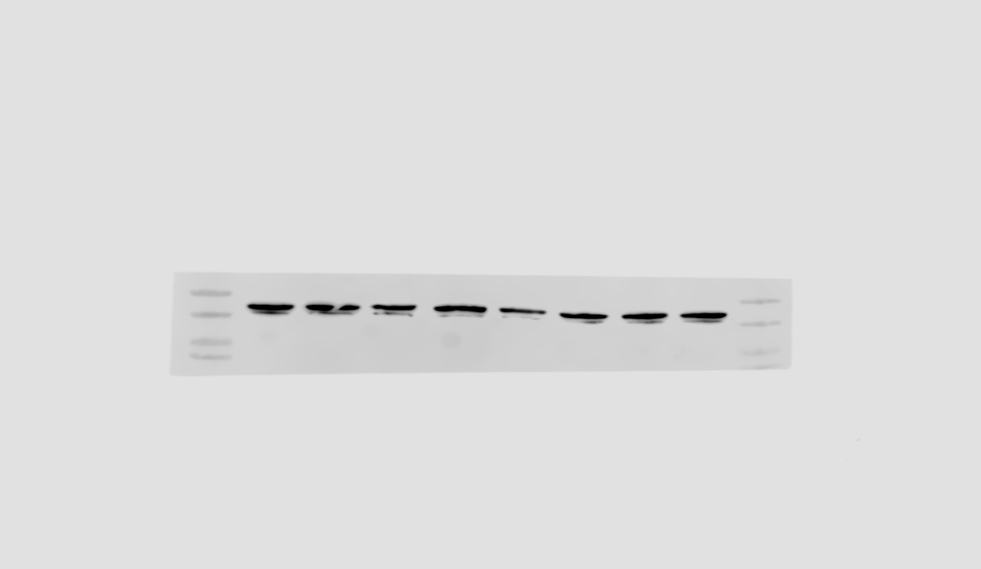

**(B)**

HN-6

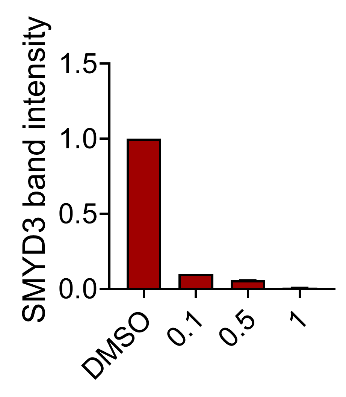

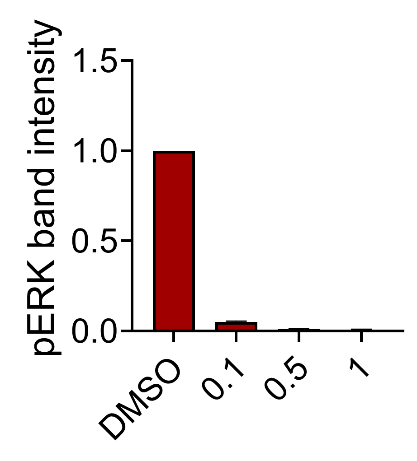

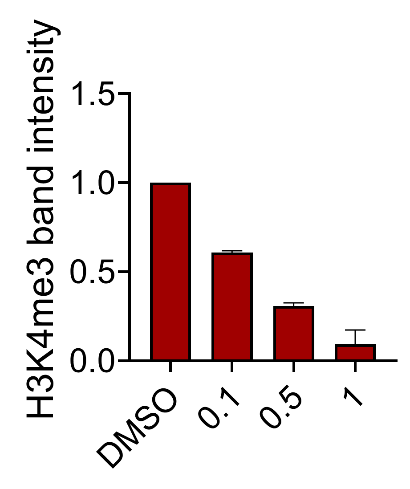

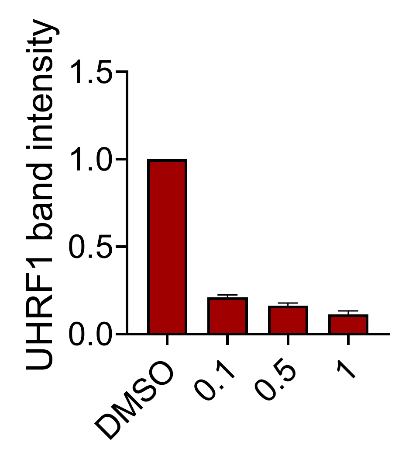

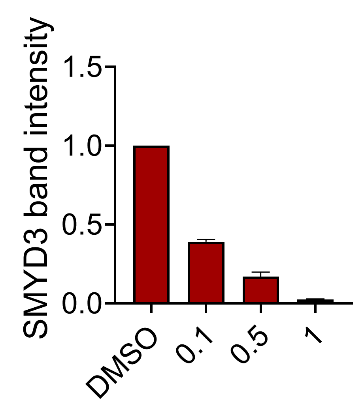

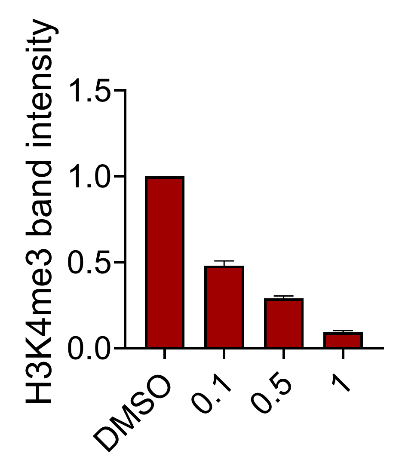

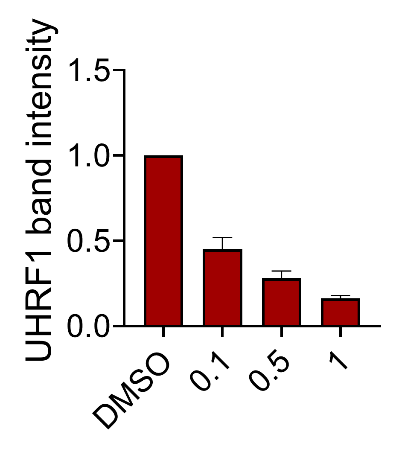

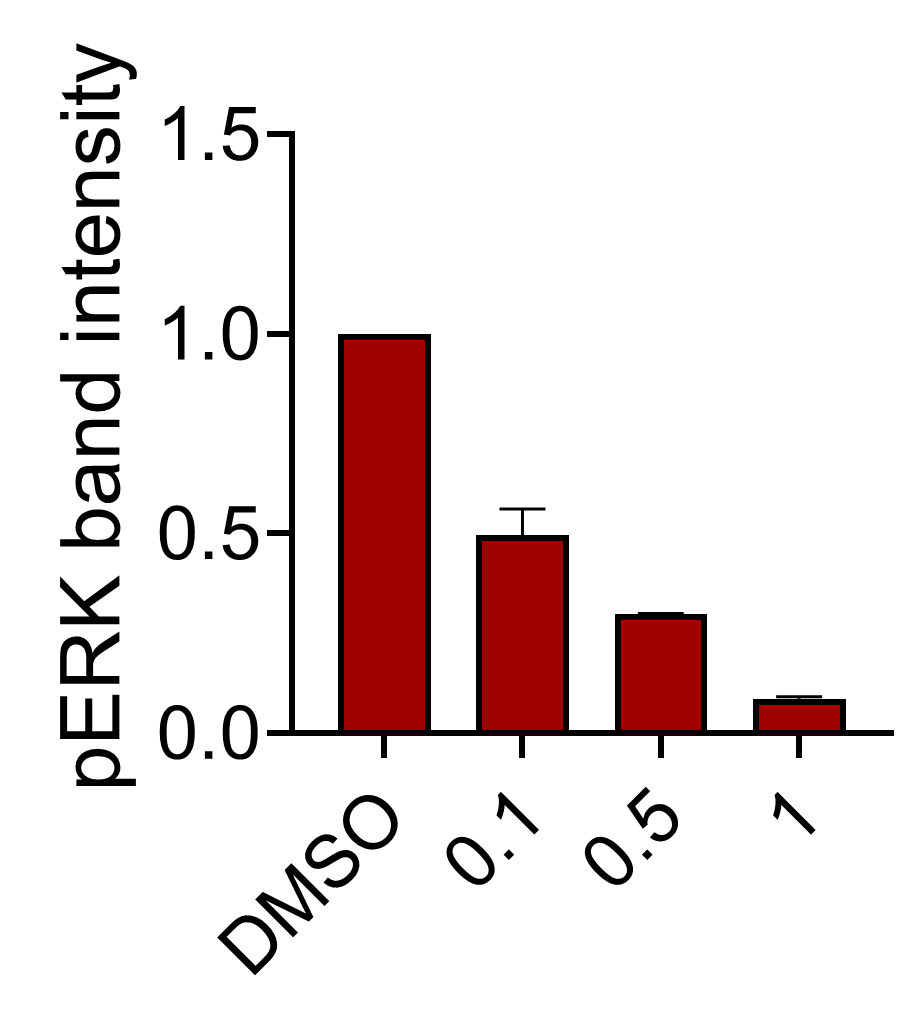

HN-SCC-151

HN-6

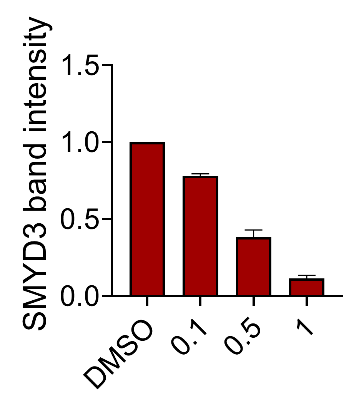

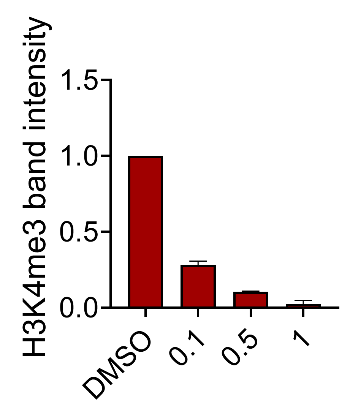

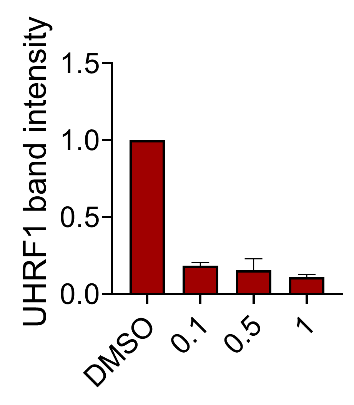

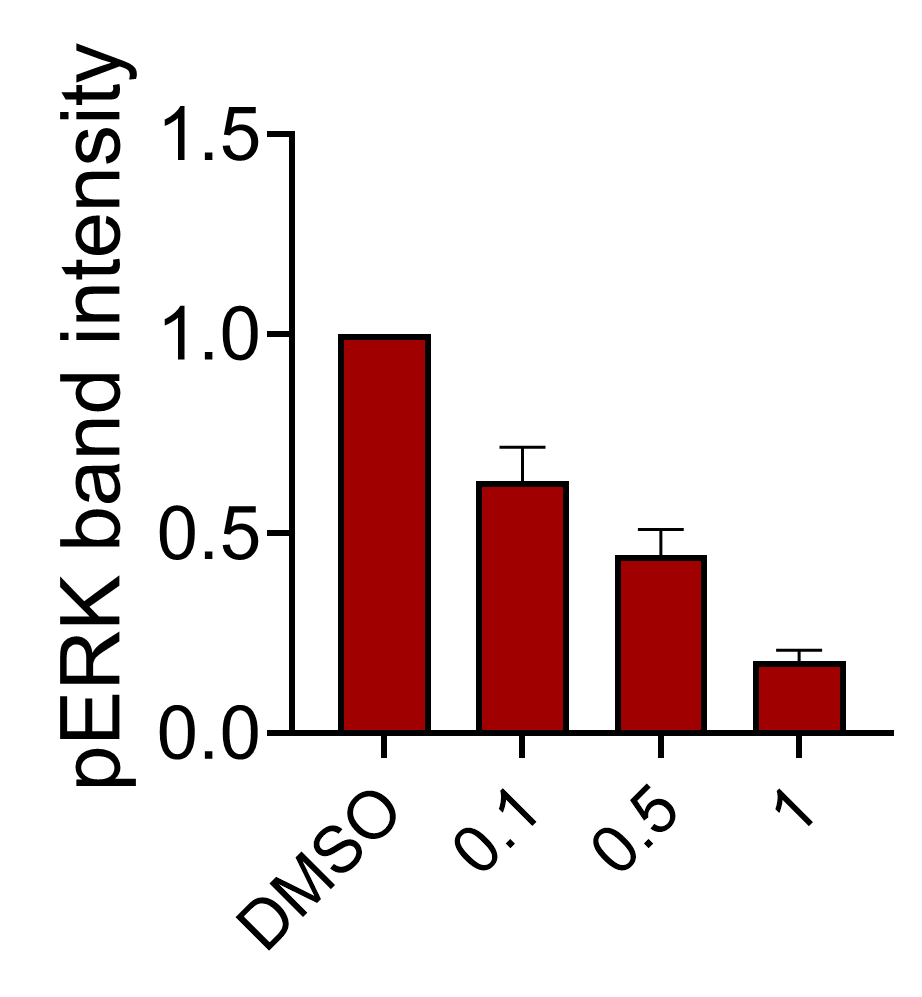

PE/CA-PJ15

HN13

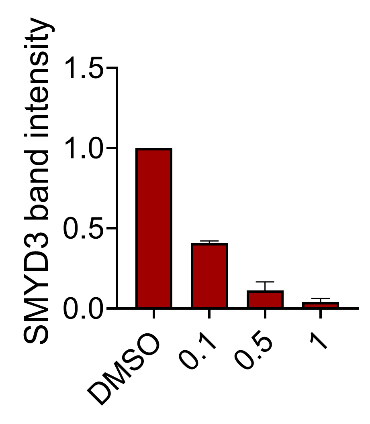

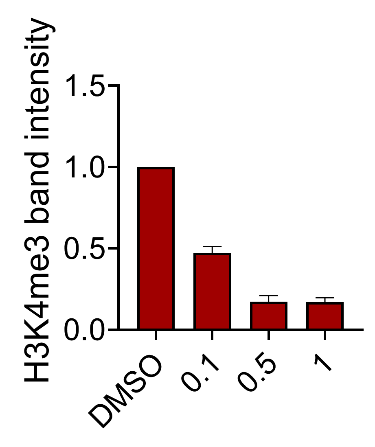

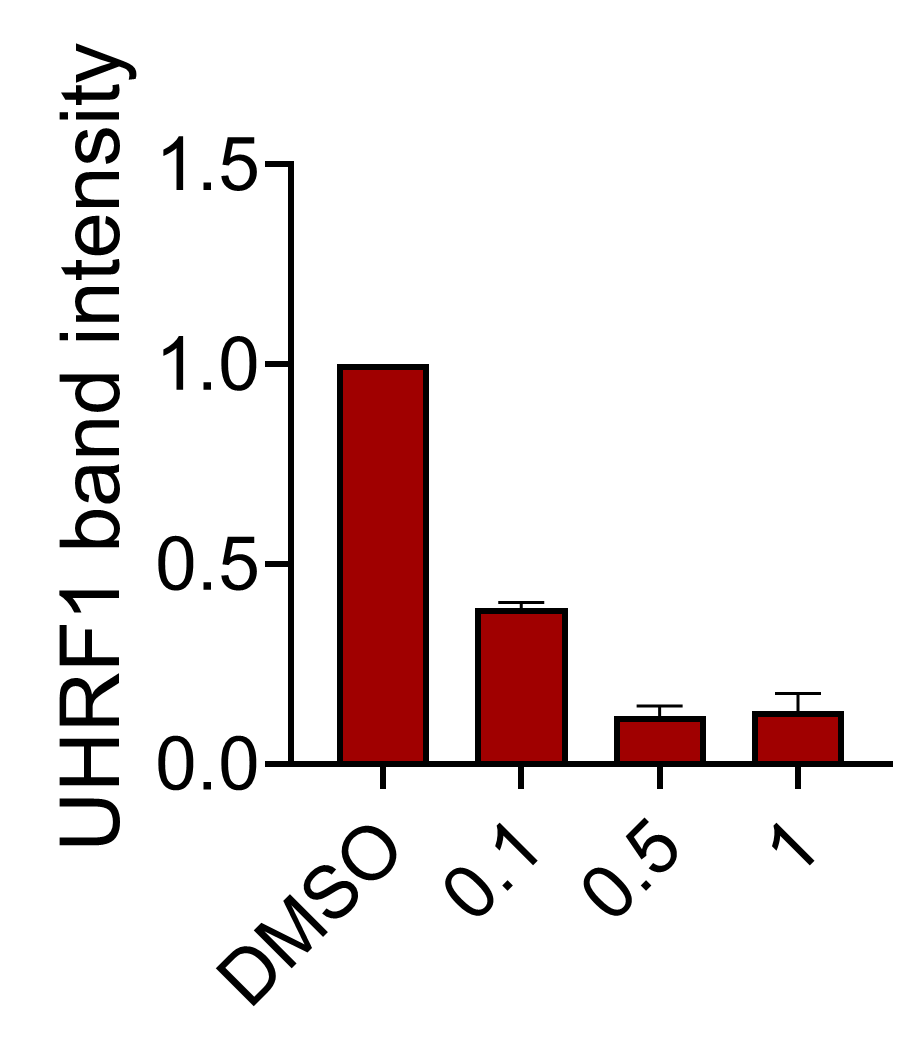

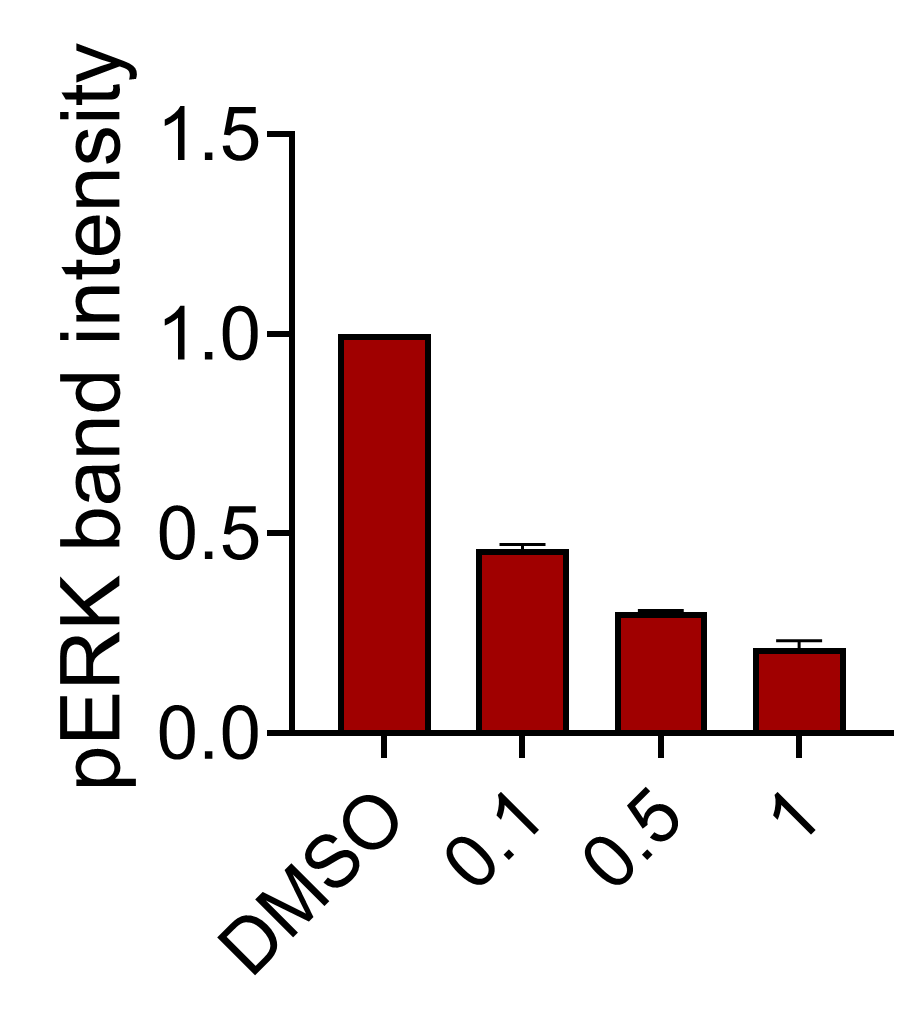

**Supplementary Figure 9.** Confirmation of expression of HiBiT-tagged SMYD3 with HiBiT-assays in HN13 cells stably transfected with a HiBiT-tagged SMYD3 expressing plasmid. Stably transfected HN13 cells were seeded in duplicates in 96-well plates (10,000 cells/well) and induced with doxycycline (1ug/ml) for 24h. The average relative bioluminescence from two biological replicates is shown. The HiBit-tagged SMYD3 transfected clones are shown in the x axis as #1-#6.

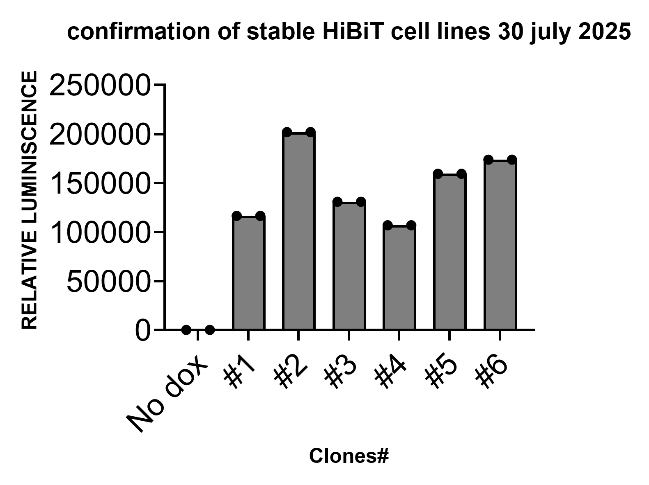

Relative Bioluminescence

Clones

**Supplementary Figure 10.** HiBiT assays in doxycycline-inducible HN13 cells treated with CRBN-, VHL- and IAP-based SMYD3 PROTACs. HN13 cells stably transfected with a doxycycline-inducible plasmid expressing HiBiT-tagged SMYD3 were exposed to doxycycline for 48h and then treated with the SMYD3 PROTACs CRBN-01 to -03 **(A)**, VHL-01 to -04 **(B)**, and IAP-01 to -05, -07 and -09 **(C)** at range of concentrations up to 10uM for 24h. The maximal level of SMYD3 degradation induced by each compound is shown as % with a blue dotted line (Dmax, %). Two biological replicates per condition are shown. The y axis represents the % bioluminescence compared to DMSO (control) and the x axis represents the concentration of the compound as log[C]M. **(D)** Summary table of the DC50 and Dmax for all compounds.

**(A)**

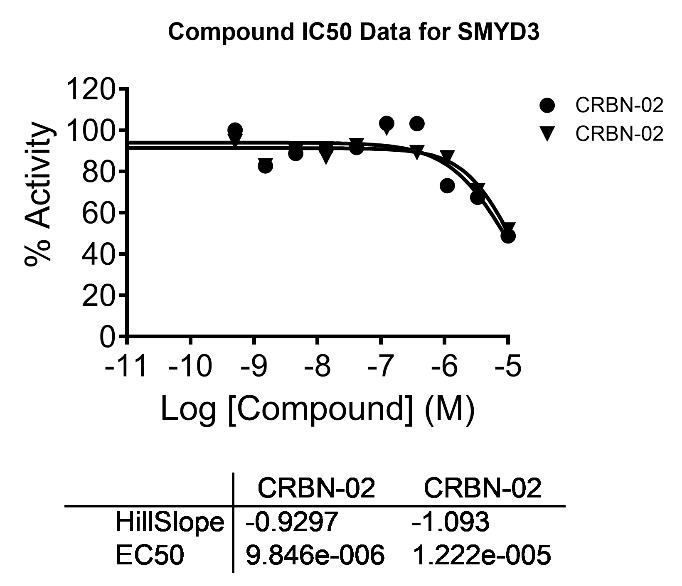

% Bioluminescence

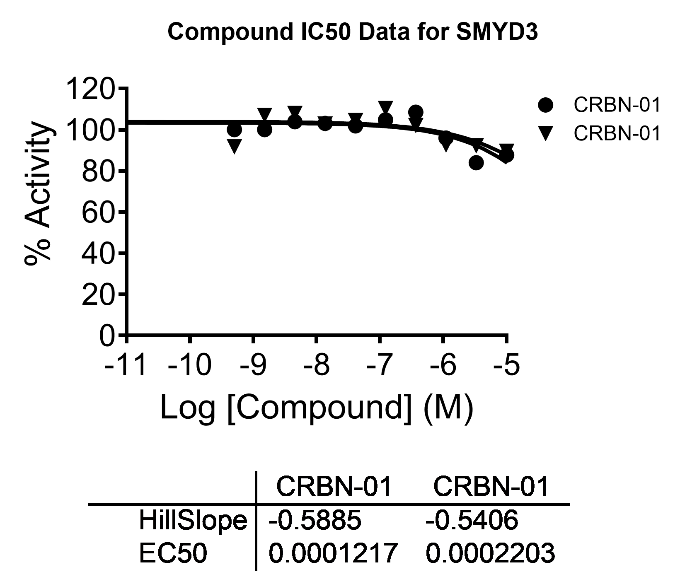

% Bioluminescence

Dmax~20%

Dmax~40%

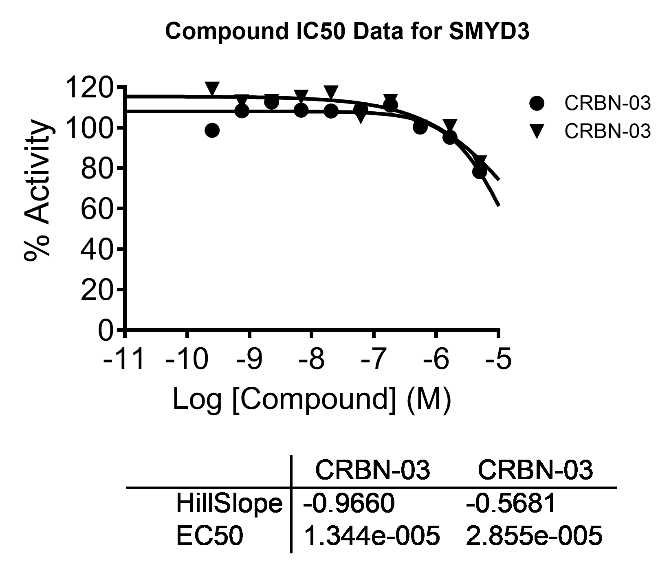

Dmax~20%

% Bioluminescence

**(B)**

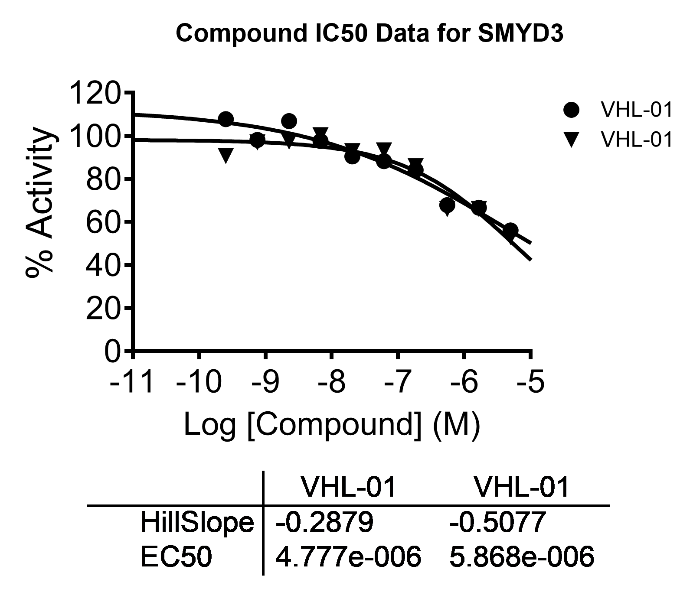

% Bioluminescence

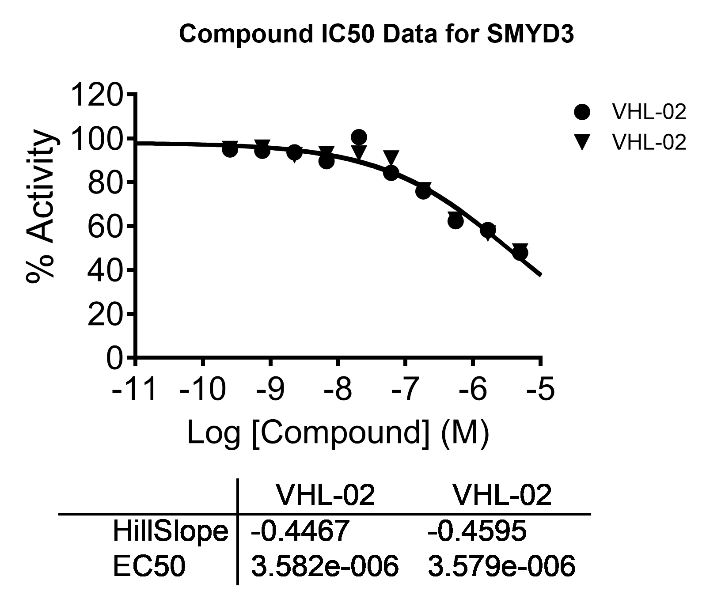

% Bioluminescence

Dmax~50%

Dmax~50%

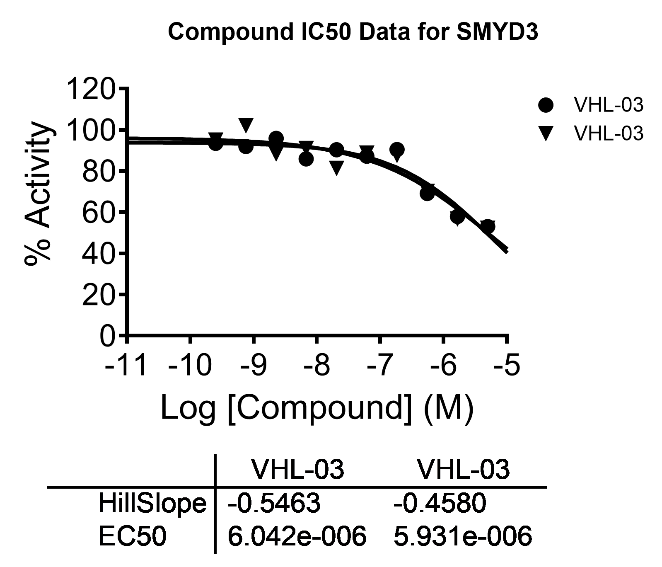

% Bioluminescence

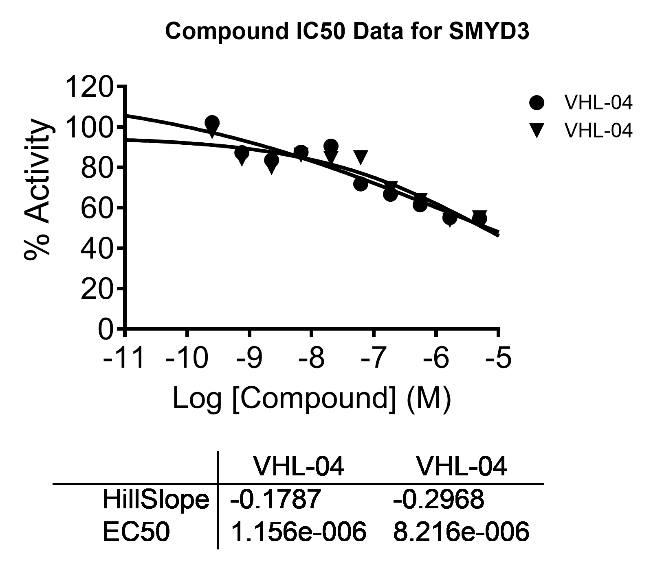

% Bioluminescence

Dmax~50%

Dmax~50%

**(C)**

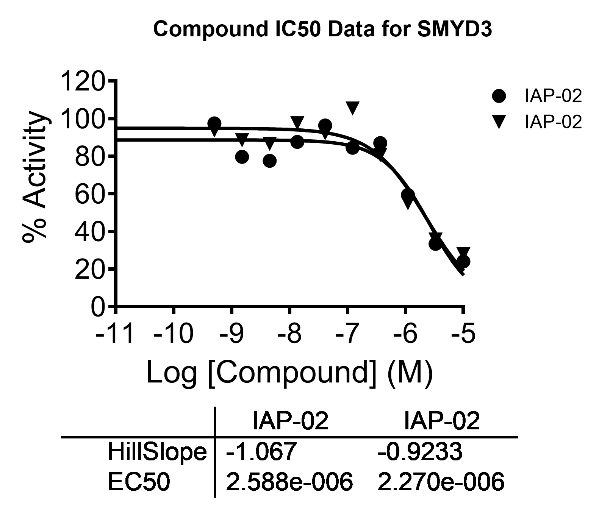

% Bioluminescence

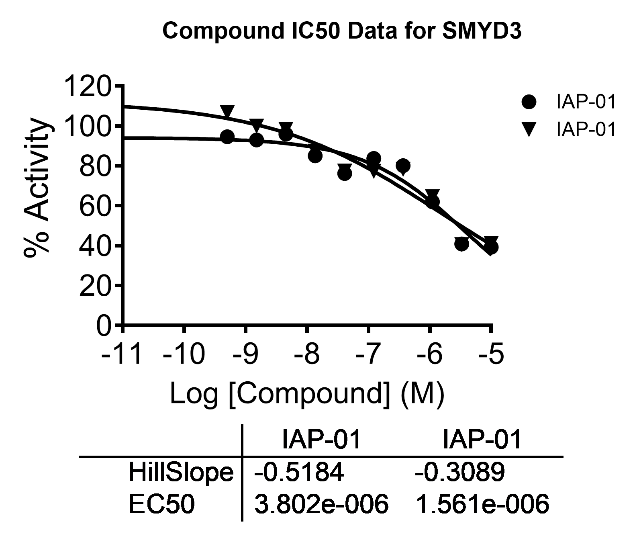

% Bioluminescence

Dmax~80%

Dmax~60%

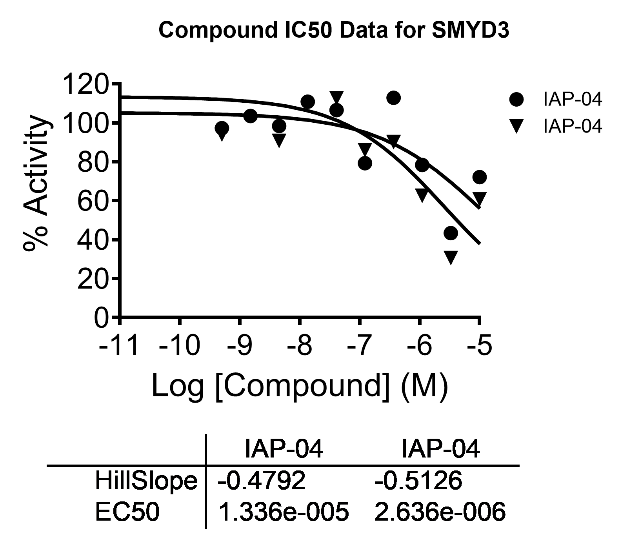

% Bioluminescence

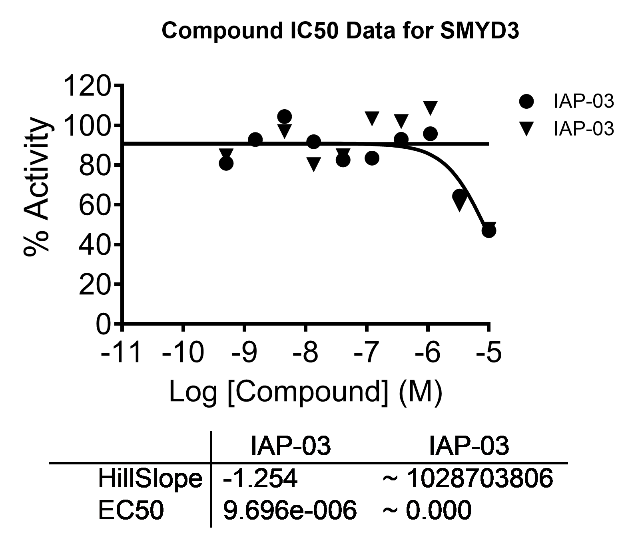

% Bioluminescence

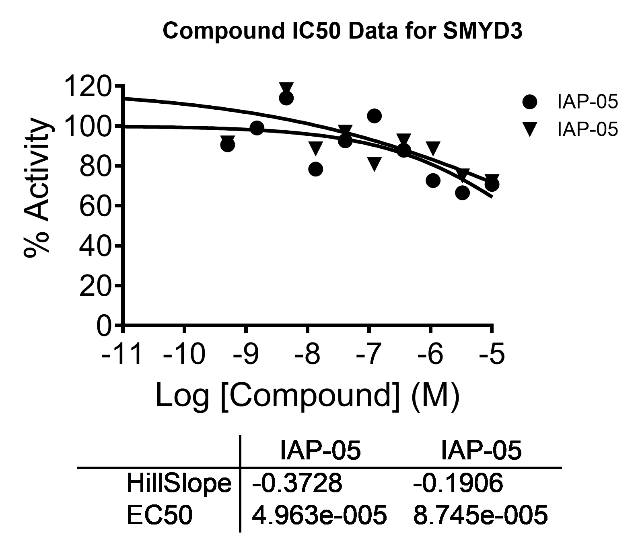

% Bioluminescence

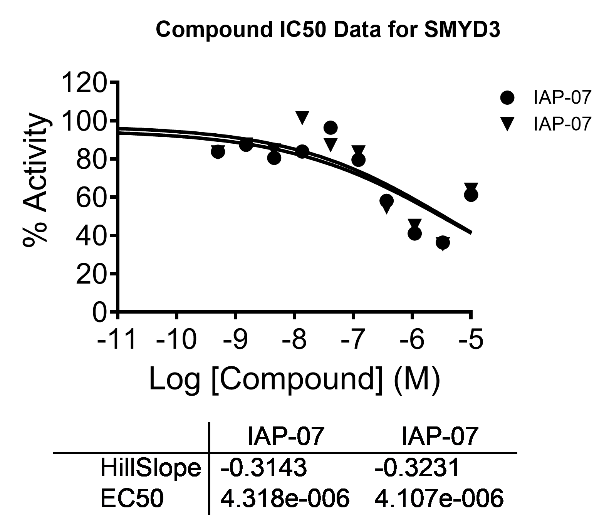

% Bioluminescence

Dmax~30%

Dmax~60%

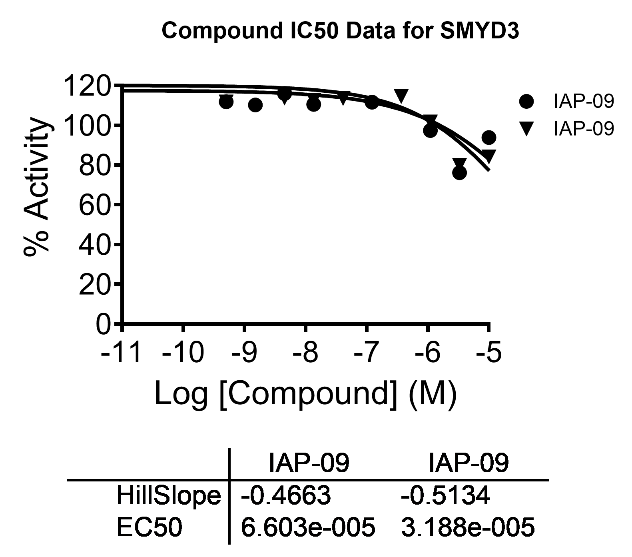

% Bioluminescence

Dmax~20%

| **Compound Code** | **DC50**  **(HiBiT assay, 24h, C at uM, HN13)** | **Dmax**  **(HiBiT assay, 24h, % degraded, HN13)** |
| --- | --- | --- |
| IAP-01 | 2.68 | 60% |
| IAP-02 | 2.43 | 80% |
| IAP-03 | repeat | pending |
| IAP-04 | 8 (repeat) | pending |
| IAP-05 | 68.5(repeat) | 30% |
| IAP-07 | 4.21 | 60% |
| IAP-08 | 0.17 | 90% |
| IAP-09 | 48.9 | 20% |
| CRBN-01 | 171 | 20% |
| CRBN-02 | 11 | 40% |
| CRBN-03 | 20 | 20% |
| VHL-01 | 5.32 | 50% |
| VHL-02 | 3.58 | 50% |
| VHL-03 | 5.98 | 50% |
| VHL-04 | 4.68 | 50% |

**(D)**
