## Supplementary Figure 14 to 16 for "SMYD3 protein degraders in HPV-negative head and neck squamous cell carcinoma"

**Supplementary Figure 14.** Pharmacokinetic (PK) assessment of the IAP-08 SMYD3 PROTAC in nude BALB/C mice. Mice (n=9) were administered a single IV dose of 12.5mg/kg of the PROTAC and plasma, urine and liver homogenates were obtained at 9 time points after the dose administration. **(A)** PK curve of IAP-08 plasma levels after a single IV dose of the PROTAC at 12.5mg/kg. Each time point represents the average of values obtained from 3 mice. **(B)** Concentration ratios of urine/plasma or liver/plasma of the SMYD3 PROTAC at ~4h, 8h and 24h after a single IV dose of the PROTAC at 12.5mg/kg. **(C)** PK parameters of IAP-08 after one single IV dose at 12.5mg/kg in mouse plasma samples. **(D)** PK parameters of IAP-08 after one single intraperitoneal (IP) dose at 10mg/kg in mouse plasma samples.

(A)
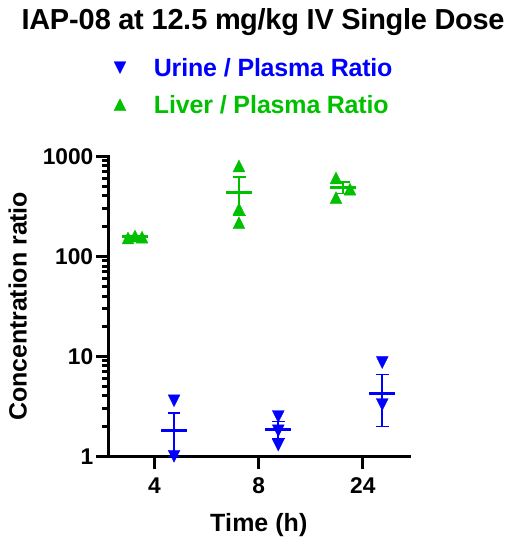

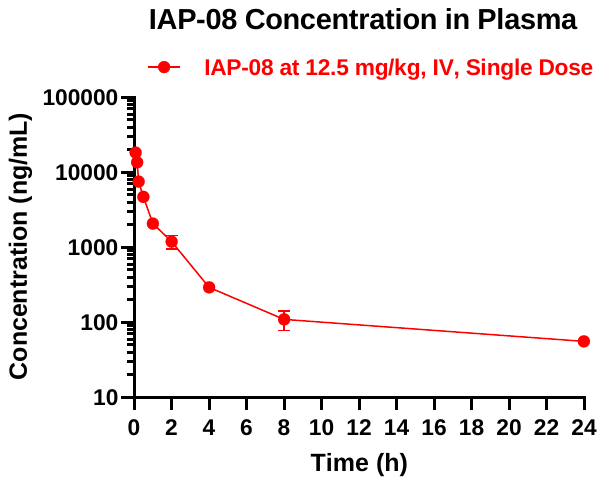
 (B)

(C)

| Time range (h) | t_1/2_ (h) | C_0_ (ng/ml) | AUC_last_ (h*ng/ml) | AUC_inf_ (h*ng/ml) | AUC/D (h*kg*ng/ml/mg) | AUC Extr (%) | MRT (h) | CL (ml/min/kg) | Vss (L/kg) |
| --- | --- | --- | --- | --- | --- | --- | --- | --- | --- |
| 4-24 | 9.71 | 24,731 | 12,542 | 13,326 | 1,066 | 5.89 | 4.47 | 15.63 | 4.19 |

(D)

| Time range (h) | T_max_  (h) | C_max_  (ng/ml) | AUC_last_ (h*ng/ml) |
| --- | --- | --- | --- |
| 4-24 | 4 | 917 | 12,808 |

**Supplementary Figure 15.** In vitro metabolic assays for IAP-08**. (A)** Protein binding assay results. Equilibrium dialysis was conducted using human or mouse plasma and the compound at a concentration of 1e-05M. Duplicate samples were incubated for 4h at 37oC and high performance liquid chromatography(HPLC)-MS/MS analysis was pursued to quantify the unbound compound. **(B)** In vitro liver microsome intrinsic clearance assay results. The compound was incubated at a concentration of 1e-07M with human or mouse liver microsomes and NAPDH at 37oC for various time periods (0, 15min, 30min, 45min, 60min) and was quantified using HPLC-MS/MS. Results are shown for each duplicate sample.

**(A)**

**Human plasma protein binding assay results:**

| **% Protein bound 1^st^** | **% Protein bound 2^nd^** | **Mean** | **% Recovery 1^st^** | **% Recovery 2^nd^** | **Mean** |
| --- | --- | --- | --- | --- | --- |
| 99.4 | 99.5 | 99.45 | 54.59 | 56.8 | 55.7 |

**Mouse plasma protein binding assay results:**

| **% Protein bound 1^st^** | **% Protein bound 2^nd^** | **Mean** | **% Recovery 1^st^** | **% Recovery 2^nd^** | **Mean** |
| --- | --- | --- | --- | --- | --- |
| 99.79 | 98.32 | 99.06 | 70.71 | 70.06 | 70.39 |

**(B)**

**Human liver microsome assay results:**

| **Incubation times (min)** | **% Compound remaining 1^st^** | **% Compound remaining 2^nd^** | **Mean** | **t 1/2 (min)**  **1^st^** | **t 1/2 (min)**  **2^nd^** | **Mean** | **Intrinsic clearance (uL/min/mg)** |
| --- | --- | --- | --- | --- | --- | --- | --- |
| 0 | 100 | 100 | 100 | 64.6 | 78.8 | 71.7 | 97.6 |
| 15 | 90.9 | 88.39 | 89.64 |  |  |  |  |
| 30 | 79.1 | 77.28 | 78.19 |  |  |  |  |
| 45 | 63.68 | 67.9 | 65.79 |  |  |  |  |
| 60 | 53.44 | 58.97 | 56.21 |  |  |  |  |

**Mouse liver microsome assay results:**

| **Incubation times (min)** | **% Compound remaining 1^st^** | **% Compound remaining 2^nd^** | **Mean** | **t 1/2 (min)**  **1^st^** | **t 1/2 (min)**  **2^nd^** | **Mean** | **Intrinsic clearance (uL/min/mg)** |
| --- | --- | --- | --- | --- | --- | --- | --- |
| 0 | 100 | 100 | 100 | 83.6 | 72.7 | 78.2 | 89.1 |
| 15 | 95.33 | 86.92 | 91.13 |  |  |  |  |
| 30 | 81.12 | 79.02 | 80.07 |  |  |  |  |
| 45 | 71.1 | 67.35 | 69.23 |  |  |  |  |
| 60 | 62.16 | 55.59 | 58.88 |  |  |  |  |

**Supplementary Figure 16.** Heatmap of top 20 type I IFN response genes (Hallmark) with the greatest log2 fold change induced by IAP-08 (0.5uM) or EPZ031686 (10uM) versus DMSO for 48h (HN-SCC-151 cells). The ratio of up- over down-regulated genes represents the one observed in the respective Hallmark gene list. FDR<0.1.

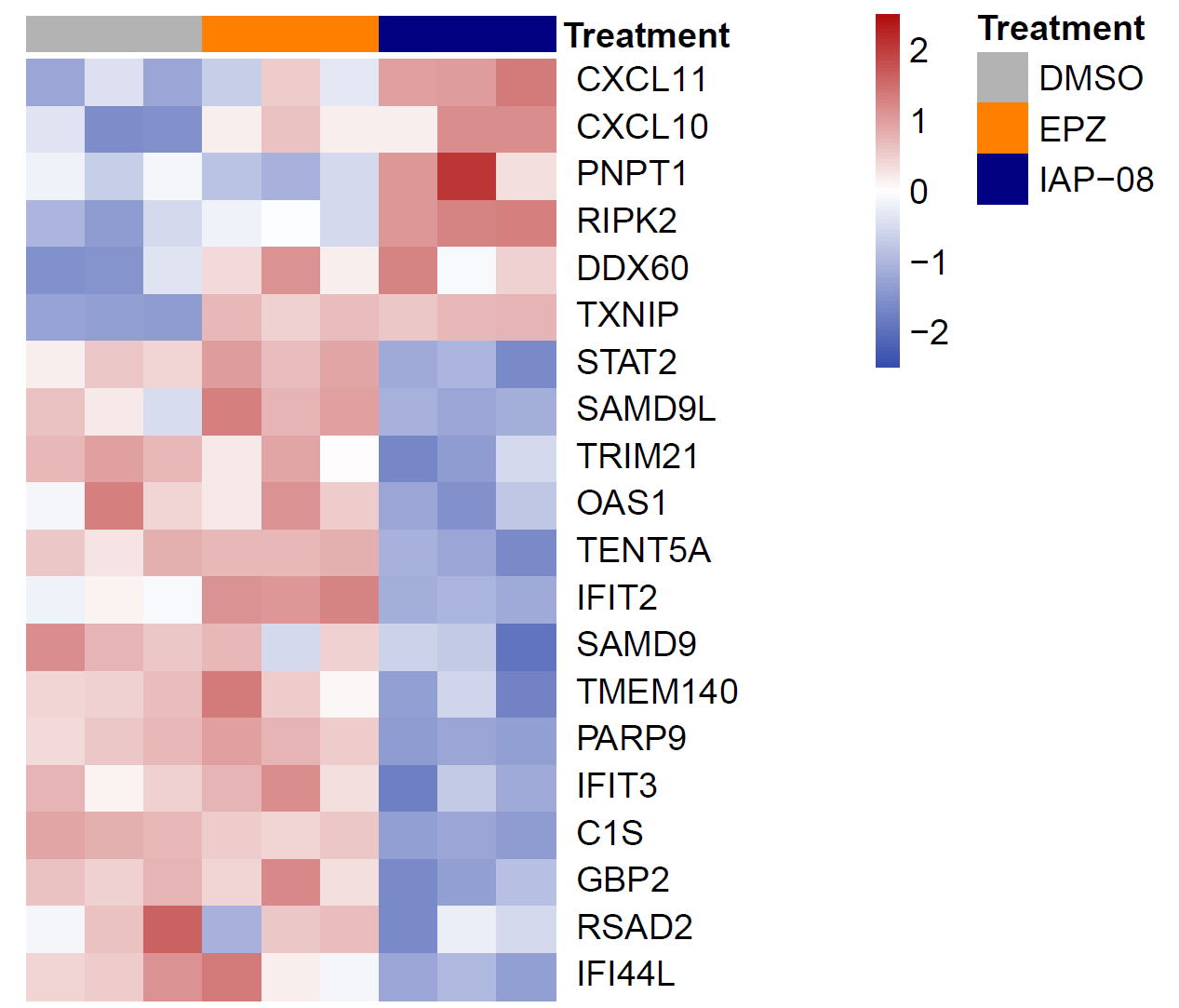
