## Supplementary Table 1 for "SMYD3 protein degraders in HPV-negative head and neck squamous cell carcinoma"

**Supplementary Table 1.** SMYD3 PROTAC structures with different E3 ligase ligands and bis-carboxyl linkers using EPZ031686 propyl amine as SMYD3 ligand.

| **Compound** | **SMYD3 ligand** | **Linker** | **E3 Ligase ligand** |
| --- | --- | --- | --- |
| IAP-01 | 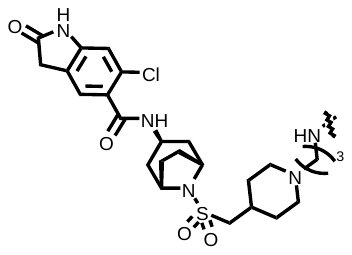 | 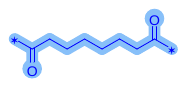 | 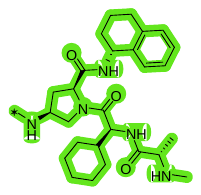  IAP |
| IAP-02 |  | 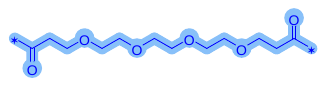 |  |
| IAP-03 |  | 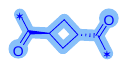 |  |
| IAP-04 |  | 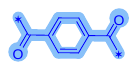 |  |
| IAP-05 |  | 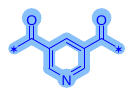 |  |
| IAP-07 |  | 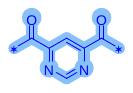 |  |
| IAP-08 |  | 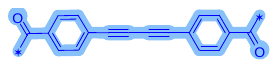 |  |
| IAP-09 |  | 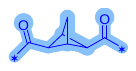 |  |
| VHL-01 |  | 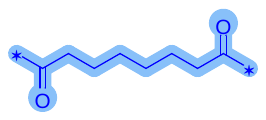 | 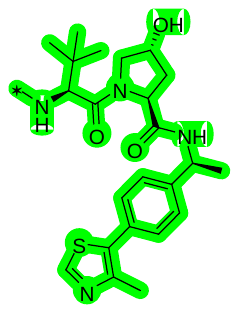  VHL |
| VHL-02 |  | 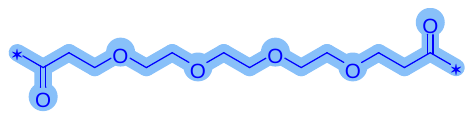 |  |
| VHL-03 |  | 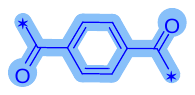 |  |
| VHL-04 |  | 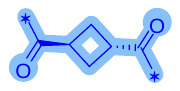 |  |
| CRBN-01 |  | 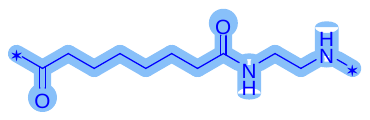 | 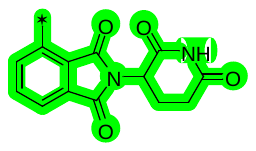  CRBN |
| CRBN-02 |  | 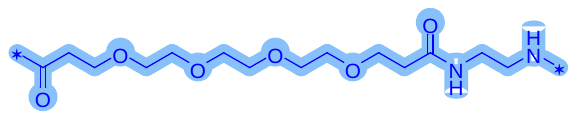 |  |
| CRBN-03 |  | 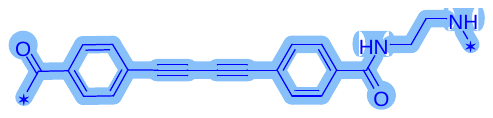 |  |
