## Supplementary Table 9 for "SMYD3 protein degraders in HPV-negative head and neck squamous cell carcinoma"

**Supplementary Table 7.** NMR spectra and UPLC-MS data of selected newly synthesized SMYD3 PROTAC compounds.

**^1^H NMR (500 MHz, DMSO-d6) spectrum of EPZ031686-IAP-01**

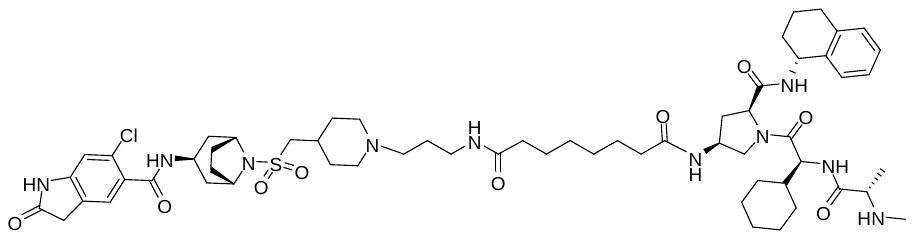

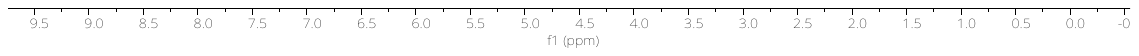

**UPLC-MS Data -EPZ031686-IAP-01**

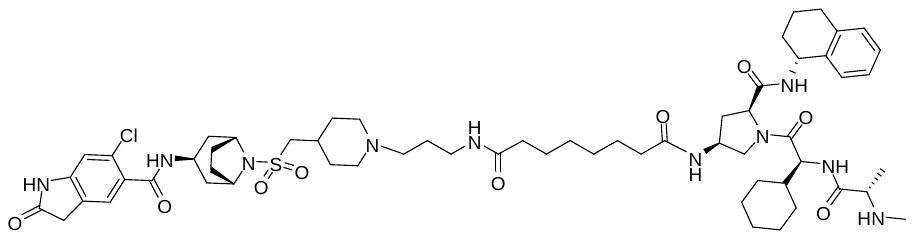

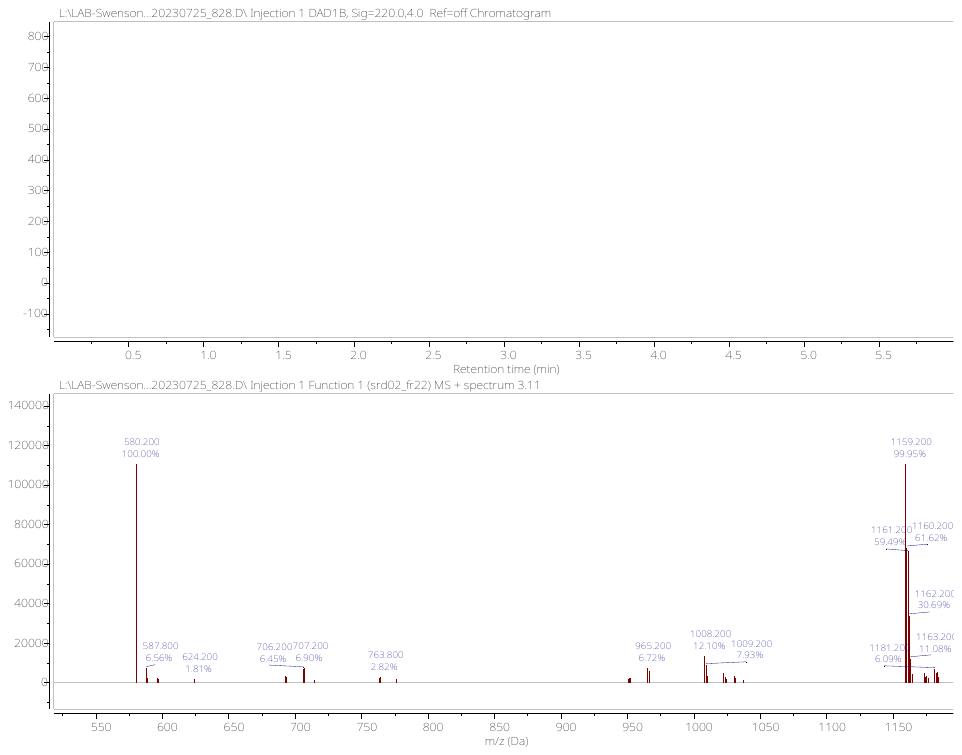

**^1^H NMR (500 MHz, MeOD-d4) spectrum of EPZ031686-IAP-02**

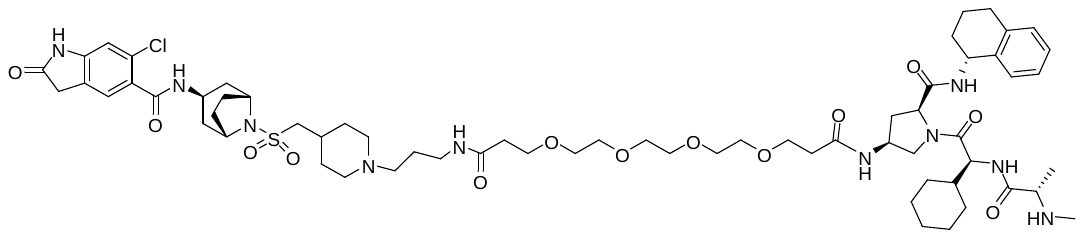

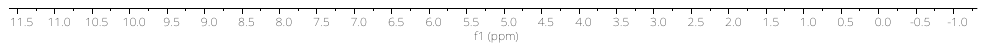

**UPLC-MS Data -EPZ031686-IAP-02**

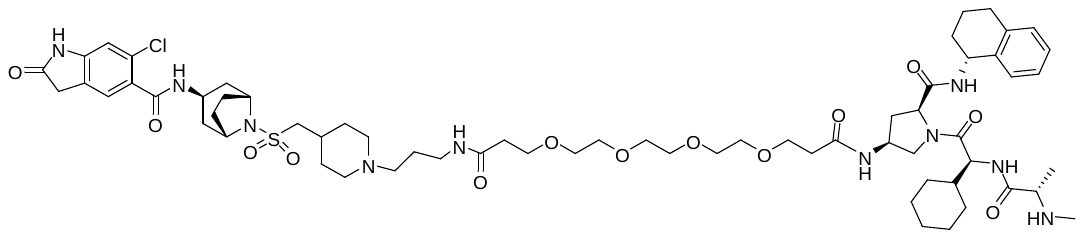

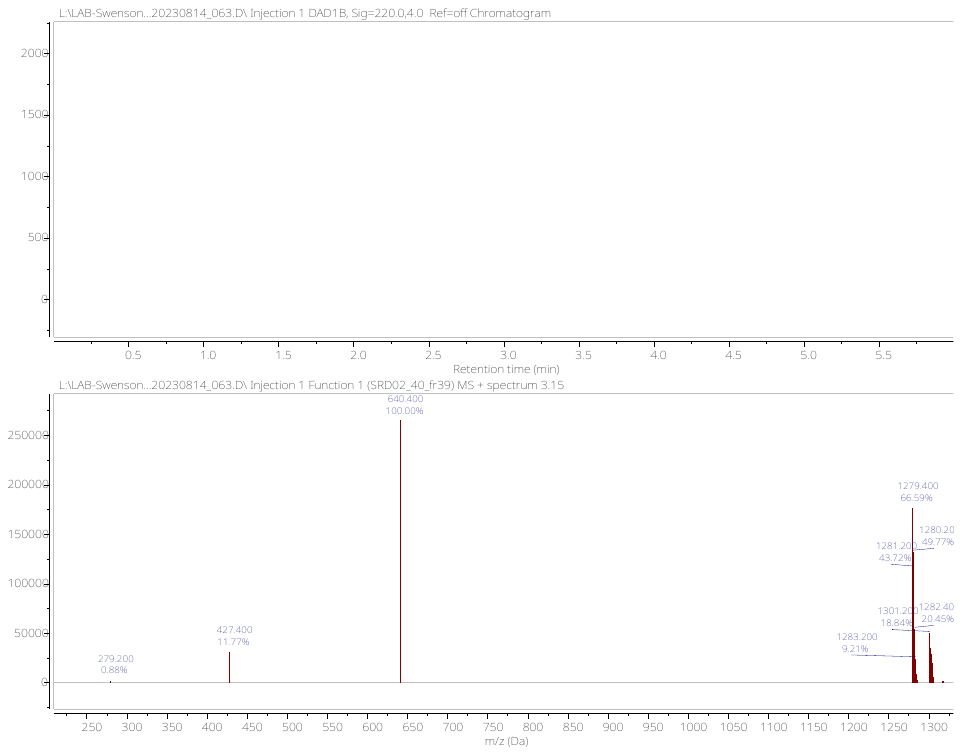

**^1^H NMR (500 MHz, MeOD-d4) spectrum of EPZ031686-IAP-03**

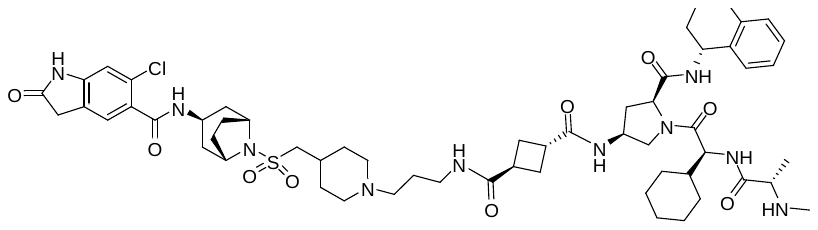

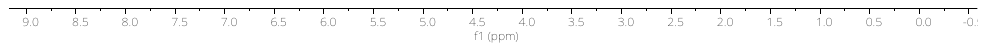

**UPLC-MS Data -EPZ031686-IAP-03**

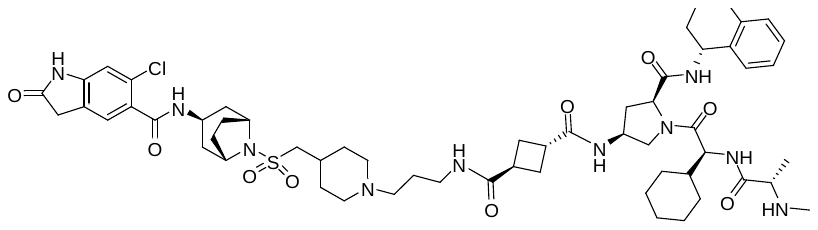

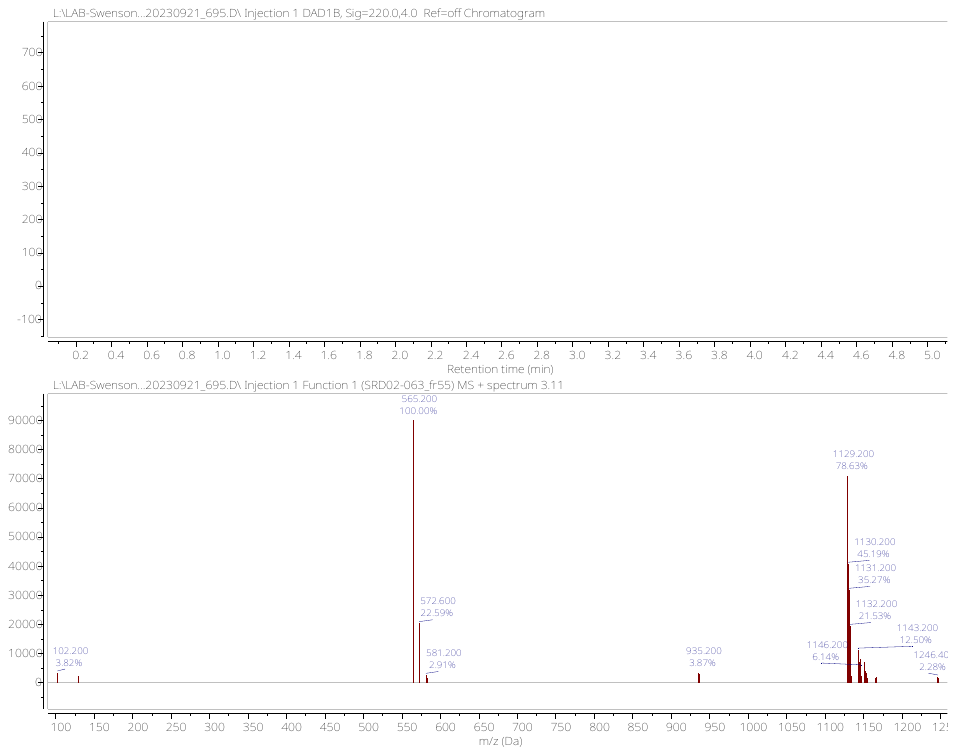

**^1^H NMR (500 MHz, MeOD-d4) spectrum of EPZ031686-IAP-04**

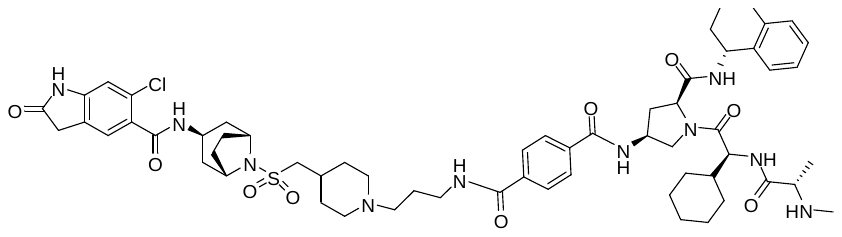

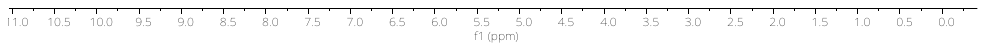

**UPLC-MS Data -EPZ031686-IAP-04**

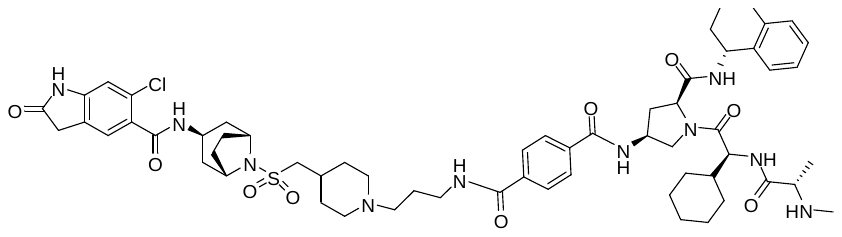

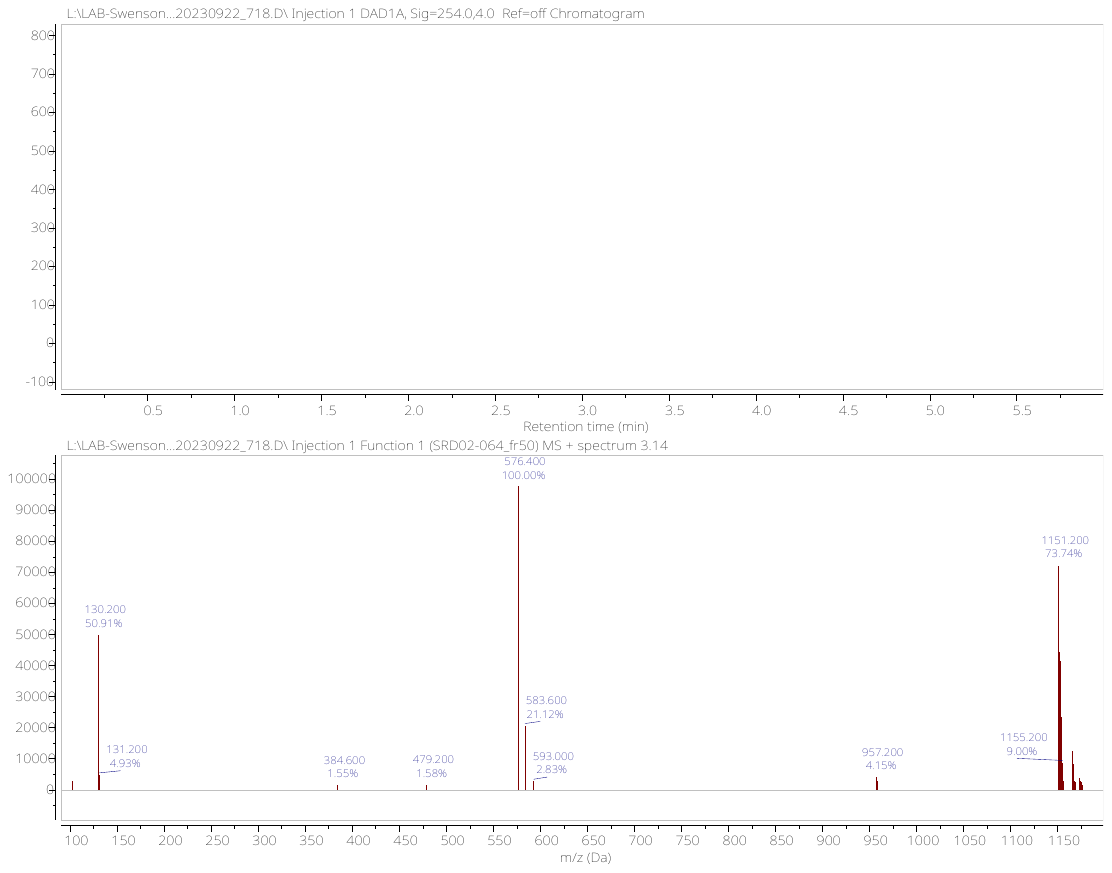

**^1^H NMR (400 MHz, MeOD-d4) spectrum of EPZ031686-IAP-05**

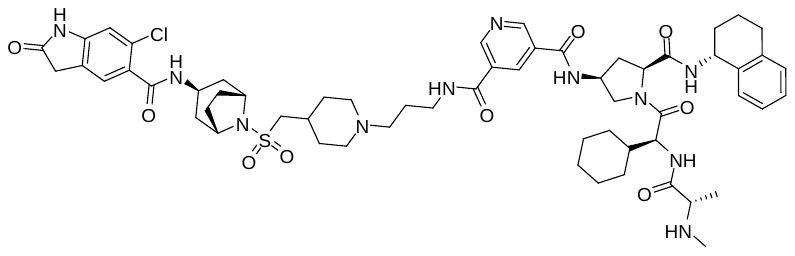

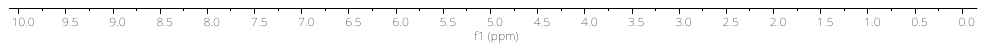

**UPLC-MS Data -EPZ031686-IAP-05**

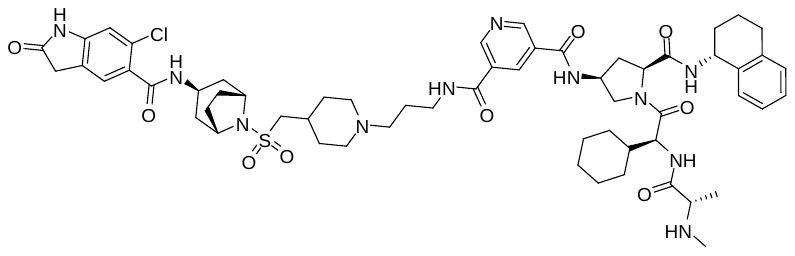

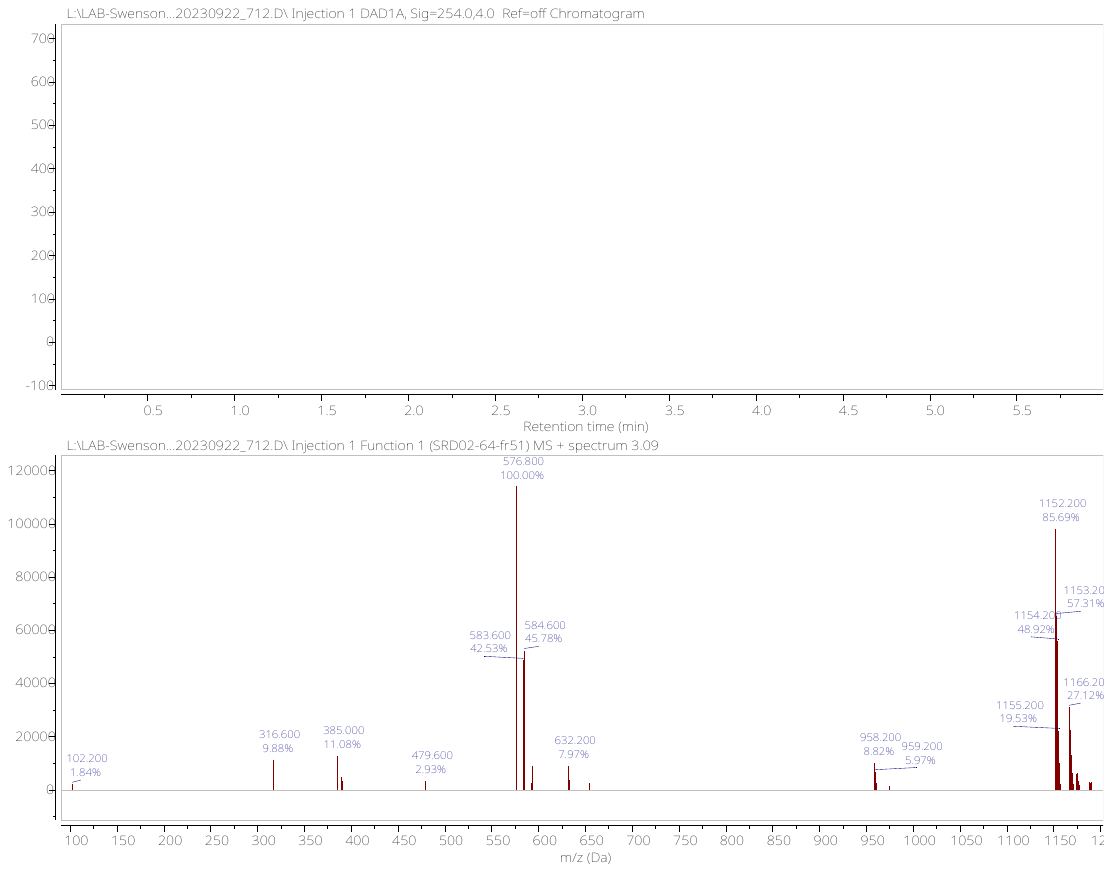

**^1^H NMR (500 MHz, DMSO-d6) spectrum of EPZ031686-IAP-07**

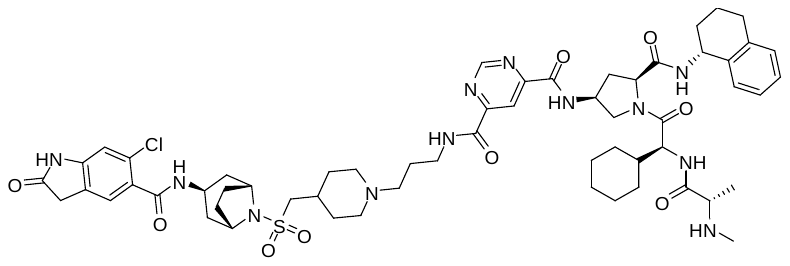

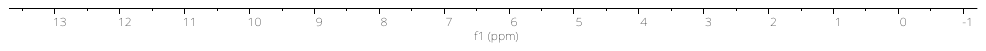

**UPLC-MS Data -EPZ031686-IAP-07**

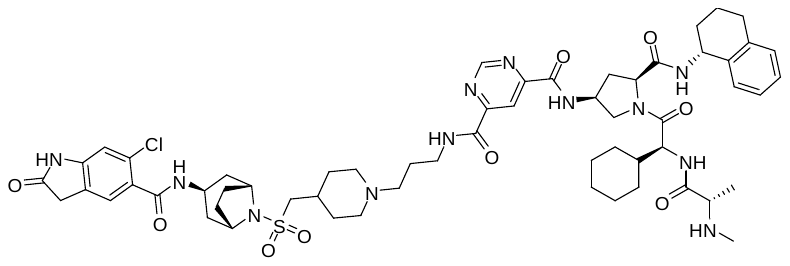

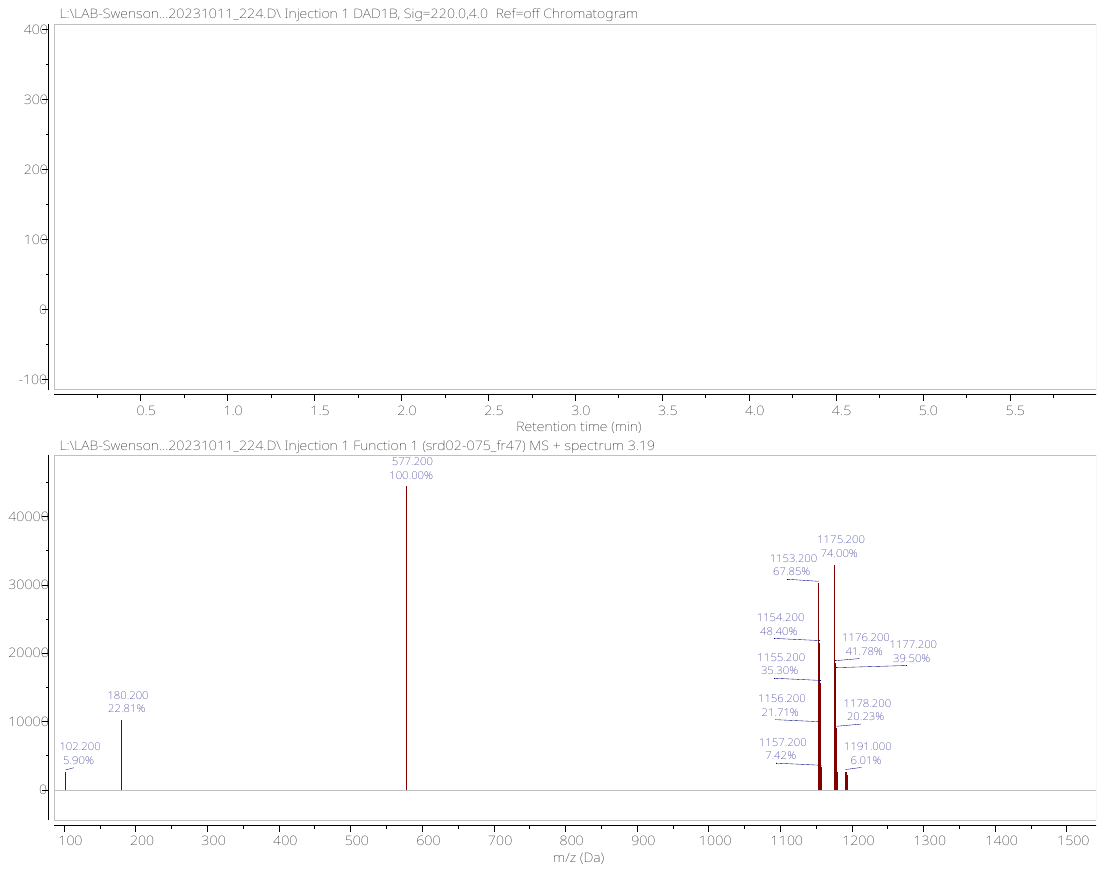

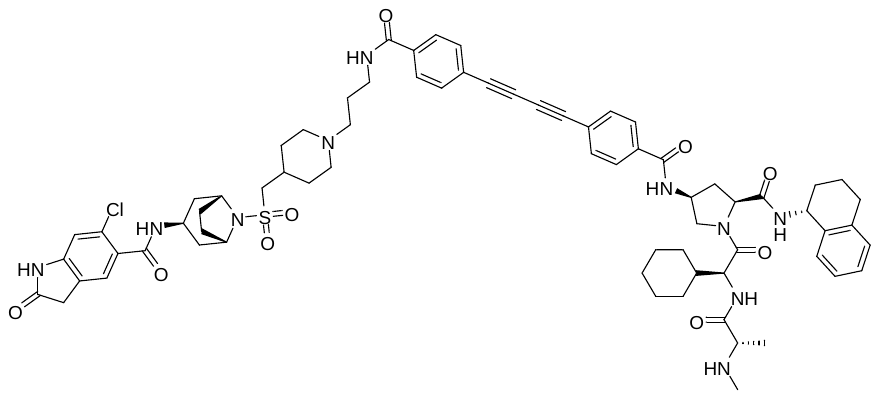
**H^1^ NMR (500 MHz, DMSO-d6) spectrum of EPZ031686-IAP-08**

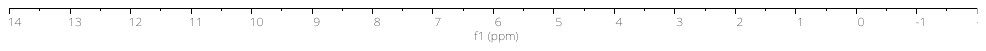

**UPLC-MS Data -EPZ031686-IAP-08**

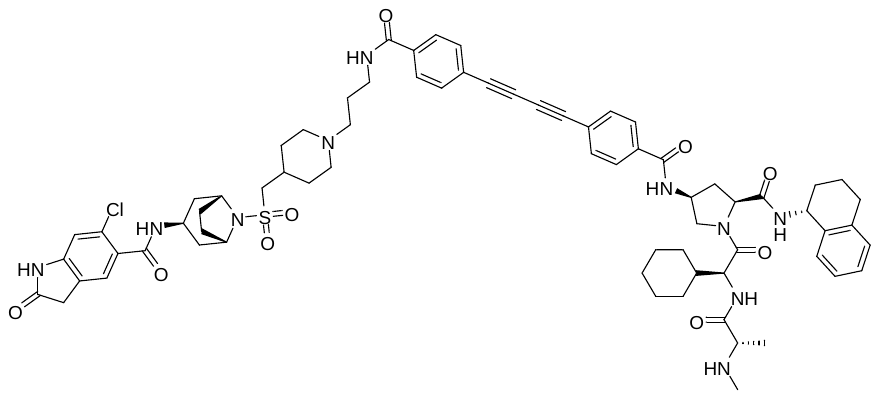

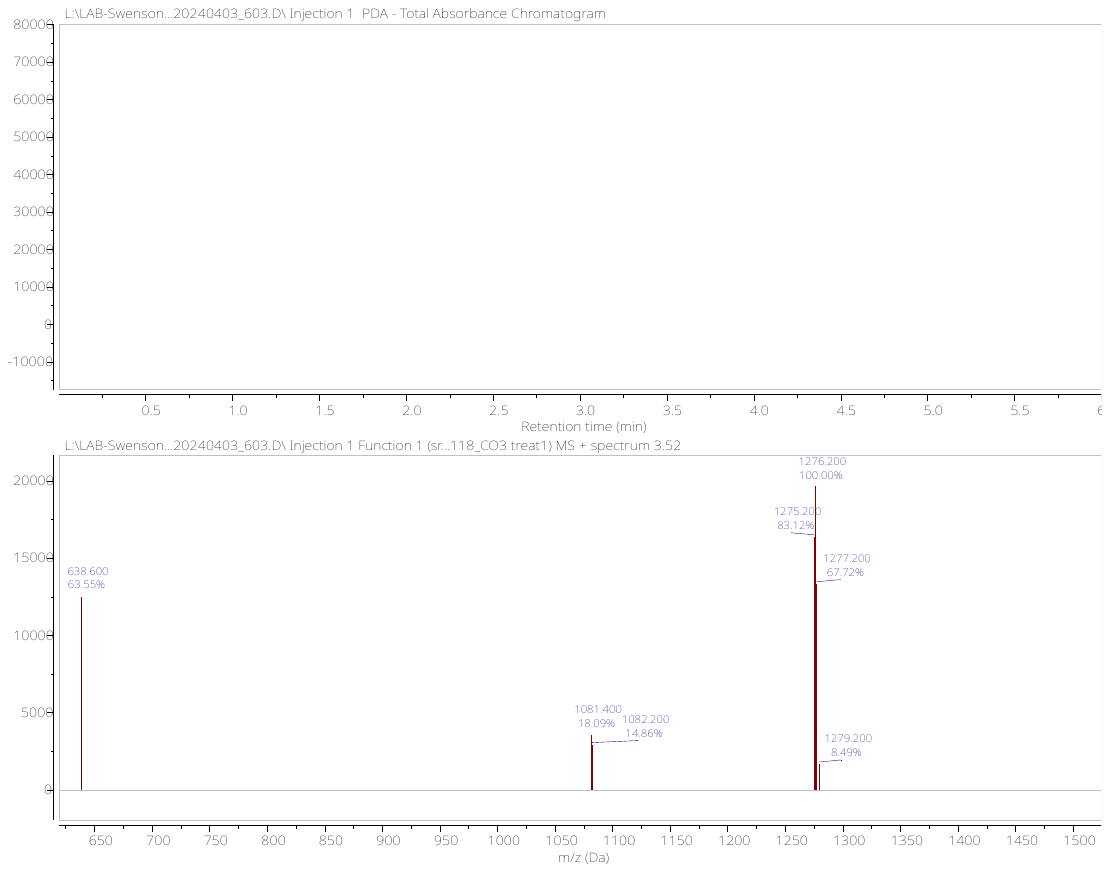

**H^1^ NMR (500 MHz, DMSO-d6) spectrum of EPZ031686-IAP-09**

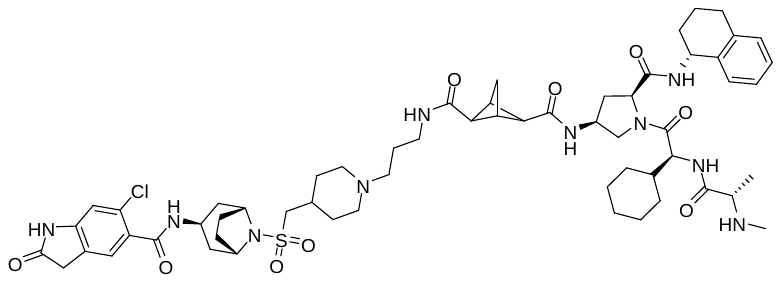

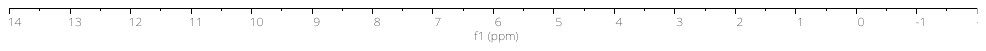

**UPLC-MS Data -EPZ031686-IAP-09**

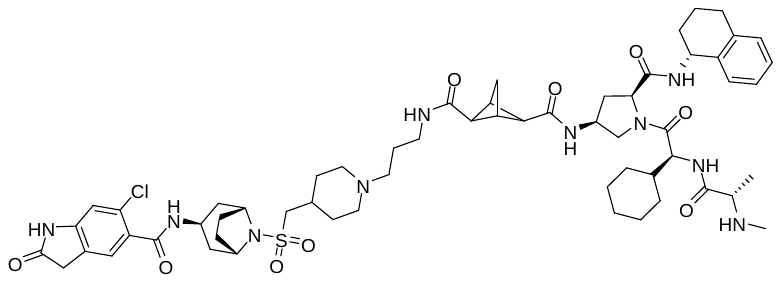

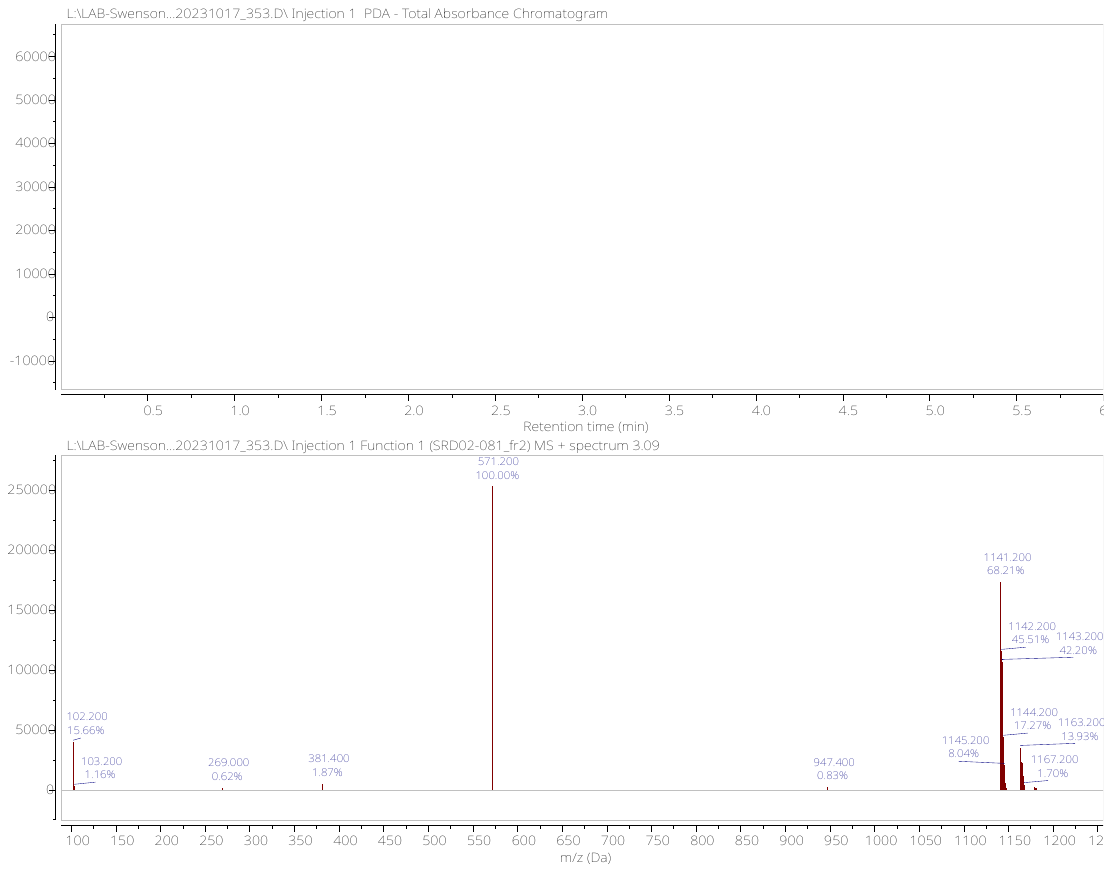

**^1^H NMR (500 MHz, MeOD-d4) spectrum of EPZ031686-IAP-10**

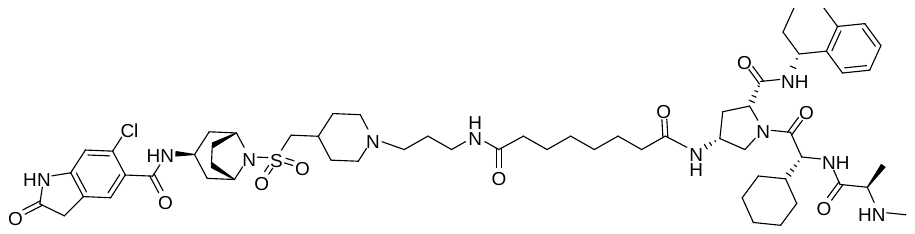

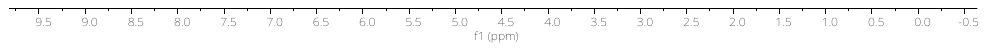

**UPLC-MS of EPZ031686-IAP-10**

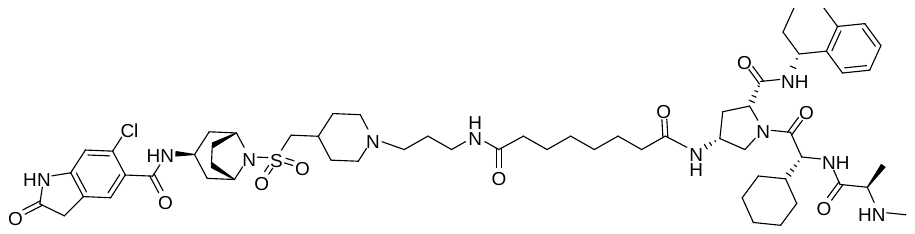

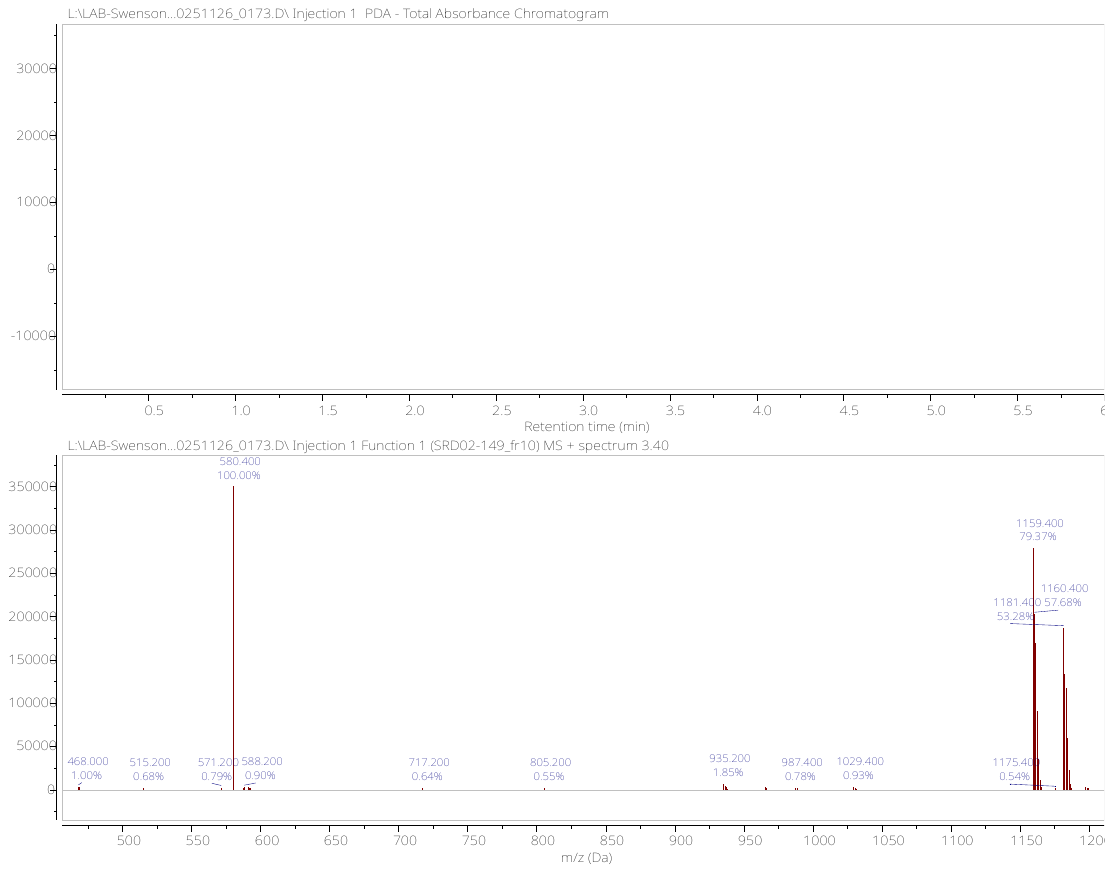

**^1^H NMR (500 MHz, DMSO-d6) spectrum of EPZ031686-IAP-11**

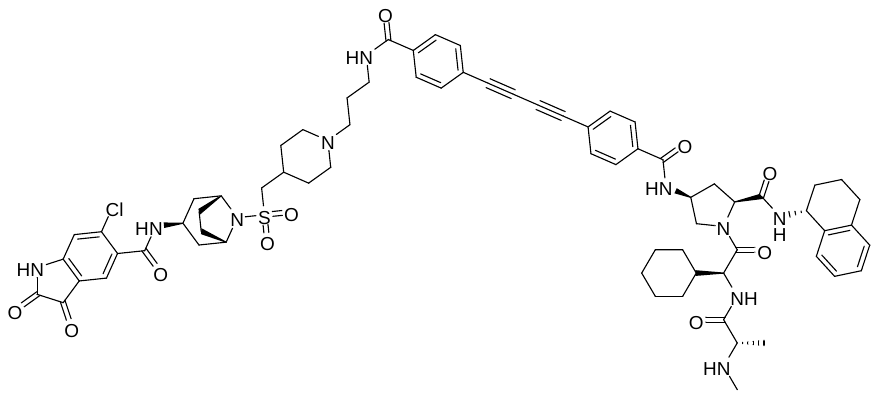

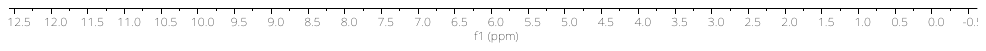

**UPLC-MS of EPZ031686-IAP-11**

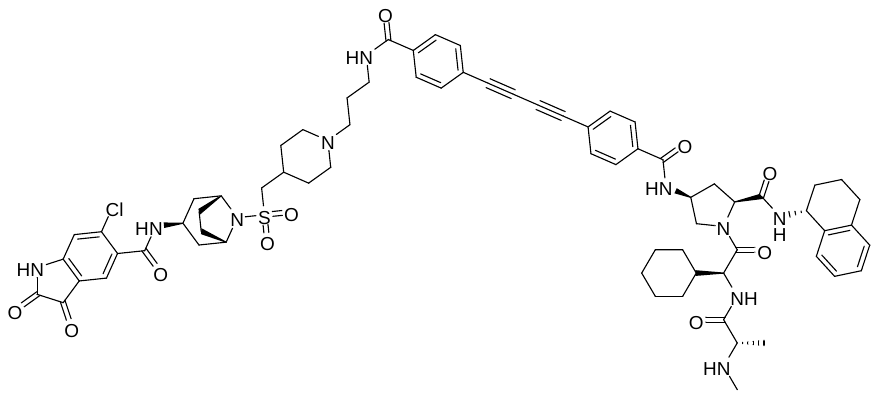

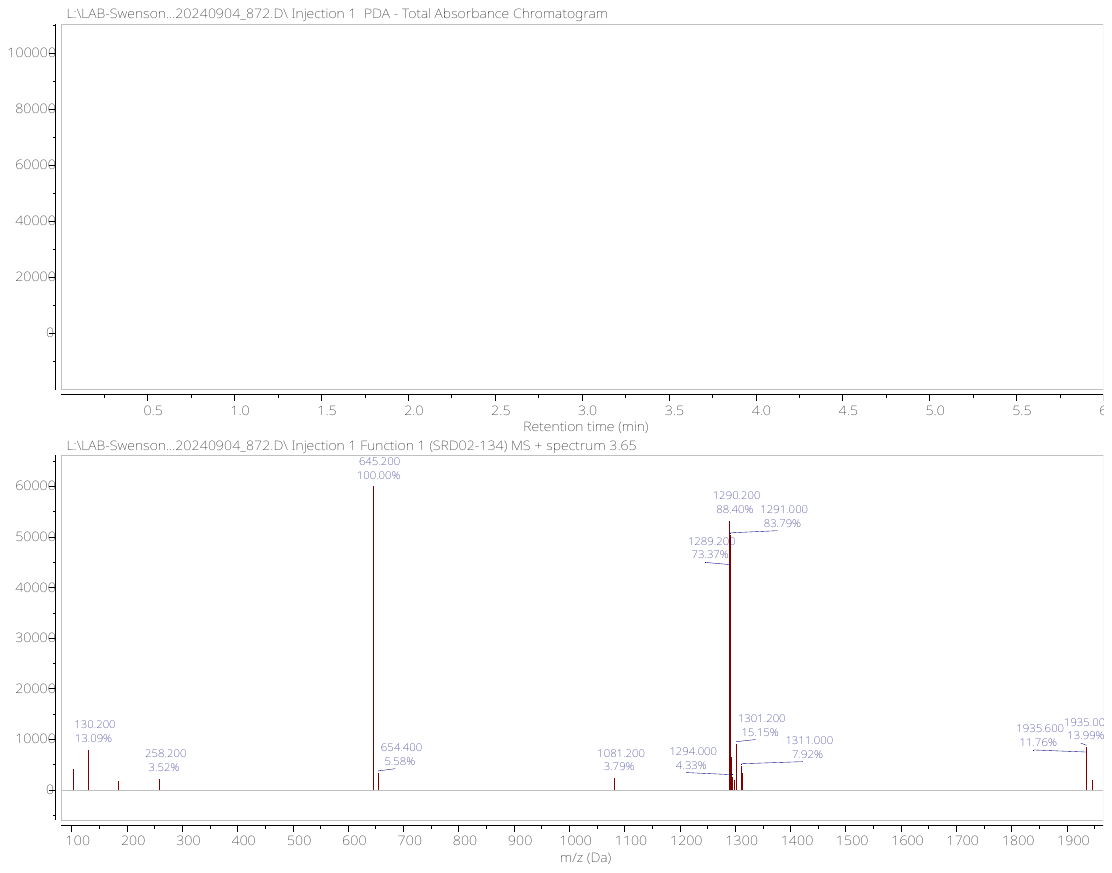

**^1^H NMR (500 MHz, MeOD-d6) spectrum of EPZ031686-CRBN-01**

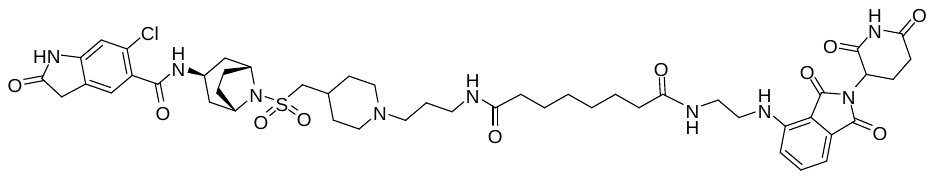

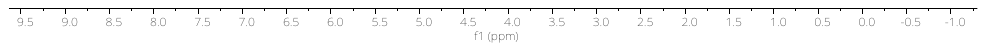

**UPLC-MS of EPZ031686-CRBN-01**

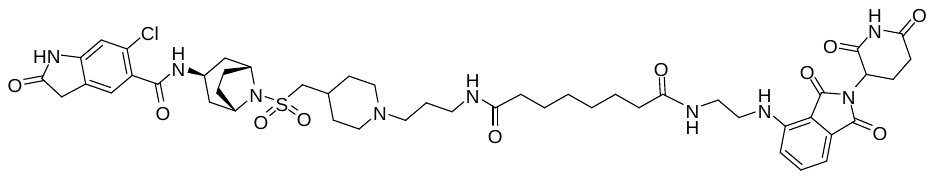

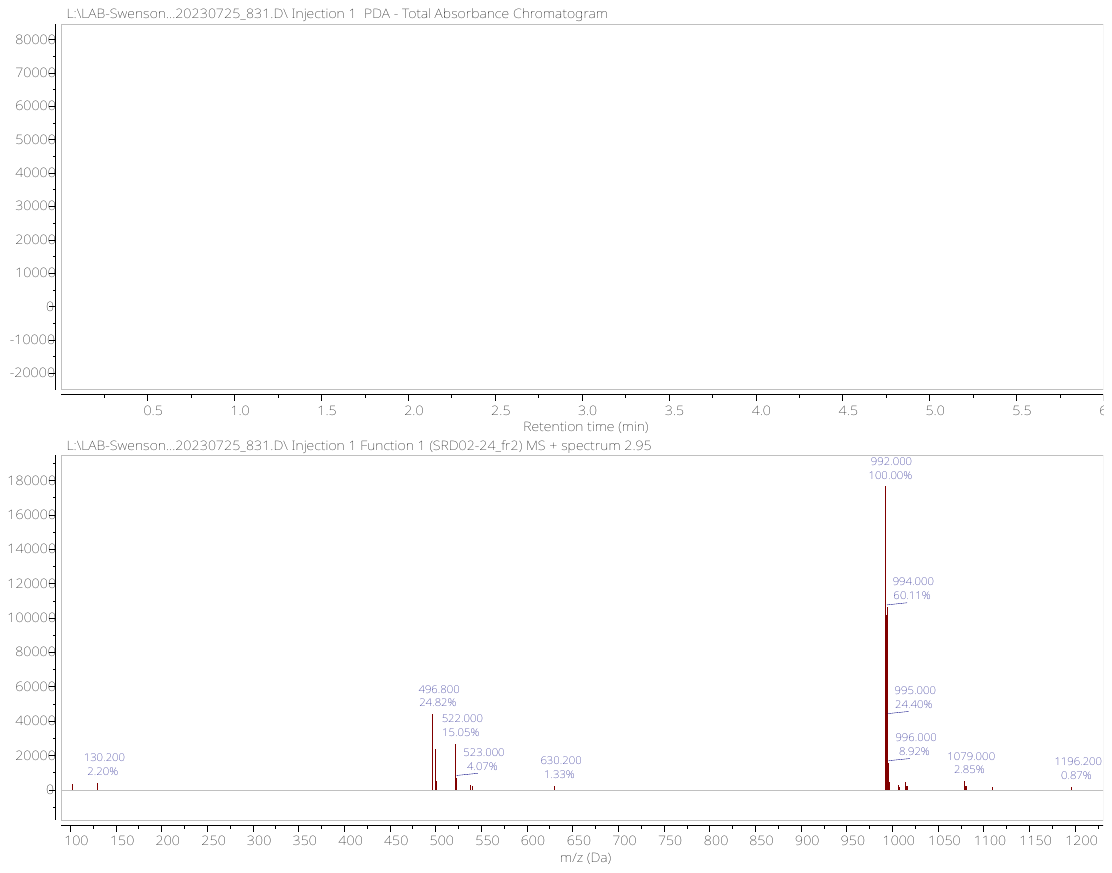

**^1^H NMR (500 MHz, MeOD-d4) spectrum of EPZ031686-CRBN-02**

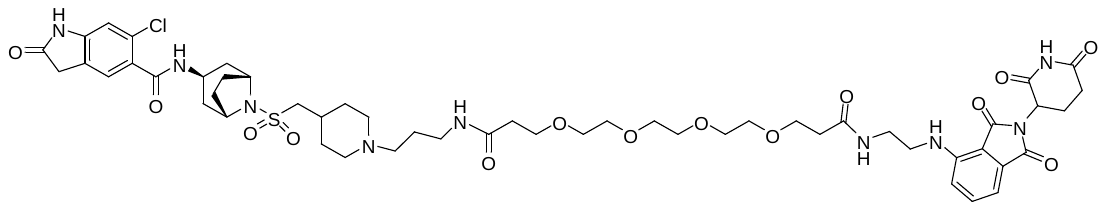

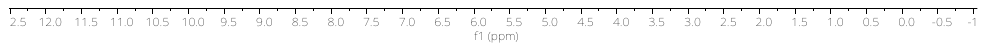

**UPLC-MS of EPZ031686-CRBN-02**

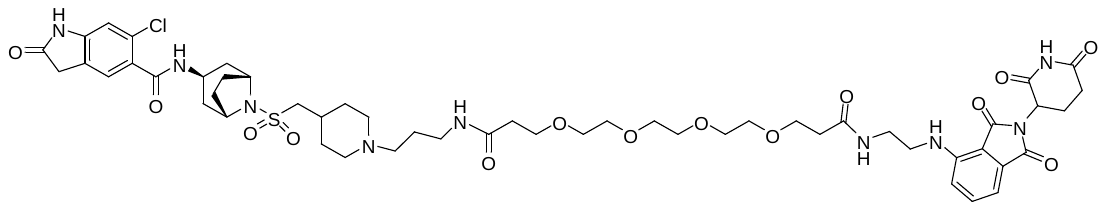

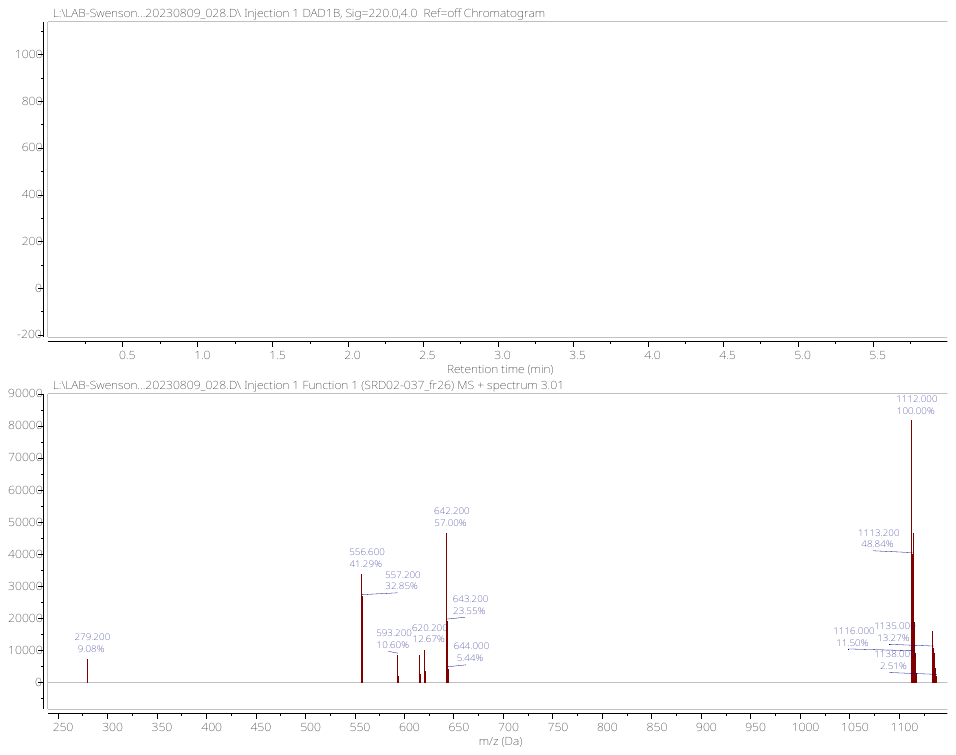

**^1^H NMR (500 MHz, DMSO-d6) spectrum of EPZ031686-CRBN-03**

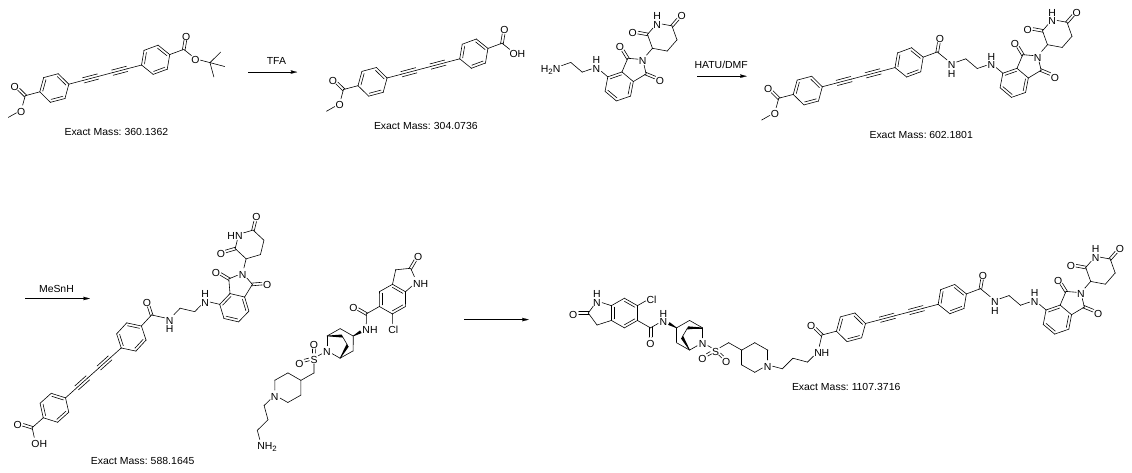

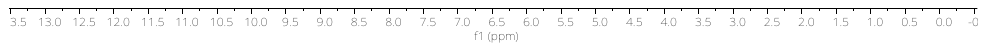

**UPLC-MS of EPZ031686-CRBN-03**

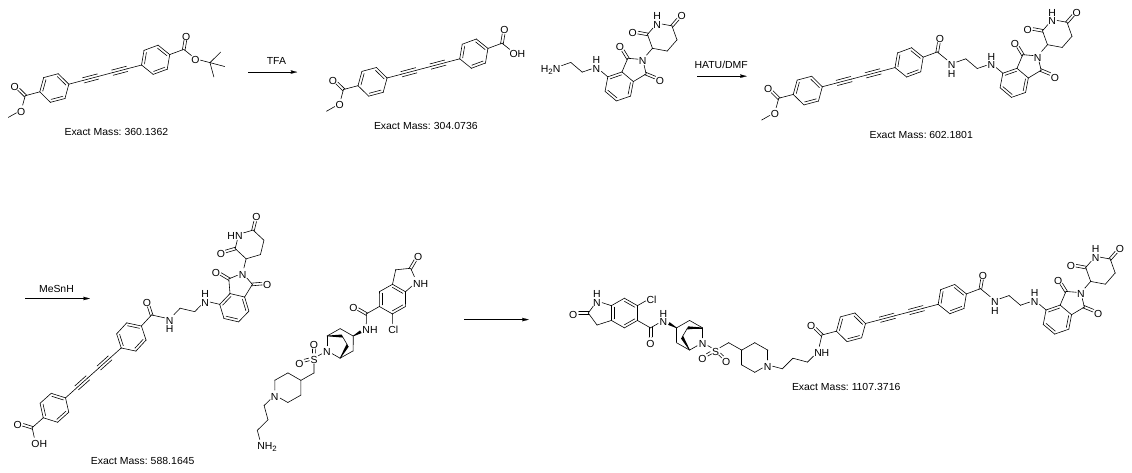

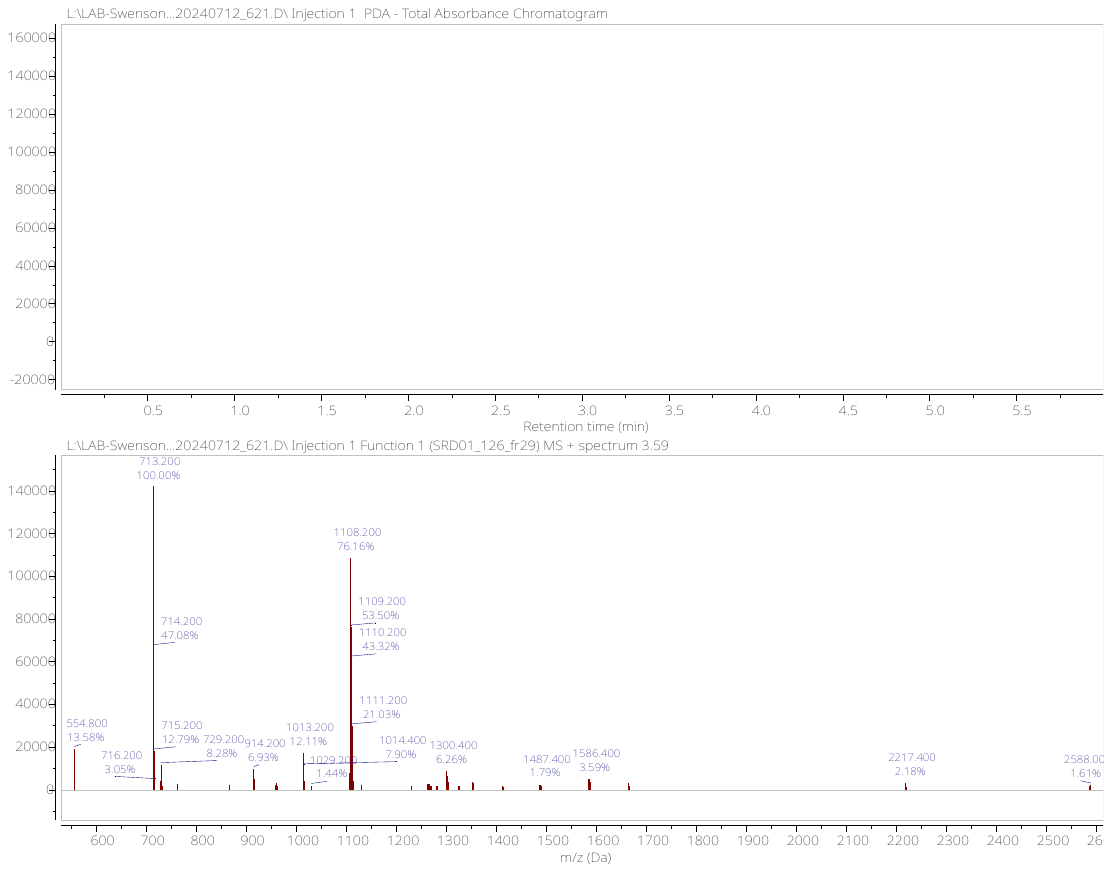

**^1^H NMR (500 MHz, MeOD-d4) spectrum of EPZ031686-CRBN-04**

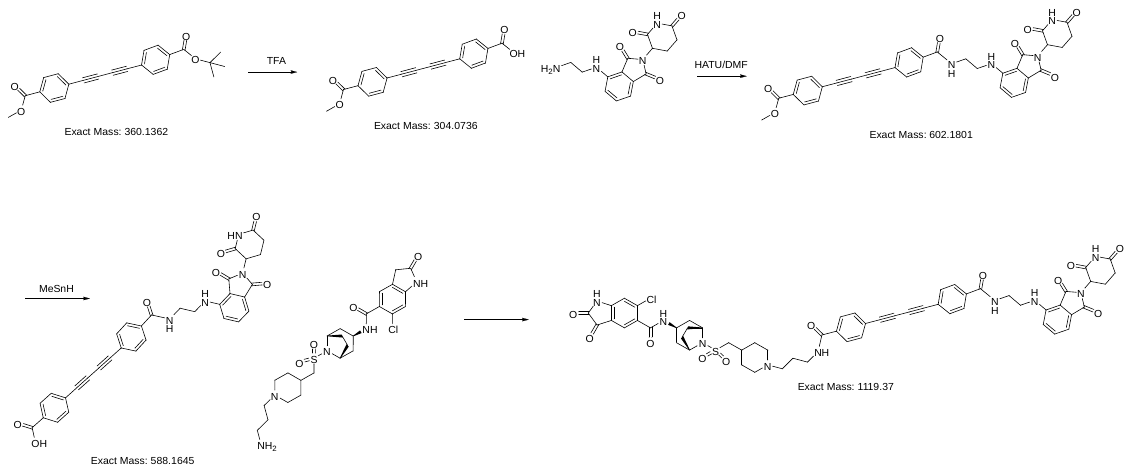

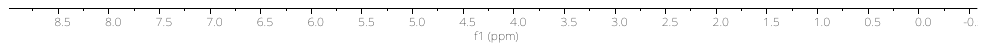

**UPLC-MS of EPZ031686-CRBN-04**

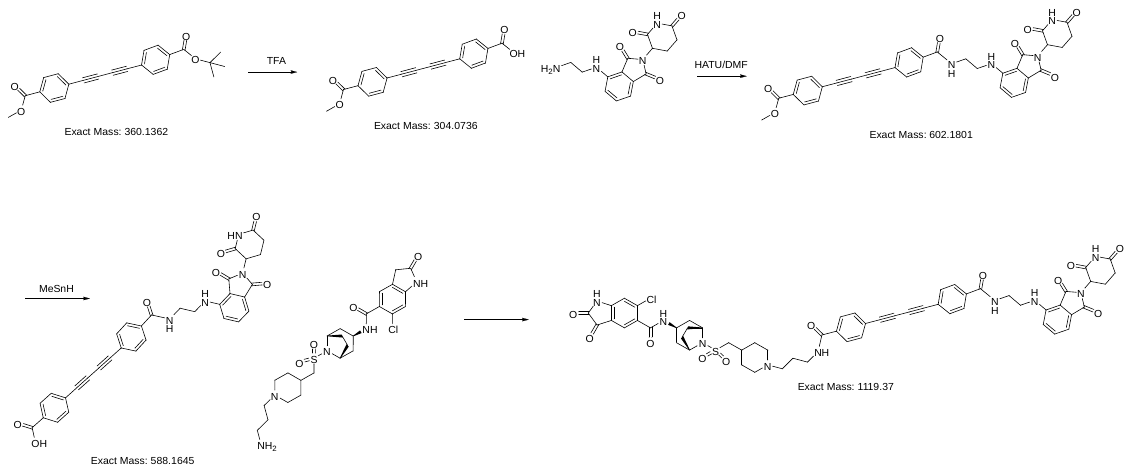

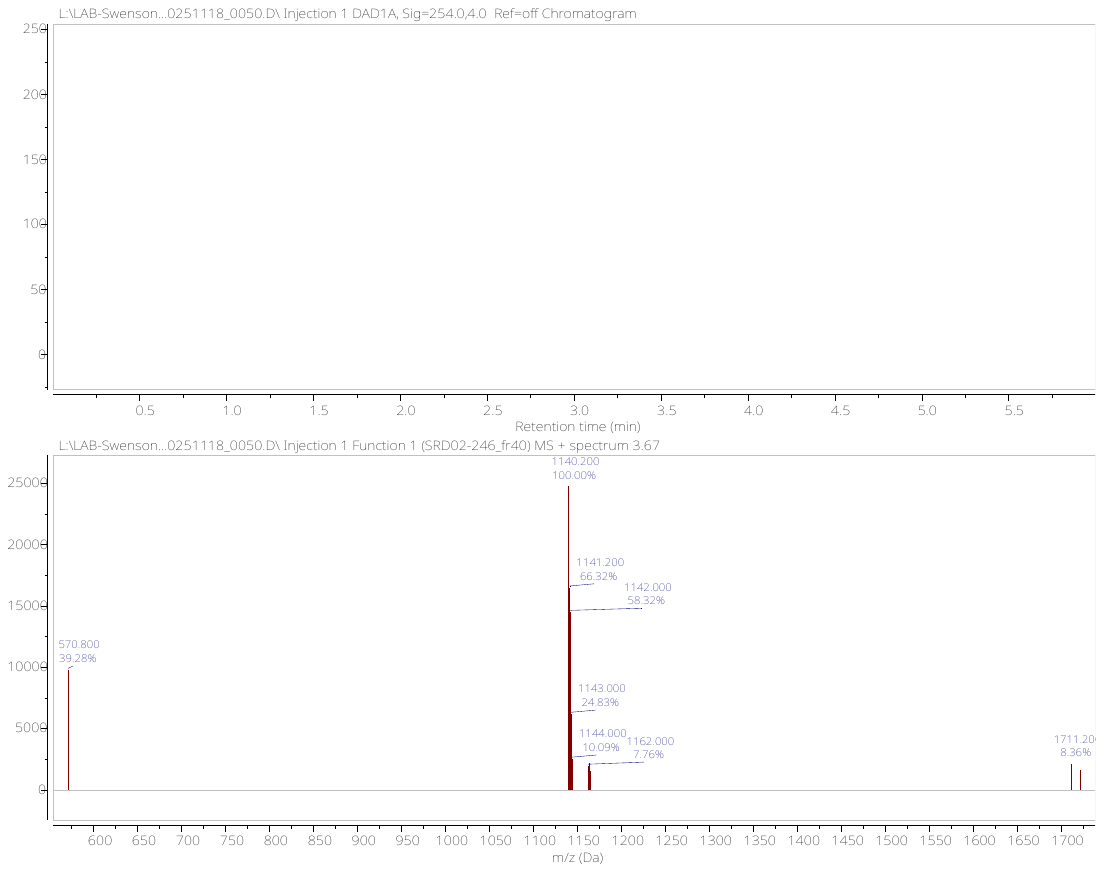

**^1^H NMR (500 MHz, DMSO-d6) spectrum of EPZ031686-CRBN-05**

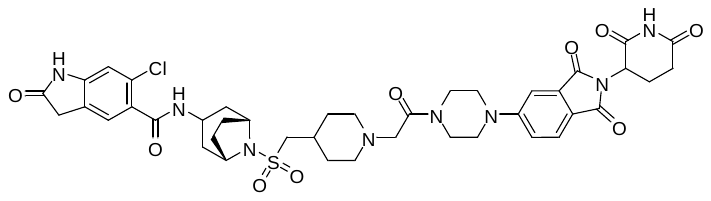

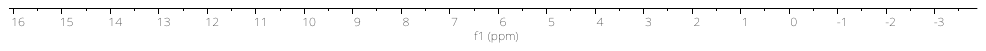

**UPLC-MS of EPZ031686-CRBN-05**

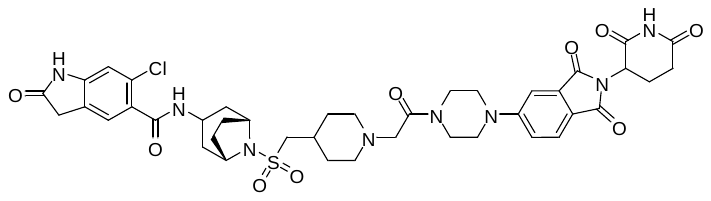

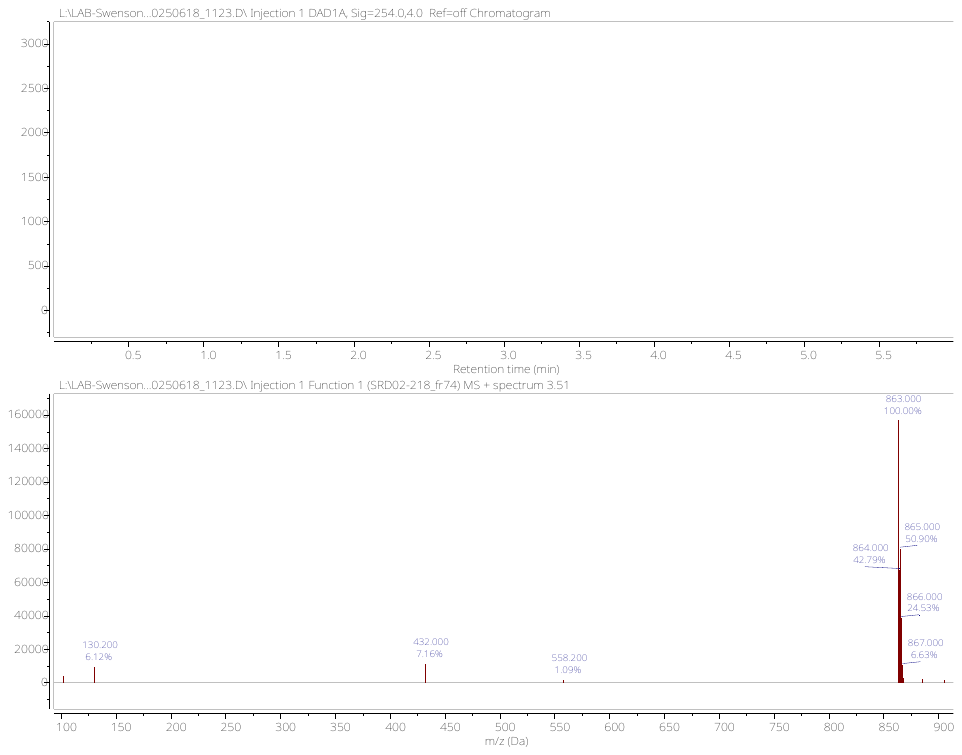

**^1^H NMR (500 MHz, MeOD-d4) spectrum of EPZ031686-VHL-01**

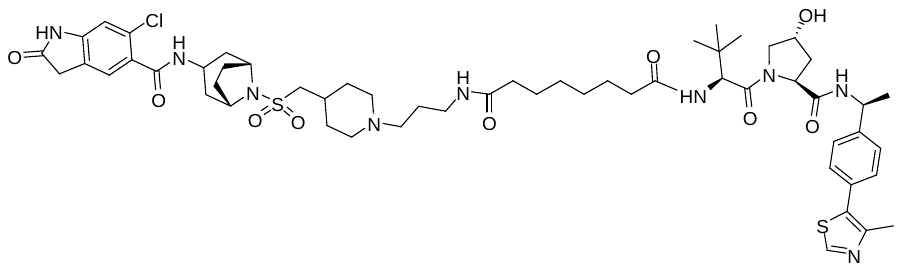

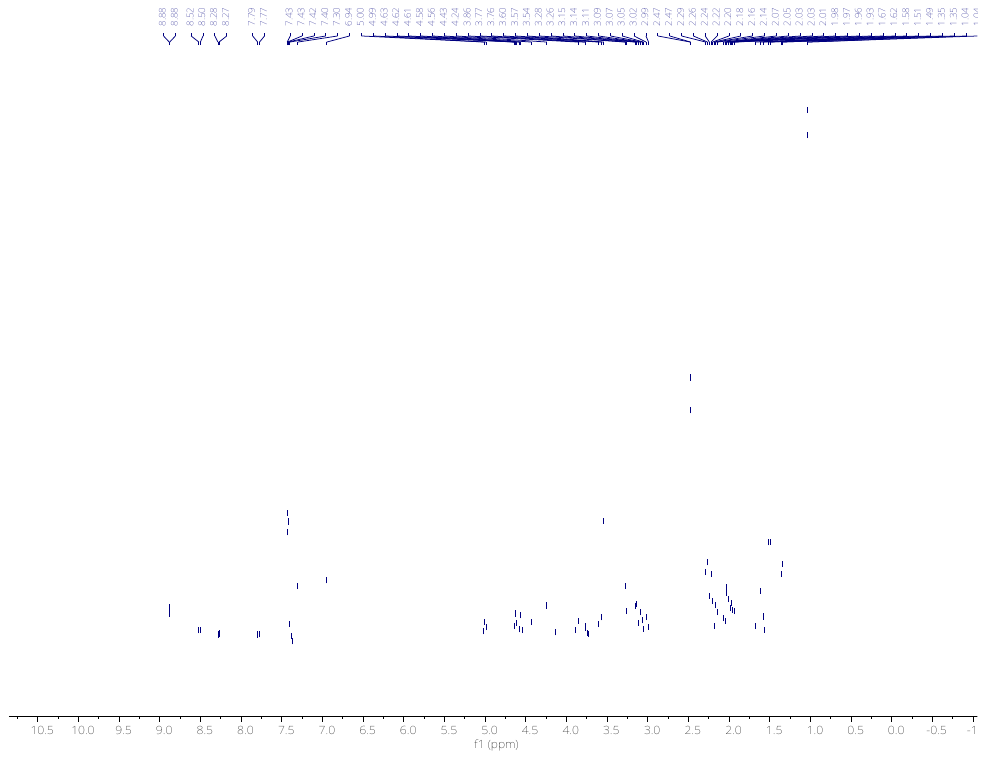

**UPLC-MS of EPZ031686-VHL-01**

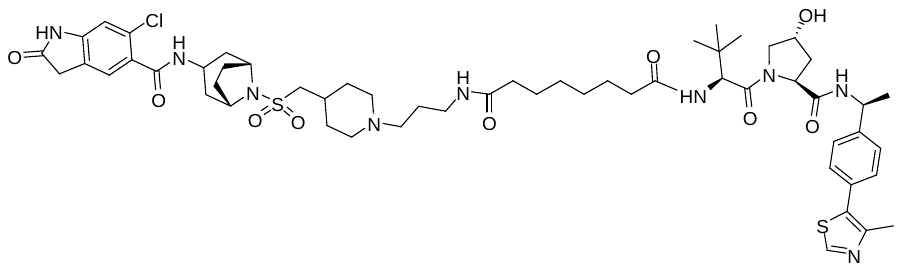

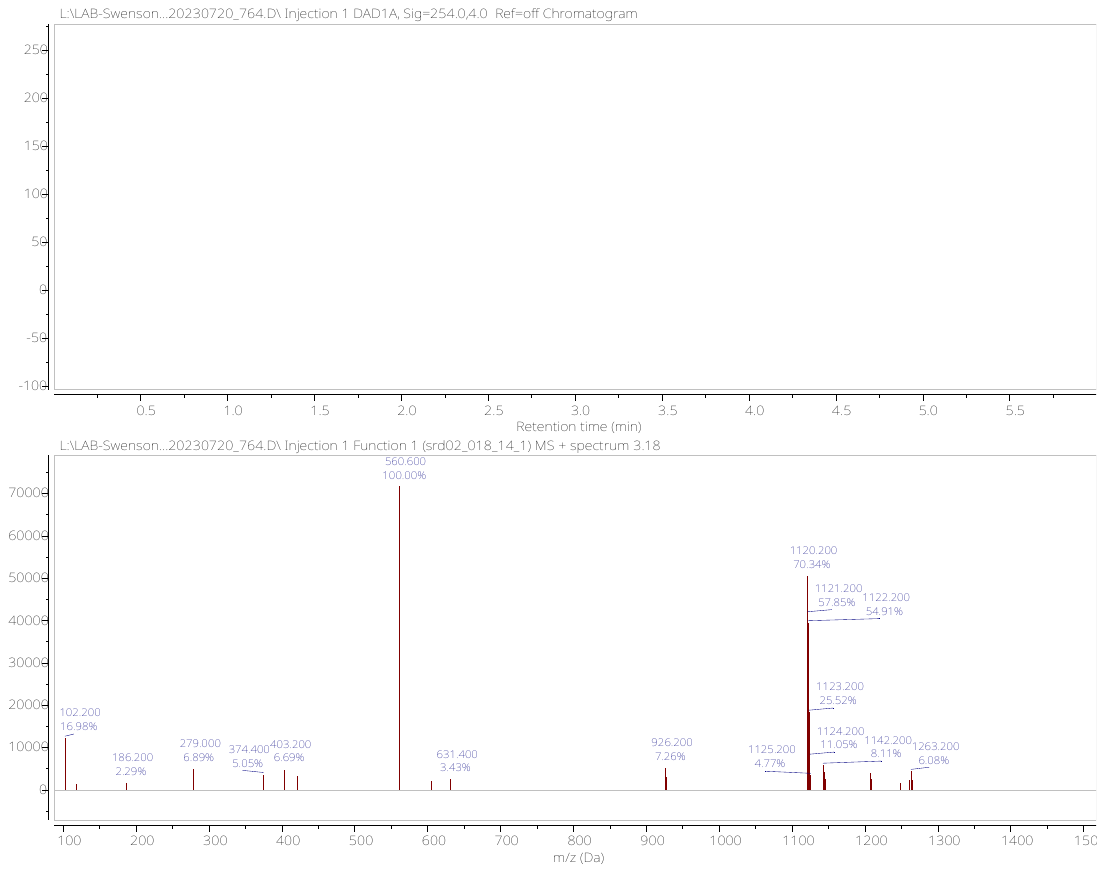

**^1^H NMR (500 MHz, MeOD-d4) spectrum of EPZ031686-VHL-02**

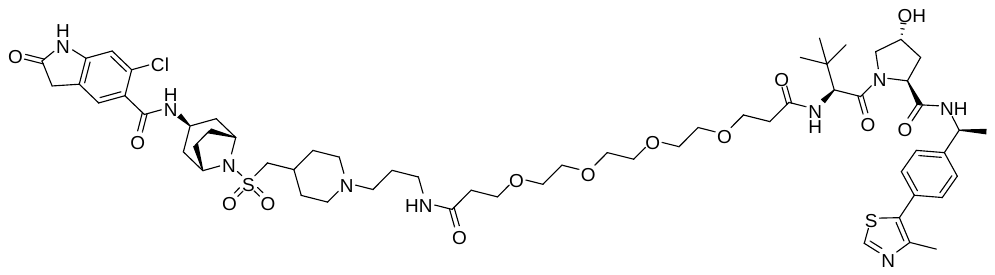

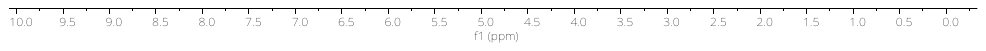

**UPLC-MS of EPZ031686-VHL-02**

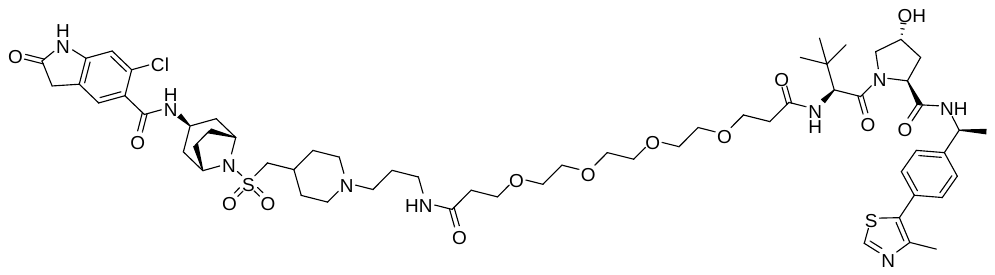

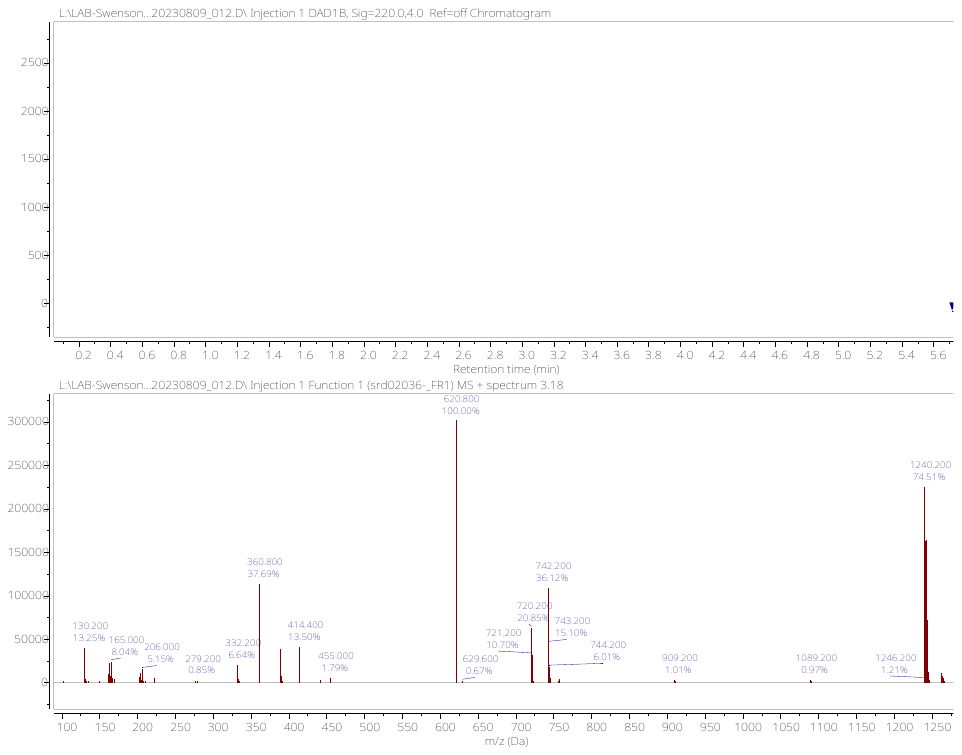

**^1^H NMR (500 MHz, MeOD-d4) spectrum of EPZ031686-VHL-03**

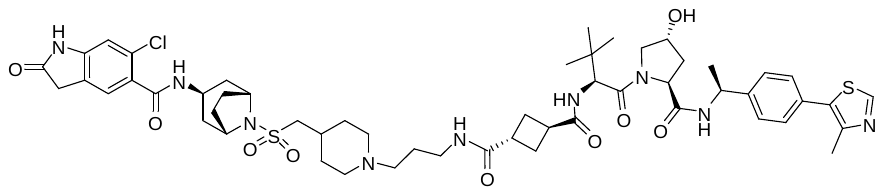

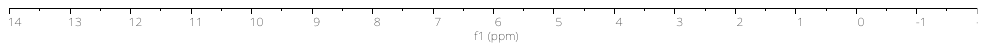

**UPLC-MS of EPZ031686-VHL-03**

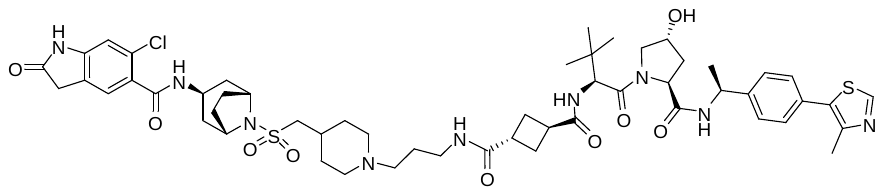

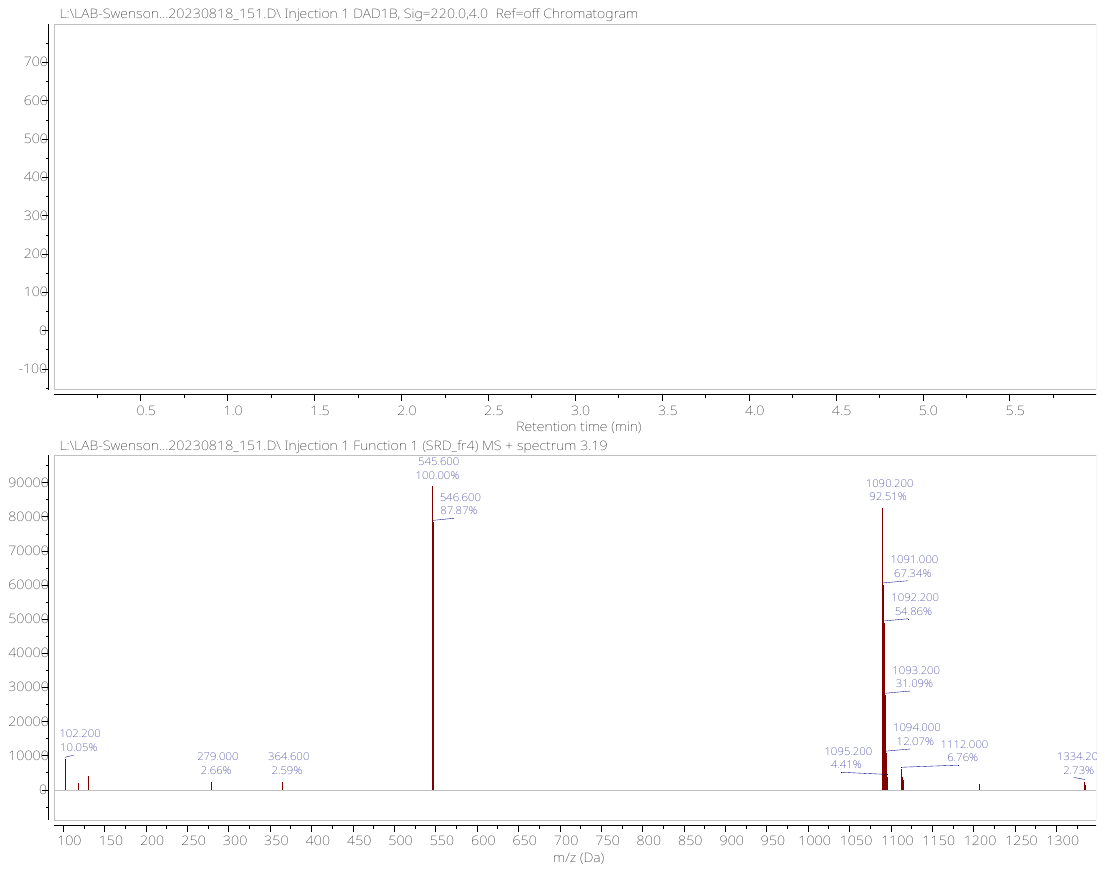

**^1^H NMR (500 MHz, MeOD-d4) spectrum of EPZ031686-VHL-04**

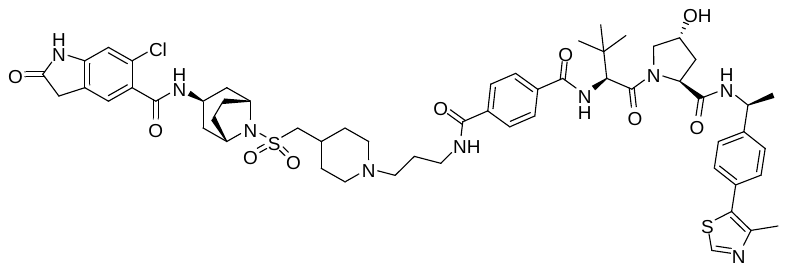

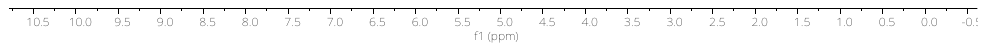

**UPLC-MS of EPZ031686-VHL-04**

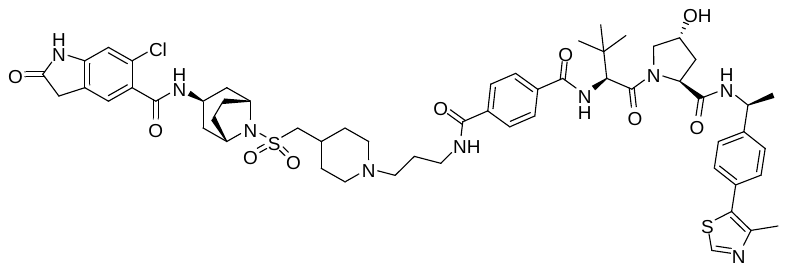

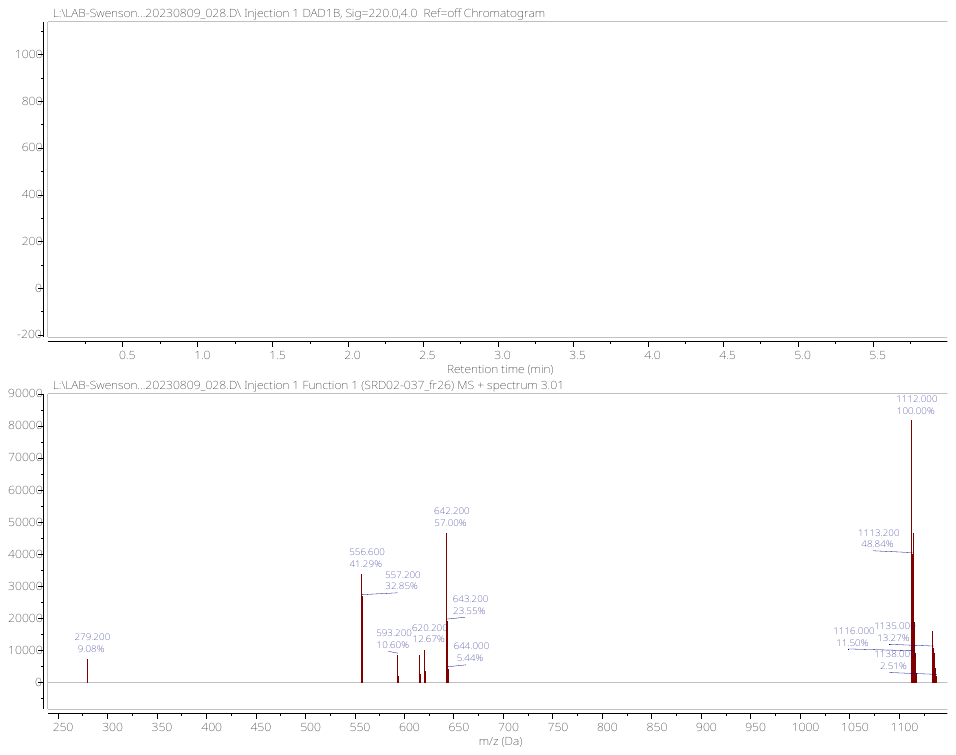

**^1^H NMR (500 MHz, DMSO-d6) spectrum of EPZ -VHL-05**

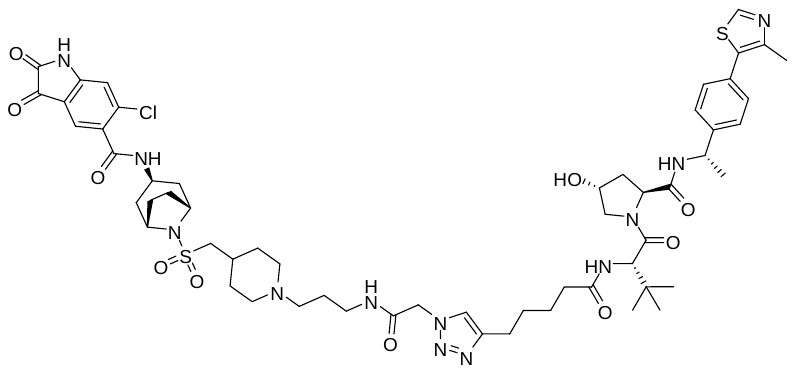

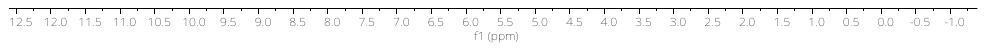

**UPLC-MS of EPZ031686-VHL-05**

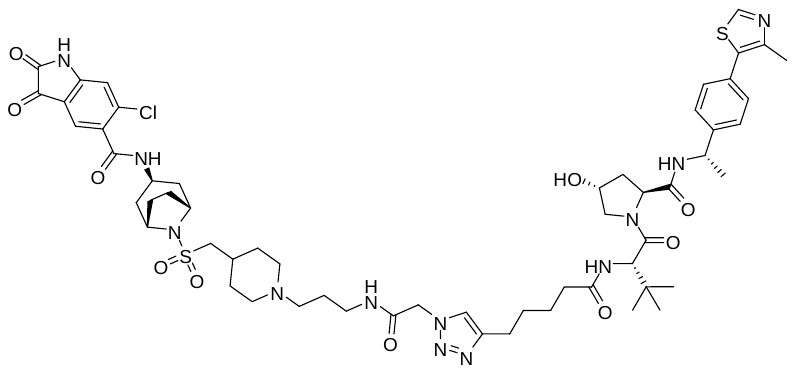

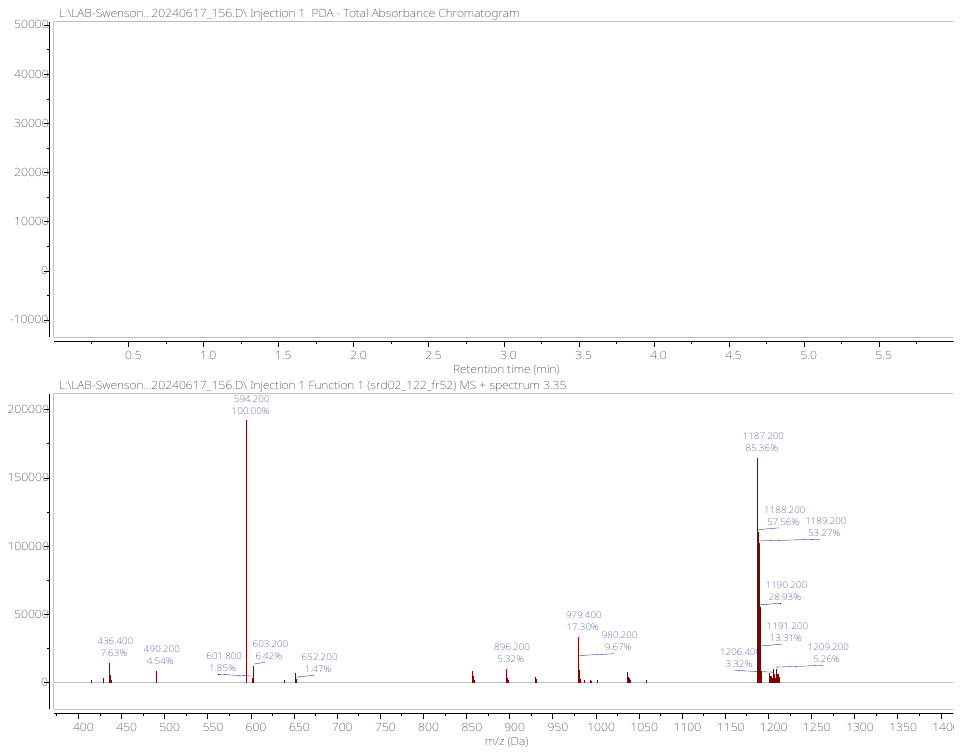
