## Supplementary Figure 11 to 13 for "SMYD3 protein degraders in HPV-negative head and neck squamous cell carcinoma"

**Supplementary Figure 11.** Dose-response curves obtained with HiBiT assays in HN13 cells stably expressing doxycycline-inducible HiBiT-tagged SMYD3. Cells were seeded in 6-well plates, and induction with doxycycline (1ug/ml) was initiated. Next day, the cells were seeded into 96-well plates in triplicates (10,000 cells/well) and 24h later, treatment with EPZ031686-IAP-08 was initiated at a range of concentrations from 0-5uM. Bioluminescence was obtained at 4h and 24h after treatment initiation.

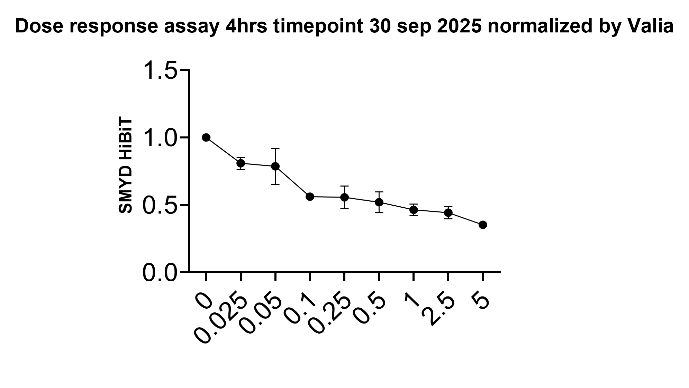

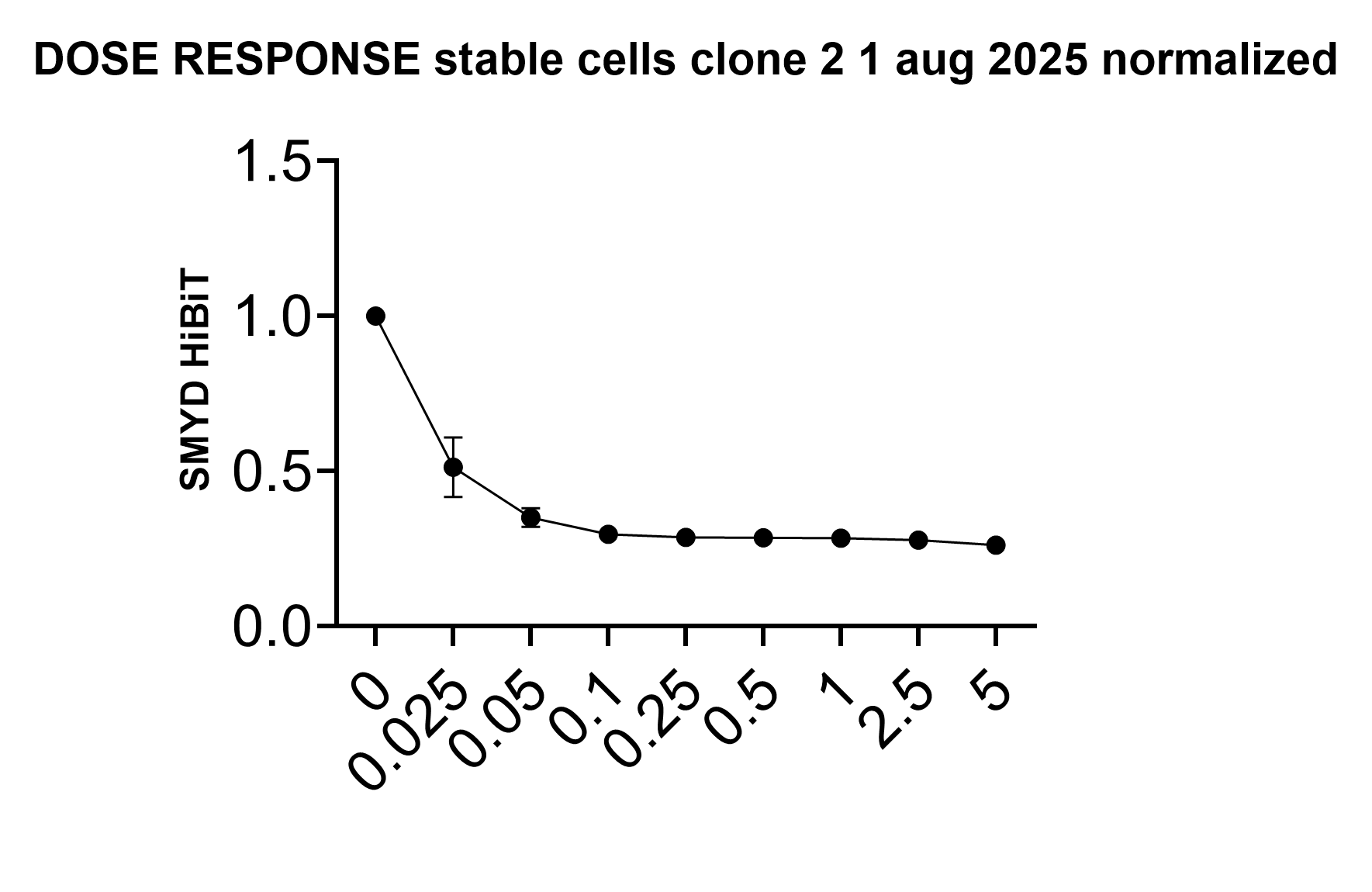

Concentration (uM)

Relative Bioluminescence

4h timepoint

24h timepoint

IAP-08

**Supplementary Fig. 12.** NanoBRET target engagement cellular SMYD3 assay with the SMYD3 inhibitor EPZ031686. The assay was performed in HEK293 cells. Standard deviations of biological duplicates are shown for each concentration.

Normalized BRET Response (%)

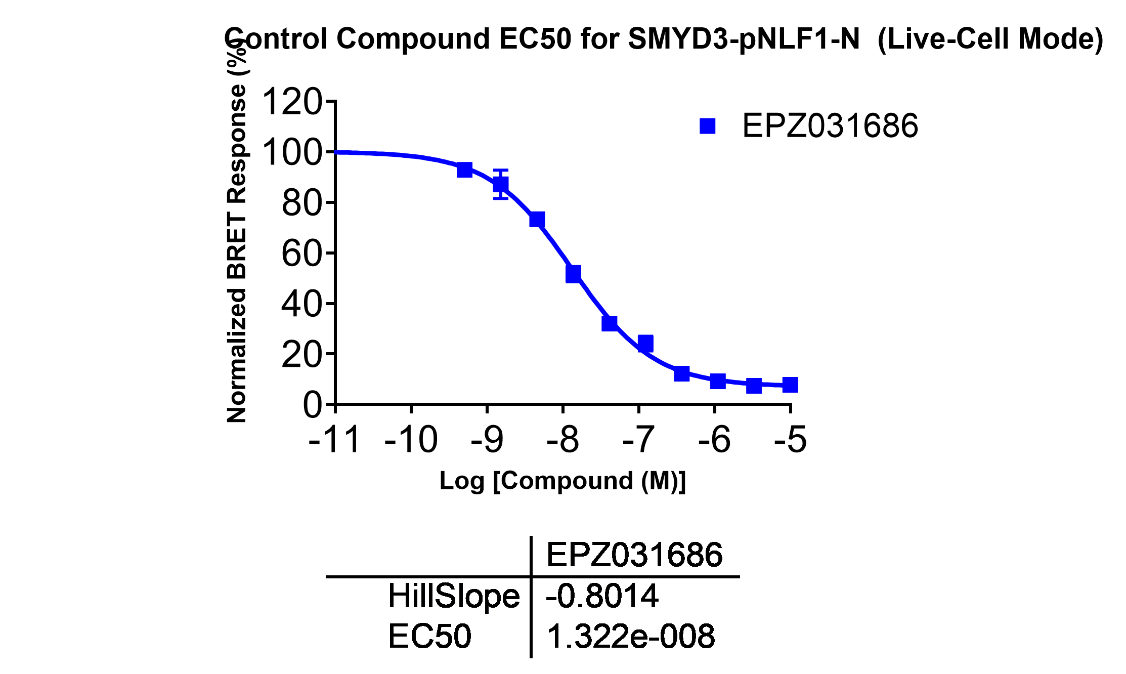

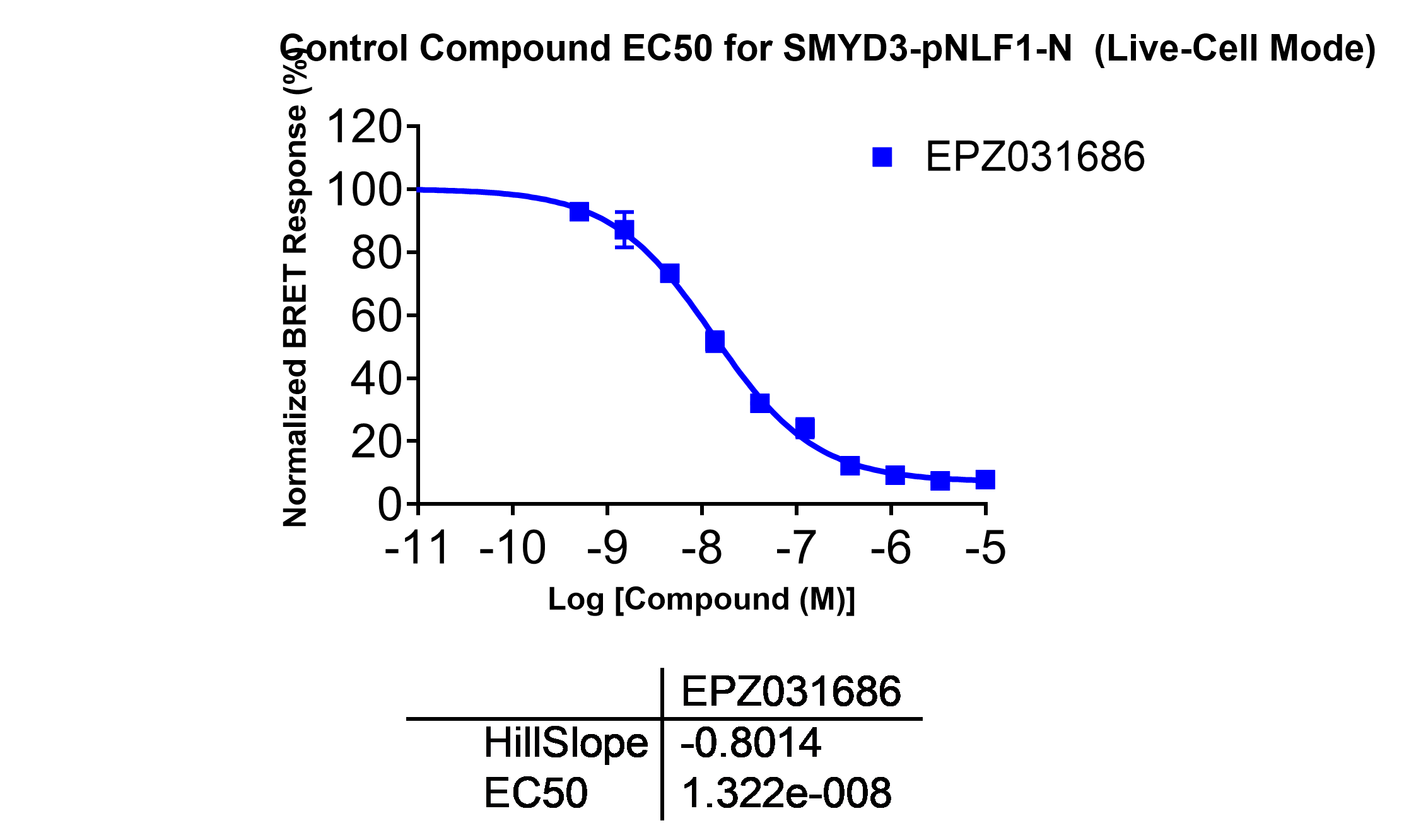

Log [Concentration (M)]

**Supplementary Figure 13.** CFAs and CCK8 assays comparing the in vitro efficacy of the XIAP inhibitor A 410099.1 and IAP-08. **(A)** CFA assays. HN-SCC-151 and HN13 cells were seeded in triplicates in 6-well plates (500 cels/well) and treated with incremental concentrations of the XIAP inhibitor A 410099.1, IAP-08 or DMSO (control). Left: images of representative wells per condition. Right: quantification graphs representing the average number of colonies per condition. **(B)** CCK8 assays. HNSCC cells were plated in 96-well plates in biological quadruples and next day, treatment was initiated with IAP-08 at the concentrations shown versus DMSO (control). Relative absorptions were assessed on the given time points. SDs are shown. Colonies were quantified using Image J. Student t-test; *p<0.05, **p<0.01, ***p<0.005, ****p<0.001.

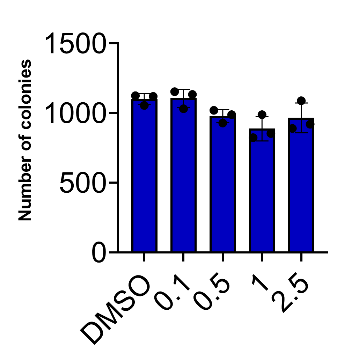

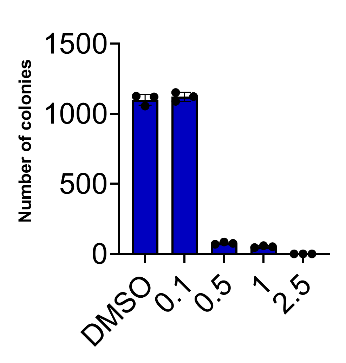

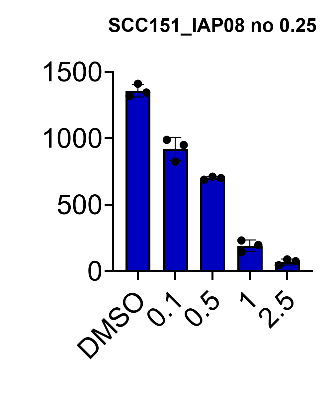

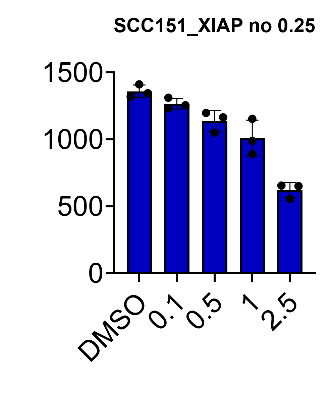
**(A)**

**HN13**

****

****

****

***

****

****

****

****

*

Average Number of Colonies

IAP-08

XIAP antagonist

Average Number of Colonies

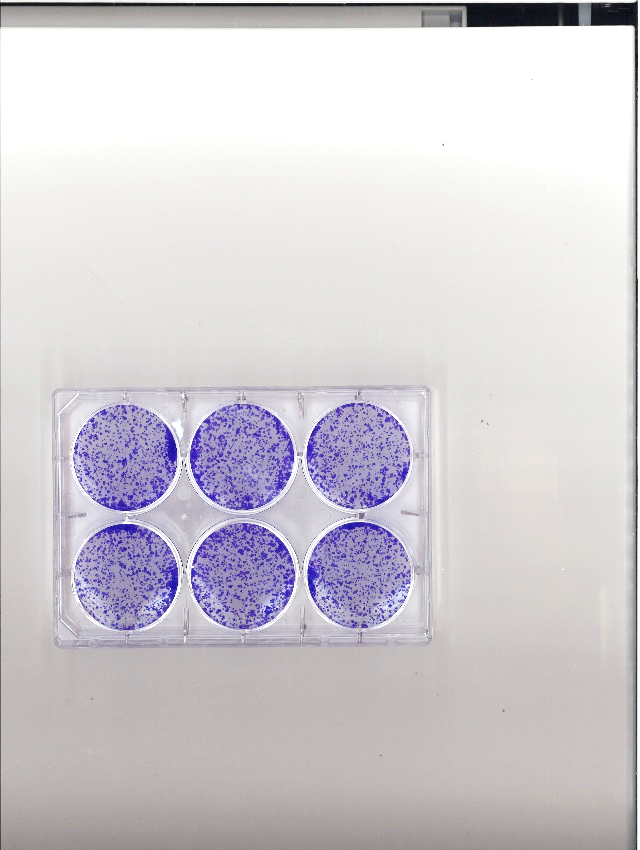

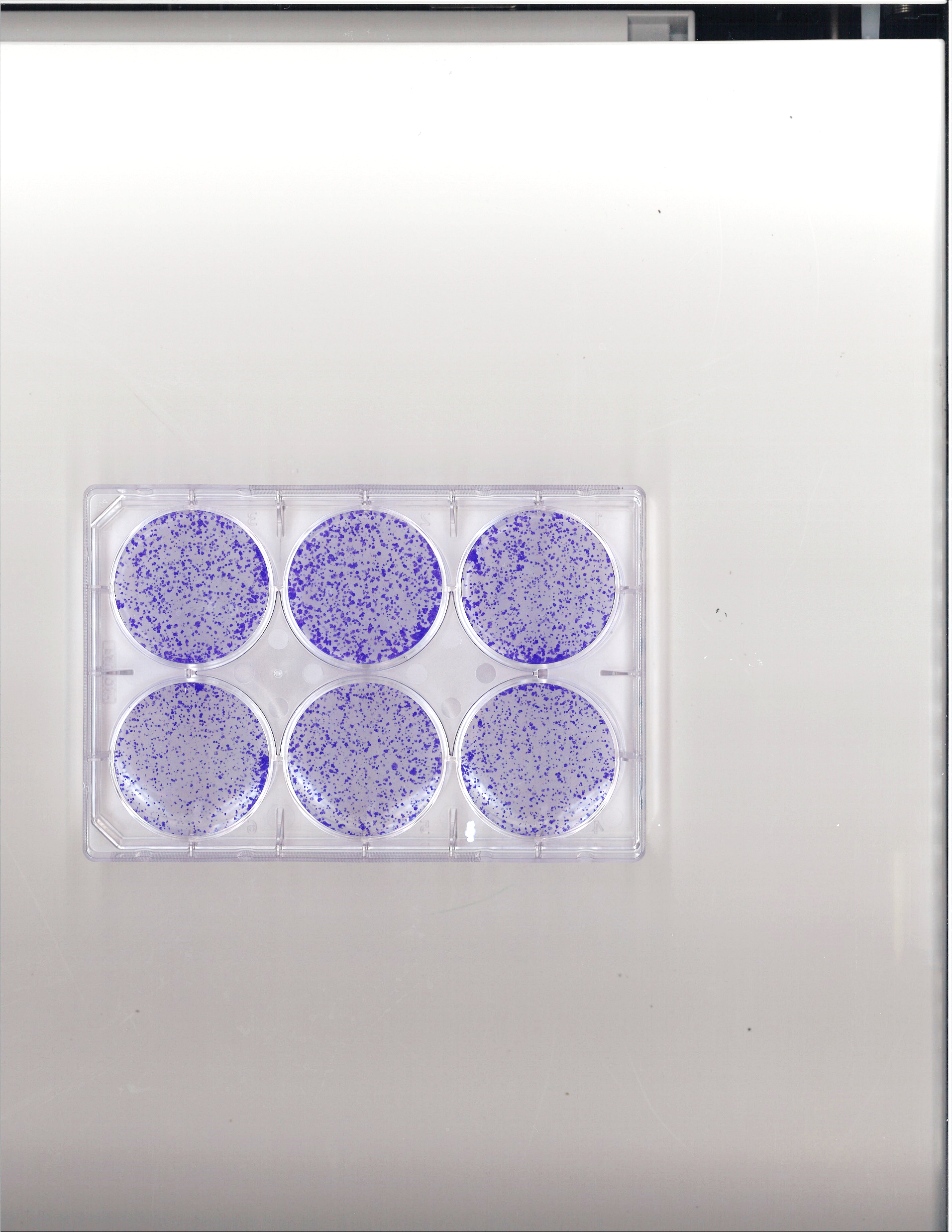

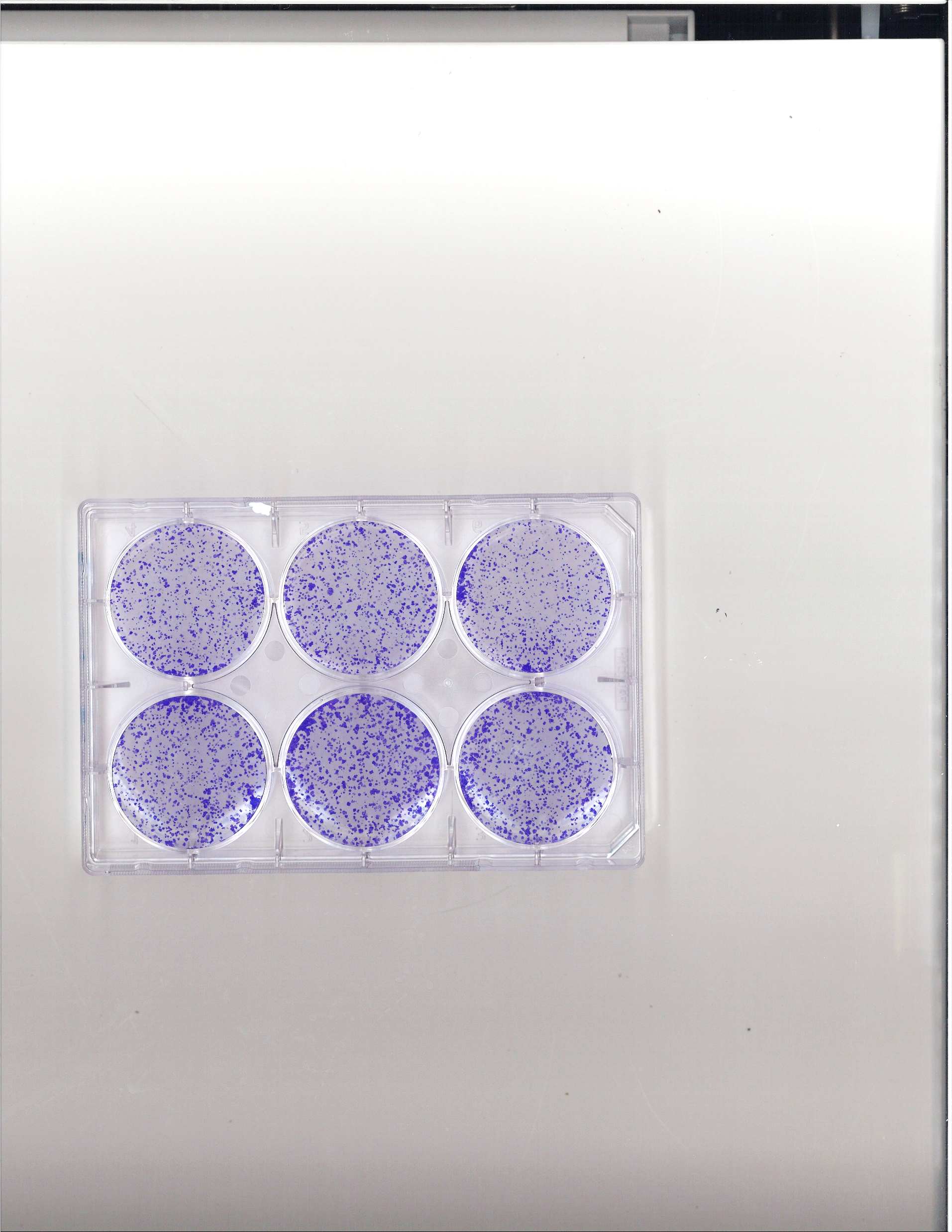

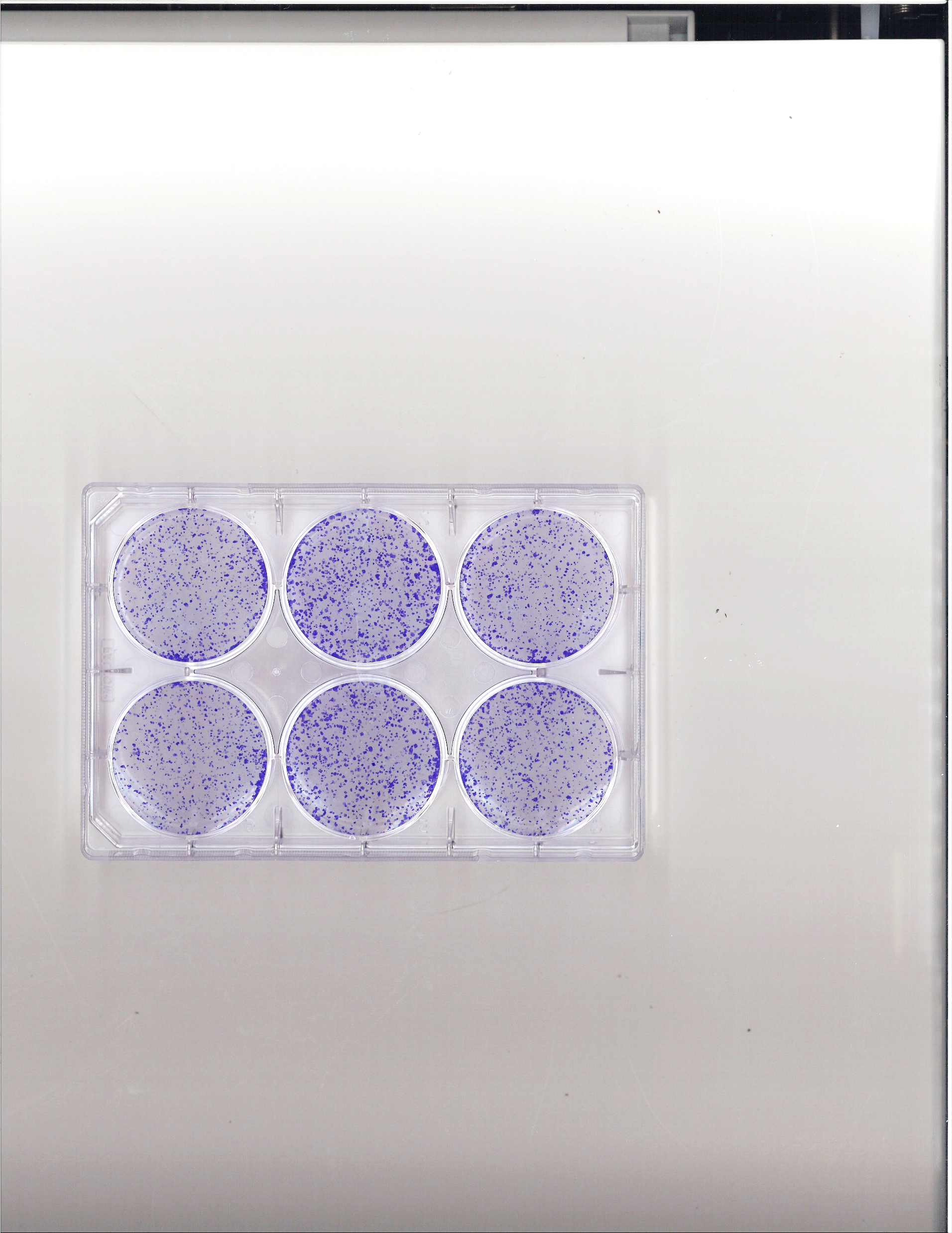

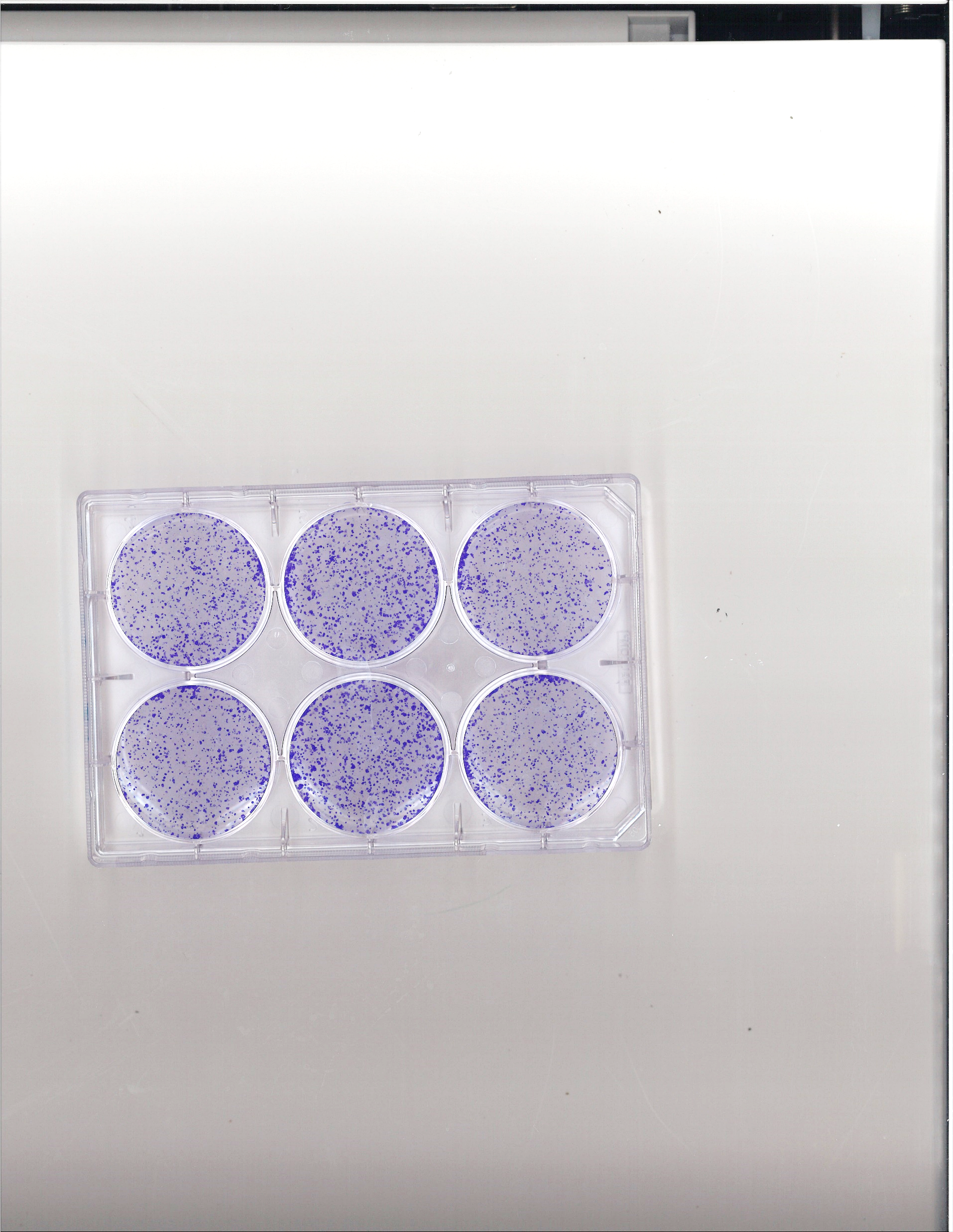

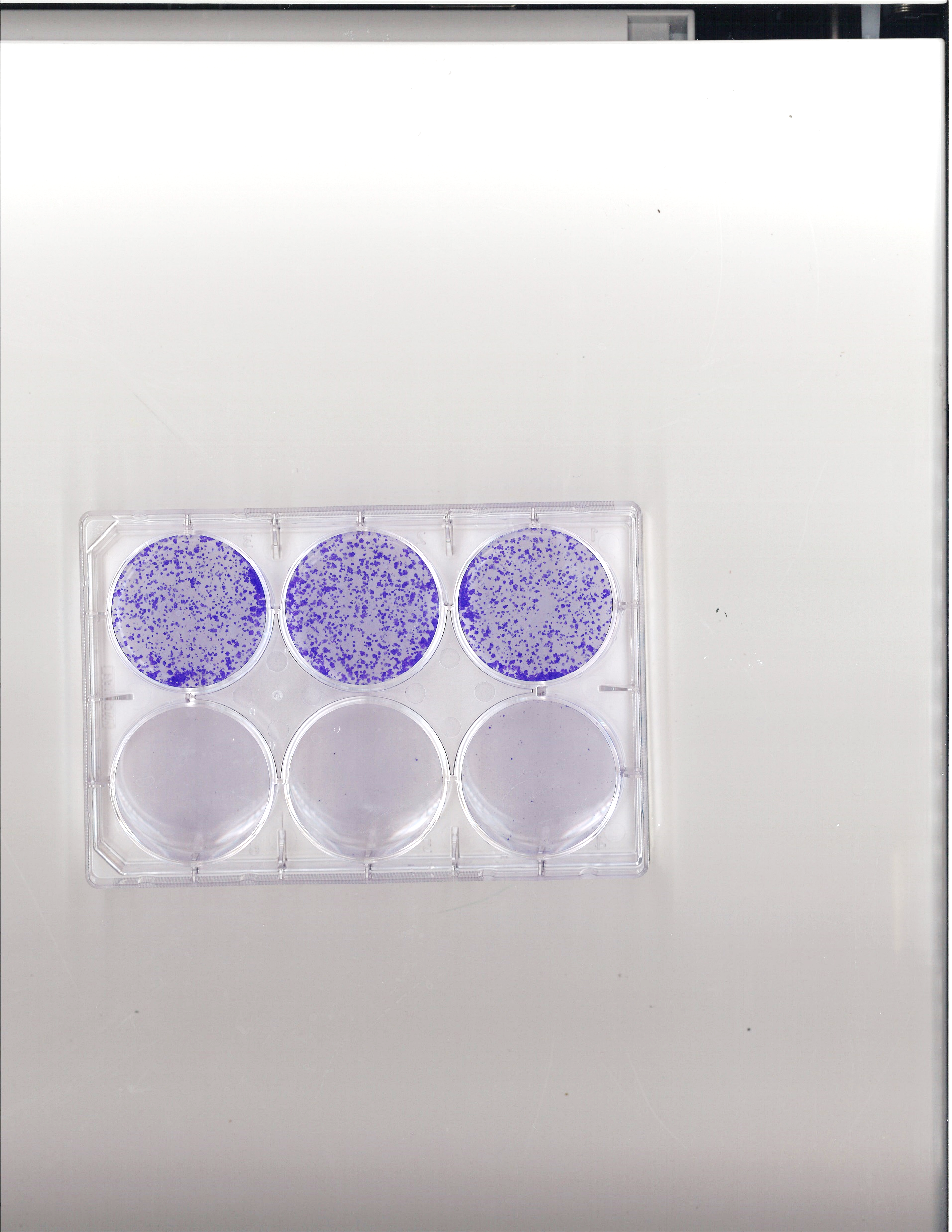

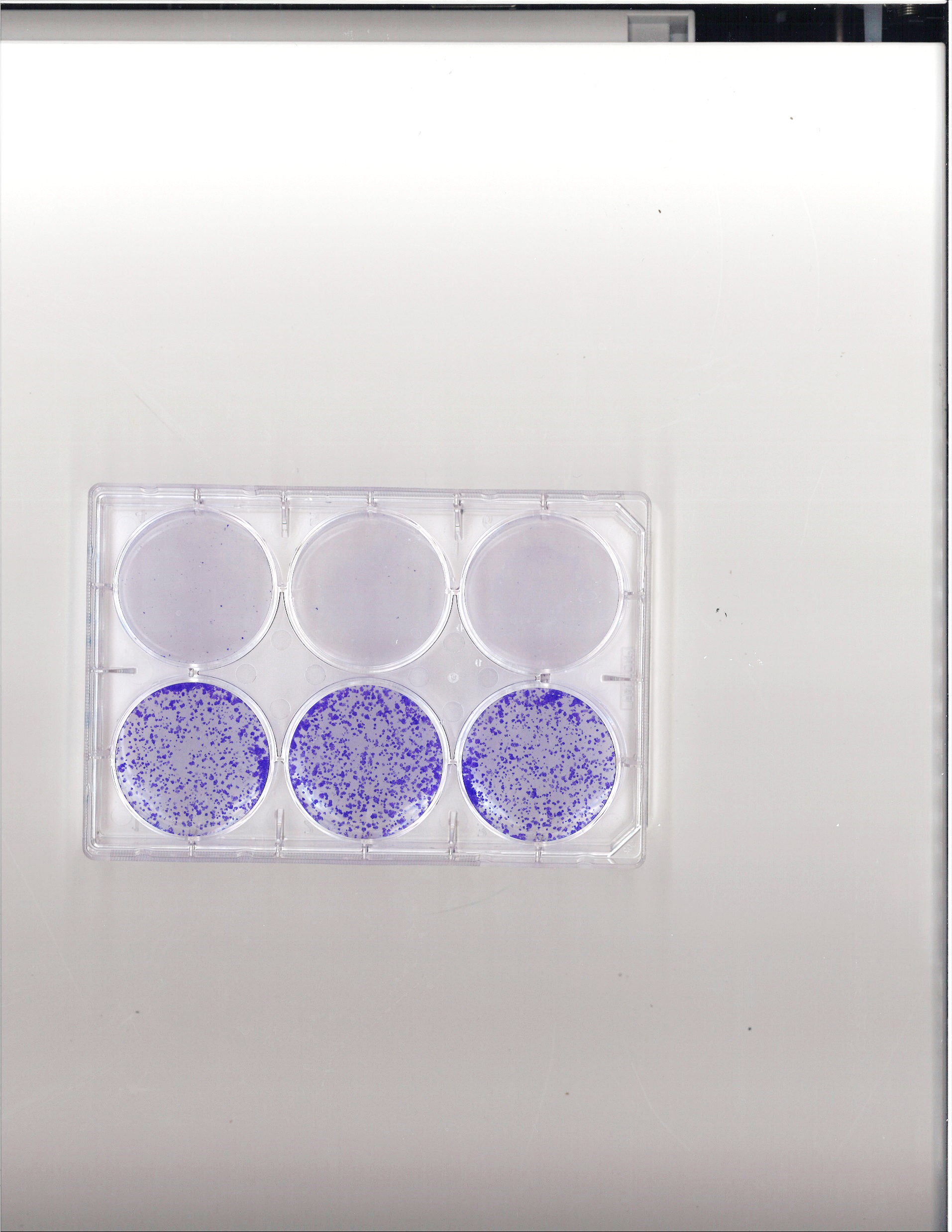

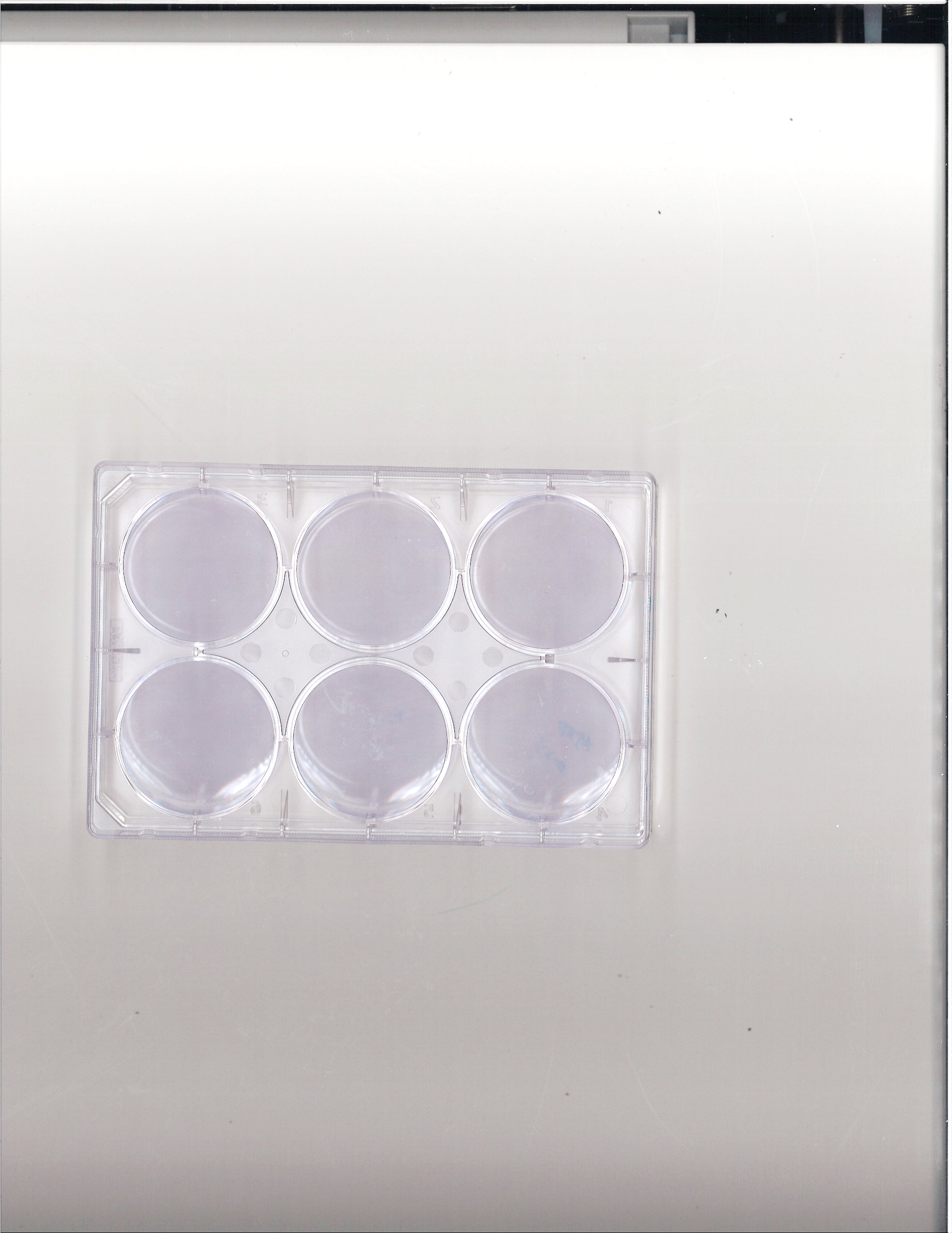

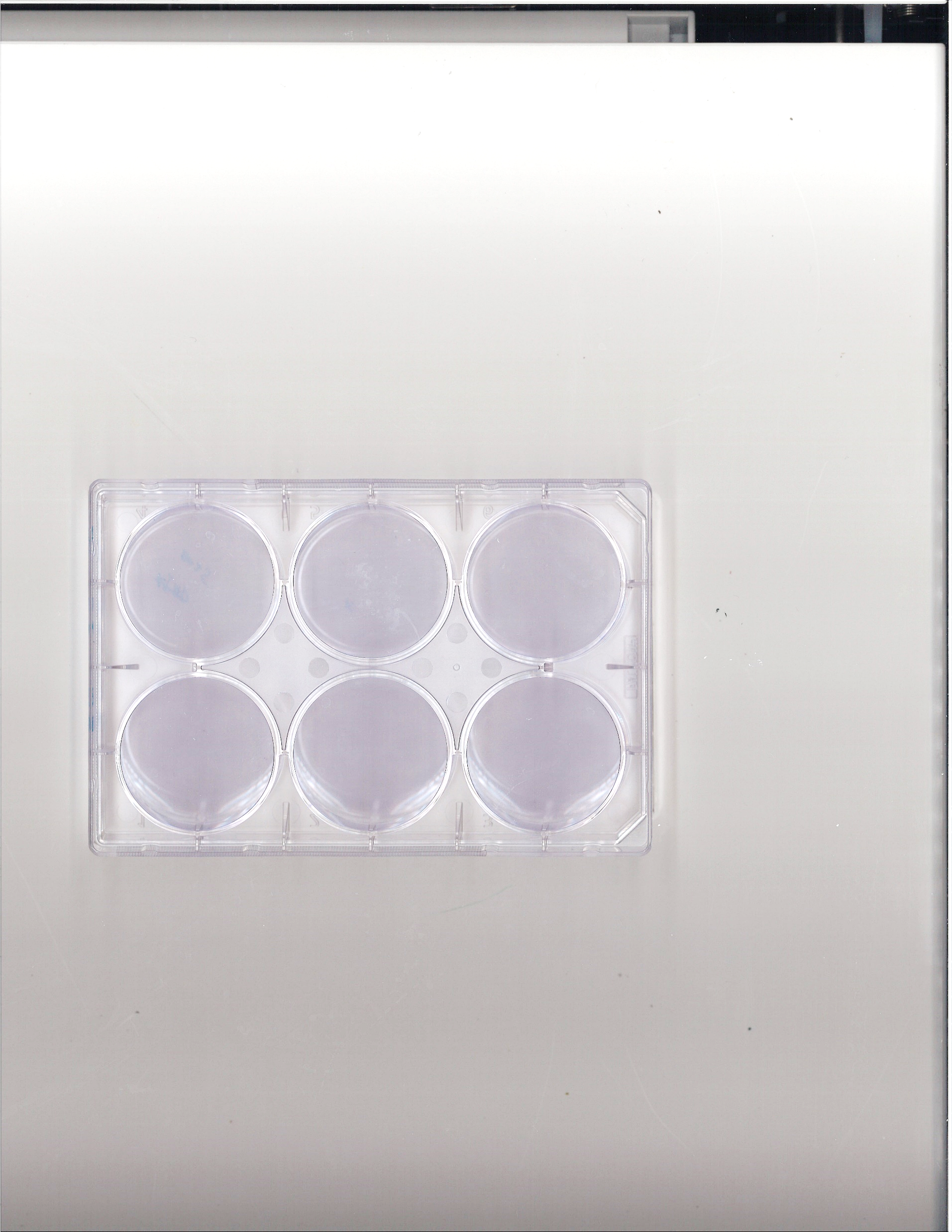

IAP-08

XIAP antagonist

**HN13**

2.5

1

0.5

0.1

DMSO

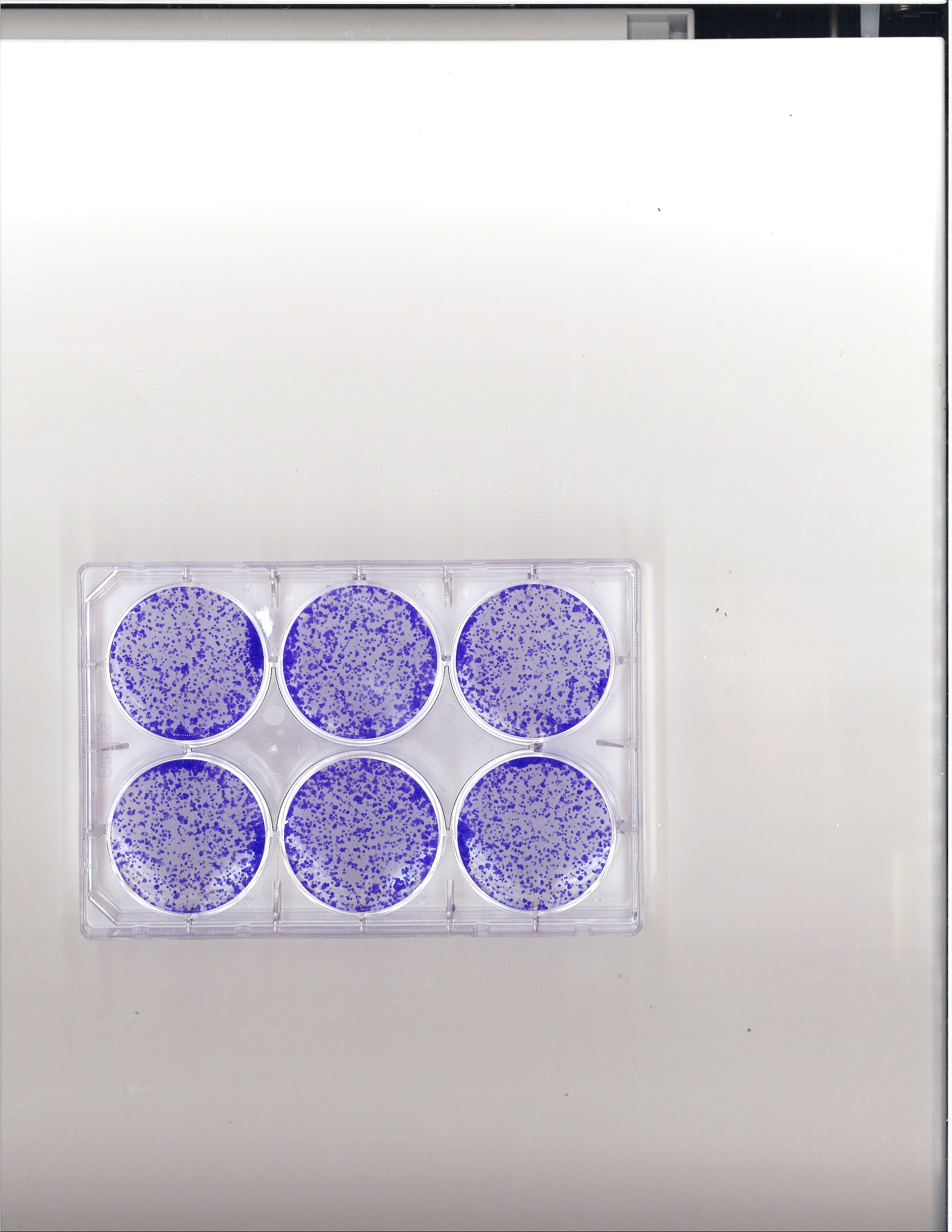

uM

uM

**HN-SCC-151**

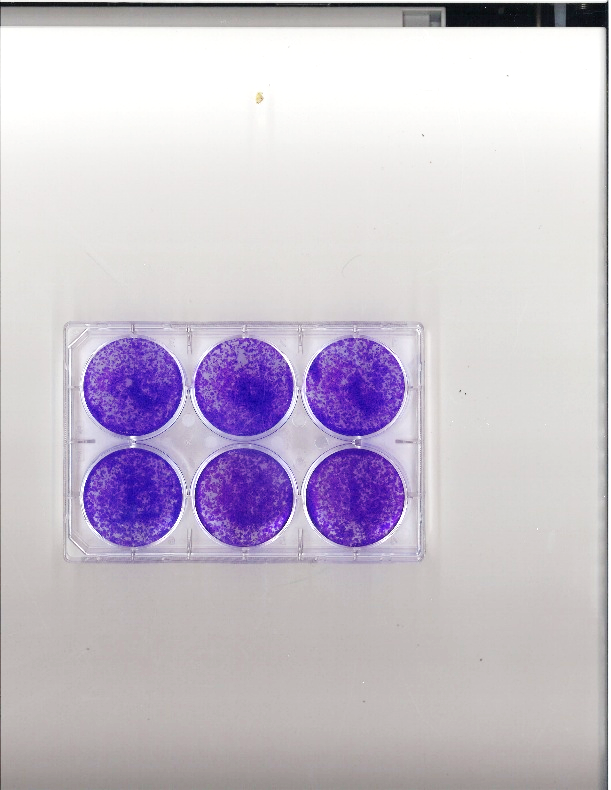

2.5

1

0.5

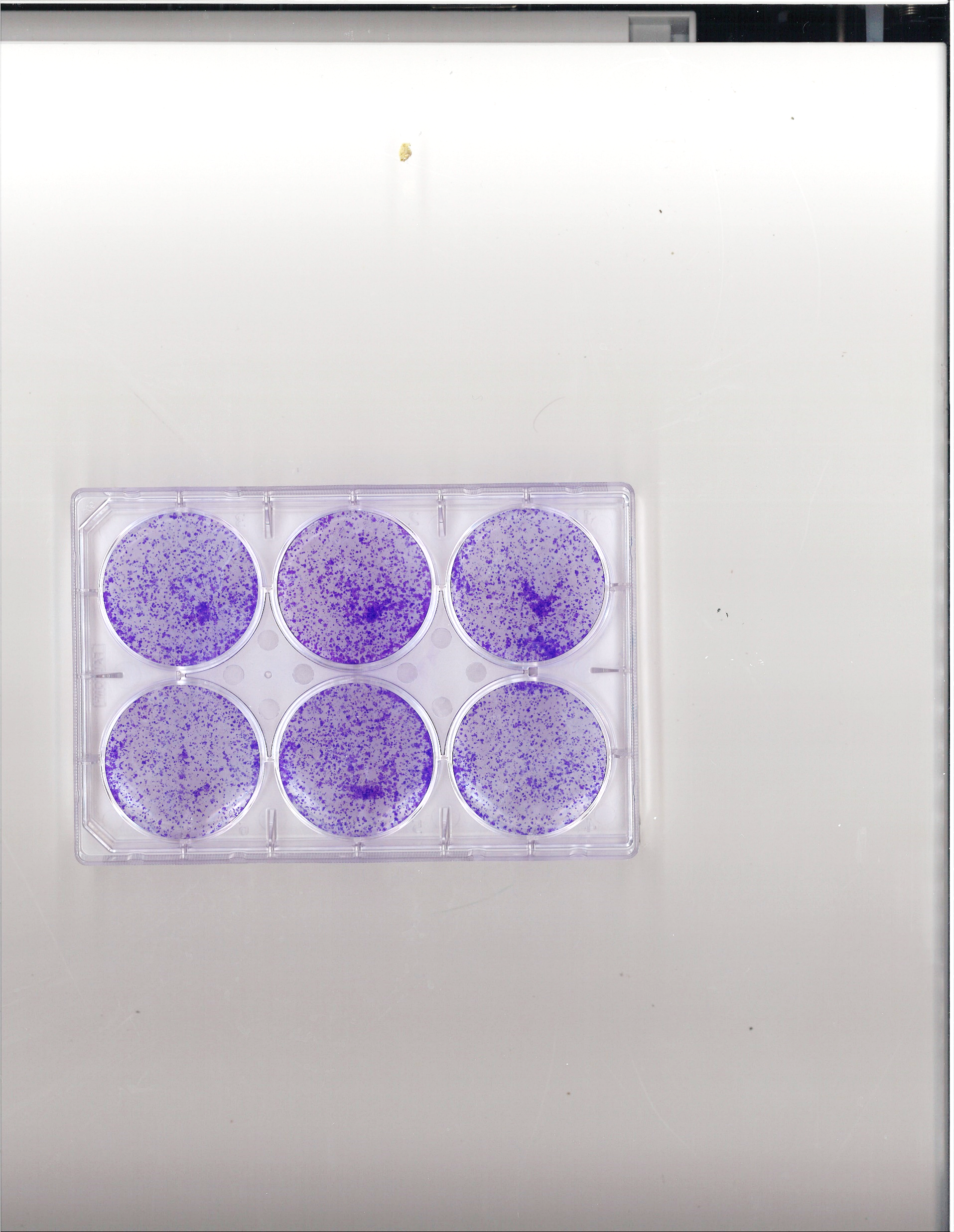

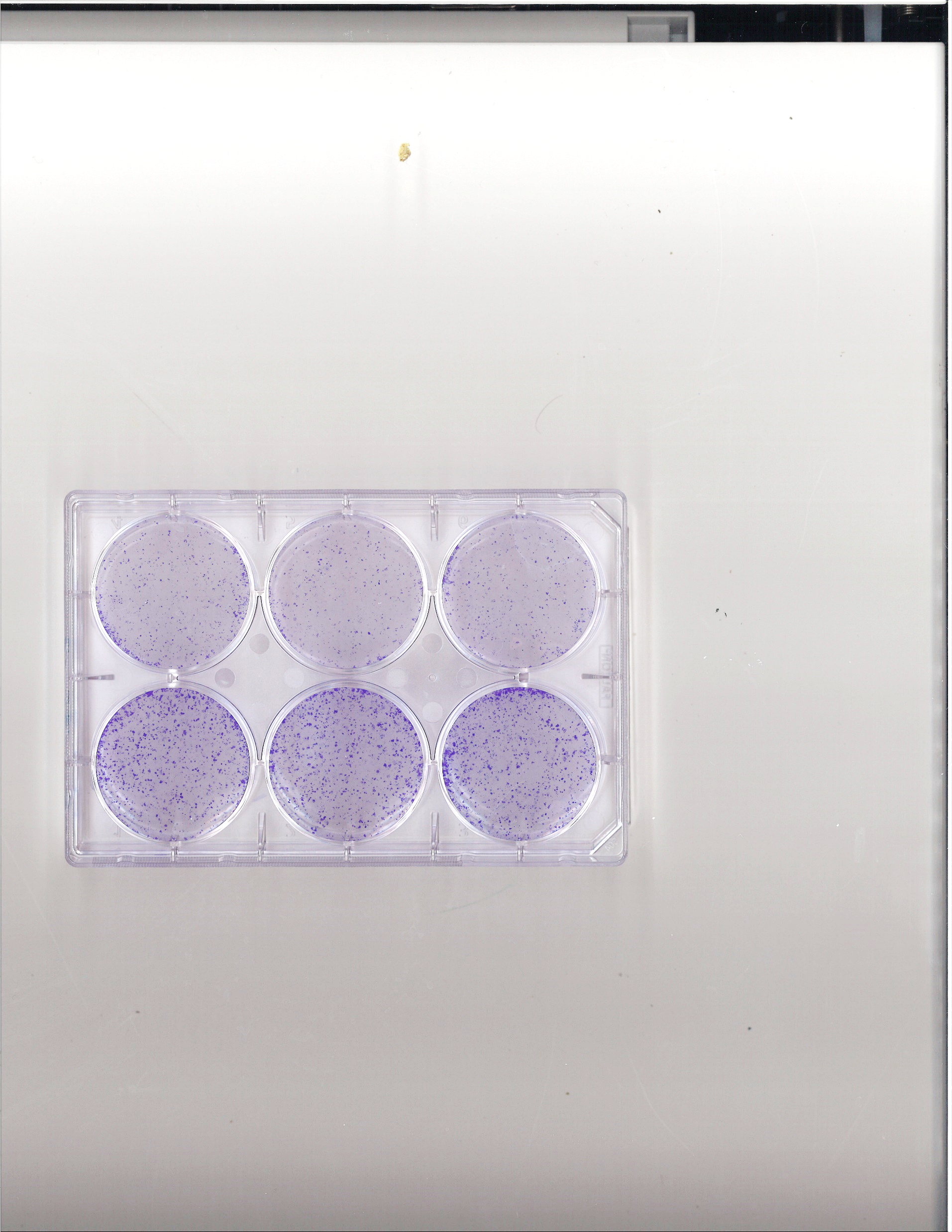

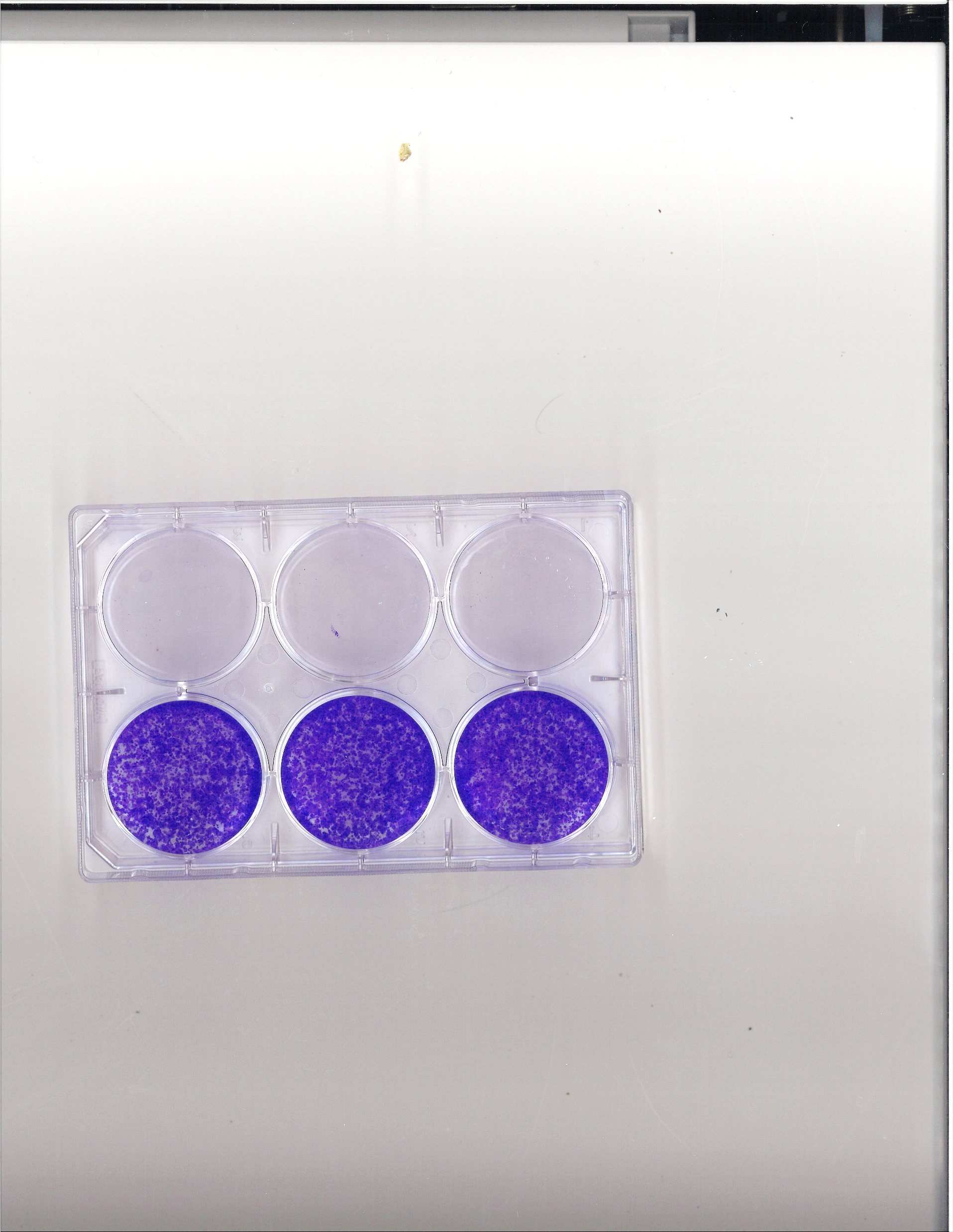

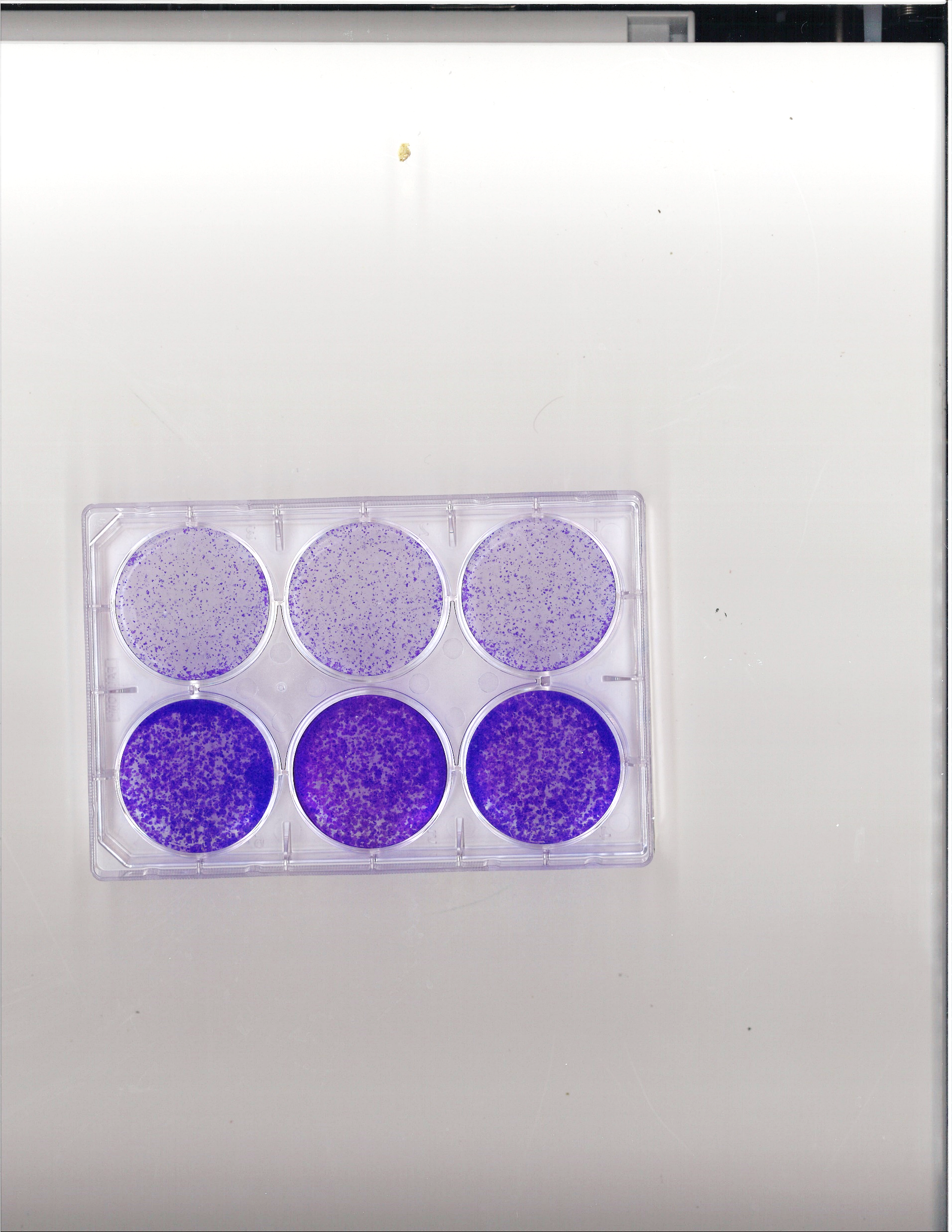

0.1

DMSO

IAP-08

XIAP antagonist

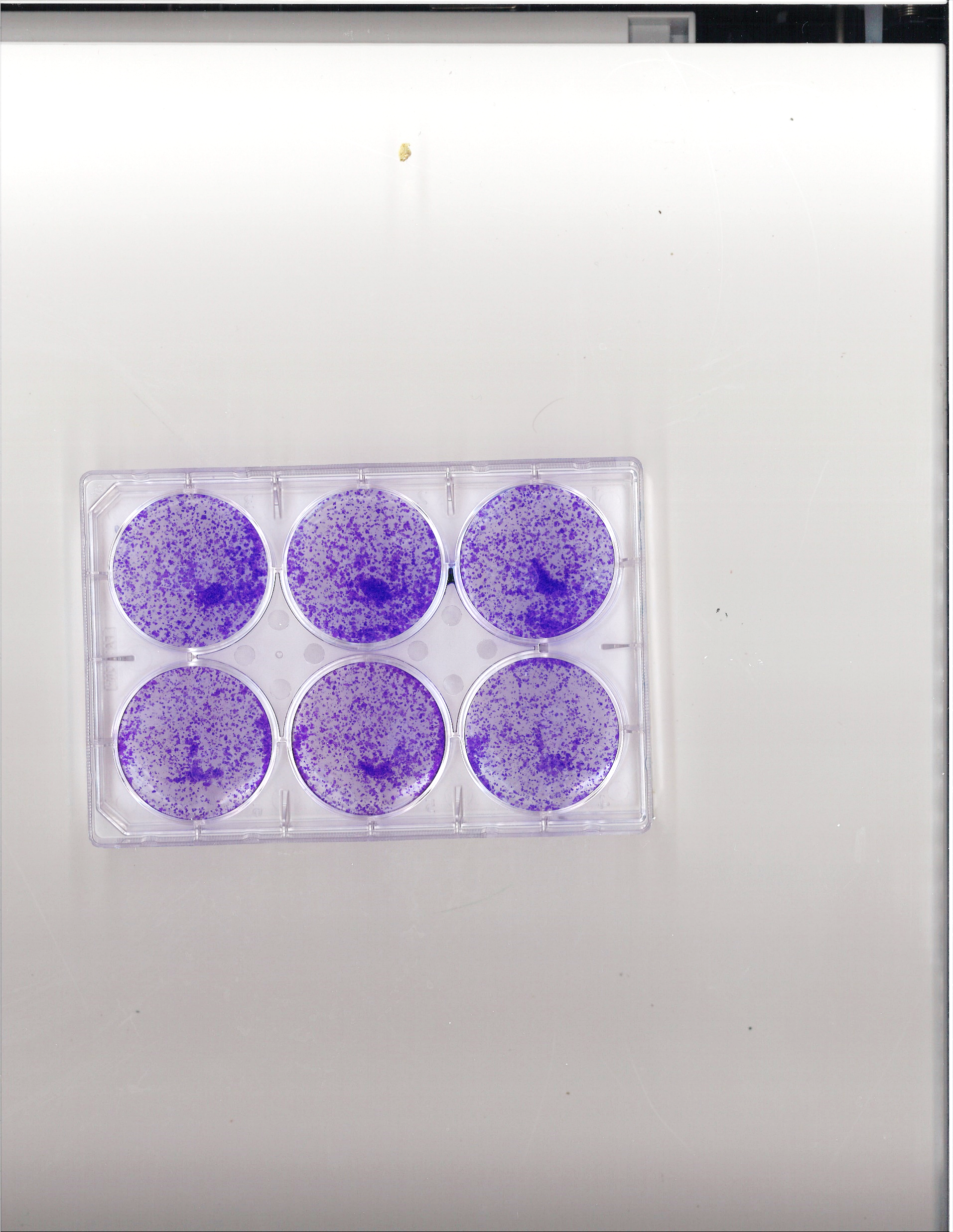

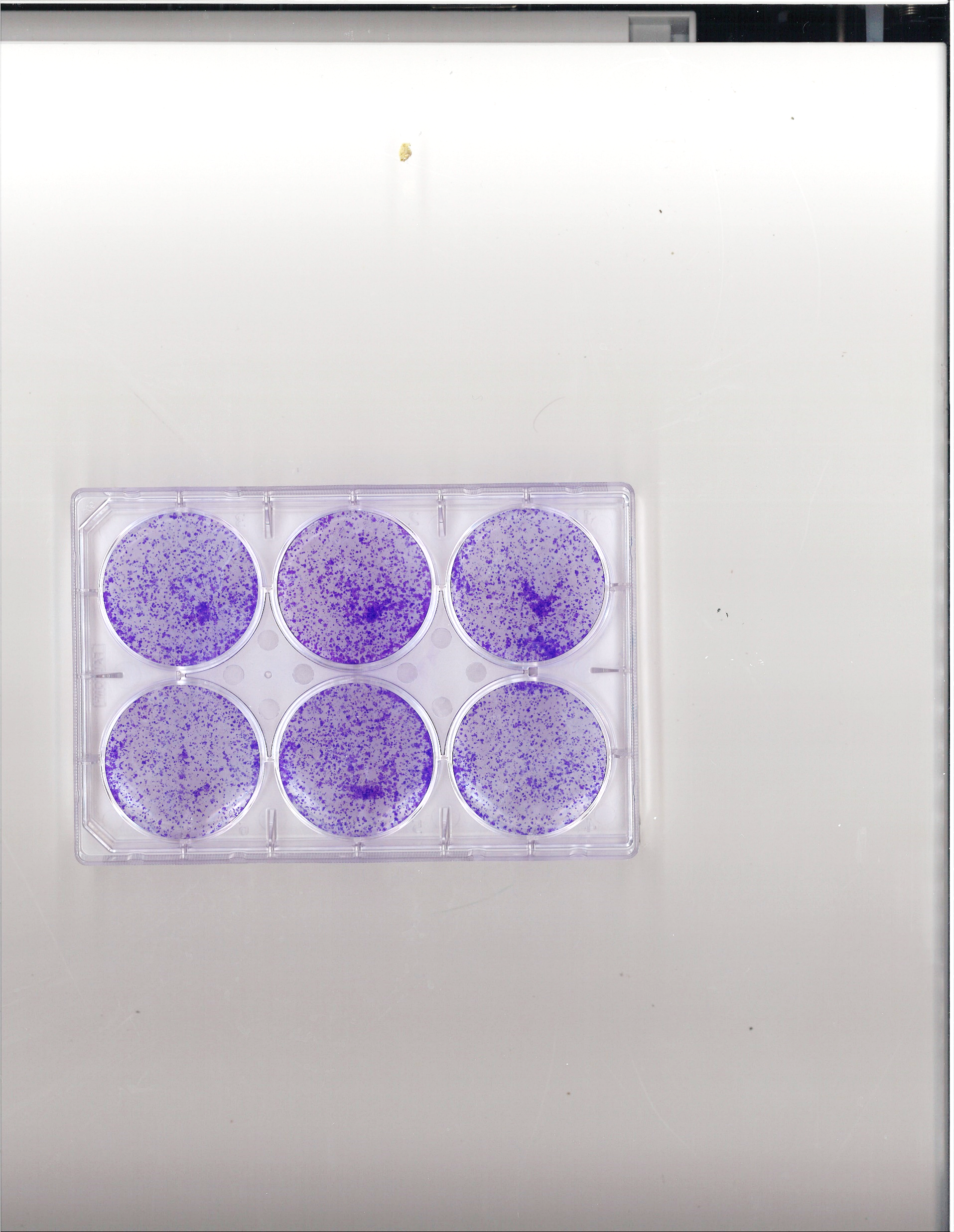

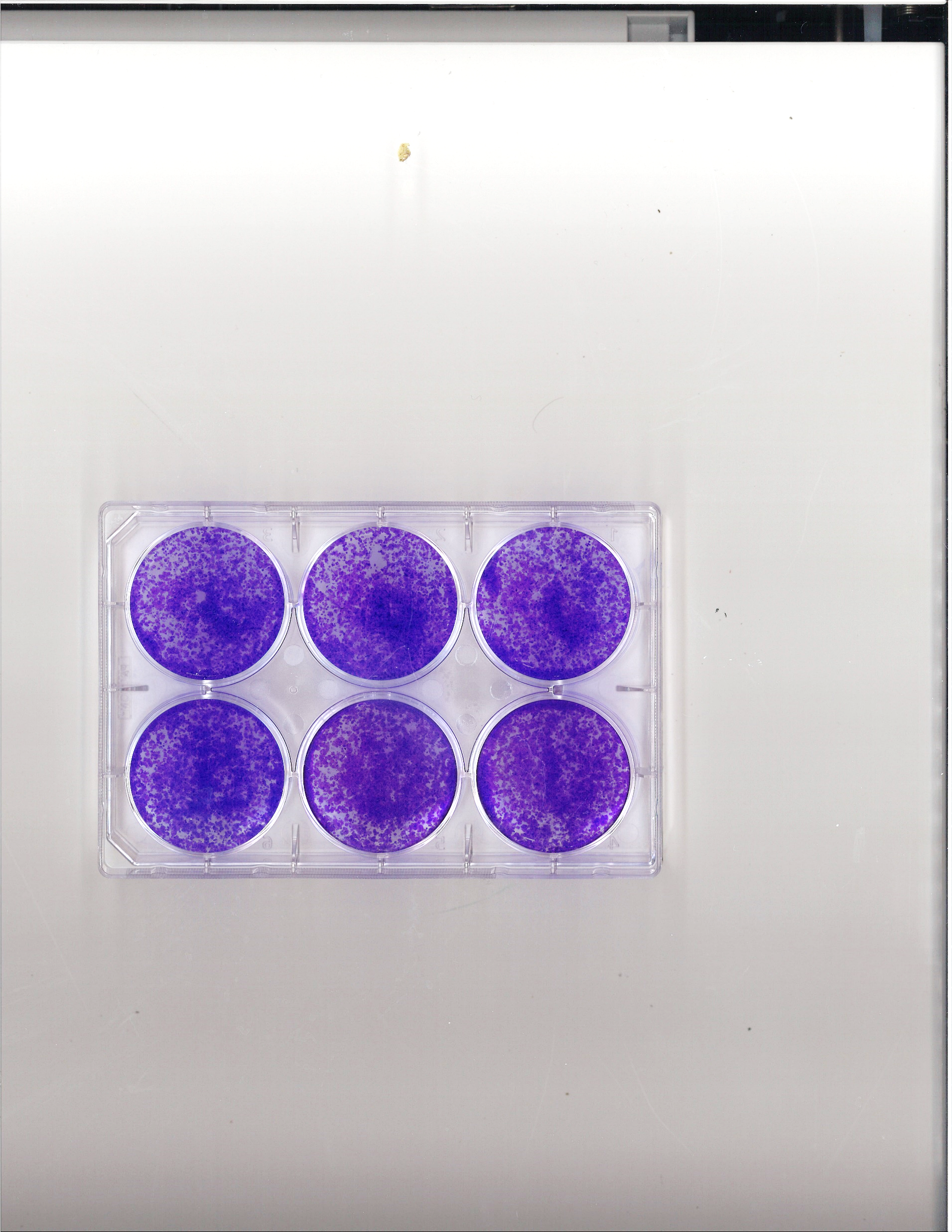

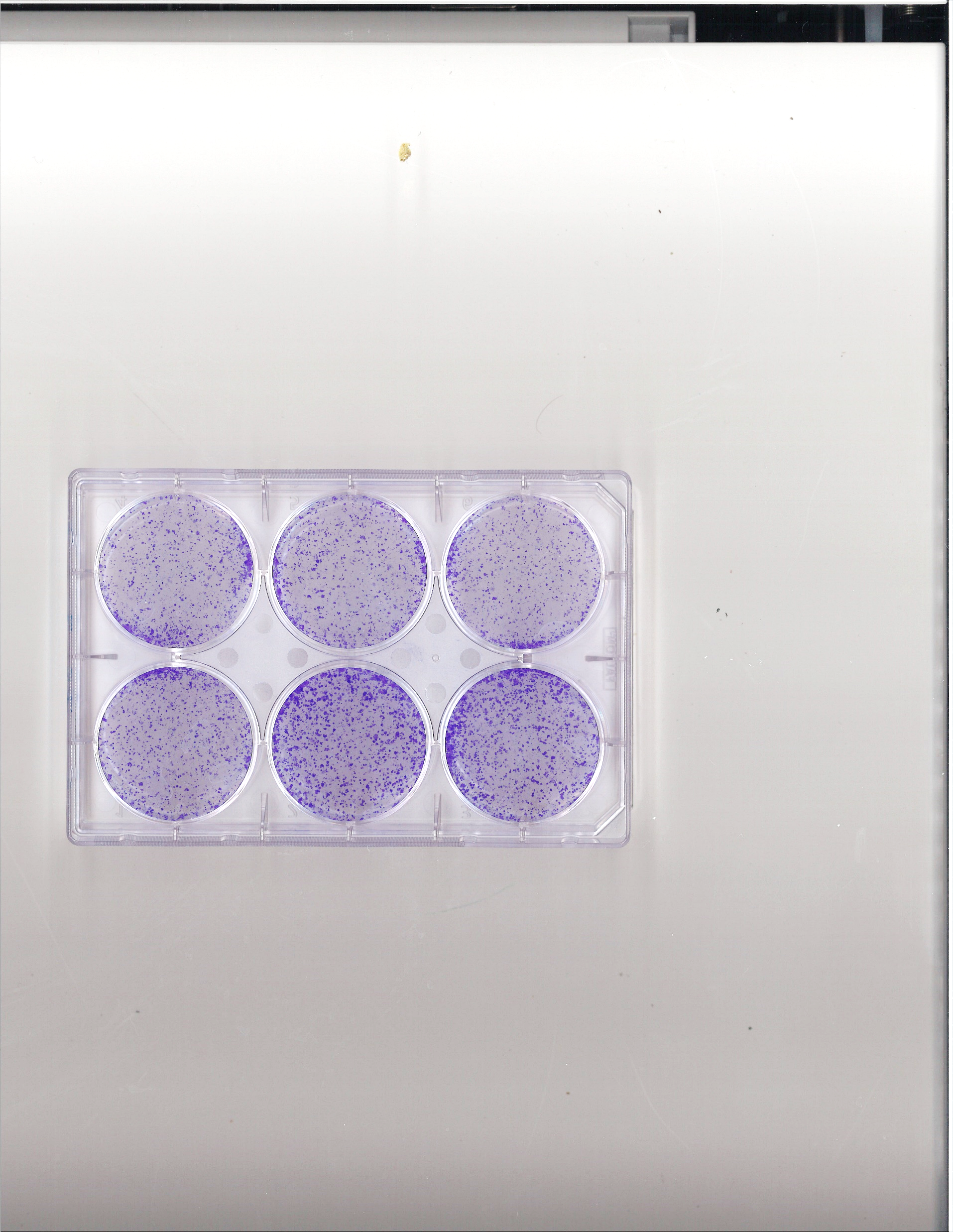

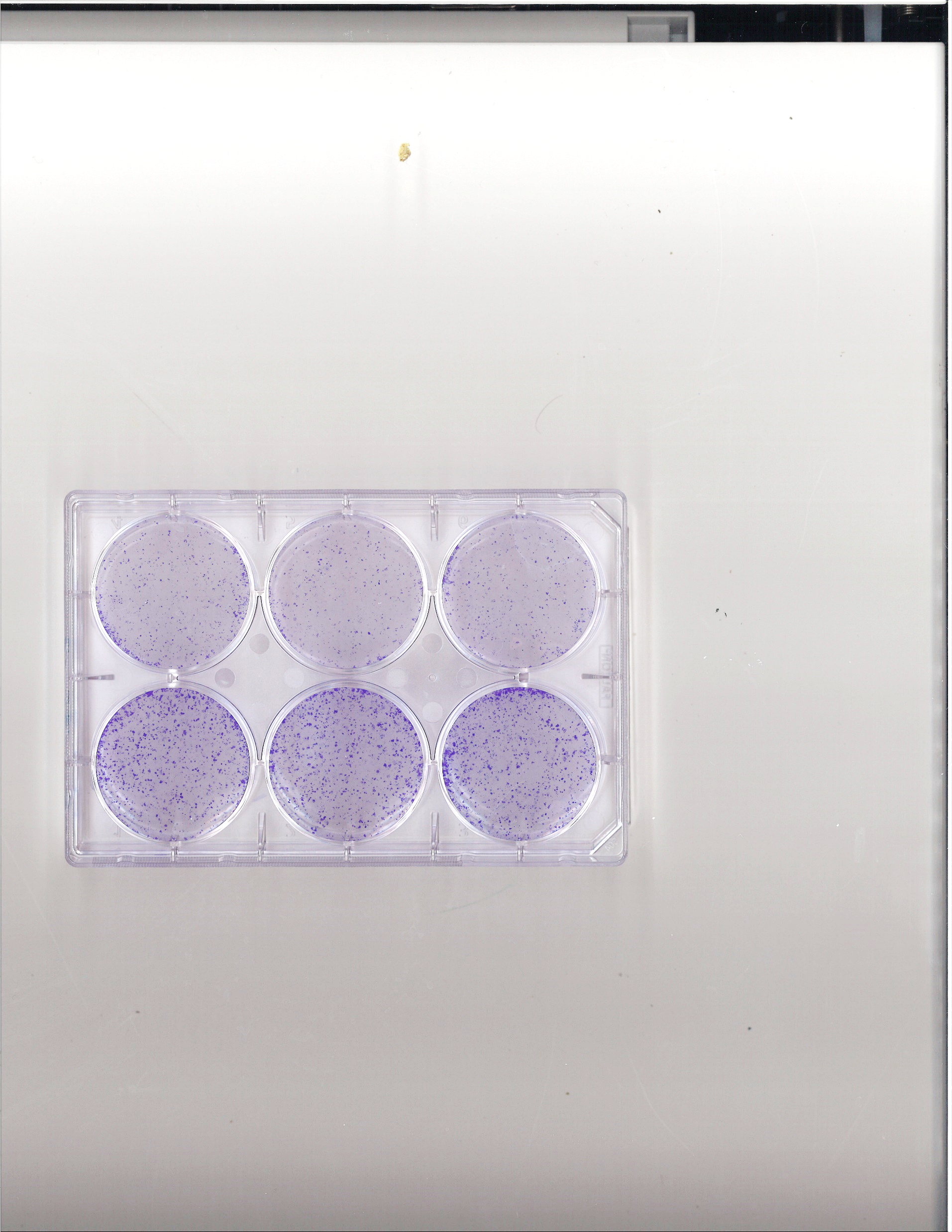

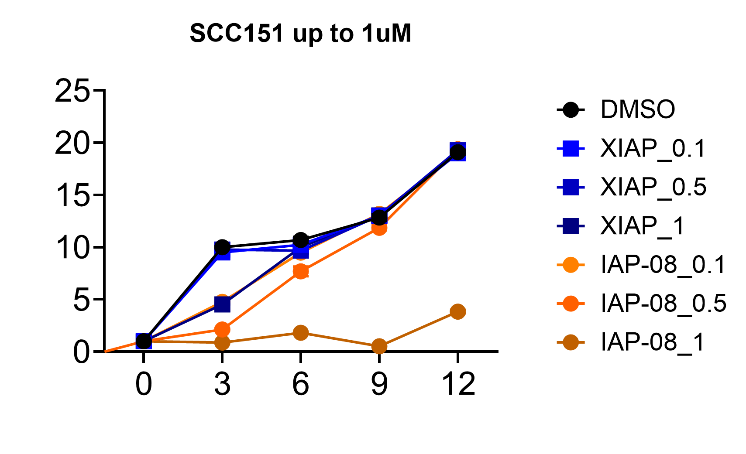

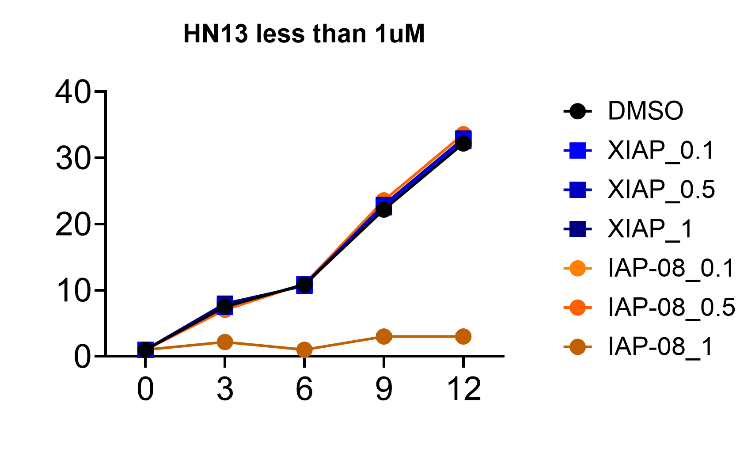
**(B)**

Days

Relative absorption compared to DMSO

**HN-SCC-151**
